# Lipid droplets protect human β cells from lipotoxic-induced stress and cell identity changes

**DOI:** 10.1101/2021.06.19.449124

**Authors:** Xin Tong, Roland Stein

**Affiliations:** Department of Molecular Physiology and Biophysics, Vanderbilt University, Nashville, TN

## Abstract

Free fatty acids (FFAs) are often stored in lipid droplet (LD) depots for eventual metabolic and/or synthetic use in many cell types, such a muscle, liver, and fat. In pancreatic islets, overt LD accumulation was detected in humans but not mice. LD buildup in islets was principally observed after roughly 11 years of age, increasing throughout adulthood under physiologic conditions, and also enriched in type 2 diabetes. To obtain insight into the role of LDs in human islet β cell function, the levels of a key LD structural protein, perilipin2 (PLIN2), were manipulated by lentiviral-mediated knock-down (KD) or over-expression (OE) in EndoCβH2-Cre cells, a human cell line with adult islet β-like properties. Glucose stimulated insulin secretion was blunted in PLIN2KD cells and improved in PLIN2OE cells. An unbiased transcriptomic analysis revealed that limiting LD formation induced effectors of endoplasmic reticulum (ER) stress that compromised the expression of critical β cell function and identity genes. These changes were aggravated by exogenous treatment with FFAs toxic to islet β cells, and essentially reversed by PLIN2OE or using the ER stress inhibitor, tauroursodeoxycholic acid. These results strongly suggest that LDs are essential for adult human islet β cell activity by preserving FFA homeostasis.

## Introduction

LDs are cellular organelles that typically play a key metabolic role by serving as a reservoir for cholesterol, acyl-glycerol and phospholipids used (for example) in signaling, energy homeostasis and membrane maintenance (1). While the exposure of pancreatic islet α and β cells to excess lipids and glucose ultimately results in their dysfunction and type 2 diabetes (T2D (2,3)), it is very difficult to detect LDs in rodent islets even under obese, pathophysiological conditions where accumulation is evident in well-recognized, peripheral LD-storage cells in liver, muscle and adipose tissue (4,5). Importantly, recent results suggest that LDs do accumulate in human islet α and β cells (5), which also differ with rodents in respect to islet cell composition, transcription factor (TF) expression, islet architecture, and glucose-stimulated insulin secretion (GSIS) (6–9).

Notably, LDs were found in human islet cells transplanted into immunocompromised mice raised on a normal or high fat diet, but not in similarly treated mouse islet cells (10). Strikingly, the compensatory mechanisms activated in response to the high fat diet-induced insulin resistant state were just observed in transplanted mouse islets (e.g., elevated β cell expansion and GSIS), while the non-responsive human islets accumulated LDs and the islet amyloid protein, the latter a hallmark of the T2D dysfunctional islet. The implication that LDs could impact islet cell function was further supported upon demonstrating that LDs were in islet α and β cells within the intact human pancreas, with this most evident being in post-juvenile aged (>11 years) islet β cells (5). Physiologic LD buildup was also found in the GSIS-responsive human embryonic stem cell-derived β-like cells produced in culture and after transplantation into immunocompromised mice. In contrast, LDs were very difficult to detect in the intact rodent pancreas under normal, aged or pathophysiological conditions (5).

In addition, LDs were enriched in human T2D islet β cells, which possibly is caused by the reduction in T2D-associated autophagic flux (5,11). We and others have proposed that LDs serve to sequester toxic FFA produced under insulin resistant conditions in human islets and that limitations in buildup and/or storage capacity results in islet β cell dysfunction and T2D susceptibility (10). However, the functional significance of LDs in human β cells has not been delineated thoroughly, and to obtain such insight, we have influenced LD formation by reducing or over-producing the key LD structural protein PLIN2 in the adult human β-like cell line, EndoCβH2-Cre. Compromising LD production induced ER stress, reduced GSIS, and generated gene expression changes in cellular identity, which are all characteristics of T2D islet β cells (12–14). Furthermore, these lipotoxic-induced changes were ameliorated by either elevating LD storage capacity by over-expressing PLIN2 or by pharmacological treatment with an ER stress inhibitor, tauroursodeoxycholic acid (TUD). These results strongly imply that LDs serve as a positive effector of adult human islet β cell activity.

## Research Design and Methods

### Human EndoCβH2-Cre cells

EndoCβH2 cells were propagated in Dulbecco’s Modified Eagle Medium (Gibco, Thermo Fisher) in presence of 5.6mM glucose, 2% BSA (Serologicals Proteins Inc., Kankakee, IL), 100 μU/ml penicillin, 100 μg/ml streptomycin, 50 μM 2-mercaptoethanol, 10 mM nicotinamide, 5 μg/ml transferrin and 6.7 ng/ml sodium selenite (Sigma-Aldrich) as described previously (15,16). Lentiviral vectors encoding short hairpin RNAs that were either Scrambled (i.e., Sham) or to PLIN2 (shPLIN2) were constructed by VectorBuilder (Chicago, IL) as was the PLIN2 protein over-expressing cassette, with each containing a puromycin selection marker. The Cre expressing lentivirus was made using the pTRIP ΔU3 CMV-nlsCre vector (17). Lentiviral particles were produced in HEK 293T cells as described earlier (18). Viral particles were isolated from the supernatants by either ultracentrifugation (18) or using the PEG-it Virus precipitation solution (System Biosciences, Mountain View, CA). The resultant pellets were resuspended in PBS or DMEM and aliquoted samples stored at -80°C. Viral particle amount was quantified using the Lenti-X™ p24 Rapid Titer kit (TAKARA Bio, Mountain View, CA). EndoCβH2 cells were infected for 16 hours with ∼50ng of shScramble, shPLIN2 or PLIN2-OE viral particles/million cells followed by puromycin selection; after that, all cells received the same viral dose of Cre expressing lentivirus for 18 days (16). The 10 mM stock solution of palmitic acid (C16:0; Sigma-Aldrich) and erucic acid (C22:1 Sigma-Aldrich) was freshly prepared by dissolving in 90% ethanol followed by fatty acid free bovine serum albumin (BSA) conjugation (Equitech-Bio, Inc.) in 0.01M NaOH solution at 55°C for 1 hour, with a working concentration of 500µM for both. The 100µM TUD (tauroursodeoxycholic acid, Sigma-Aldrich) and N-acetyl-L-cysteine (NAC; Sigma-Aldrich) stocks were made in DMSO.

#### Immunofluorescence analysis and LD quantification

EndoCβH2-Cre cells cultured on chamber slides were treated at room temperature for 12 mins with 4% paraformaldehyde-PBS, 8 mins with 0.5% Triton-PBS, 30 mins with 0.5% BSA-PBS followed by a 4°C overnight incubation with one of the following primary antibodies: insulin (guinea pig; 1:500; Dako, Santa Clara, CA), glucagon (mouse; 1:400; Sigma-Aldrich, St. Louis, MO), somatostatin (goat, 1:400, Santa Cruz Biotechnology, Santa Cruz, CA), FEV (rabbit, 1:400, Thermo Fisher), MAFA (rabbit, 1:500, Novus (NBP1-00121) and NKX2.2 (goat, 1:400, Santa Cruz). Species-matched antibodies conjugated with the Cy2, Cy3 or Cy5 fluorophores were used for secondary detection (1:1000; Jackson ImmunoResearch, West Grove, PA). BODIPY 493/503 (5 μM in PBS, Thermo Fisher Scientific, Waltham, MA) was used to detect neutral lipid enriched LDs by incubating at room temperature for 30 mins following the secondary antibody treatment. Images were acquired on a Zeiss Axio Imager M2 wide-field microscope with Apotome. Quantification of the LD level was calculated as the BODIPY 493/503 area divided by the DAPI^+^ nuclear cell number using ImageJ software. Normalization was to the Sham control; at least five distinct areas of the slide from several independently generated sample sets were quantified per condition.

#### RNA isolation, reverse transcription, and RT-PCR

Total RNA was collected from EndoCβH2-Cre cells using the Trizol reagent (Life Technologies) as described by the manufacturer. The iScript cDNA synthesis kit (Bio-Rad Laboratories, Inc.) was used for cDNA synthesis. Quantitative real time (RT)-PCR reactions were performed with the primers described in **Supp. Table 1** on a LightCycler 480 II instrument (Roche), and analyzed by the ΔΔCT method. Significance was calculated by comparing the ΔCT values.

#### Bulk RNA sequencing (RNA-seq) analysis

The RNeasy mini plus kit (QIAGEN) was used to isolate total RNA from treated EndoCβH2-Cre cells (n=3) and RNA quality control analyzed on an Agilent 2100 Bioanalyzer. Only samples with an RNA Integrity Number >8.0 were used for library preparation. cDNA libraries were constructed and paired-end sequencing performed on an Illumina NovaSeq6000 (150 nucleotide reads). The generated FASTQ files were processed and interpreted using the Genialis visual informatics platform (https://www.genialis.com) with previously described process (15). Sequence quality checks were determined using raw and trimmed reads with FastQC (http://www.bioinformatics.babraham.ac.uk/projects/fastqc), and Trimmomatic was utilized to trim adapters and filter out poor-quality reads. Trimmed reads were then mapped to the University of California, Santa Cruz, mm10 reference genome using the HISAT2 aligner. Gene expression levels were quantified with HTSeq-count and differential gene expression analyses performed with DESeq2. All detectable genes were included in the pathway analysis (KEGG and GO term) as part of the Enrichr bioinformatic analysis platform (https://maayanlab.cloud/Enrichr/) to prevent bias. Some of the heatmap selected genes were manually curated based on GESA (https://www.gsea-msigdb.org/gsea/index.jsp) and published gene or RNAseq datasets (19–21). However, poorly expressed genes, which had average expression count in all samples of TPM (Transcripts Per Million) < 5, were filtered out of the gene lists and heatmaps.

### Static glucose regulated insulin secretion

Static insulin secretion was assessed as described earlier in EndoCβH2-Cre cells (22), which involved a 1-hour incubation at 37°C in 2.5mM glucose secretion assay buffer, and then with either freshly prepared 2.8 or 16.7 mmol/L glucose secretion assay buffer for another hour. The outcome was presented as the secreted insulin (Lumit Insulin Immunoassay, Promega and Human insulin ELISA, Crystal Chem) relative to the total insulin content. The secreted insulin data was normalized to the basal secretion in the Sham at 2.5mM glucose. Insulin content was presented as the concentration of insulin normalized to total DNA content (Quant-iT™ PicoGreen™, Invitrogen) in each condition (ng/ng DNA) relative to the Sham. At least 3 independently generated sets of samples were analyzed under each condition.

### Statistical Analysis

Significance was determined using the two-tailed Student *t* test. Data are presented as the mean ± SEM. A threshold of *P* < 0.05 was considered significant.

### Data and Resource Sharing and Availability

The datasets generated during and/or analyzed during the current study are available from the corresponding author upon reasonable request. The bulkRNAseq full dataset will be uploaded to GEO/SRA depository upon manuscript acceptance.

## Results

### PLIN2 regulates LD levels in EndoCβH2-Cre β cells

To obtain insight into the functional role of LDs in human β cells, the mRNA levels of core structural perilipins was first determined in proliferating EndoCβH2 and non-proliferating EndoCβH2-Cre cells. Lentiviral-mediated Cre treatment and its expression in EndoCβH2 cells removes the endogenous floxed SV40 virus Tag transforming protein to prevent cell proliferation and impart functional maturation (**Fig. 1A**). The resulting EndoCβH2-Cre cell line produces high insulin levels and adult islet β-like GSIS-responsivity (16). Compared to other family members, PLIN2 was the principally produced in human and mouse islets (**Fig. 1B**; (5,23)). Lentiviral knock-down of PLIN2 (PLIN2KD) treatment reduced PLIN2 mRNA by roughly 70%, while PLIN2 overexpression (PLIN2OE) induced by 50% (**Fig. 1C**). PLIN2KD or PLIN2OE had no significant effect on cell proliferation, cell death, or cell morphology in EndoCβH2 or EndoCβH2-Cre cells (data not shown). PLIN2 mRNA levels were elevated upon inducing GSIS in EndoCβH2-Cre cells (**Fig. 1B**), a property also observed in rodent β cell lines (24).

**Figure 1.**
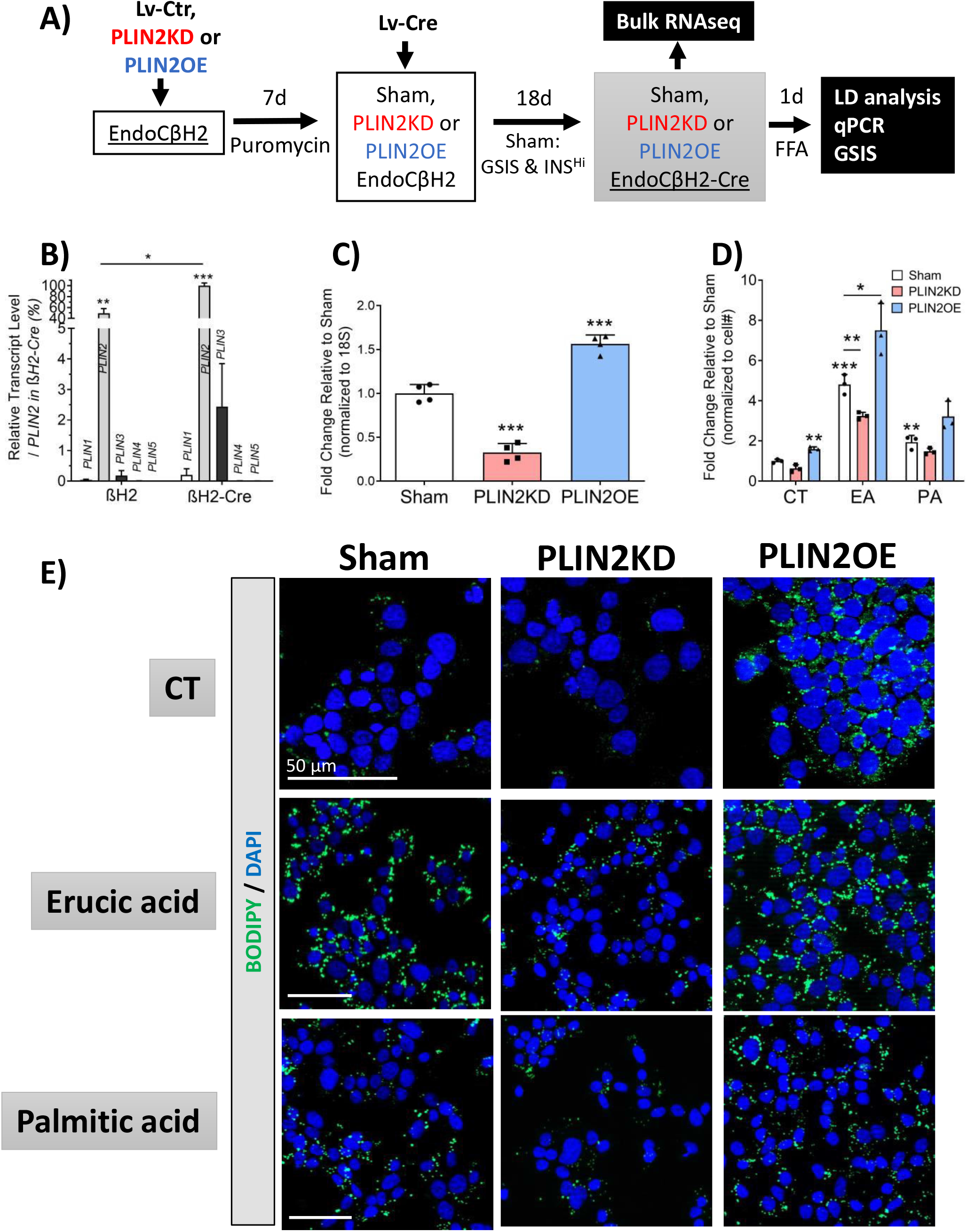
PLIN2 influences LD accumulation in human EndoCβH2-Cre β cells. (**A**) Experimental workflow and depiction of how maturation changes upon Cre mediated SV40 Tag removal. (**B**) *PLIN* mRNA levels before and after Cre treatment. Both *PLIN2* and *PLIN3* levels increased significantly. (**C**) *PLIN2* mRNA levels decreased upon PLIN2KD and increased in PLIN2OE. (**D**) Quantitation of the change in LD numbers between Sham and PLIN2KD or PLIN2OE cells incubated without (CT) or with 500μM of Erucic acid (EA) or Palmitic acid (PA) for 24 hours. (**E**) Representative images of (**D**) effects. Notably, FFA induced LD accumulation appears to positively correlate with PLIN2 levels, which was statistically significant in the EA treated group. All error bars indicate SD, n=3-4, * P<0.05, ** P<0.01, *** P<0.05 vs βH2 PLIN1 (B), Sham (C) or CT (D) unless specified.

The impact of PLIN2KD and PLIN2OE on LD accumulation was evaluated in EndoCβH2-Cre cells treated with lipotoxic levels of two FFA, palmitic acid (PA, C16:0) and erucic acid (EA, C22:1). PA induces a much milder cell stress response in relation to EA in EndoCβH1 cells (25), a closely related human β cell line. As expected from studies in other contexts (26), BODIPY 493/503 detected LD accumulation was decreased significantly following PLIN2KD treatment and enhanced by PLIN2OE (**Fig. 1D-E**). Because of the higher toxicity profile of EA than PA in EndoCβH1 (25) and EndoCβH2-Cre (see below) cells, EA was the FFA of choice to impart lipotoxicity in our EndoCβH2-Cre experiments.

### PLIN2 levels regulates insulin secretion

PLIN2KD prevented GSIS in EndoCβH2-Cre cells (**Fig. 2A**), while PLIN2OE enhanced in comparison to the Sham. The loss of GSIS was not due to reduced insulin mRNA production in PLIN2KD nor a difference in insulin protein levels between PLIN2KD and PLIN2OE cells (**Fig. 2B, C**).

**Figure 2.**
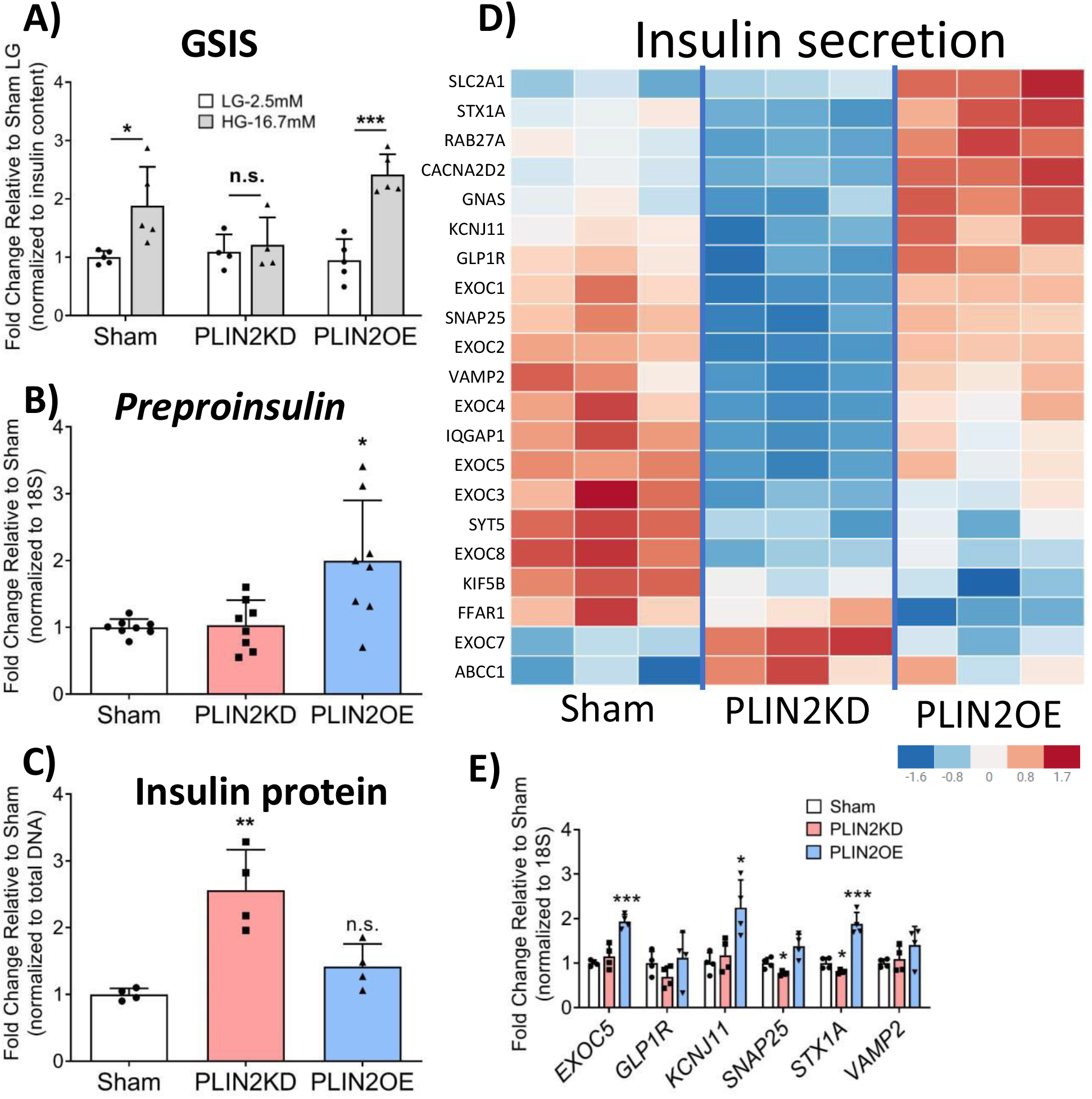
PLIN2/LDs regulate GSIS in EndoβCH2-Cre cells. **(A)** GSIS was blocked in PLIN2KD cells compared to Sham, whereas this was enhanced in PLIN2OE. Levels were first normalized to DNA content and then to Sham at 2.5 mM glucose. **(B)** *Preproinsulin* mRNA trended higher in both PLIN2OE and PLIN2KD cells, although (**C**) insulin protein levels were only elevated in PLIN2KD. Normalization was to (**B**) 18S or (**C**) DNA content before Sham. (**D**) The heatmap shows that insulin secretion-related gene expression was generally decreased in PLIN2KD cells, while elevated in PLIN2OE cell. The color keys represent the row-wise Z-score. FDR<0.05 (**E**) qRT-PCR analysis of selected secretory gene changes. All error bars indicate SD, n=3-8, * P<0.05, ** P<0.01, *** P<0.001 vs corresponding LG (A), or Sham (B-E).

Bulk RNA-seq analysis performed on Sham, PLIN2KD and PLIN2OE EndoCβH2-Cre cells revealed that a large number of positive effectors of insulin secretion were impacted upon PLIN2/LD manipulation. Not surprisingly, expression of these were reduced in PLIN2KD and boosted by PLIN2OE (**Fig. 2D**). These differences were also verified upon qPCR analysis of various regulatory candidates (**Fig. 2E**), including *EXOC5* (27) that is involved in vesicle docking and fusion, *KCNJ11* (28) encoding the ATP-sensitive inward rectifier potassium channel 11, *SNAP 25* (29) of the SNARE complex, and *STX1A* (14) which is important in vesicle membrane docking. Taken together, our data suggest that LD accumulation affects the insulin secretory machinery and GSIS in human β cells.

### PLIN2KD induces ER stress and compromises Ca^2+^ homeostasis

The RNA-seq results revealed that PLIN2KD in EndoCβH2-Cre cells had 1479 upregulated and 493 downregulated genes compared to Sham, while 237 were elevated and 48 decreased in PLIN2OE cells (**Supp. Tables 2-3** provides the full list of differentially expressed genes (i.e., DEGs)). We next focused on obtaining a more thorough understanding why reducing LD accumulation eliminated GSIS (**Fig. 2A**). KEGG and gene ontology (GO term) analysis revealed that the PLIN2KD cells manifested gene expression profiles characteristic of proinflammatory cytokines treatment and glucolipotoxicity (**Supp. Table 4** (19)). For example, the positively affected “AGE-RAGE signaling in diabetic complications” and “inflammation-like” pathways mediate proinflammatory signaling and ER stress in other systems (30), while β cell dysfunction would also have resulted upon down-regulating proteins essential to pathways associated with “insulin secretion”, “cAMP signaling”, “maturity onset diabetes of the young”, “voltage-gated cation channel activity”, “gap junction channel activity” and “cytoskeleton molecule binding.”

In contrast, while the PLIN2OE cells more closely resembled the Sham, differentially expressed genes defined by KEGG analysis in elevating “gap junction activity” and “microtubule motor activity” would be predicted to enhance β cell activity, as would reducing of “TNF signaling” and “cytokine/chemokine activity” pathway gene encoded proteins (**Supp. Table 5**). Importantly, these properties are opposite to PLIN2KD cells (**Supp. Table 4**). In addition, genes important in “transforming growth factor β (TGFβ) receptor signaling”, a pathway negatively regulating β cell function and identity (31), were only elevated after PLIN2KD treatment (**Supp. Fig. 1**).

To counteract ER stress conditions that can result in β cell death, 3 arms of a network of signaling pathways may be activated to restore ER homeostasis: 1) ATF6, 2) IRE1-XBP1, and 3) PERK-eIF2α which collectively orchestrate the selective transcription and translation of ER chaperones, elimination of misfolded proteins by ER-associated degradation and autophagy, and reduce incoming protein load by decreasing global transcription and translation (32). Our RNA-seq and qPCR results indicate that the IRE-XBP1 pathway was primarily activated in PLIN2KD EndoCβH2-Cre cells, and not the ATF4/6 (i.e., CHOP) or PERK pathways (**Fig. 3A, C**). In addition, many ER regulators essential to Ca^2+^ homeostasis and insulin secretion in islet β cells were mis-regulated in PLIN2KD cells (e.g. **Fig. 3B**), an effect that would contribute to blunting the GSIS response (33). Significantly, many genes involved in these pathways were amongst the most upregulated in PLIN2KD cells (**Supp. Table 6**; e.g., ERN1 (IRE1), DDIT4 (CHOP), TGFβ2 and BMP2). On the contrary, a key TGFβ signaling regulatory gene was down regulated genes in PLIN2OE cells (**Supp. Table 7**; SMAD7).

**Figure 3.**
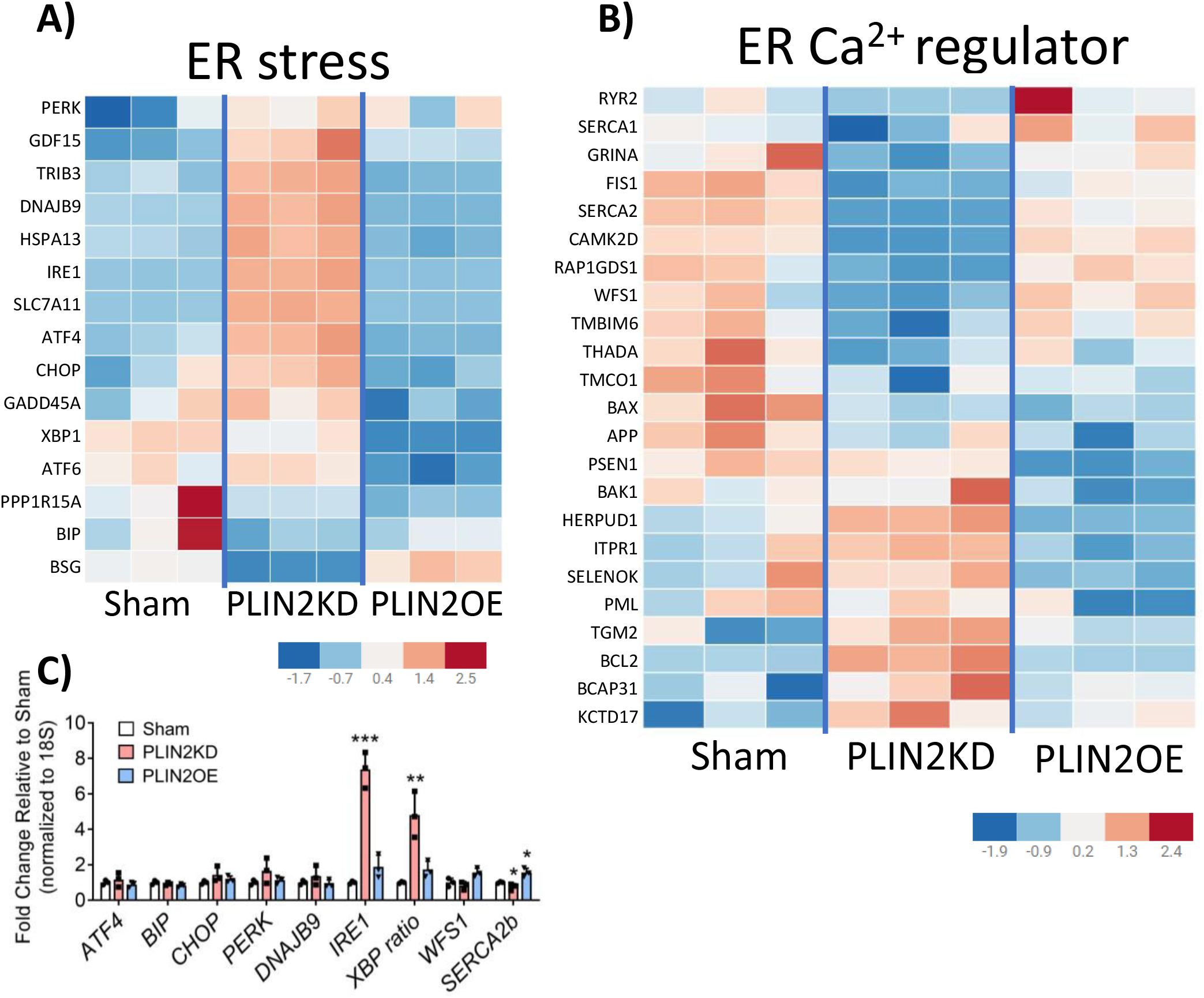
PLIN2KD induces ER stress. Heatmaps showing how PLIN2KD influences the expression of genes involved in (**A**) ER stress and (**B**) ER Ca^2+^ homeostasis. The color keys represent the row-wise Z-score. (**C**) qRT-PCR analysis of selected ER stress and ER Ca^2+^ regulators from panels (**A**) and (**B**). All error bars indicate SD, n=3, * P<0.05, ** P<0.01, *** P<0.001 vs Sham.

### PLIN2KD influences β cell identity

Cellular stress is believed to be a driver of T2D islet cell dysfunction and β cell identity (12,13), which is manifested by decreased expression of essential TF genes, misexpression of non-islet β hormones, induced production of gene products incompatible with adult function (i.e., termed the disallowed genes (21)), and the synthesis of progenitor/dedifferentiation markers (34). Strikingly, these same characteristics were witnessed upon simply reducing LD accumulation in PLIN2KD EndoCβH2-Cre cells (**Fig. 4A, B**), as exhibited, for example, by increased non-β cell POMC, SST, and NPY hormone levels, elevated islet progenitor FEV, FOS, and SOX9 TF marker levels, misexpression of disallowed genes, and reduced expression of islet-enriched MAFA, NKX2.2, PDX1, and NKX6.1 TF genes. However, none of changes were observed in PLIN2OE cells (**Fig. 4B, data not shown**). Interestingly, immunostaining revealed that only a fraction of PLIN2KD EndoCβH2-Cre cells produced the α cell glucagon and δ cell somatostatin hormones (**Fig. 5A**) or the FEV protein (35) that is normally principally produced in Neurogenin 3^+^ TF islet progenitor cell population (**Fig. 5B**). In contrast, islet cell-enriched MAFA and NKX2.2 TF levels appeared to be decreased throughout the β cell population (**Fig. 5C**). As expected, PLIN2OE prevented these changes found upon PLIN2KD (**Fig. 5**). Collectively, these results imply that LD accumulation regulates both the cell identity and function of human islet β cells.

**Figure 4.**
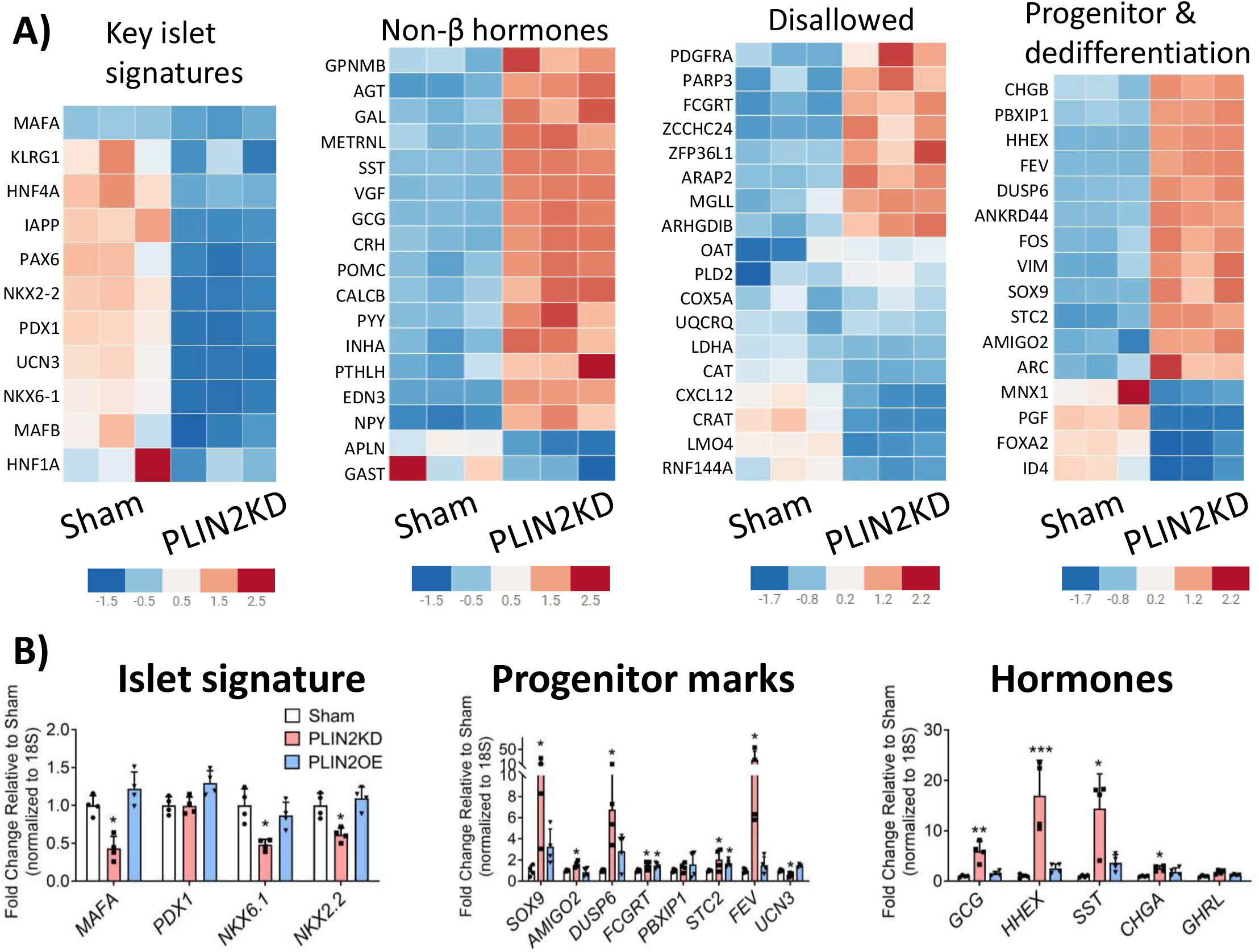
PLIN2KD influences β cell identity. **(A)** Heatmaps illustrating the changes in PLIN2KD cells in expression of key β cell signature genes, non-β cell hormone genes, disallowed genes and genes enriched in embryonic progenitors and dedifferentiated β cells. The color keys represent the row-wise Z-score. (**B**) qRT-PCR results of candidates in Sham, PLIN2KD, and PLIN2OE cells. All error bars indicate SD, n=3-4, * P<0.05, ** P<0.01, *** P<0.001 vs Sham.

**Figure 5.**
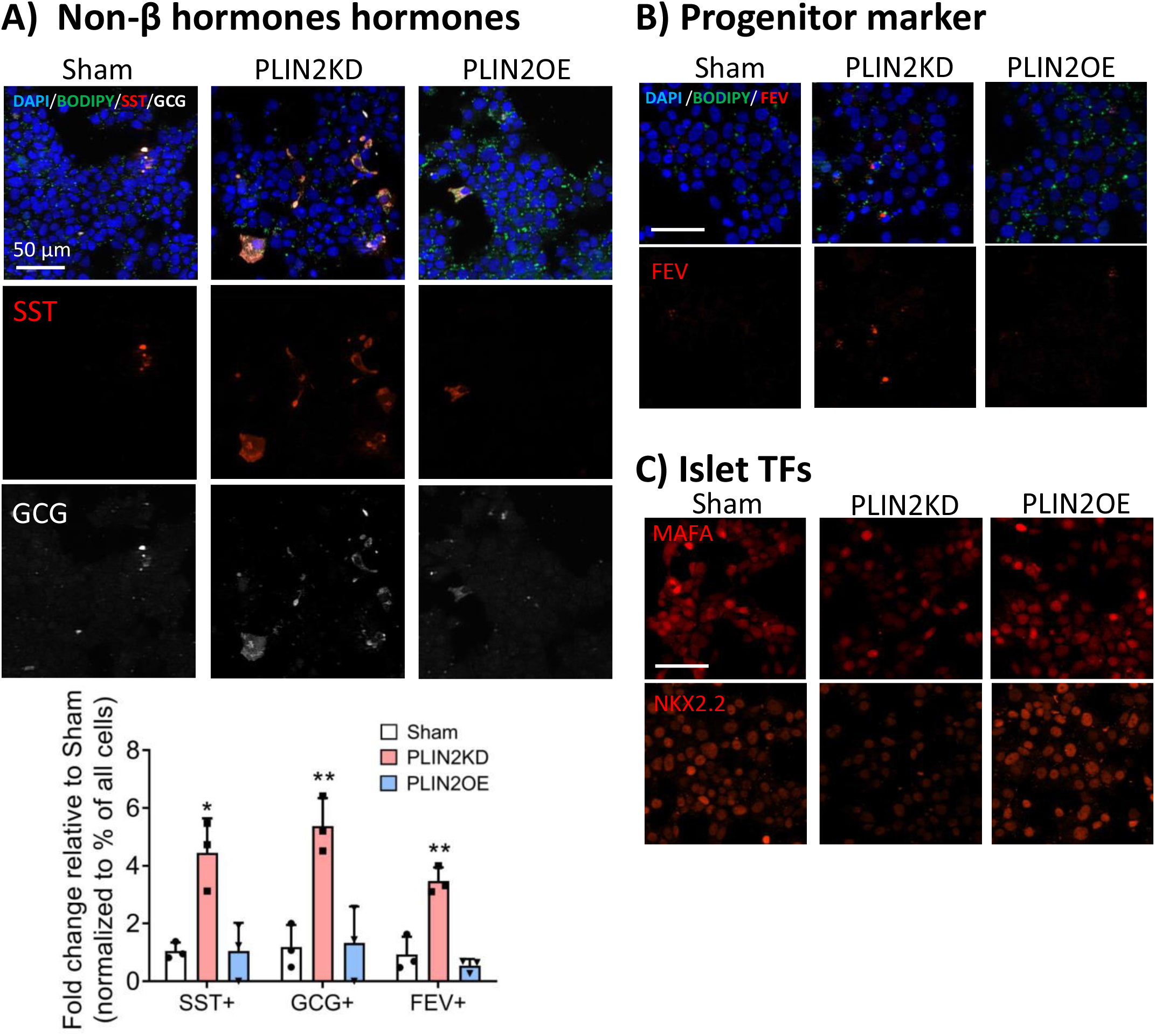
Cellular distribution of proteins linked to β cell identity in PLIN2KD cells. Immunofluorescent staining illustrates that (**A**) somatostatin (SST, red), glucagon (GCG, white), with quantitation of the change in SST or GCG positive cell percentages between Sham and PLIN2KD or PLIN2OE cells, all error bars indicate SD, n=3, * P<0.05, ** P<0.01 vs Sham; (**B**) FEV (red) protein production is only found in a small fraction of PLIN2KD cells. In contrast, and (**C**) MAFA (red, top panel) and NKX2.2 (red) protein levels were lower in most PLIN2KD cells. The remaining MAFA^+^ and NKX2.2^+^ cells in the PLIN2KD population probably represent uninfected cells. DAPI (blue) and BODIPY (green) counterstaining. Scale bar=50 µm.

### ER stress is a dominant effector of PLIN2KD induced dysfunction

Given the pronounced elevation in multiple markers of ER stress in PLIN2KD EndoCβH2-Cre cells, a bile acid analog commonly used to inhibit this response, tauroursodeoxycholic acid (TUD), was tested in this context. GSIS (**Fig. 6A**) and insulin levels (**Fig. 6B**) were rescued by TUD treatment, with levels now indistinguishable from Sham. As anticipated from these results, the expression of a representative panel of genes that affected insulin secretion, ER stress, and islet TF returned to Sham-like levels after TUD treatment of PLIN2KD cells (**Fig. 6C**). In contrast, a general oxidative stress inhibitor, N-acetyl cysteine, appears to have further compromised PLIN2KD cells (**Supp. Fig. 2**). Notably, those genes influenced to a lesser extent by TUD were produced in a small subfraction of the β cell population (e.g., FEV (**Fig. 5B**), and likely the HHEX (36) TF of the somatostatin gene). Moreover, the toxic effect of EA treatment on GSIS and ER stress-responsive targets genes was essential completely blunted by PLIN2OE (**Fig. 6A-C**). EA also induced a much more robust stress response than PA in EndoCβH2-Cre cells (**Supp. Fig 3**), as originally described in EndoCβH1 cells (25). In total, our data strongly indicate that LDs are beneficial to human islet β cells, with our mechanistic analysis implying their ability to sequester toxic FFA prevents ER stress, loss of cell activity, changes in cell identity, and decreased cell health (**Fig. 7**).

**Figure 6.**
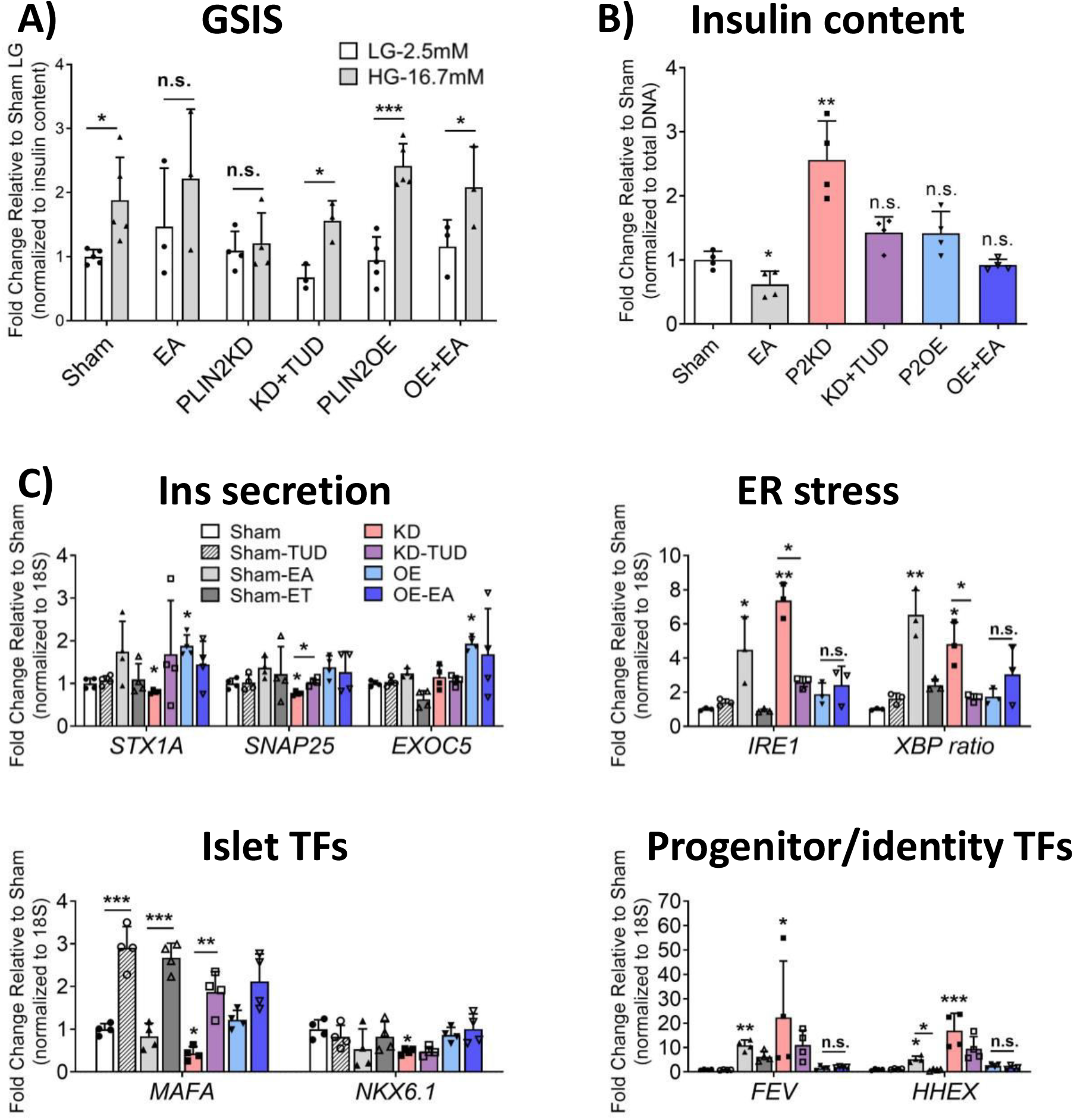
TUD rescues PLIN2KD induced dysfunction, while PLIN2OE prevents EA toxicity. Sham, PLIN2KD and PLIN2OE cells were treated with TUD (100 µM) and/or EA (500 µM) for 24 hours. The toxic effects of PLIN2KD and EA on (**A**) GSIS and (**B**) insulin levels were rescued by TUD and PLIN2OE treatment, respectively. (**C**) Expression of the majority of candidate regulatory genes were corrected to Sham by TUD or PLIN2OE. All error bars indicate SD, n=3-4, * P<0.05, ** P<0.01, *** P<0.001 vs corresponding LG (A), or Sham (B-C) unless specified.

**Fig. 7.**
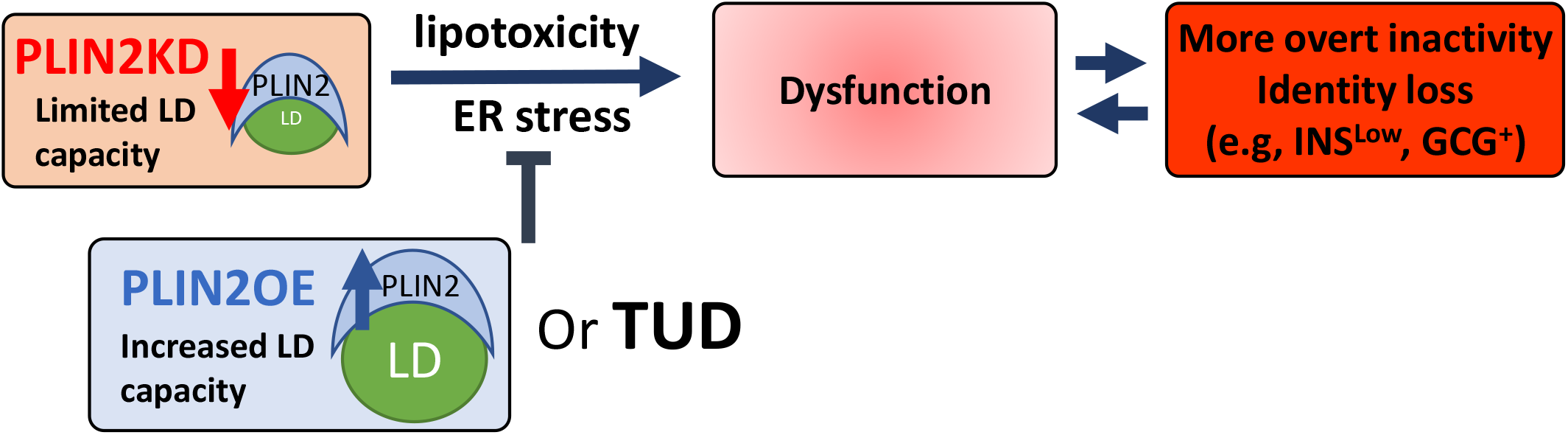
Model illustrating the proposed importance of FFA storage in LDs to human β cell health.

## Discussion

The functional role of LDs in cells has been studied extensively under both healthy and diseased conditions in many contexts, including in muscle, liver and adipose tissues (1). The general consensus has been that LDs normally serve as a storage depot for neutral lipids and cholesterol for anabolic and catabolic needs, with lipotoxicity produced under pathological conditions due (at least in part) to limitations in FFA storage and/or breakdown in LDs (4). In contrast, studies in mice suggest that LD accumulation is detrimental to islet β cell health, since mice totally lacking the LD PLIN2 structural protein or upon knockdown of PLIN2 in rodent β cells have improved autophagic flux, reduced ER stress, and decreased β cell apoptosis (24,37). However, we and others have found that LDs are very difficult to find in the rodent pancreas, whereas they are readily detected in adult humans and are even enriched in T2D islets (5,11). Because human and rodent islets differ substantially in architecture, cell composition, proliferative capacity, islet amyloid formation, and antioxidant enzyme levels (7,10,38,39), the EndoCβH2-Cre cell line was used to examine the influence of LDs in human β cells, a well-established model system whose molecular composition and GSIS properties are quite similar to adults (16,40). Our data strongly suggest that LDs function, in part, to insulate human β cells from noxious FFA exposure until a threshold capacity for LDs is reached, after which these cells are vulnerable to lipotoxicity induced ER stress, dysfunction, loss of identity and overall compromised cell health (**Fig. 7**). Consequently, differences in LD formation and/or degradation could be a contributing factor to T2D islet β cell dysfunction.

### Limiting LD accumulation induces lipotoxic-like ER stress and dysfunction

Manipulating PLIN2 levels significantly altered LD accumulation levels in EndoCβH2-Cre cells (**Fig. 1D**,**E**), a result consistent with many studies showing that LD formation is dependent on perilipin structural protein levels (23). This resulted in the reduction in GSIS in PLIN2KD cells, and improvement in PLIN2OE (**Fig. 2A**). RNA-seq analysis suggested that limiting LD formation induced IRE1 activated ER stress, which reduced production of a variety effectors of insulin secretion. This not only included the expression of mRNA encoding proteins directly involved in this process (**Fig. 2D, E**), but effectors mediating critical GSIS-dependent signaling pathways (e.g., ER Ca^2+^ (**Fig. 3B**) and TGFβ (**Supp Fig. 1**)). These results are not consistent with the current published work in rodents (24,37). However, they do partly agree with preprint findings in INS-1 cells (41). It is now unclear if this reflects species differences or the methodological approach. Although this recent work had only limited molecular detail in how compromising LD formation negatively impacted β cell function, their physiological results showed that mitochondrial health and activity was negatively impacted in both rodent β cells and human islets (41). Notably, our RNA data also suggests that the integrity of PLIN2KD mitochondria are compromised (**Supp Fig. 4**).

Our results suggest that ER stress facilitates lipotoxicity in human β cells with limited LD storage capacity (**Figs. 3 and 6**). Indeed, cellular stress appears to be a mediator of this condition in many tissues (42). For example, LDs were generated in mouse embryonic fibroblasts to protect against mitochondrial dysfunction induced upon FFA release from autophagic degradation of membranous organelles, a diacylglycerol acyltransferase 1 (DGAT1)-dependent process. This enzyme, which catalyzes the terminal step in triacylglycerol synthesis, is also necessary in preventing ER stress during adipose tissue inflammation (43). In addition, reducing LD formation or storage capacity (e.g., by overexpression of the lipolysis enzyme adipose triglyceride lipase), results in the generation of proinflammatory and ER stress markers in liver and cardiomyocytes (44,45), while improving LD levels by perilipin protein over-expression lowered their levels in skeletal muscle (46). The overall protective role of LDs in preventing stress was also observed in many cancer cells during metastasis (47).

### LD maintenance is essential for sustaining β cell identity

The loss of islet β cell identity produced by glucolipotoxic induced ER stress (48) is a characteristic of T2D islets (12,13), as manifested by fewer insulin^+^ cells, limited islet enriched TF gene expression, induction of disallowed gene production, and gain in non-islet β cell hormone synthesis. Significantly, these same properties were observed in PLIN2KD EndoCβH2-Cre cells (**Fig. 4**). Moreover, changing perilipin levels also influenced adipose cell identity, with PLIN2 deletion or PLIN1 over-expression promoting the browning of white adipose (49).

Here we observed that MAFA and NKX2.2 protein levels were dysregulated throughout the PLIN2KD cell population, while glucagon, somatostatin and FEV were only produced in a small fraction (**Fig. 5**). This may represent the temporal progression of T2D, with broad limitations in islet β cell function induced under prediabetic circumstances that become more severe and eventually result in a reduction in insulin production as well as the gain (for example) of non-β cell hormone and progenitor cell markers. However, we did not observe induction of the very early islet cell progenitor signatures first observed upon FoxO1 TF β cell deficiency in mice, like Nanog, Neurogenin 3, Oct4 and L-Myc (34). Notably, these markers are not a feature in all human T2D studies (50), which may reflect both the heterogeneity of the disease and/or methodological differences (51). Progenitor cell marker expression in PLIN2KD cells was represented by FEV, which is normally expressed at high levels in islet Neurogenin 3 ^+^ progenitors developmentally as well as later at lower levels in immature insulin^+^ cells and adult islet β cells (35). FEV levels are also elevated in T2D islets (52). It is presently unclear if *bona fide* markers of dedifferentiation, like Neurogenin 3, are simply not regulated by LD/lipotoxicity levels in human β cells or represent a limitation of the conditions and/or model used for experimentation.

### There are many similarities in the signaling pathways regulated in human islets and PLIN2KD cells

As hoped of the human PLIN2KD model, there were many differentially regulated genes shared with palmitate treated human islets (19). These included genes controlling ER stress, extracellular matrix and metabolic signaling pathways (**Supp Fig. 5**). As dyslipidemia is one of the highest T2D risk factors (53), it was also not surprising that there was overlap between PLIN2KD and T2D islet dysregulated genes (**Supp Fig. 6A**). In addition, overlap was found with T2D associated genes/pathways found in genome-wide association studies (**Supp. Table 2, 4** and **Supp Fig. 6B**), including a FFA desaturase that were found to maintain ω-6 and ω-3 polyunsaturated fatty acid level balance and their associated proinflammatory phenotype in liver (i.e., *FADS1*), a G protein-coupled receptor for medium- and long-chain unsaturated fatty acids protective against lipotoxicity-induced pancreatic β cell dysfunction (*GPR120*), and a master lipid TF regulator that plays a role in β cell function (*PPARγ*, (54)). In addition, a number of PLIN2KD DEGs (108 out of 1972) overlapped with genes critical to FFA uptake, lipid synthesis, breakdown and storage (**Supp Fig. 7**). This evidence supports a strong relationship between FFA and LD homeostasis in human β cell function and health. Furthermore, many amino acid metabolism-related genes altered in PLIN2KD cells are changed in individuals with T2D (55) (i.e., in the “Glycine, serine, threonine, cysteine and methionine metabolism” pathway (**Supp. Table 4**)).

While the positive impact of either TUD or PLIN2OE treatment on EndoCβH2-Cre cells on GSIS strongly indicates that ER stress is the primary driver of lipotoxicity in PLIN2KD cells (**Figs. 3 and 6**), it is still unclear precisely how limiting LD formation causes adult human islet β cell dysfunction. For example, does this result from FFA toxicity and/or the reduction in critical signaling molecule interactions mediated by the large number of proteins binding to this organelle (56)? Future studies examining whether acute changes in LD levels influence adult human islet β cells differently than EndoCβH2-Cre cells, and whether accumulation effects transplanted islet human islet β cell function *in vivo*, which are normally unable to mount a compensatory proliferation or secretion response to high fat diet induced stressors (10) are needed. Importantly, this study supports such effort by providing a linkage between LD homeostasis and human β cell integrity.

## Acknowledgements

This research was performed using resources and/or funding provided by NIH grants to R.S. (DK090570, DK050203, DK126482) and the Vanderbilt Diabetes Research and Training Center (DK20593). X.T. was supported by a JDRF Fellowship (3-PDF-2019-738-A-N). We also thank Drs. Raphaël Scharfmann and Phillippe Ravassard for generously providing EndoC-βH2 cells, Dr. Rotonya Carr for providing LD and PLIN2-related background knowledge and suggestions on evaluating PLIN2 manipulation, and Dr. Jeeyeon Cha for reading and editing the manuscript. R.S. is the guarantor of this work and, as such, had full access to all the data in the study and takes responsibility for the integrity of the data and the accuracy of the data analysis. All authors have no conflict of interest to declare.

## Author Contributions

X.T. and R.S. designed the experiments. X.T. executed and analyzed the experiments with input from R.S. The manuscript was written by X.T. and R.S. The guarantor of this work is R.S. and, as such, had full access to all data in the study and takes responsibility for the integrity of the data and the accuracy of the data analysis.

