## Supplemental Figures and legends for "Lipid droplets protect human β cells from lipotoxic-induced stress and cell identity changes"

#### Supplemental Figure and Legends

**Supplemental Figure 1. PLIN2KD significantly altered TGFβ/SMAD pathway genes.**

(**A**) Heat maps showing significantly elevated levels of TGFβ/SMAD pathway-related genes in PLIN2KD EndoCβH2-Cre β cells. The color keys represent the row-wise Z-score. (**B**) Notably, *GDF15* level appears to be PLIN2 dependent (directly extracted from bulk RNA-seq data), TPM: Transcripts Per Million. All error bars indicate SD, n=3, * P<0.05 vs Sham.

**Supplemental Figure 2. NAC cannot rescue EA induced effectors of ER stress.**

EndoCβH2-Cre cells (termed Sham) were treated with EA (500 μM, 24h) in the presence of absence of N-acetyl cysteine (NAC, 1mM). The expression of stress induced IRE1 and XBP-1 ratio (i.e., spliced/total) mRNA levels were analyzed by qPCR. All error bars indicate SD, n=3, * P<0.05, ** P<0.01, *** P<0.05 vs Sham

**Supplemental Figure 3. EA provoked a stronger stress gene response than PA.**

Sham cells were treated with PA (500 μM, 48 hours) or EA (500 μM, 24 hours) and then analyzed by qRT-PCR for FEV, IRE1 and XBP-1 ratio (i.e., spliced/total). All of these PLIN2KD and ER stress markers were more highly elevated in EA cells. All error bars indicate SD, n=3, * P<0.05, ** P<0.01, *** P<0.05 vs Sham.

**Supplemental Figure 4. PLIN2 manipulation induced many changes in the expression of gene important to mitochondria function and health.**

Heat maps showing changes in many different mitochondrial genes in PLIN2KD and PLIN2OE cells, including (**A**) electron transport chain and mitochondrial encoding, (**B**) fusion and fission, (**C**) ion balance and (**D**) mitophagy. The color keys represent the row-wise Z-score.

**Supplemental Figure 5. PLIN2KD DEGs partly overlapped with those altered in human islets under lipotoxic condition.**

Venn diagram and pathway analysis uncovered 149 out of 1972 DEGs in PLIN2KD cell overlapped with the 1196 DEGs found in human islets treated with palmitic acid (1). Those involved in β cell function and stress response pathways are highlighted in red.

**Supplemental Figure 6. PLIN2KD DEGs share genes dysregulated in T2D in human islets.**

(**A**) Venn diagram and heatmaps demonstrated 19 of 1972 DEGs in PLIN2KD overlapped with signature genes found T2D islet based on human GWAS datasets (2,3). (**B**) T2D signatures genes include two FFA handling genes, *FADS1* (4) and *GPR120* (5), which were down regulated in PLIN2KD. The color keys represent the row-wise Z-score.

**Supplemental Figure 7. Expression of many genes essential to lipid homeostasis are impacted by PLIN2KD.**

Venn diagram showed 104 out of 1972 PLIN2KD DEGs overlapped with the master list of 1108 major lipid handling and homeostasis genes (Gene Ontology online database: http://amigo.geneontology.org/amigo/landing (6)). GO term analysis highlighted in red depict some enriched processes, including effects on phospholipid metabolism, lipid transport/remodeling, and long-chain FA metabolism.

### Supplemental Fig. 1

A)

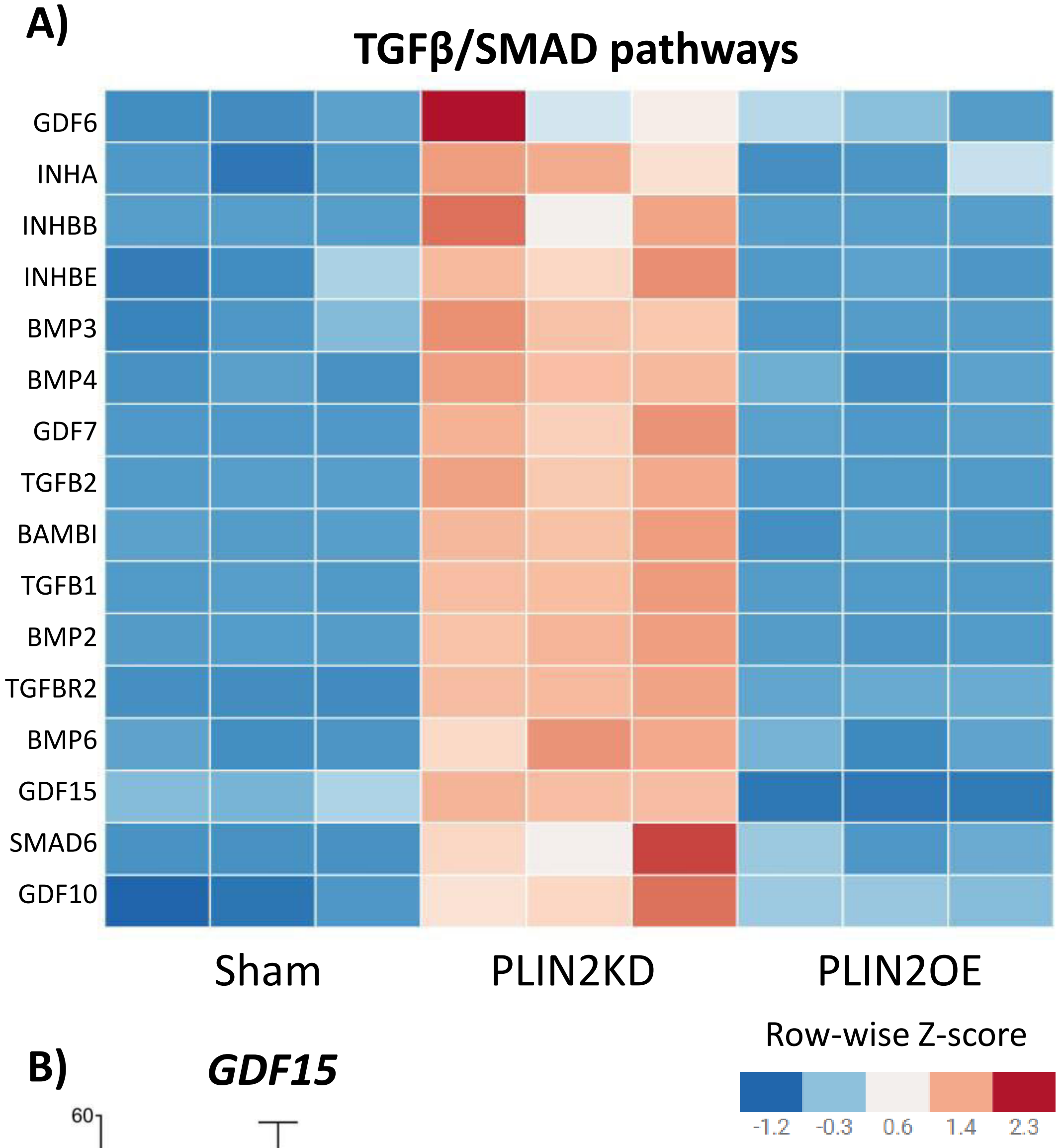


B)

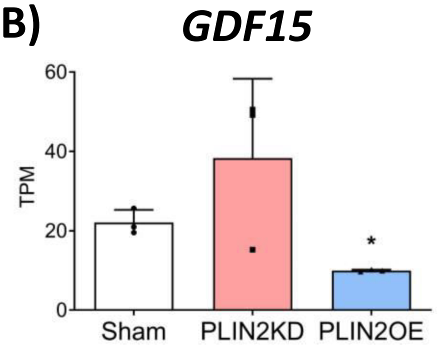

### Supplemental Fig. 2

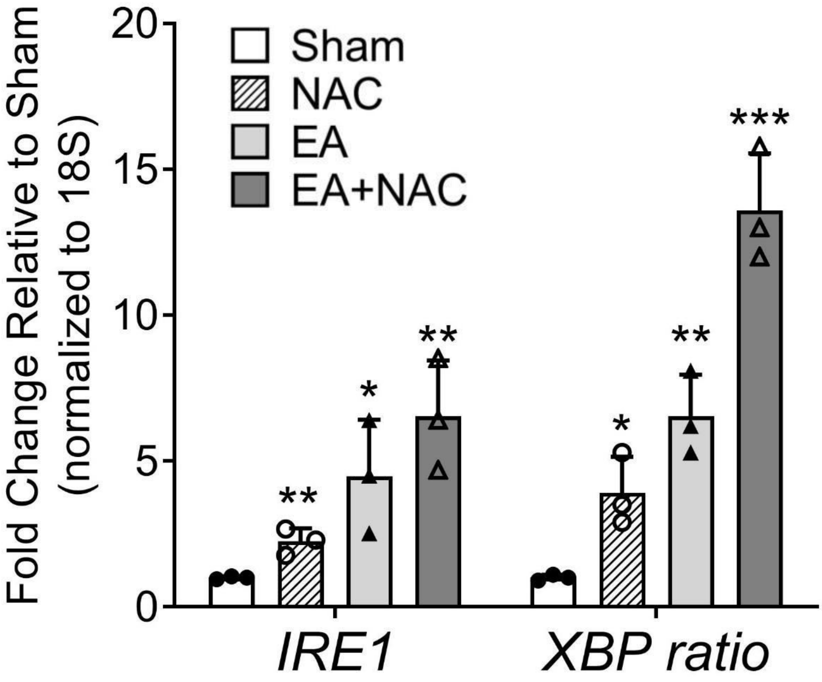

### Supplemental Fig. 3

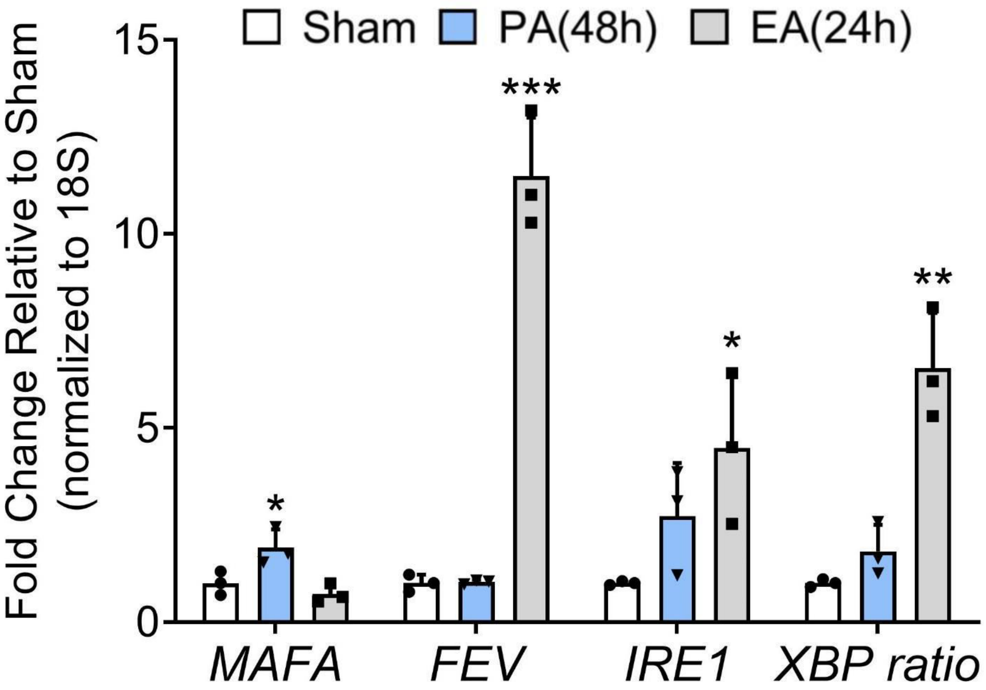

### Supplemental Fig. 4

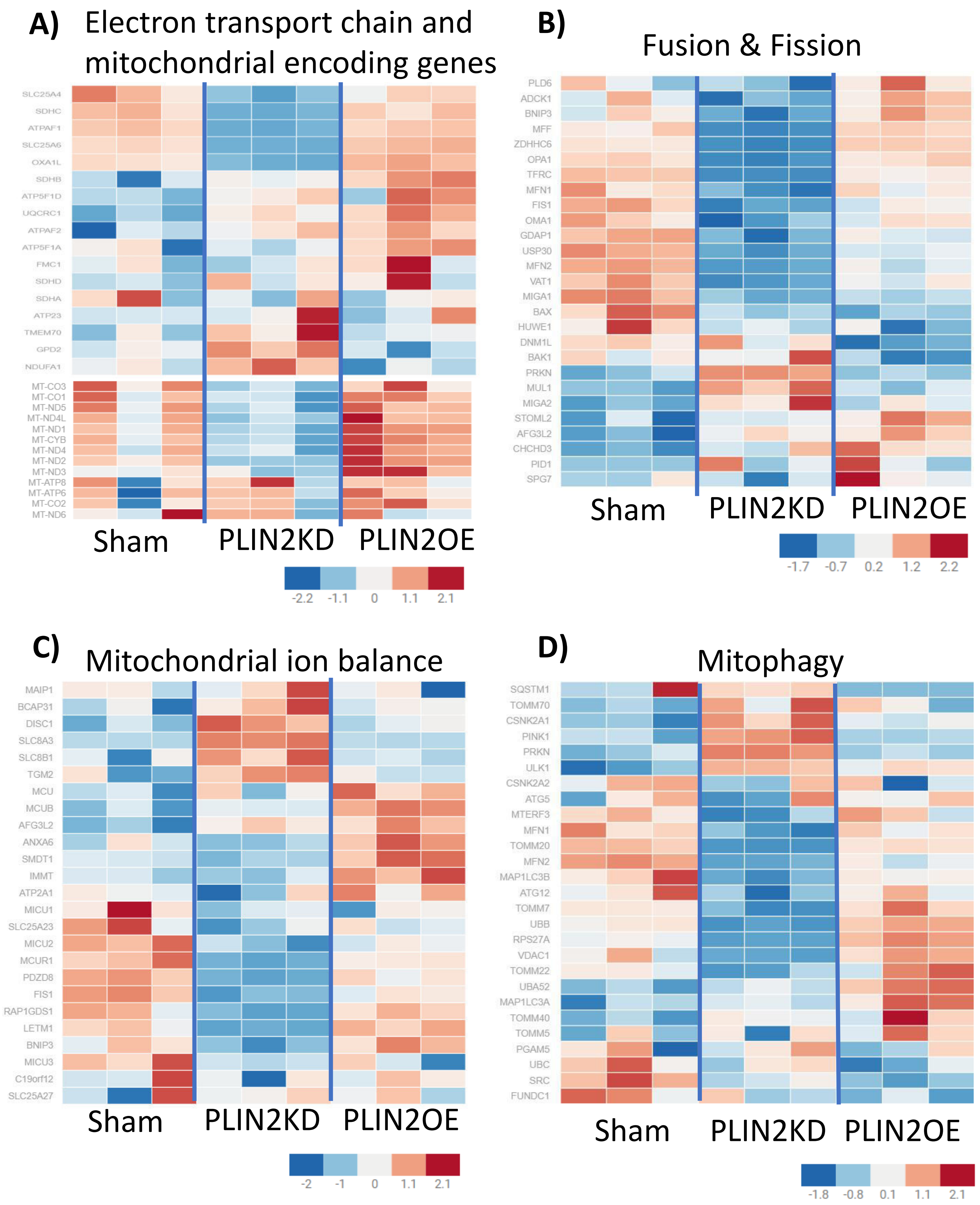

### Supplemental Fig. 5

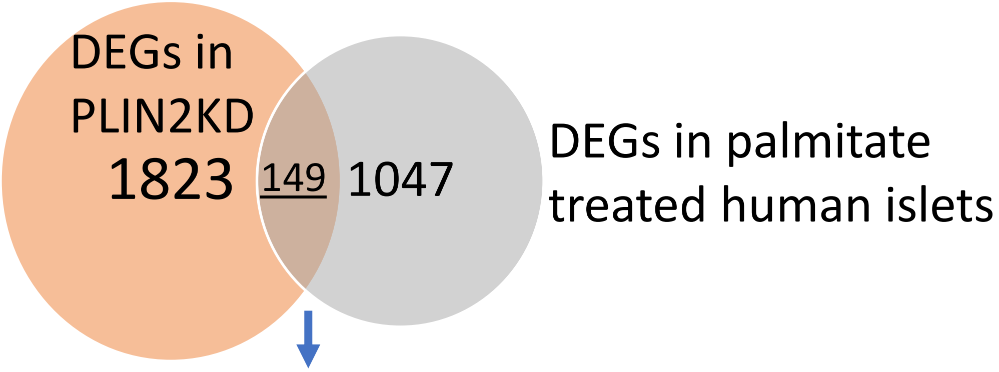


| KEGG | P-value | Genes |
|---|---|---|
| Pathways in cancer | 1.72E-04 | NOTCH2;PTGER4;TGFB2;FOS;PLD1;FGF2;RASGRP1;AGT;SMO;LPAR6;MYC;FGF18;HMOX1 |
| Ferroptosis | 2.19E-04 | ACSL1;HMOX1;SLC7A11;GCLM |
| MAPK signaling pathway | 3.77E-04 | DUSP4;TGFB2;DUSP3;MYC;HSPA6;FGF18;FOS;FGF2;RASGRP1 |
| Legionellosis | 7.51E-04 | EEF1A2;HSPA6;CXCL1;TLR5 |
| Breast cancer | 8.25E-04 | NOTCH2;SHC4;MYC;FGF18;FOS;FGF2 |
| Neuroactive ligand-receptor interaction | 9.95E-04 | PTGER4;PYY;EDN3;LPAR6;LEPR;IAPP;ADM;GCG;AGT |
| Parathyroid hormone synthesis, secretion and action | 0.001189 | EGR1;CASR;MMP14;FOS;PLD1 |
| Circadian rhythm | 0.001563 | BHLHE40;BHLHE41;NPAS2 |
| Adipocytokine signaling pathway | 0.001757 | ACSL1;LEPR;PPARGC1A;PCK2 |
| MicroRNAs in cancer | 0.001836 | NOTCH2;SHC4;TGFB2;MYC;DDIT4;HMOX1;SPRY2;VIM |
| PPAR signaling pathway | 0.002272 | FABP5;ACSL1;PLIN2;PCK2 |
| cAMP signaling pathway | 0.005176 | PDE10A;EDN3;GCG;FOS;PLD1;MYL9 |
| Gastric cancer | 0.005203 | SHC4;TGFB2;MYC;FGF18;FGF2 |
| AGE-RAGE signaling pathway in diabetic complications | 0.006675 | EGR1;TGFB2;COL1A2;AGT |
| Insulin resistance | 0.008716 | PPARGC1A;PPARGC1B;AGT;PCK2 |
| Renin-angiotensin system | 0.012589 | CTSA;AGT |
| Renin secretion | 0.014838 | PTGER4;EDN3;AGT |
| Maturity onset diabetes of the young | 0.015938 | IAPP;BHLHA15 |
| PI3K-Akt signaling pathway | 0.016991 | COL1A2;LPAR6;MYC;DDIT4;FGF18;FGF2;PCK2 |
| Vascular smooth muscle contraction | 0.017153 | EDN3;ADM;MYL9;AGT |

| GO (biological process) Term | P-value | Genes |
|---|---|---|
| insulin-like growth factor II binding (GO:0031995) | 1.39E-05 | IGFBP1;IGFBP4;IGFBP6 |
| hormone activity (GO:0005179) | 3.64E-05 | PYY;EDN3;VGF;GPNMB;GCG;AGT |
| insulin-like growth factor I binding (GO:0031994) | 1.10E-04 | IGFBP1;IGFBP4;IGFBP6 |
| insulin-like growth factor binding (GO:0005520) | 1.39E-04 | IGFBP1;IGFBP4;IGFBP6 |
| activating transcription factor binding (GO:0033613) | 0.001665 | MYC;BHLHE40;BHLHE41;FOS |
| MAP kinase phosphatase activity (GO:0033549) | 0.001918 | DUSP4;DUSP3 |
| E-box binding (GO:0070888) | 0.002047 | MYC;BHLHE40;BHLHE41 |
| G-protein coupled receptor binding (GO:0001664) | 0.003213 | PRKN;PYY;GPRC5B;ADM;GCG |
| fibroblast growth factor receptor binding (GO:0005104) | 0.003465 | FGF18;FGF2 |
| protein homodimerization activity (GO:0042803) | 0.004125 | ERN1;TRIM9;CASR;TGFB2;STC2;BHLHE40;BHLHE41;AGR2;HMOX1;RASGRP1;ATF3;MGLL |
| transcription corepressor activity (GO:0003714) | 0.004199 | ZNF85;BHLHE40;BHLHE41;PBXIP1;MEIS2;ATF3 |
| growth factor receptor binding (GO:0070851) | 0.004978 | ERN1;AGR2;TLR5;FGF2 |
| transcription factor activity, RNA polymerase II core promoter proximal region sequence-specific binding (GO:0000982) | 0.005081 | TSHZ3;MYC;BHLHE40;BHLHE41;FOS;MEIS2;ATF3 |
| RNA polymerase II activating transcription factor binding (GO:0001102) | 0.00517 | BHLHE40;BHLHE41;FOS |
| cytokine activity (GO:0005125) | 0.006134 | TGFB2;GDF15;TNFRSF11B;CXCL1;FGF2 |
| transcription regulatory region DNA binding (GO:0044212) | 0.007055 | PRKN;EGR1;MYC;BHLHE40;BHLHE41;FOS;ATF3;NPAS2 |
| bHLH transcription factor binding (GO:0043425) | 0.01155 | BHLHE40;BHLHE41 |
| ligand-dependent nuclear receptor transcription coactivator activity (GO:0030374) | 0.012639 | FGF2;PPARGC1A;PPARGC1B |
| carboxy-lyase activity (GO:0016831) | 0.015938 | DDC;PCK2 |
| Hsp70 protein binding (GO:0030544) | 0.01713 | PRKN;ERN1 |

### Supplemental Fig. 6

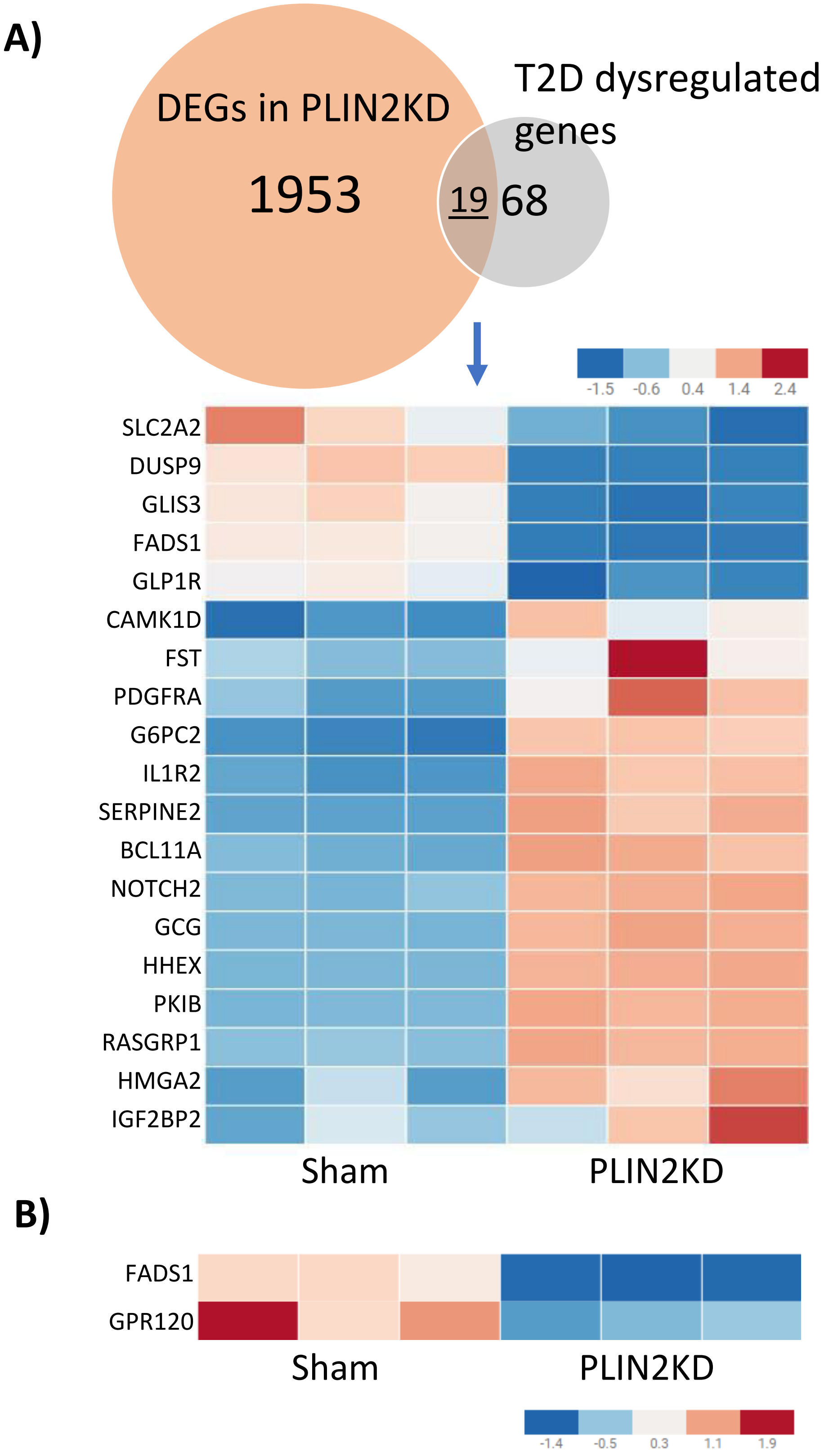

### Supplemental Fig. 7

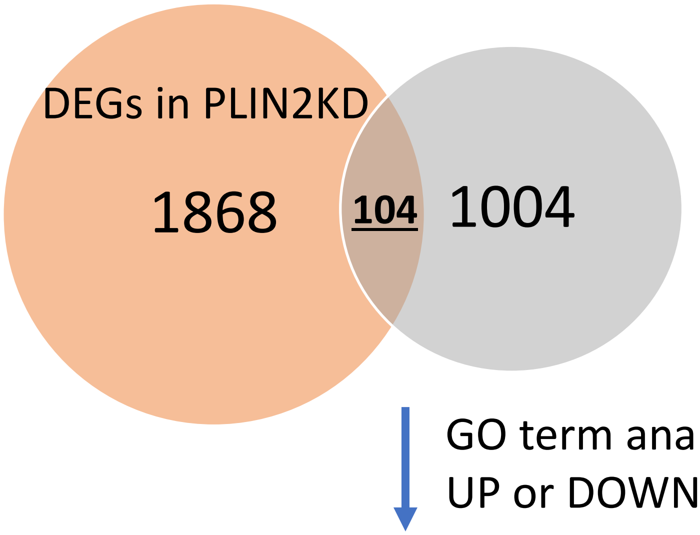


Master gene list including genes involved in:

- LD homeostasis
- Lipid metabolism
- Phospholipid metabolism
- Lipase

GO term analysis on
UP or DOWN regulated DEGs

| rank | Top 10 GO term (biological process) on UP regulated DEGs in P2KD | P value |
|---|---|---|
| 1 | phospholipid transport (GO:0015914) | 3.75E-13 |
| 2 | regulation of phosphatidylinositol 3-kinase activity (GO:0043551) | 1.39E-09 |
| 3 | phosphatidylcholine metabolic process (GO:0046470) | 4.75E-09 |
| 4 | cholesterol transport (GO:0030301) | 8.60E-09 |
| 5 | positive regulation of phosphatidylinositol 3-kinase activity (GO:0043552) | 2.12E-08 |
| 6 | lipid transport (GO:0006869) | 2.74E-08 |
| 7 | positive regulation of lipid kinase activity (GO:0090218) | 3.21E-08 |
| 8 | positive regulation of phospholipid metabolic process (GO:1903727) | 3.90E-08 |
| 9 | regulation of lipid metabolic process (GO:0019216) | 5.29E-08 |
| 10 | high-density lipoprotein particle remodeling (GO:0034375) | 2.98E-07 |

| rank | Top 10 GO term (biological process) on DOWN regulated DEGs in P2KD | P value |
|---|---|---|
| 1 | glycerophospholipid biosynthetic process (GO:0046474) | 3.08E-07 |
| 2 | fatty acid catabolic process (GO:0009062) | 4.38E-06 |
| 3 | long-chain fatty acid transport (GO:0015909) | 1.01E-05 |
| 4 | phosphatidylinositol biosynthetic process (GO:0006661) | 2.25E-05 |
| 5 | phosphatidylinositol metabolic process (GO:0046488) | 3.8E-05 |
| 6 | fatty acid homeostasis (GO:0055089) | 0.0001 |
| 7 | long-chain fatty acid metabolic process (GO:0001676) | 0.000111 |
| 8 | carnitine shuttle (GO:0006853) | 0.000153 |
| 9 | fatty acid transmembrane transport (GO:1902001) | 0.000153 |
| 10 | lipid biosynthetic process (GO:0008610) | 0.000247 |
