## Supplemental Tables and legends for "Lipid droplets protect human β cells from lipotoxic-induced stress and cell identity changes"

### Supplemental Table and Legends

**Supplemental Table 1. Primers used in qPCR**

**Supplemental Table 2. Differentially expressed genes in PLIN2KD cells**

The 1972 differentially expressed gene (DEGs) ID, name and description of PLIN2KD cells were listed in alphabetical order. “Fold-change” was calculated as average TPM in PLIN2KD divided by average TPM in Sham. Log2 (fold-change) were highlighted and average TPM in PLIN2OE were italic as a reference. TPM: Transcripts Per Million

**Supplemental Table 3. Differentially expressed genes in PLIN2OE cells**

The 285 DEGs ID, name and description in PLIN2OE cells were listed in alphabetical order. “Fold-change” was calculated as average TPM in PLIN2OE divided by average TPM in Sham. Log2 (fold-change) were highlighted and average TPM in PLIN2KD were italic as a reference.

**Supplemental Table 4. Pathway analysis of up or down regulated PLIN2KD DEGs**

Kyoto Encyclopedia of Genes and Genomes (KEGG) pathway (Human 2019) and GO molecular function (2018) analyses of PLIN2KD genes was performed using the Enrichr online server on 1479 upregulated and 493 down regulated DEGs. Top 20 enriched pathways/molecular function terms were listed in ascending order of the p-value. DEGs included in each term were also listed. Highlighted terms were mentioned in the main text.

**Supplemental Table 5. Pathway analysis of up or down regulated PLIN2OE DEGs**

KEGG pathway (Human 2019) and GO molecular function (2018) analyses of PLIN2OE genes was performed using Enrichr online server on 237 upregulated and 48 down regulated DEGs. Top 20 enriched pathways/molecular function terms were listed in ascending order of the p-value. DEGs included in each term were also listed. Highlighted terms were mentioned in the main text.

**Supplemental Table 6. Top up and down regulated PLIN2KD genes**

Top 20 most up or down regulated genes in PLIN2KD (readily detectable with average TPM>5) were listed in descending order of the absolute value of the log2 (Fold-change). “Fold-change” was calculated as average TPM in PLIN2KD divided by average TPM in Sham. Highlighted genes belong to pathways that were discussed in the main text.

**Supplemental Table 7. Top up and down regulated genes in PLIN2OE genes**

Top 20 most up or down regulated genes in PLIN2OE (readily detectable with average TPM>5) were listed in descending order of the absolute value of log2 (Fold-change). “Fold-change” was calculated as average TPM in PLIN2OE divided by average TPM in Sham. Highlighted genes belong to pathways that were discussed in the main text.

**Supplemental Table 1**

| **Gene** | **Forward 5'-3'** | **Reverse 3'-5'** |
|---|---|---|
| GCG | CTGAAGGGACCTTTACCAGTGA | CCTGGCGGCAAGATTATCAAG |
| HHEX | ACCCGACGCCCTTTTACATC | GAGAAGGCTGGATGGATCGG |
| SST | CGCTGTCCATCGTCCTG | GGGCATCATTCTCCGTCT |
| CHGA | TGACCTCAACGATGCATTTC | CTGTCCTGGCTCTTCTGCTC |
| GHRL | CAACGCCCCCTTTGATGTTG | CTGCTGGTACTGAACCCCTG |
| EXOC5 | CTTCAGTAATCCAGAAACAGTCCT | TGCTCTGCATCGGACTTCCTAC |
| GLP1R | GGCTTGAGCACTTGCACATC | GGCACTGCCCATTTACCTGT |
| KCNJ11 | TGTGTCACCAGCATCCACTCCT | GTTCTGCACGATGAGGATCAGG |
| SNAP25 | CGTCGTATGCTGCAACTGGTTG | GGTTCATGCCTTCTTCGACACG |
| STX1A | GGAACACGCGGTAGACTATGT | CTGGAGTGGAGTGGCAGTTT |
| VAMP2 | CTCCAAACCTCACCAGTAACAGG | AGCTCCGACAGCTTCTGGTCTC |
| MAFA | GGAACGGAGAACCACGTTCAACGT | TGAGCGGAGAACGGTGATTTCTAAGG |
| PDX1 | TACTGGATTGGCGTTGTTTGTGGC | AGGGAGCCTTCCAATGTGTATGGT |
| NKX6.1 | ATTCGTTGGGGATGACAGAG | CGAGTCCTGCTTCTTCTTGG |
| NKX2.2 | CCTTCTACGACAGCAGCGACAA | ACTTGGAGCTTGAGTCCTGAGG |
| UCN3 | CAGCCACAAGTTCATGGGGA | ATCTCTCCCCGAGAGTGGAC |
| SOX9 | AGGAAGCTCGCGGACCAGTAC | GGTGGTCCTTCTTGTGCTGCAC |
| AMIGO2 | GGTACTTCTGCTCCAGGATAGC | GTCTTGCTGTCACAGTGGACCA |
| DUSP6 | CTCGGATCACTGGAGCCAAAAC | GTCACAGTGACTGAGCGGCTAA |
| FCGRT | GTCAAAAGTGGCGATGAGCACC | CGTGAGTAGCAAGACACCGATG |
| PBXIP1 | GACTCTGATAATAGCTGGGTGCT | GTCCCTTCTCCATCCATGGTC |
| STC2 | GCATGACTTTTCTGCACAACGCT | GGCTTATGCAGCCGAACCTGTG |
| FEV | CATTACCACTAGACGGGGCG | GATTGAGGGAGCTTCGGTCC |
| ATF4 | GCTAAGGCGGGCTCCTCCGA | ACCCAACAGGGCATCCAAGTCG |
| BIP/HSP5A | TGACATTGAAGACTTCAAAGCT | CTGCTGTATCCTCTTCACCAGT |
| CHOP/DDIT3 | GGAGCATCAGTCCCCCACTT | TGTGGGATTGAGGGTCACATC |
| PERK/EIF2AK3 | AATGCCTGGGACGTGGTGGC | TGGTGGTGCTTCGAGCCAGG |
| DNAJB9 | TCGGCATCAGAGCGCCAAATCA | ACCACTAGTAAAAGCACTGTGTCCAAG |
| IRE1/ERN1 | TGCTTAAGGACATGGCTACCATCA | CTGGAACTGCTGGTGCTGGA |
| Total XBP-1 | GGCATCCTGGCTTGCCTCCA | GCCCCCTCAGCAGGTGTTCC |
| Spliced XBP-1 | CGCTTGGGGATGGATGCCCTG | CCTGCACCTGCTGCGGACT |
| WFS1 | TGAGGACCTGCCACTGCGTCT | GAAGATGAGCGCGTTGATGTGG |
| SERCA2b/ATP2/ | ACAATGGCGCTCTCTGTTCT | ATCACCAGGGGCATTATGAG |
| P21/CDKN1A | CCTGTCACTGTCTTGTACCCT | GCGTTTGGAGTGGTAGAAATCT |
| IGFBP3 | CGCTACAAAGTTGACTACGAGTC | GTCTTCCATTTCTCTACGGCAGG |
| ID2 | CGCATCCCACTATTGTCAGC | ATTCAGAAGCCTGCAAGGACA |
| LAMC1 | CTTCTGAGGACACTGGCAGG | CTTTGTCACCGGCCCTTTTG |
| PLIN1 | TGTGCAATGCCTATGAGAAGG | AGGGCGGGGATCTTTTCCT |
| PLIN2 | TTGCAGTTGCCAATACCTATGC | CCAGTCACAGTAGTCGTCACA |
| PLIN3 | TATGCCTCCACCAAGGAGAG | ATTCGCTGGCTGATGCAATCT |
| PLIN4 | GGCACCAAGAACACTGTCTG | TCGTACCCATGACCATAGACTT |
| PLIN5 | AAGGCCCTGAAGTGGGTTC | GCATGTGGTCTATCAGCTCCA |
| 18S | GGATGTAAAGGATGGAAAATACA | TCCAGGTCTTCACGGAGCTTGTT |

**Supplemental Table 2**

| **Gene ID** | **Gene name** | **Log2(fold-change)** | **Average TPM in Sham** | **Average TPM in PLIN2KD** | ***Average TPM in PLIN2OE*** |
|---|---|---|---|---|---|
| ENSG00000245105 | A2M-AS1 | **3.012865719** | 0.12460112 | 8.324611328 | *0.47640881* |
| ENSG00000240602 | AADACP1 | **4.795163272** | 0 | 0.20105822 | *0.013474925* |
| ENSG00000165029 | ABCA1 | **1.766517312** | 1.364279519 | 3.404202956 | *2.231433114* |
| ENSG00000167972 | ABCA3 | **-1.308866068** | 67.10300456 | 17.69383034 | *73.63544689* |
| ENSG00000198691 | ABCA4 | **4.274029283** | 0.024080549 | 1.729928519 | *0.110322708* |
| ENSG00000125257 | ABCC4 | **-1.213142632** | 1.779345554 | 9.603918887 | *1.893291931* |
| ENSG00000069431 | ABCC9 | **3.751434874** | 0.031270584 | 0.453263936 | *0.031629089* |
| ENSG00000118777 | ABCG2 | **1.062018557** | 3.377453361 | 4.944101145 | *4.540040669* |
| ENSG00000172350 | ABCG4 | **1.605523148** | 0.292247989 | 0.705847189 | *0.364527716* |
| ENSG00000143921 | ABCG8 | **2.125466902** | 0.074618614 | 2.516241852 | *0.159264508* |
| ENSG00000114786 | ABHD14A-ACY1 | **2.868078234** | 0.005019864 | 0.338454682 | *0.00643356* |
| ENSG00000229107 | ABHD17AP4 | **1.046924482** | 5.559573457 | 7.222502234 | *6.049422639* |
| ENSG00000140526 | ABHD2 | **-1.160133808** | 60.23747818 | 17.93476665 | *71.44479063* |
| ENSG00000173210 | ABLIM3 | **1.956058621** | 0.017087596 | 4.229576903 | *0.017001287* |
| ENSG00000278727 | AC000403.1 | **1.13374155** | 1.800621603 | 11.16732415 | *1.554481062* |
| ENSG00000274898 | AC001226.1 | **1.979866466** | 0.909683583 | 2.273610682 | *2.083181627* |
| ENSG00000248636 | AC002070.1 | **4.244277406** | 0 | 1.620496735 | *0.061765895* |
| ENSG00000265975 | AC002091.1 | **-1.589810814** | 4.72164006 | 2.092460328 | *4.172180876* |
| ENSG00000266389 | AC002091.2 | **-1.753775625** | 5.468794911 | 1.199148539 | *5.030607688* |
| ENSG00000231170 | AC002451.1 | **4.334843559** | 0.08658557 | 2.071690466 | *0.142182371* |
| ENSG00000284130 | AC002472.3 | **2.050967779** | 0.807179539 | 2.444292985 | *1.698266941* |
| ENSG00000232759 | AC002480.1 | **2.903998468** | 0.39500518 | 2.430574601 | *0.554426119* |
| ENSG00000232949 | AC002480.2 | **4.128380617** | 0 | 1.738952833 | *0* |
| ENSG00000271133 | AC004130.2 | **2.286396307** | 0.38674623 | 2.214338215 | *0.553937335* |
| ENSG00000263571 | AC004147.2 | **-1.432087365** | 1.134598122 | 1.055925433 | *0.767210415* |
| ENSG00000272079 | AC004233.2 | **1.200117325** | 16.48153541 | 24.61424586 | *24.58921818* |
| ENSG00000281530 | AC004461.3 | **-1.384098399** | 4.23999504 | 1.548045919 | *1.655809406* |
| ENSG00000285081 | AC004593.1 | **3.891359957** | 0.130015489 | 14.68759875 | *0.118454819* |
| ENSG00000272768 | AC004854.2 | **1.473750581** | 3.094213603 | 5.743538009 | *4.055707262* |
| ENSG00000272812 | AC004908.2 | **1.190346985** | 5.237415205 | 8.894777948 | *6.553463541* |
| ENSG00000230882 | AC005077.4 | **2.262992901** | 0.487693778 | 4.126968233 | *0.245534891* |
| ENSG00000274712 | AC005332.4 | **1.20410917** | 18.59803398 | 29.60178284 | *22.80476176* |
| ENSG00000259065 | AC005520.4 | **1.106675758** | 2.13674575 | 3.675811276 | *2.413351934* |
| ENSG00000235852 | AC005540.1 | **2.76552293** | 45.35880011 | 207.1559759 | *63.08332307* |
| ENSG00000267131 | AC005746.2 | **1.517956592** | 6.185327386 | 14.36894259 | *9.906857887* |
| ENSG00000247011 | AC005920.1 | **1.161665508** | 1.176439174 | 114.9956378 | *1.136166584* |
| ENSG00000278627 | AC005962.1 | **-1.342420138** | 5.539459332 | 6.553853203 | *4.011899204* |
| ENSG00000231916 | AC006033.1 | **1.521275991** | 3.538221274 | 7.635421848 | *3.692717208* |
| ENSG00000255966 | AC006064.3 | **1.13690649** | 5.116178347 | 8.720789503 | *6.525784983* |
| ENSG00000278847 | AC006157.1 | **1.363230581** | 3.598330898 | 9.374346379 | *3.636564639* |
| ENSG00000236213 | AC006369.1 | **1.272849393** | 86.14887045 | 140.7193666 | *113.6872903* |
| ENSG00000275532 | AC006449.2 | **3.182299946** | 0.284744858 | 4.650805987 | *0.121329775* |
| ENSG00000277969 | AC006449.7 | **1.126811072** | 6.645513034 | 76.49910234 | *8.928785445* |
| ENSG00000261770 | AC006504.1 | **1.064441339** | 1.73279041 | 3.422131471 | *1.457128099* |

| | | | | | |
|---|---|---|---|---|---|
| ENSG00000258881 | AC007040.2 | **-1.280253008** | 2.795492304 | 6.033108111 | *1.756154566* |
| ENSG00000277196 | AC007325.2 | **-1.482180306** | 53.07611059 | 13.04530006 | *47.08485122* |
| ENSG00000279722 | AC007342.7 | **-1.100922709** | 3.354338235 | 1.403751746 | *2.071458781* |
| ENSG00000266473 | AC007448.4 | **1.222841511** | 4.129742734 | 13.18731002 | *3.124812871* |
| ENSG00000260086 | AC007611.1 | **2.028557066** | 0.703580692 | 2.289577155 | *1.084294494* |
| ENSG00000270953 | AC007938.3 | **-1.178017351** | 28.54711762 | 10.83354571 | *15.81706632* |
| ENSG00000233483 | AC008105.2 | **1.229610682** | 0.620903084 | 2.043000688 | *0.938839951* |
| ENSG00000285188 | AC008397.2 | **1.437798082** | 0.657401349 | 5.836037352 | *1.308613084* |
| ENSG00000260686 | AC008669.1 | **-1.474854039** | 18.91155064 | 4.596089329 | *16.87485656* |
| ENSG00000253357 | AC008708.1 | **1.527703402** | 13.86231753 | 27.4880593 | *23.83101584* |
| ENSG00000270442 | AC008725.1 | **2.440646547** | 0.148128651 | 3.00696136 | *0.211819558* |
| ENSG00000248918 | AC008808.1 | **1.659298444** | 5.19349059 | 24.10435795 | *8.562258312* |
| ENSG00000276259 | AC009118.3 | **-1.313302263** | 14.39903639 | 4.548018833 | *15.85576284* |
| ENSG00000276261 | AC009509.5 | **-2.789174914** | 5.161225694 | 4.911799286 | *3.459976852* |
| ENSG00000272986 | AC009570.1 | **1.806078548** | 3.915820404 | 10.01393426 | *4.310618955* |
| ENSG00000263823 | AC009831.1 | **1.762028061** | 4.522799174 | 10.10267025 | *4.817090626* |
| ENSG00000272425 | AC009902.3 | **2.391370293** | 0.319724348 | 6.258732386 | *0.442137515* |
| ENSG00000271259 | AC010201.1 | **1.838844276** | 1.421941047 | 8.923178145 | *0.90226817* |
| ENSG00000271327 | AC010201.2 | **1.589756093** | 1.165184672 | 2.539343353 | *1.392860492* |
| ENSG00000277383 | AC010331.1 | **1.209595607** | 9.519459409 | 14.78288789 | *7.032045464* |
| ENSG00000271714 | AC010501.2 | **1.593741956** | 0.938705753 | 2.99300644 | *1.664097221* |
| ENSG00000270006 | AC010531.6 | **1.11756882** | 7.827290505 | 20.53785123 | *6.147006059* |
| ENSG00000226180 | AC010536.1 | **1.528054513** | 1.54073565 | 4.093833081 | *1.210805364* |
| ENSG00000277504 | AC010536.3 | **1.393246993** | 2.846315641 | 10.05405274 | *1.219018811* |
| ENSG00000213976 | AC010615.1 | **1.86514938** | 3.319260688 | 9.109414253 | *3.639200836* |
| ENSG00000265474 | AC010761.4 | **-4.407133342** | 2.375821413 | 1.887369243 | *2.295447426* |
| ENSG00000273008 | AC010864.1 | **1.324237585** | 2.300866628 | 7.922835072 | *3.299337244* |
| ENSG00000234936 | AC010883.1 | **1.617270379** | 3.380059323 | 6.600900212 | *4.821653276* |
| ENSG00000276524 | AC010999.2 | **1.712786169** | 1.39799604 | 5.028597266 | *1.712511172* |
| ENSG00000283458 | AC011139.1 | **2.963292831** | 0.264688084 | 5.36087373 | *1.657137114* |
| ENSG00000266826 | AC011195.3 | **-1.606954517** | 1.673985739 | 1.832935369 | *0.806987101* |
| ENSG00000279739 | AC011369.2 | **1.534344672** | 0.783420948 | 2.021193439 | *0.683947529* |
| ENSG00000268583 | AC011466.1 | **-2.094837477** | 3.303491872 | 0.485973258 | *2.354893421* |
| ENSG00000280239 | AC011498.7 | **1.045565088** | 1.709380227 | 3.10173307 | *4.191496536* |
| ENSG00000227799 | AC012358.3 | **1.120996955** | 8.498592779 | 12.65034917 | *7.940688892* |
| ENSG00000260163 | AC012508.1 | **1.22619671** | 0.565803312 | 2.085761291 | *0.79439326* |
| ENSG00000272564 | AC012511.1 | **2.988887361** | 1.061226667 | 11.12481379 | *2.44626347* |
| ENSG00000279348 | AC012513.3 | **-1.487623242** | 22.83224245 | 5.670600005 | *21.63440113* |
| ENSG00000234773 | AC012618.3 | **1.149857712** | 1.143650885 | 4.931245641 | *1.55340004* |
| ENSG00000259793 | AC013726.2 | **2.023481746** | 0.278336169 | 3.247373908 | *0.4869842* |
| ENSG00000234595 | AC013733.2 | **1.772491489** | 0.97405216 | 2.688378489 | *1.561530128* |
| ENSG00000272384 | AC016405.3 | **2.595575509** | 0.329427889 | 1.844691096 | *0.499867329* |
| ENSG00000267422 | AC016582.2 | **1.312858866** | 2.156821533 | 4.757952934 | *1.061007026* |
| ENSG00000226380 | AC016831.1 | **-1.70549998** | 0.344945848 | 0.767452758 | *0.301429956* |
| ENSG00000271930 | AC016877.3 | **1.539507806** | 2.21530481 | 5.485302736 | *3.472214966* |
| ENSG00000272777 | AC019131.2 | **-2.572055163** | 2.22978769 | 0.229624831 | *1.39805154* |
| ENSG00000270988 | AC019257.2 | **2.144246376** | 1.237937888 | 6.316650673 | *2.371843262* |

| | | | | | |
|---|---|---|---|---|---|
| ENSG00000272267 | AC021242.3 | **1.427453789** | 1.955280576 | 3.317033354 | *2.152994786* |
| ENSG00000271966 | AC021321.1 | **1.549837178** | 4.053732277 | 10.01701421 | *4.054835445* |
| ENSG00000266521 | AC021549.2 | **1.931221365** | 1.117282837 | 5.227673304 | *2.88370575* |
| ENSG00000267780 | AC021594.2 | **2.28040448** | 0.780139087 | 6.043855987 | *0.247499281* |
| ENSG00000275091 | AC022098.4 | **1.506444621** | 3.142986631 | 6.65465995 | *6.928578037* |
| ENSG00000254777 | AC022182.1 | **1.196046306** | 0.907739231 | 2.565559176 | *1.066936559* |
| ENSG00000256802 | AC022613.1 | **2.551020838** | 2.991306706 | 14.42605488 | *4.192714964* |
| ENSG00000271870 | AC024060.2 | **-1.424538828** | 23.440565 | 6.530203185 | *24.64458033* |
| ENSG00000272969 | AC024243.1 | **1.900483727** | 2.038676168 | 11.23866433 | *4.16618556* |
| ENSG00000269514 | AC024257.3 | **1.06054152** | 0.784111482 | 3.857337676 | *0.990410995* |
| ENSG00000279668 | AC024610.2 | **-1.950681001** | 0.69517721 | 2.475794735 | *0.502782425* |
| ENSG00000258302 | AC025034.1 | **4.210246086** | 0 | 0.891228836 | *0.103389136* |
| ENSG00000272382 | AC025171.5 | **3.053905306** | 0.529403761 | 2.677605725 | *0.725129012* |
| ENSG00000233912 | AC026202.2 | **2.403356209** | 0.41672618 | 1.338812183 | *0.445615747* |
| ENSG00000256694 | AC026369.2 | **-3.81085066** | 0.738275186 | 1.700911751 | *0.112790492* |
| ENSG00000280132 | AC026471.6 | **2.459152667** | 0.283116519 | 1.839544654 | *0.470590198* |
| ENSG00000271869 | AC026979.2 | **1.081472303** | 17.27139474 | 22.97378024 | *24.93859053* |
| ENSG00000268798 | AC027307.3 | **-1.03124224** | 6.997323734 | 2.651448505 | *8.051856399* |
| ENSG00000275155 | AC027348.1 | **2.382448315** | 2.287990737 | 20.9012311 | *3.143325114* |
| ENSG00000253622 | AC027419.2 | **3.025379148** | 0.94634436 | 7.007892993 | *3.126938641* |
| ENSG00000185332 | AC027601.1 | **3.757228456** | 0.037328694 | 4.22737261 | *0.034667538* |
| ENSG00000275542 | AC027601.3 | **1.322041098** | 7.397008313 | 14.06044997 | *9.217206334* |
| ENSG00000253171 | AC037450.1 | **4.905085373** | 0.112287743 | 2.383207276 | *0.091137976* |
| ENSG00000279369 | AC046185.3 | **1.252478903** | 17.76878163 | 34.65293709 | *21.17115669* |
| ENSG00000279146 | AC063926.2 | **1.087112893** | 2.465583753 | 5.054661284 | *1.555858965* |
| ENSG00000253844 | AC064807.2 | **2.044044635** | 4.80916417 | 26.69064476 | *6.399610958* |
| ENSG00000260274 | AC068338.2 | **-1.14829585** | 30.44028421 | 10.0187011 | *24.89073067* |
| ENSG00000283674 | AC068587.4 | **2.141442374** | 0.026365068 | 6.400266883 | *0.098758002* |
| ENSG00000259200 | AC068722.1 | **1.555080236** | 0.94823333 | 6.430179106 | *1.786735926* |
| ENSG00000260473 | AC068987.2 | **1.580756438** | 0.427584119 | 0.881260115 | *0.54457923* |
| ENSG00000256364 | AC069234.2 | **-2.440658687** | 1.268742476 | 1.309357865 | *0.874817931* |
| ENSG00000277423 | AC069234.5 | **-1.785210275** | 7.230263828 | 1.550112203 | *4.097021145* |
| ENSG00000241634 | AC069499.2 | **1.99894095** | 1.188748134 | 3.022603387 | *4.557090811* |
| ENSG00000259826 | AC072061.1 | **1.092053175** | 4.094231146 | 6.643153931 | *4.348669125* |
| ENSG00000271855 | AC073195.2 | **1.036186241** | 7.755713778 | 13.33218935 | *9.103397778* |
| ENSG00000226087 | AC073283.2 | **-4.524867576** | 0.597468452 | 2.926943037 | *1.973621394* |
| ENSG00000257605 | AC073611.1 | **1.547251544** | 2.089919551 | 7.861117954 | *3.148300688* |
| ENSG00000272368 | AC074032.1 | **1.056718134** | 3.073649904 | 4.069565787 | *2.856631724* |
| ENSG00000259605 | AC074212.1 | **1.559857415** | 0.953460945 | 4.284040266 | *1.136445199* |
| ENSG00000274444 | AC078909.2 | **-2.29196923** | 4.014537221 | 2.506218489 | *1.968401685* |
| ENSG00000280002 | AC078925.4 | **-2.608380615** | 6.757431768 | 1.558364223 | *7.241189868* |
| ENSG00000246528 | AC079089.1 | **1.046186942** | 0.959597997 | 1.748076828 | *0.985189962* |
| ENSG00000227210 | AC079145.1 | **-1.506147595** | 21.46060277 | 5.284862562 | *25.67752812* |
| ENSG00000222043 | AC079305.2 | **1.634203121** | 3.849552786 | 8.230663621 | *3.38883782* |
| ENSG00000270028 | AC079315.1 | **1.009141414** | 8.332491163 | 13.85012517 | *7.521920746* |
| ENSG00000243144 | AC079760.2 | **3.893641074** | 0.080722411 | 5.099741596 | *0.536080617* |
| ENSG00000254554 | AC080023.1 | **3.104749519** | 0.266879405 | 6.89130322 | *0.305242764* |

| | | | | | |
|---|---|---|---|---|---|
| ENSG00000263893 | AC080037.2 | **-1.465782616** | 11.00932727 | 3.166545643 | *8.997284324* |
| ENSG00000279675 | AC080100.1 | **5.082022085** | 0 | 0.880723097 | *0* |
| ENSG00000243491 | AC082651.3 | **2.850103979** | 0.138407451 | 1.856555717 | *0.428614787* |
| ENSG00000254577 | AC087276.2 | **-2.341769917** | 13.51963108 | 1.753351954 | *6.18906594* |
| ENSG00000283415 | AC087280.2 | **3.802001717** | 0.042252898 | 0.724851376 | *0.124540632* |
| ENSG00000271851 | AC087501.4 | **-1.083260612** | 10.28456165 | 3.964647043 | *11.4344193* |
| ENSG00000197376 | AC089987.1 | **4.574292268** | 0 | 0.503986535 | *0.021760497* |
| ENSG00000254102 | AC090136.3 | **4.263682403** | 0.026575707 | 2.103865193 | *0.055834637* |
| ENSG00000257228 | AC090531.1 | **1.280918007** | 6.926864223 | 10.3185253 | *10.9755697* |
| ENSG00000257729 | AC090679.2 | **1.028315826** | 90.13828993 | 119.5990856 | *122.9714975* |
| ENSG00000274528 | AC090970.2 | **1.776051915** | 2.37172278 | 11.73225902 | *2.813436363* |
| ENSG00000187951 | AC091057.1 | **1.635219581** | 1.605739931 | 65.31669565 | *2.794935601* |
| ENSG00000257060 | AC091078.1 | **-2.896764316** | 1.258467235 | 2.8128254 | *1.252901834* |
| ENSG00000263958 | AC091138.1 | **1.600760608** | 1.388187031 | 4.740314131 | *1.143719583* |
| ENSG00000251127 | AC091173.1 | **3.132603247** | 0.048996238 | 0.295410894 | *0.109484052* |
| ENSG00000271320 | AC091488.1 | **4.227002331** | 0 | 1.711622365 | *0* |
| ENSG00000279130 | AC091925.1 | **1.11532783** | 4.644570754 | 6.134228388 | *5.673618732* |
| ENSG00000276417 | AC092111.2 | **-1.870298626** | 1.820918224 | 0.704361318 | *0.994316046* |
| ENSG00000276791 | AC092117.1 | **1.338441559** | 4.410549895 | 11.40716798 | *3.517893817* |
| ENSG00000187185 | AC092118.1 | **1.011429512** | 4.103144259 | 5.391187311 | *4.701545807* |
| ENSG00000284670 | AC092128.1 | **4.335732087** | 0 | 4.005892098 | *0.051923615* |
| ENSG00000279294 | AC092135.1 | **-1.654534757** | 9.983458723 | 4.707146472 | *10.75153959* |
| ENSG00000277954 | AC092376.2 | **1.588942377** | 0.333189393 | 0.843709174 | *0.408866196* |
| ENSG00000261226 | AC092384.3 | **1.428258737** | 0.567417017 | 2.216555023 | *0.241484551* |
| ENSG00000224593 | AC092427.1 | **-1.284054328** | 4.364996913 | 1.579513947 | *0.914876362* |
| ENSG00000230393 | AC092667.1 | **3.417118172** | 0.081460872 | 1.337445338 | *0.300531354* |
| ENSG00000239268 | AC092691.1 | **4.98925813** | 0.386830617 | 8.544516737 | *0.428218028* |
| ENSG00000256625 | AC092747.2 | **-2.026126954** | 2.458195385 | 0.661221337 | *1.457364607* |
| ENSG00000279746 | AC092850.1 | **-1.736297419** | 2.960138123 | 4.342481027 | *2.927964166* |
| ENSG00000236412 | AC092941.2 | **-2.676564648** | 2.414189219 | 0.42624217 | *1.591081269* |
| ENSG00000232533 | AC093673.1 | **1.003182022** | 26.97054906 | 36.50759242 | *24.82979505* |
| ENSG00000245293 | AC096564.1 | **-1.360524843** | 2.475320751 | 0.622276299 | *1.624359509* |
| ENSG00000261428 | AC097461.1 | **-1.452394836** | 2.530079574 | 17.87964391 | *3.63008916* |
| ENSG00000235024 | AC097468.2 | **1.746248216** | 2.594271416 | 5.651185079 | *1.868744837* |
| ENSG00000261220 | AC103706.1 | **1.272977404** | 3.347921424 | 5.622713574 | *4.280157385* |
| ENSG00000271938 | AC103724.4 | **2.864400761** | 0.922874338 | 8.72213964 | *1.515018775* |
| ENSG00000272431 | AC104117.5 | **1.290138649** | 2.749569052 | 6.76946548 | *2.558415798* |
| ENSG00000271862 | AC104118.1 | **-1.486526644** | 3.237266367 | 2.056979581 | *2.315654332* |
| ENSG00000271941 | AC104184.1 | **2.370225149** | 0.675522136 | 4.581923777 | *1.183442939* |
| ENSG00000233392 | AC104809.2 | **-1.978633537** | 0.472592487 | 0.494895773 | *0.19441781* |
| ENSG00000203392 | AC105020.1 | **3.613382321** | 0.032947923 | 1.72301606 | *0* |
| ENSG00000270460 | AC106900.1 | **1.226642331** | 10.07350628 | 15.28297384 | *11.64539409* |
| ENSG00000281392 | AC107204.1 | **2.088668664** | 0.104303348 | 0.408721808 | *0.227233069* |
| ENSG00000233654 | AC108047.1 | **1.037154005** | 18.83899905 | 32.68208417 | *19.53892151* |
| ENSG00000262223 | AC110285.1 | **2.498693799** | 0.131149859 | 0.557168277 | *0.117152389* |
| ENSG00000279692 | AC110285.6 | **2.667475958** | 5.760819665 | 37.11226186 | *7.298896441* |
| ENSG00000273077 | AC110609.1 | **1.850356505** | 2.315953723 | 5.682797243 | *1.014293509* |

| | | | | | |
|---|---|---|---|---|---|
| ENSG00000253389 | AC113133.1 | **2.426493678** | 1.095051531 | 16.78191753 | *0.596587703* |
| ENSG00000272457 | AC113194.1 | **6.708935325** | 0 | 6.857675059 | *0.391296363* |
| ENSG00000246323 | AC113382.1 | **-2.510663678** | 0.979443152 | 1.775437978 | *0.706785402* |
| ENSG00000280422 | AC115284.2 | **2.862954131** | 0.210445312 | 4.858275206 | *0.482363882* |
| ENSG00000277511 | AC116407.2 | **2.217488351** | 0.715852892 | 2.254841865 | *1.629163725* |
| ENSG00000284727 | AC116562.3 | **1.204893167** | 1.49801364 | 2.576360055 | *3.908182405* |
| ENSG00000283183 | AC116565.1 | **5.437023877** | 0 | 1.56632686 | *0* |
| ENSG00000249926 | AC117500.1 | **-1.631156168** | 2.694337759 | 1.850975505 | *2.930809216* |
| ENSG00000260369 | AC120024.1 | **1.263382072** | 1.455739468 | 2.695563875 | *0.494318834* |
| ENSG00000242568 | AC121764.2 | **-3.816604102** | 1.401439444 | 0.309433998 | *0.930478846* |
| ENSG00000248285 | AC122707.1 | **4.304305308** | 0 | 4.047640032 | *0.29779023* |
| ENSG00000279674 | AC123769.1 | **-1.563945351** | 0.484047992 | 0.105882673 | *0.115704797* |
| ENSG00000272677 | AC124016.2 | **1.017016151** | 2.680751773 | 4.370672331 | *2.942979133* |
| ENSG00000279613 | AC124283.5 | **1.344974771** | 2.135077698 | 3.146877766 | *1.626826637* |
| ENSG00000267655 | AC125437.1 | **1.21774739** | 2.671535741 | 6.106126407 | *2.694901605* |
| ENSG00000262873 | AC127496.5 | **2.569090825** | 0.220884678 | 3.188261268 | *0.292453025* |
| ENSG00000275011 | AC129492.6 | **1.088251545** | 6.463978813 | 10.28851918 | *5.73128968* |
| ENSG00000274370 | AC130371.2 | **4.475142647** | 0.118903715 | 1.631082762 | *0.106736597* |
| ENSG00000280182 | AC131888.1 | **2.925091665** | 0.125980468 | 5.529096171 | *0.084478137* |
| ENSG00000272710 | AC133041.1 | **4.770966834** | 0 | 3.105642642 | *1.291231874* |
| ENSG00000278716 | AC133540.1 | **-1.646839141** | 11.75511006 | 2.325025473 | *12.28741876* |
| ENSG00000261693 | AC134682.1 | **1.215180949** | 1.122506141 | 2.437064798 | *1.024352746* |
| ENSG00000251661 | AC136475.1 | **1.08070448** | 1.08805653 | 3.194192031 | *1.195401958* |
| ENSG00000254910 | AC136475.2 | **2.338004811** | 0.890417896 | 4.201403339 | *0.910185436* |
| ENSG00000278928 | AC136621.1 | **1.17786163** | 1.011745784 | 2.361062862 | *3.522230912* |
| ENSG00000182376 | AC138028.1 | **1.410664687** | 0.472685963 | 1.907462047 | *0.472861532* |
| ENSG00000261819 | AC138932.4 | **1.522699494** | 0.688390146 | 1.849241335 | *1.503718193* |
| ENSG00000233836 | AC139769.1 | **4.517651771** | 0 | 0.623874201 | *0* |
| ENSG00000273117 | AC144652.1 | **1.681007484** | 12.39904236 | 26.1770301 | *14.14141967* |
| ENSG00000261888 | AC144831.1 | **5.099431097** | 0.079313936 | 1.766514639 | *0.046142684* |
| ENSG00000223855 | AC147651.1 | **1.779108022** | 0.197140129 | 14.10290279 | *0.382340893* |
| ENSG00000229178 | AC233280.1 | **1.287661148** | 2.087556213 | 4.281260629 | *2.32382902* |
| ENSG00000236491 | AC234771.4 | **4.128009008** | 0 | 0.748253327 | *0.089337069* |
| ENSG00000167315 | ACAA2 | **-1.399564996** | 43.59254361 | 12.50630116 | *43.70689778* |
| ENSG00000196177 | ACADSB | **-1.54657172** | 70.67340364 | 15.82584078 | *68.45735699* |
| ENSG00000176244 | ACBD7 | **1.263061644** | 15.24609966 | 28.98226643 | *17.72453673* |
| ENSG00000153086 | ACMSD | **3.02077533** | 0.053638188 | 8.706382241 | *0.118525136* |
| ENSG00000167107 | ACSF2 | **-1.00187775** | 0.499578945 | 12.67784924 | *0.693537845* |
| ENSG00000151726 | ACSL1 | **1.142803753** | 3.399666955 | 5.092943651 | *3.793642946* |
| ENSG00000111058 | ACSS3 | **2.216258528** | 0.439344777 | 1.490605964 | *0.517026316* |
| ENSG00000107796 | ACTA2 | **3.240385595** | 0.684631733 | 6.753199961 | *0.332340127* |
| ENSG00000196839 | ADA | **3.122660074** | 0.115859736 | 1.310909571 | *0.198583776* |
| ENSG00000148848 | ADAM12 | **3.359022385** | 0.24193714 | 3.825863587 | *0.345970618* |
| ENSG00000104755 | ADAM2 | **4.777255016** | 0 | 0.305457769 | *0.004472224* |
| ENSG00000168594 | ADAM29 | **1.746883323** | 0.062600244 | 0.874217994 | *0.045296749* |
| ENSG00000154734 | ADAMTS1 | **4.901918508** | 0.187484702 | 3.683116745 | *0.465900344* |
| ENSG00000166106 | ADAMTS15 | **1.091466143** | 6.342963558 | 8.691253621 | *11.93747095* |

| | | | | | |
|---|---|---|---|---|---|
| ENSG00000145536 | ADAMTS16 | **-2.779894733** | 0.403969692 | 1.883114226 | *0.650159426* |
| ENSG00000140873 | ADAMTS18 | **2.227128545** | 0.042121484 | 4.848764616 | *0.077750523* |
| ENSG00000156140 | ADAMTS3 | **2.498692406** | 0.031720683 | 0.159808497 | *0.04294931* |
| ENSG00000154736 | ADAMTS5 | **1.431837836** | 60.49228584 | 107.1237911 | *68.22104112* |
| ENSG00000136378 | ADAMTS7 | **2.933763968** | 0.170142989 | 0.882908489 | *0.310987696* |
| ENSG00000134917 | ADAMTS8 | **2.394276472** | 0.28597634 | 54.82868657 | *0.543351598* |
| ENSG00000164742 | ADCY1 | **-1.639375969** | 174.6221145 | 36.94565133 | *133.3909901* |
| ENSG00000138031 | ADCY3 | **1.681434473** | 1.687604768 | 3.931730779 | *2.884146079* |
| ENSG00000121753 | ADGRB2 | **1.504709353** | 0.141678306 | 19.10469834 | *0.311224196* |
| ENSG00000111452 | ADGRD1 | **-1.454327585** | 0.595011403 | 1.978797993 | *0.614341395* |
| ENSG00000256151 | ADGRD1-AS1 | **-2.639190227** | 3.7902195 | 0.617595349 | *4.469961497* |
| ENSG00000205336 | ADGRG1 | **1.066558058** | 0.251013236 | 0.425330144 | *0.280079098* |
| ENSG00000173698 | ADGRG2 | **1.528953932** | 0.354607127 | 0.791081548 | *0.477257078* |
| ENSG00000156920 | ADGRG4 | **-1.430324979** | 0.460384877 | 0.296885409 | *0.278976564* |
| ENSG00000159618 | ADGRG5 | **-1.027215204** | 4.104397413 | 1.54376952 | *4.417135518* |
| ENSG00000150471 | ADGRL3 | **1.196372271** | 0.269641217 | 0.457907779 | *0.523925402* |
| ENSG00000148926 | ADM | **1.864400765** | 1.366513146 | 4.018908631 | *2.43891044* |
| ENSG00000128165 | ADM2 | **1.57705995** | 19.0585057 | 37.28153423 | *16.77074127* |
| ENSG00000169252 | ADRB2 | **1.635761473** | 0.737459078 | 3.228375093 | *0.91387009* |
| ENSG00000270076 | AF131215.7 | **2.464116662** | 0.684811633 | 21.46691147 | *1.794938219* |
| ENSG00000155966 | AFF2 | **3.048605372** | 0.381097448 | 2.725723871 | *0.285627845* |
| ENSG00000144218 | AFF3 | **2.284852956** | 0.36268705 | 2.578091398 | *0.680899103* |
| ENSG00000106541 | AGR2 | **2.554682457** | 0.06744808 | 1.285614164 | *0.045263328* |
| ENSG00000188157 | AGRN | **1.260308104** | 30.6848842 | 49.22221074 | *39.72752816* |
| ENSG00000135744 | AGT | **1.80842616** | 4.170396197 | 9.550761361 | *6.574763639* |
| ENSG00000129474 | AJUBA | **1.95955004** | 0.095654144 | 24.17109191 | *0.079941967* |
| ENSG00000154027 | AK5 | **1.577010361** | 0.240947444 | 5.56677809 | *0.382682274* |
| ENSG00000228172 | AL020996.1 | **1.027601785** | 1.90414111 | 2.676428979 | *2.016161693* |
| ENSG00000236166 | AL021408.1 | **4.320140903** | 0 | 0.802551513 | *0.346847523* |
| ENSG00000228201 | AL022341.1 | **1.042941483** | 24.28549254 | 31.97642044 | *23.17948289* |
| ENSG00000216316 | AL022722.1 | **1.187619822** | 6.344136591 | 10.41956992 | *6.078610105* |
| ENSG00000275438 | AL031008.1 | **1.696291501** | 1.405133297 | 21.58411887 | *1.497800458* |
| ENSG00000272432 | AL031432.3 | **3.322539429** | 0.299588329 | 5.7184032 | *1.641079278* |
| ENSG00000272402 | AL031775.2 | **-2.059164099** | 7.819207911 | 2.201384249 | *7.501300469* |
| ENSG00000279930 | AL032819.2 | **6.153688823** | 3.166575694 | 146.4642927 | *4.350971173* |
| ENSG00000225129 | AL035250.1 | **-2.105379415** | 4.777302321 | 1.357185261 | *7.065677258* |
| ENSG00000205562 | AL049775.1 | **5.218779722** | 0 | 75.03126924 | *0* |
| ENSG00000258670 | AL049874.3 | **3.256818891** | 0.561119193 | 4.235706368 | *1.886540754* |
| ENSG00000202343 | AL078581.1 | **-2.185788453** | 23.91194055 | 4.266340068 | *19.80264331* |
| ENSG00000258399 | AL117190.1 | **-2.14192356** | 1.459596926 | 2.029556485 | *1.069291353* |
| ENSG00000273759 | AL117379.1 | **-2.122976889** | 2.763532449 | 1.693679658 | *1.583080912* |
| ENSG00000268628 | AL121761.1 | **1.312761038** | 5.519406212 | 9.295450996 | *4.512073019* |
| ENSG00000275620 | AL121827.2 | **2.025923723** | 0.430816944 | 1.391276315 | *0.140956777* |
| ENSG00000273108 | AL121929.3 | **1.33263912** | 19.89160556 | 36.94945632 | *22.16390169* |
| ENSG00000258940 | AL132639.2 | **-1.861214099** | 1.523848055 | 0.804350054 | *1.504392961* |
| ENSG00000261120 | AL133299.1 | **1.209869146** | 12.08120923 | 35.48600307 | *14.2305764* |
| ENSG00000233492 | AL133325.2 | **-1.352155997** | 9.274245922 | 2.437676598 | *11.70535024* |

| | | | | | |
|---|---|---|---|---|---|
| ENSG00000227220 | AL133346.1 | **2.23273975** | 0.705139021 | 10.92767859 | *1.554135211* |
| ENSG00000260461 | AL133355.1 | **-1.351936786** | 8.0918929 | 3.323099802 | *7.704643543* |
| ENSG00000273783 | AL136040.1 | **-1.366519752** | 17.627511 | 6.312592844 | *15.65428723* |
| ENSG00000278784 | AL136295.7 | **-1.023167691** | 15.49547236 | 6.060022182 | *13.92274559* |
| ENSG00000274270 | AL137060.3 | **1.176532683** | 7.764496701 | 12.94020386 | *8.342747069* |
| ENSG00000261135 | AL137802.2 | **2.004923046** | 0.501291566 | 3.882029307 | *0.715815987* |
| ENSG00000255872 | AL138752.2 | **1.687171522** | 0.428251365 | 6.688581315 | *0.635349573* |
| ENSG00000259775 | AL138976.2 | **1.068499957** | 4.233270392 | 7.235080433 | *3.961736492* |
| ENSG00000277763 | AL138995.1 | **1.590588486** | 1.833221629 | 4.157421618 | *1.915041148* |
| ENSG00000272449 | AL139246.5 | **1.396557987** | 3.838853569 | 9.102345004 | *5.712907875* |
| ENSG00000254718 | AL157756.1 | **1.631299232** | 0.46730188 | 2.945349357 | *0.474232178* |
| ENSG00000282787 | AL157888.1 | **1.675017262** | 5.881194802 | 15.49753908 | *6.134978586* |
| ENSG00000279513 | AL157902.2 | **1.475298703** | 0.880510132 | 2.094577784 | *0.517796014* |
| ENSG00000261671 | AL158211.1 | **3.118072287** | 0.185653987 | 6.986209677 | *0.534576423* |
| ENSG00000272366 | AL158211.3 | **2.928853377** | 0.854066242 | 6.576465517 | *1.96587091* |
| ENSG00000274330 | AL160191.3 | **4.278387852** | 0.031322661 | 1.034714622 | *0.029603296* |
| ENSG00000224842 | AL161908.1 | **1.63943574** | 6.692244281 | 14.36259586 | *13.37459897* |
| ENSG00000260475 | AL353719.1 | **2.629208834** | 1.287165005 | 5.723688373 | *2.206318018* |
| ENSG00000224764 | AL353768.1 | **1.4758089** | 2.328518787 | 11.64425804 | *2.392640083* |
| ENSG00000225339 | AL354740.1 | **-1.136011582** | 1.345901497 | 2.604824978 | *1.284588205* |
| ENSG00000225811 | AL354771.1 | **4.312409756** | 0 | 2.589336189 | *0.304233558* |
| ENSG00000278177 | AL354811.1 | **1.144599164** | 17.33037231 | 25.43145993 | *14.62540758* |
| ENSG00000258793 | AL355102.4 | **-1.834029918** | 5.953958637 | 1.772511936 | *8.265578136* |
| ENSG00000271551 | AL355297.3 | **1.320368715** | 31.75810432 | 63.32292198 | *32.90781643* |
| ENSG00000273221 | AL355816.2 | **-1.041808092** | 10.70645289 | 4.299088849 | *9.857778847* |
| ENSG00000228395 | AL356481.1 | **-1.018219142** | 9.531952986 | 30.70298608 | *6.818227704* |
| ENSG00000226334 | AL359182.1 | **2.692995303** | 0.861874115 | 4.60402948 | *0.781017493* |
| ENSG00000258957 | AL359317.2 | **-1.286763423** | 4.094106702 | 2.634709259 | *3.525055002* |
| ENSG00000259834 | AL365361.1 | **1.719054697** | 26.59116032 | 59.3176558 | *25.08432078* |
| ENSG00000237976 | AL391069.2 | **2.82027117** | 0.788721385 | 4.585881377 | *1.476098106* |
| ENSG00000284418 | AL391117.1 | **3.01935425** | 0.227216953 | 30.20566415 | *0.294195227* |
| ENSG00000225905 | AL391244.1 | **1.279241121** | 9.902013441 | 16.35966337 | *9.917749839* |
| ENSG00000272842 | AL391834.1 | **-1.459937499** | 11.13326267 | 3.679869105 | *8.856193343* |
| ENSG00000271387 | AL445228.3 | **1.45845869** | 2.262438126 | 13.54549809 | *3.317120193* |
| ENSG00000272320 | AL445309.1 | **1.680967676** | 0.415430518 | 1.88037623 | *0.249738223* |
| ENSG00000273062 | AL449106.1 | **1.958402306** | 3.694804589 | 10.62418267 | *4.677051049* |
| ENSG00000232590 | AL451127.1 | **3.664620406** | 1.234550391 | 11.31960602 | *0.924108677* |
| ENSG00000224984 | AL512363.1 | **1.1220026** | 3.568896342 | 9.28405486 | *2.776760608* |
| ENSG00000260063 | AL512408.1 | **1.269990201** | 3.150629601 | 10.05026698 | *2.381797681* |
| ENSG00000229422 | AL512625.2 | **1.037186836** | 45.62335718 | 64.3350021 | *42.89268412* |
| ENSG00000269621 | AL589765.7 | **-1.341455345** | 24.51949932 | 8.404115183 | *24.72738106* |
| ENSG00000216809 | AL589993.1 | **2.02201865** | 0.357452243 | 32.13288018 | *0.756000708* |
| ENSG00000259357 | AL590133.2 | **-1.693616613** | 5.947057857 | 4.404744548 | *6.920499903* |
| ENSG00000279140 | AL590326.1 | **1.672122764** | 17.41127203 | 34.39423139 | *20.61563179* |
| ENSG00000233926 | AL591368.1 | **1.557964465** | 0.730038251 | 1.73353911 | *0.745268381* |
| ENSG00000233765 | AL591479.1 | **1.308849821** | 2.406339022 | 24.84416181 | *4.807441132* |
| ENSG00000233875 | AL592078.1 | **-1.831734193** | 9.616312583 | 2.61238781 | *6.083148071* |

| | | | | | |
|---|---|---|---|---|---|
| ENSG00000225643 | AL606491.1 | **4.051751299** | 0.15041443 | 3.804560366 | *0.632926649* |
| ENSG00000237372 | AL606807.1 | **-1.606194136** | 5.095803078 | 1.972946697 | *3.112747192* |
| ENSG00000235121 | AL645504.1 | **-1.830809725** | 1.383992053 | 1.068097159 | *1.205164125* |
| ENSG00000225173 | AL662890.1 | **4.497132974** | 0 | 1.342800192 | *0.083168224* |
| ENSG00000227766 | AL671277.1 | **1.500289837** | 4.967754368 | 8.687088482 | *5.237763101* |
| ENSG00000225936 | AL731557.1 | **4.109425167** | 0 | 1.043896516 | *0.077068426* |
| ENSG00000276742 | AL731566.3 | **1.469601132** | 4.730634736 | 12.83553023 | *5.76051931* |
| ENSG00000136010 | ALDH1L2 | **2.035628924** | 0.286557809 | 1.017394539 | *0.23825859* |
| ENSG00000108602 | ALDH3A1 | **4.100538865** | 0 | 5.657894179 | *0.012089553* |
| ENSG00000112294 | ALDH5A1 | **1.172591497** | 13.08625724 | 19.92988217 | *14.98641665* |
| ENSG00000119711 | ALDH6A1 | **-1.375221593** | 12.82328844 | 3.266989926 | *12.52328658* |
| ENSG00000171094 | ALK | **2.87337802** | 0.263213429 | 10.76498068 | *0.318268277* |
| ENSG00000073331 | ALPK1 | **1.537236826** | 0.027991583 | 1.66553712 | *0.040373622* |
| ENSG00000162551 | ALPL | **1.24100511** | 0.522546887 | 1.474202059 | *0.371958784* |
| ENSG00000139211 | AMIGO2 | **1.349684293** | 27.20982328 | 44.68747837 | *35.93228153* |
| ENSG00000166025 | AMOTL1 | **1.991146616** | 2.732958468 | 7.583593642 | *5.241805127* |
| ENSG00000114019 | AMOTL2 | **-4.234734182** | 0.125913804 | 23.71525205 | *0.033545237* |
| ENSG00000078053 | AMPH | **2.282446746** | 3.051899934 | 13.11520371 | *4.512744624* |
| ENSG00000091879 | ANGPT2 | **3.360519721** | 0.010533744 | 0.070997374 | *0.015174937* |
| ENSG00000029534 | ANK1 | **1.133729563** | 2.303442416 | 8.307356753 | *4.190795886* |
| ENSG00000180071 | ANKRD18A | **1.387575524** | 1.802338011 | 3.207461064 | *3.087395989* |
| ENSG00000230453 | ANKRD18B | **1.628964026** | 1.086009419 | 3.942746239 | *1.627622089* |
| ENSG00000172014 | ANKRD20A4P | **1.695064282** | 0.125005635 | 1.753764276 | *0.219815468* |
| ENSG00000180777 | ANKRD30B | **4.600757854** | 0 | 1.162954493 | *0.022735838* |
| ENSG00000167612 | ANKRD33 | **1.478785083** | 0.557011015 | 1.164215136 | *1.160055927* |
| ENSG00000065413 | ANKRD44 | **1.12975275** | 1.962210727 | 2.886861622 | *1.797328742* |
| ENSG00000241859 | ANOS2P | **1.359320036** | 4.825582954 | 8.909288203 | *7.123225057* |
| ENSG00000166825 | ANPEP | **1.021708076** | 1.505200068 | 3.273148818 | *2.17402722* |
| ENSG00000169604 | ANTXR1 | **3.691837863** | 0.168820402 | 5.281727095 | *0.892249262* |
| ENSG00000163297 | ANTXR2 | **5.49642528** | 0.043250567 | 2.455547728 | *0.038550423* |
| ENSG00000104537 | ANXA13 | **4.407664921** | 0 | 0.769611411 | *0.188797427* |
| ENSG00000260583 | AP000223.1 | **-1.649599155** | 5.559497869 | 1.67673689 | *5.143077974* |
| ENSG00000227757 | AP000282.1 | **2.353785023** | 1.727602275 | 5.689683944 | *2.24445265* |
| ENSG00000227755 | AP000344.1 | **3.368251697** | 0.09458393 | 1.346135638 | *0.513370778* |
| ENSG00000254528 | AP000757.1 | **-1.932570944** | 11.7933653 | 5.030759125 | *14.47580931* |
| ENSG00000261098 | AP000766.1 | **-1.200397652** | 4.412838966 | 1.602276367 | *3.099258167* |
| ENSG00000245552 | AP000787.1 | **1.004946887** | 0.888598946 | 2.122612601 | *0.84937214* |
| ENSG00000266456 | AP001178.3 | **-1.84367417** | 1.201700407 | 0.862036087 | *1.10928047* |
| ENSG00000248027 | AP001351.1 | **4.176159557** | 0.072246463 | 1.595416293 | *0.0665857* |
| ENSG00000235023 | AP001626.1 | **2.312594666** | 2.090414879 | 6.950444418 | *1.987676568* |
| ENSG00000279117 | AP001972.5 | **-2.15630178** | 22.74234203 | 3.864537718 | *22.34147506* |
| ENSG00000250230 | AP002754.1 | **-1.293462143** | 2.555490807 | 3.968047821 | *4.249256832* |
| ENSG00000270179 | AP002840.2 | **-1.570808455** | 3.997560019 | 2.541942823 | *4.265062317* |
| ENSG00000255301 | AP002893.1 | **1.633806431** | 1.733765794 | 3.969579967 | *2.728510998* |
| ENSG00000279878 | AP003108.5 | **-1.113662231** | 15.42150043 | 5.813568432 | *16.10115731* |
| ENSG00000261578 | AP003119.3 | **1.620644627** | 2.302940959 | 6.281521319 | *2.514549422* |
| ENSG00000256196 | AP003721.1 | **-1.039305698** | 176.2877584 | 58.16445004 | *137.3233695* |

| | | | | | |
|---|---|---|---|---|---|
| ENSG00000279711 | AP004782.1 | **1.434890345** | 0.362574219 | 3.225609523 | *0.447935542* |
| ENSG00000279042 | AP005242.4 | **1.439862486** | 0.917257764 | 29.66927897 | *1.580356667* |
| ENSG00000081014 | AP4E1 | **-1.146055796** | 11.31956288 | 3.650283754 | *9.570572431* |
| ENSG00000034053 | APBA2 | **1.028820009** | 0.233400743 | 1.239356212 | *0.348797793* |
| ENSG00000115266 | APC2 | **1.292058974** | 0.110813329 | 1.853697926 | *0.173534868* |
| ENSG00000171388 | APLN | **-1.460743997** | 2.898689328 | 0.942028787 | *4.271096965* |
| ENSG00000118137 | APOA1 | **1.481863488** | 0.631943356 | 1.278960882 | *1.073142933* |
| ENSG00000124701 | APOBEC2 | **-1.873504096** | 3.619485838 | 1.099953316 | *3.848933357* |
| ENSG00000130208 | APOC1 | **1.896757524** | 0.284501427 | 1.135629809 | *0.636664295* |
| ENSG00000143595 | AQP10 | **1.879016067** | 0.834247352 | 2.167383959 | *0.81576637* |
| ENSG00000165272 | AQP3 | **1.67429341** | 12.38012935 | 26.35054063 | *11.79645311* |
| ENSG00000047365 | ARAP2 | **1.991499157** | 0.626417429 | 2.604164495 | *0.795871908* |
| ENSG00000198576 | ARC | **2.906240812** | 0.181436408 | 13.98869523 | *0.143768324* |
| ENSG00000137727 | ARHGAP20 | **2.602769409** | 0.830957091 | 4.182594969 | *0.732842674* |
| ENSG00000275832 | ARHGAP23 | **1.937893468** | 0.099052476 | 0.636349667 | *0.110567081* |
| ENSG00000165895 | ARHGAP42 | **3.185445861** | 0.165923612 | 2.640977618 | *0.240274712* |
| ENSG00000180448 | ARHGAP45 | **4.019459457** | 0.001894459 | 0.16392241 | *0.008536242* |
| ENSG00000258655 | ARHGAP5-AS1 | **-1.04250321** | 45.79925603 | 15.15467309 | *36.03653278* |
| ENSG00000047648 | ARHGAP6 | **2.340291948** | 0.292429736 | 0.983666928 | *0.41431349* |
| ENSG00000123329 | ARHGAP9 | **3.579392319** | 0.027028431 | 7.548091804 | *0.012346622* |
| ENSG00000111348 | ARHGDIB | **2.318932394** | 0.120819629 | 0.86104757 | *0.182584044* |
| ENSG00000214694 | ARHGEF33 | **-1.018566178** | 2.42655644 | 0.865192604 | *2.414341214* |
| ENSG00000129675 | ARHGEF6 | **3.622821308** | 0.021774932 | 0.411515615 | *0.015558082* |
| ENSG00000196843 | ARID5A | **1.035275972** | 4.140021793 | 5.880226509 | *6.253398807* |
| ENSG00000170540 | ARL6IP1 | **-1.321800611** | 182.6695575 | 47.36753621 | *195.1433045* |
| ENSG00000172995 | ARPP21 | **2.278384816** | 0.01562538 | 2.94850235 | *0.018366816* |
| ENSG00000137486 | ARRB1 | **-1.875244612** | 16.91179951 | 27.85959156 | *15.85437903* |
| ENSG00000105643 | ARRDC2 | **1.731947393** | 0.324007526 | 0.722821824 | *0.212030617* |
| ENSG00000113369 | ARRDC3 | **1.069028176** | 9.104017663 | 13.69272366 | *5.276157475* |
| ENSG00000141337 | ARSG | **1.470156365** | 0.21976967 | 0.803103746 | *0.278640742* |
| ENSG00000180801 | ARSJ | **4.090183072** | 0.16147323 | 8.582655179 | *0.123035271* |
| ENSG00000157399 | ARSL | **1.289428987** | 9.3947268 | 15.10679863 | *14.54066066* |
| ENSG00000224060 | ARSLP1 | **7.328460296** | 0.060849931 | 7.339983698 | *0* |
| ENSG00000167311 | ART5 | **4.681908278** | 0.013275072 | 7.914577151 | *0.103561741* |
| ENSG00000102048 | ASB9 | **-1.052522781** | 12.60058405 | 7.902799213 | *11.67499749* |
| ENSG00000139352 | ASCL1 | **4.086504685** | 3.929105984 | 42.70080571 | *4.931854648* |
| ENSG00000183734 | ASCL2 | **-1.314087264** | 281.263017 | 77.10533849 | *318.9720522* |
| ENSG00000141505 | ASGR1 | **1.225363283** | 0.442324109 | 24.14674673 | *0.553828067* |
| ENSG00000161944 | ASGR2 | **2.75312857** | 0.071157637 | 36.75276998 | *0.235757926* |
| ENSG00000248498 | ASNSP1 | **2.018551439** | 0.892072022 | 2.624112509 | *0.600770322* |
| ENSG00000148219 | ASTN2 | **1.410364327** | 1.967378482 | 3.581884338 | *1.881549054* |
| ENSG00000226864 | ATE1-AS1 | **1.435368958** | 6.46310803 | 12.931751 | *6.714576494* |
| ENSG00000162772 | ATF3 | **1.211368976** | 0.980800509 | 3.175326197 | *0.666554278* |
| ENSG00000169136 | ATF5 | **1.137048789** | 43.35227883 | 68.15936994 | *43.03074045* |
| ENSG00000152348 | ATG10 | **-1.093621013** | 8.508771136 | 3.535329068 | *7.943427781* |
| ENSG00000172238 | ATOH1 | **4.327929547** | 0 | 31.88984108 | *0* |
| ENSG00000179774 | ATOH7 | **1.629513982** | 5.378377994 | 11.75624496 | *7.452421414* |

| | | | | | |
|---|---|---|---|---|---|
| ENSG00000129244 | ATP1B2 | **2.852398273** | 4.382804997 | 21.07326641 | *4.286684414* |
| ENSG00000069849 | ATP1B3 | **-1.424602694** | 97.31391331 | 29.47159042 | *126.8379382* |
| ENSG00000157087 | ATP2B2 | **1.270436471** | 0.019704624 | 10.17791305 | *0.040400298* |
| ENSG00000081923 | ATP8B1 | **1.324193015** | 0.752875635 | 13.39922018 | *1.275022999* |
| ENSG00000143515 | ATP8B2 | **-1.37913469** | 199.4216876 | 50.31023728 | *199.407481* |
| ENSG00000104043 | ATP8B4 | **1.16478194** | 0.135855592 | 0.861352052 | *0.192948755* |
| ENSG00000183778 | B3GALT5 | **1.982390376** | 0.261861735 | 26.12090905 | *0.369487004* |
| ENSG00000184809 | B3GALT5-AS1 | **4.024510747** | 0.015580903 | 0.248823051 | *0.056229625* |
| ENSG00000187676 | B3GLCT | **1.048527544** | 6.541303295 | 8.488210874 | *8.271535569* |
| ENSG00000176383 | B3GNT4 | **1.2878069** | 0.860767355 | 1.477180133 | *0.827757853* |
| ENSG00000139044 | B4GALNT3 | **1.408467555** | 11.77604153 | 25.99297758 | *16.44404091* |
| ENSG00000233554 | B4GALT1-AS1 | **-1.909614511** | 1.518609763 | 1.062625512 | *1.241423233* |
| ENSG00000164929 | BAALC | **2.835253445** | 15.36113358 | 80.67288014 | *13.88902575* |
| ENSG00000247081 | BAALC-AS1 | **2.509272403** | 0.391579595 | 1.608042502 | *0.621949721* |
| ENSG00000236939 | BAALC-AS2 | **5.231681377** | 0.508927808 | 50.08418891 | *0.974350339* |
| ENSG00000007516 | BAIAP3 | **2.654620917** | 0.756721314 | 3.922236918 | *0.756658035* |
| ENSG00000095739 | BAMBI | **3.025593717** | 2.260061982 | 18.80385819 | *1.739541897* |
| ENSG00000153064 | BANK1 | **1.422490297** | 0.187224741 | 1.841632673 | *0.309389365* |
| ENSG00000179941 | BBS10 | **-1.16433599** | 87.03418633 | 31.33853342 | *74.09997445* |
| ENSG00000187244 | BCAM | **-1.148492483** | 384.1389165 | 116.7997511 | *430.2322841* |
| ENSG00000060982 | BCAT1 | **3.920971334** | 0.013451229 | 13.6496505 | *0.033604792* |
| ENSG00000119866 | BCL11A | **1.45282394** | 0.676266661 | 55.60239403 | *0.773628728* |
| ENSG00000171791 | BCL2 | **3.093944736** | 0.234530524 | 1.318194196 | *0.243816828* |
| ENSG00000126453 | BCL2L12 | **1.612095381** | 1.383654508 | 3.378031157 | *1.733642353* |
| ENSG00000176697 | BDNF | **2.972001064** | 0.046455833 | 0.957753344 | *0.089943609* |
| ENSG00000245573 | BDNF-AS | **-1.189535583** | 1.443250873 | 1.781280782 | *1.14085982* |
| ENSG00000188848 | BEND4 | **1.298540004** | 0.196293113 | 0.438783431 | *0.187044634* |
| ENSG00000151917 | BEND6 | **2.013865755** | 0.114726065 | 0.50430266 | *0.057300214* |
| ENSG00000184515 | BEX5 | **1.245124795** | 4.22494377 | 6.851778316 | *6.144488151* |
| ENSG00000182492 | BGN | **5.097592435** | 0.016936238 | 0.491786186 | *0.030961582* |
| ENSG00000180535 | BHLHA15 | **1.328046637** | 6.241030675 | 13.44566761 | *6.612026344* |
| ENSG00000180828 | BHLHE22 | **3.814072799** | 1.10597878 | 10.49383432 | *2.081110634* |
| ENSG00000134107 | BHLHE40 | **3.021034265** | 3.030430906 | 21.5995741 | *5.40539948* |
| ENSG00000123095 | BHLHE41 | **2.989828294** | 0.059054195 | 5.420293895 | *0.063043596* |
| ENSG00000132840 | BHMT2 | **1.141836091** | 0.894435369 | 9.228433965 | *0.834977944* |
| ENSG00000023445 | BIRC3 | **-6.165082507** | 10.42018447 | 0.168160085 | *0.144706032* |
| ENSG00000125845 | BMP2 | **3.329680761** | 5.688195534 | 37.02159049 | *5.184149415* |
| ENSG00000152785 | BMP3 | **3.222837975** | 0.092853615 | 0.6707181 | *0.083597429* |
| ENSG00000125378 | BMP4 | **1.485262484** | 4.333331281 | 28.15389228 | *4.625646431* |
| ENSG00000153162 | BMP6 | **2.569607213** | 0.74862983 | 3.1341707 | *0.881970634* |
| ENSG00000164619 | BMPER | **4.450753566** | 0.023898343 | 4.247993207 | *0.020197217* |
| ENSG00000152430 | BOLL | **2.922475237** | 0.044874297 | 1.734120109 | *0.099081664* |
| ENSG00000185515 | BRCC3 | **-1.115700994** | 21.36982799 | 6.569908614 | *20.85718319* |
| ENSG00000162670 | BRINP3 | **1.21832453** | 105.5275936 | 161.0174272 | *121.7938588* |
| ENSG00000145741 | BTF3 | **-1.414052487** | 230.3476356 | 60.89859196 | *254.9671821* |
| ENSG00000124557 | BTN1A1 | **7.344540845** | 0 | 82.02153908 | *0.012636323* |
| ENSG00000124508 | BTN2A2 | **3.092400598** | 0.886896168 | 32.73570673 | *1.187720565* |

| | | | | | |
|---|---|---|---|---|---|
| ENSG00000124549 | BTN2A3P | **2.245841416** | 0.632953519 | 2.386096213 | *0.962952471* |
| ENSG00000186470 | BTN3A2 | **1.138538881** | 0.395430137 | 3.127825735 | *0.444723217* |
| ENSG00000111801 | BTN3A3 | **1.186751577** | 0.139934032 | 1.213712014 | *0.191832248* |
| ENSG00000240240 | BX664727.3 | **1.044167001** | 0.799483291 | 1.478376685 | *1.015577177* |
| ENSG00000154493 | C10orf90 | **1.394026315** | 0.109834298 | 0.300888117 | *0.17136714* |
| ENSG00000137720 | C11orf1 | **-1.013911711** | 11.70403388 | 4.19655137 | *10.21244111* |
| ENSG00000150750 | C11orf53 | **5.646984417** | 0.025927969 | 0.837146976 | *0* |
| ENSG00000187479 | C11orf96 | **6.326005458** | 0 | 2.639924269 | *0* |
| ENSG00000186073 | C15orf41 | **-1.449544289** | 1.622465274 | 0.940156137 | *1.466130057* |
| ENSG00000166920 | C15orf48 | **1.398881613** | 1.420106394 | 2.923615829 | *1.759440065* |
| ENSG00000173372 | C1QA | **-1.942406831** | 0.962425301 | 1.980128829 | *1.425817142* |
| ENSG00000165985 | C1QL3 | **-1.60254728** | 48.24229814 | 86.06002979 | *55.62528669* |
| ENSG00000182795 | C1orf116 | **2.200493939** | 104.4490393 | 18.68900863 | *80.77757989* |
| ENSG00000175262 | C1orf127 | **1.455095469** | 0.243654611 | 45.3217184 | *0.355046157* |
| ENSG00000221953 | C1orf229 | **1.450991605** | 0.246427652 | 9.919992167 | *0.134666169* |
| ENSG00000166278 | C2 | **1.843684143** | 0.042082768 | 0.110691664 | *0.036707541* |
| ENSG00000149609 | C20orf144 | **1.414355452** | 0.916539537 | 1.659978883 | *1.409578234* |
| ENSG00000157617 | C2CD2 | **-1.082260159** | 0.452476701 | 0.205873451 | *0.543924083* |
| ENSG00000225556 | C2CD4D | **2.222023221** | 0.859517301 | 3.300490596 | *1.957540725* |
| ENSG00000234614 | C2CD4D-AS1 | **1.404251798** | 22.24923069 | 39.76946086 | *17.55614408* |
| ENSG00000187699 | C2orf88 | **1.852946383** | 0.725097658 | 3.406872558 | *1.051276293* |
| ENSG00000187068 | C3orf70 | **1.194787951** | 33.81125617 | 69.40498208 | *31.97524195* |
| ENSG00000106804 | C5 | **2.144484974** | 21.12557796 | 61.68051646 | *24.61624298* |
| ENSG00000197261 | C6orf141 | **1.451743295** | 0.172632623 | 25.82655737 | *0.437314342* |
| ENSG00000112936 | C7 | **4.666756125** | 0.974141153 | 47.17558104 | *0.995773139* |
| ENSG00000196366 | C9orf163 | **1.429020985** | 1.62977652 | 2.780844925 | *1.545823527* |
| ENSG00000135045 | C9orf40 | **1.65545516** | 12.35874563 | 35.28132279 | *17.16300356* |
| ENSG00000063180 | CA11 | **1.025215957** | 10.57864674 | 16.17349123 | *11.22412206* |
| ENSG00000074410 | CA12 | **2.114514452** | 0.055787422 | 11.79924816 | *0.169816557* |
| ENSG00000169239 | CA5B | **-1.519820498** | 3.235065111 | 7.569708466 | *2.552520277* |
| ENSG00000134508 | CABLES1 | **-1.045478179** | 5.540483361 | 1.770072998 | *4.786181815* |
| ENSG00000175544 | CABP4 | **2.098376055** | 0.150000864 | 0.749636338 | *0.190170053* |
| ENSG00000158966 | CACHD1 | **-1.322650194** | 5.239142597 | 2.36616597 | *5.038349738* |
| ENSG00000141837 | CACNA1A | **1.083546971** | 0.737210085 | 1.281825001 | *1.161056531* |
| ENSG00000198216 | CACNA1E | **-1.096929958** | 1.848819454 | 1.220939319 | *1.595704591* |
| ENSG00000157445 | CACNA2D3 | **-1.686618391** | 0.165174443 | 0.513254189 | *0.103674109* |
| ENSG00000175161 | CADM2 | **2.341818394** | 0.1622738 | 0.793562203 | *0.217039652* |
| ENSG00000162706 | CADM3 | **2.96150332** | 0.507987817 | 2.712480389 | *1.404142173* |
| ENSG00000270419 | CAHM | **1.708516439** | 3.240043714 | 6.745606872 | *4.509453726* |
| ENSG00000110680 | CALCA | **4.776740443** | 0 | 1.294219596 | *0* |
| ENSG00000175868 | CALCB | **2.588471584** | 0.354366869 | 5.342387555 | *0.299989193* |
| ENSG00000004948 | CALCR | **4.68277657** | 0.058801011 | 1.045595897 | *0.068603069* |
| ENSG00000064989 | CALCRL | **3.961850068** | 0.042825897 | 1.211331029 | *0.042146705* |
| ENSG00000183128 | CALHM3 | **2.835542422** | 0.171493937 | 1.189317039 | *0.438533656* |
| ENSG00000183049 | CAMK1D | **1.415629807** | 0.299564879 | 0.746715763 | *0.774141233* |
| ENSG00000145349 | CAMK2D | **-1.31907631** | 22.90462024 | 6.351019054 | *23.11589105* |
| ENSG00000163888 | CAMK2N2 | **1.3355216** | 20.03815467 | 32.8387734 | *41.48105803* |

| | | | | | |
|---|---|---|---|---|---|
| ENSG00000112186 | CAP2 | **1.238035994** | 0.700190537 | 4.282930554 | *0.892366063* |
| ENSG00000042493 | CAPG | **1.733314666** | 0.73844479 | 18.97276539 | *1.076110658* |
| ENSG00000164305 | CASP3 | **-1.393837007** | 17.89823606 | 4.974469578 | *19.08217959* |
| ENSG00000196954 | CASP4 | **1.062680446** | 0.301564423 | 1.361378875 | *0.214923355* |
| ENSG00000165806 | CASP7 | **1.559338874** | 0.432675819 | 3.2064958 | *0.519212816* |
| ENSG00000036828 | CASR | **-1.193575497** | 77.24792621 | 22.18084902 | *79.86959777* |
| ENSG00000160200 | CBS | **2.268986136** | 0.363409545 | 1.499155631 | *0.483616806* |
| ENSG00000274276 | CBSL | **2.076240268** | 0.690576007 | 13.17920319 | *0.794817834* |
| ENSG00000183287 | CCBE1 | **5.025898718** | 0.099978073 | 2.745732186 | *0.406446204* |
| ENSG00000105479 | CCDC114 | **4.366067372** | 0.007576407 | 1.128061834 | *0.079066123* |
| ENSG00000128596 | CCDC136 | **1.271758567** | 0.487295195 | 1.858477992 | *0.662089155* |
| ENSG00000144395 | CCDC150 | **2.227740812** | 1.521563588 | 4.763491597 | *3.092579189* |
| ENSG00000120262 | CCDC170 | **2.636660514** | 0.088275623 | 0.773031321 | *0.128952701* |
| ENSG00000228544 | CCDC183-AS1 | **1.043322134** | 8.952286789 | 14.63903284 | *12.94755765* |
| ENSG00000260220 | CCDC187 | **-1.210239694** | 0.234239888 | 0.223679837 | *0.456787516* |
| ENSG00000140481 | CCDC33 | **-1.211570354** | 5.544778639 | 7.484138918 | *4.425173318* |
| ENSG00000165972 | CCDC38 | **1.503033143** | 0.167225193 | 0.316599542 | *0.144551188* |
| ENSG00000253276 | CCDC71L | **1.901394659** | 5.862488976 | 15.01168065 | *7.280610452* |
| ENSG00000205476 | CCDC85C | **-1.965263517** | 28.24964187 | 4.884120215 | *32.98705461* |
| ENSG00000115009 | CCL20 | **-22.46188546** | 2.967713822 | 7.343181951 | *0* |
| ENSG00000133101 | CCNA1 | **-2.73439882** | 1.025301942 | 2.569709761 | *1.005051503* |
| ENSG00000135083 | CCNJL | **4.645248129** | 0.002195864 | 0.038633037 | *0.003564532* |
| ENSG00000090061 | CCNK | **-1.024535837** | 34.91784202 | 11.36961351 | *33.14840658* |
| ENSG00000103540 | CCP110 | **-1.166058125** | 22.07719562 | 6.599604539 | *20.10469188* |
| ENSG00000184451 | CCR10 | **1.011311159** | 2.518593142 | 8.980514636 | *2.025956012* |
| ENSG00000170458 | CD14 | **3.233384988** | 0.126727441 | 3.894589721 | *0.333533899* |
| ENSG00000120217 | CD274 | **1.410785839** | 0.425872051 | 2.393159952 | *0.472976911* |
| ENSG00000135218 | CD36 | **2.147806558** | 0.101491029 | 0.703961965 | *0.071419034* |
| ENSG00000254126 | CD8B2 | **-3.02296494** | 0.223603395 | 0.390578779 | *0.423744297* |
| ENSG00000010278 | CD9 | **-1.267359139** | 0.283268436 | 0.221960227 | *0.457599881* |
| ENSG00000040731 | CDH10 | **1.929471636** | 2.739272431 | 6.647382927 | *3.537074817* |
| ENSG00000166589 | CDH16 | **-1.30057843** | 4.21670042 | 1.07229992 | *9.484052678* |
| ENSG00000107736 | CDH23 | **1.281327952** | 1.980872269 | 6.788481623 | *3.938360004* |
| ENSG00000081138 | CDH7 | **1.868432549** | 1.931786299 | 5.210048397 | *2.313704027* |
| ENSG00000138395 | CDK15 | **4.323481085** | 0.010227377 | 1.742445017 | *0.020637995* |
| ENSG00000124762 | CDKN1A | **2.387539678** | 133.6668482 | 465.7650914 | *64.92046721* |
| ENSG00000172216 | CEBPB | **1.390280702** | 99.68336012 | 168.5043716 | *74.83799051* |
| ENSG00000277449 | CEBPB-AS1 | **1.531189687** | 1.948632122 | 231.4856529 | *0.748718543* |
| ENSG00000099954 | CECR2 | **1.450821983** | 8.623698615 | 105.4478495 | *7.736720693* |
| ENSG00000241832 | CECR3 | **4.94593679** | 0 | 2.39222146 | *0.034055384* |
| ENSG00000219073 | CELA3B | **4.366038018** | 0.026418244 | 8.13597148 | *0.023714931* |
| ENSG00000161082 | CELF5 | **1.174376162** | 0.450781264 | 0.78528962 | *0.305754595* |
| ENSG00000205923 | CEMP1 | **-2.015140561** | 1.389904164 | 0.452807903 | *0.602333102* |
| ENSG00000111860 | CEP85L | **1.281256762** | 6.792415115 | 11.09500298 | *5.546027653* |
| ENSG00000143418 | CERS2 | **-1.578934107** | 228.2029019 | 49.514523 | *228.7495559* |
| ENSG00000227617 | CERS6-AS1 | **-1.000803939** | 0.879722618 | 5.810782508 | *0.742288621* |
| ENSG00000171811 | CFAP46 | **-1.080540144** | 0.552655097 | 26.14697614 | *0.740785018* |

| | | | | | |
|---|---|---|---|---|---|
| ENSG00000231233 | CFAP58-DT | **1.386133382** | 2.373901163 | 4.185022654 | *1.808532484* |
| ENSG00000089101 | CFAP61 | **-1.115067223** | 0.226966556 | 0.139145629 | *0.196911476* |
| ENSG00000188523 | CFAP77 | **2.313271475** | 0.143460892 | 2.518351944 | *0.252153306* |
| ENSG00000143375 | CGN | **1.057143553** | 15.43210149 | 20.84464903 | *20.26426912* |
| ENSG00000128965 | CHAC1 | **1.460288776** | 37.44371846 | 67.29264433 | *17.8719427* |
| ENSG00000170004 | CHD3 | **-1.482326995** | 17.33030008 | 14.96538819 | *13.56497399* |
| ENSG00000016391 | CHDH | **-1.174041945** | 16.81426459 | 39.53938359 | *15.27637554* |
| ENSG00000106069 | CHN2 | **1.848094927** | 0.138758623 | 2.375911849 | *0.084554487* |
| ENSG00000080644 | CHRNA3 | **1.174081103** | 27.2776433 | 42.98958177 | *28.95545051* |
| ENSG00000175344 | CHRNA7 | **1.912154869** | 0.010001891 | 0.185147832 | *0.018120855* |
| ENSG00000160716 | CHRNB2 | **-1.242086428** | 161.9376073 | 65.76748964 | *151.9094644* |
| ENSG00000135902 | CHRND | **1.739846767** | 0.209845058 | 0.489766917 | *0.430300089* |
| ENSG00000108556 | CHRNE | **1.377623609** | 0.25826294 | 22.64847806 | *0.346399037* |
| ENSG00000183196 | CHST6 | **-2.156058554** | 1.911562007 | 0.516368524 | *1.738531951* |
| ENSG00000124302 | CHST8 | **1.072779855** | 7.898019701 | 11.18587963 | *10.49632989* |
| ENSG00000258289 | CHURC1 | **-1.035085089** | 40.56478259 | 12.53440637 | *32.21303465* |
| ENSG00000136425 | CIB2 | **1.006100244** | 11.01707719 | 19.89504866 | *16.25334873* |
| ENSG00000016490 | CLCA1 | **4.128009008** | 0 | 7.198559382 | *0.008920179* |
| ENSG00000112782 | CLIC5 | **1.987119544** | 0.599073458 | 8.901133913 | *0.638442096* |
| ENSG00000106665 | CLIP2 | **-1.167079981** | 67.69741046 | 19.21191104 | *65.90970632* |
| ENSG00000165959 | CLMN | **1.153879928** | 0.483526878 | 1.603027111 | *0.477267984* |
| ENSG00000166250 | CLMP | **1.841939342** | 4.425971123 | 21.10717515 | *7.512093746* |
| ENSG00000140931 | CMTM3 | **2.49670379** | 1.441510034 | 5.662564112 | *2.171971025* |
| ENSG00000153551 | CMTM7 | **2.30102765** | 0.584751719 | 7.12719572 | *0.964187201* |
| ENSG00000132259 | CNGA4 | **-1.061816225** | 6.329890633 | 4.484744082 | *5.919893407* |
| ENSG00000119946 | CNNM1 | **-1.94863825** | 158.6078302 | 26.50151553 | *154.4769933* |
| ENSG00000122756 | CNTFR | **2.75254196** | 0.031473662 | 1.372259147 | *0.070160126* |
| ENSG00000149972 | CNTN5 | **1.048851311** | 0.06513378 | 15.04918896 | *0.05581731* |
| ENSG00000106714 | CNTNAP3 | **1.368799929** | 0.494554327 | 0.887467675 | *0.652138163* |
| ENSG00000154529 | CNTNAP3B | **1.59576215** | 0.568642984 | 1.12438297 | *0.755465304* |
| ENSG00000276386 | CNTNAP3P2 | **1.00155479** | 2.477713494 | 3.559560396 | *2.449026145* |
| ENSG00000155052 | CNTNAP5 | **2.881187989** | 0.186961999 | 1.517995032 | *0.367862713* |
| ENSG00000181924 | COA4 | **-1.39459314** | 61.29068609 | 17.24385923 | *68.59527736* |
| ENSG00000183513 | COA5 | **-1.121516504** | 58.56870297 | 17.88736847 | *60.13400103* |
| ENSG00000187955 | COL14A1 | **3.064457751** | 0.027100959 | 7.683357982 | *0.042763991* |
| ENSG00000164692 | COL1A2 | **5.179577707** | 0.082140663 | 11.13559345 | *0.174835164* |
| ENSG00000124749 | COL21A1 | **2.13199591** | 0.085163045 | 0.317698378 | *0.319802945* |
| ENSG00000196739 | COL27A1 | **1.384842732** | 10.18630527 | 18.79897132 | *18.91211322* |
| ENSG00000139219 | COL2A1 | **5.274356494** | 0 | 0.189954767 | *0.010128563* |
| ENSG00000130635 | COL5A1 | **1.019388136** | 5.067719256 | 15.39872139 | *7.622672837* |
| ENSG00000196167 | COLCA1 | **1.702645084** | 0.444928205 | 0.977536807 | *0.727929591* |
| ENSG00000198756 | COLGALT2 | **2.520035474** | 0.553845492 | 5.374101369 | *0.942263454* |
| ENSG00000188243 | COMMD6 | **-1.086632498** | 44.59831442 | 14.27192895 | *46.52653636* |
| ENSG00000110880 | CORO1C | **-1.401430977** | 31.95537283 | 8.937209095 | *36.35270036* |
| ENSG00000103647 | CORO2B | **4.443638469** | 0.029462967 | 7.341448647 | *0.025112798* |
| ENSG00000124772 | CPNE5 | **1.529733373** | 0.317260412 | 4.636079795 | *0.628722545* |
| ENSG00000103381 | CPPED1 | **-1.114460885** | 23.52955201 | 7.451766293 | *21.15196444* |

| | | | | | |
|---|---|---|---|---|---|
| ENSG00000088882 | CPXM1 | **3.124570855** | 0.144299085 | 1.179029924 | *0.079213045* |
| ENSG00000270533 | CR382285.1 | **1.274541566** | 0.726622284 | 4.598510728 | *0.858816072* |
| ENSG00000284471 | CR769775.2 | **4.410399148** | 0.016671617 | 0.555531368 | *0.04195315* |
| ENSG00000177685 | CRACR2B | **1.185986029** | 0.106602466 | 0.778943281 | *0.199767427* |
| ENSG00000157613 | CREB3L1 | **1.47081722** | 0.884698461 | 1.762222924 | *0.526682709* |
| ENSG00000147571 | CRH | **4.568853328** | 0.831209968 | 12.72255523 | *1.181522719* |
| ENSG00000146215 | CRIP3 | **2.645146826** | 0.071264597 | 1.126910316 | *0.25667557* |
| ENSG00000214960 | CRPPA | **1.777125657** | 0.103331043 | 6.770348337 | *0.157878196* |
| ENSG00000095713 | CRTAC1 | **1.636159879** | 0.399051706 | 0.886165618 | *1.323316609* |
| ENSG00000170275 | CRTAP | **1.674801416** | 11.47920123 | 23.9631502 | *16.38022862* |
| ENSG00000172346 | CSDC2 | **1.697517877** | 0.194680092 | 0.841879632 | *0.238103381* |
| ENSG00000164796 | CSMD3 | **4.910088091** | 0.003407489 | 12.34986649 | *0.006698774* |
| ENSG00000175183 | CSRP2 | **1.016586862** | 13.45145706 | 17.66407613 | *18.21697057* |
| ENSG00000175315 | CST6 | **1.099607783** | 3.384217052 | 4.96265723 | *4.464518565* |
| ENSG00000234215 | CT66 | **1.449021568** | 2.951816054 | 14.08244992 | *2.716980315* |
| ENSG00000175215 | CTDSP2 | **-1.154156611** | 120.7099504 | 37.49237027 | *102.6351732* |
| ENSG00000164932 | CTHRC1 | **3.109138738** | 3.178020111 | 21.60431013 | *4.722365539* |
| ENSG00000064601 | CTSA | **1.03712268** | 127.5419545 | 190.0137959 | *147.5607324* |
| ENSG00000164733 | CTSB | **-1.186744715** | 12.88629591 | 12.69879497 | *14.69912024* |
| ENSG00000109861 | CTSC | **1.537428439** | 2.159879351 | 90.67840756 | *2.965148429* |
| ENSG00000135047 | CTSL | **1.029261125** | 57.17879337 | 78.34785483 | *51.27729384* |
| ENSG00000280913 | CTSLP3 | **-1.958295062** | 2.785673068 | 2.536609647 | *2.115985914* |
| ENSG00000233932 | CTXN2 | **1.058870843** | 0.976214864 | 39.86491114 | *1.39728557* |
| ENSG00000259417 | CTXND1 | **6.706435201** | 0.010544292 | 1.097934728 | *0.025438527* |
| ENSG00000154639 | CXADR | **-1.33423616** | 41.01249655 | 11.45724726 | *34.16833744* |
| ENSG00000163739 | CXCL1 | **-6.27036975** | 232.8164847 | 2.264359039 | *1.46196609* |
| ENSG00000163734 | CXCL3 | **-4.77786535** | 10.32456113 | 5.592935727 | *0.424059091* |
| ENSG00000051523 | CYBA | **2.077889379** | 0.238005314 | 1.778816891 | *0.23369989* |
| ENSG00000140465 | CYP1A1 | **-1.536482089** | 11.66429648 | 2.671475844 | *9.598428429* |
| ENSG00000232973 | CYP1B1-AS1 | **-3.845013223** | 0.036869842 | 0.320077959 | *0.015934885* |
| ENSG00000135929 | CYP27A1 | **3.212977833** | 1.41839305 | 10.12414629 | *2.468159685* |
| ENSG00000146233 | CYP39A1 | **2.112826286** | 10.14423453 | 28.0480129 | *5.583730927* |
| ENSG00000226562 | CYP4F26P | **1.813273264** | 0.41419907 | 4.970479572 | *0.306623442* |
| ENSG00000186529 | CYP4F3 | **2.016974563** | 0.059648728 | 15.4856393 | *0.219427634* |
| ENSG00000197872 | CYRIA | **-1.138974127** | 1.040703379 | 1.021849833 | *1.425326909* |
| ENSG00000152207 | CYSLTR2 | **2.728887087** | 0.049703972 | 0.294326837 | *0.10169442* |
| ENSG00000173406 | DAB1 | **1.464159574** | 1.519541281 | 2.860244447 | *2.039831265* |
| ENSG00000153071 | DAB2 | **3.614026588** | 0.021836603 | 0.27718036 | *0.066669958* |
| ENSG00000165617 | DACT1 | **1.1770378** | 0.795481593 | 2.650307545 | *1.561704425* |
| ENSG00000187323 | DCC | **1.573696734** | 1.388463351 | 2.879310117 | *3.533571063* |
| ENSG00000133083 | DCLK1 | **4.444567441** | 0.01617286 | 0.789609741 | *0.025986832* |
| ENSG00000163673 | DCLK3 | **2.028250612** | 0.52282636 | 2.668521389 | *0.681972103* |
| ENSG00000232093 | DCST1-AS1 | **1.033126361** | 3.350986419 | 5.054479766 | *5.438748207* |
| ENSG00000077279 | DCX | **-1.038532715** | 51.28220301 | 17.07237702 | *35.05781756* |
| ENSG00000132437 | DDC | **-1.026929386** | 34.66375192 | 13.07561877 | *39.14214817* |
| ENSG00000168209 | DDIT4 | **3.334456178** | 12.61119974 | 90.860591 | *11.10956415* |
| ENSG00000145358 | DDIT4L | **4.015023141** | 0.121678267 | 6.654708848 | *0.11916196* |

| | | | | | |
|---|---|---|---|---|---|
| ENSG00000229816 | DDX50P1 | **-1.394075838** | 2.052112602 | 43.99887587 | *1.585598584* |
| ENSG00000166153 | DEPDC4 | **-1.33008632** | 1.303256251 | 1.198392751 | *1.033681471* |
| ENSG00000165507 | DEPP1 | **2.238317243** | 121.0339613 | 354.6940987 | *58.61968406* |
| ENSG00000062282 | DGAT2 | **-1.098895275** | 6.293265507 | 2.077158636 | *5.674502474* |
| ENSG00000058866 | DGKG | **-1.228367858** | 10.60510299 | 212.8757942 | *9.813039625* |
| ENSG00000157680 | DGKI | **3.777910539** | 0.005632941 | 1.014161599 | *0.015138479* |
| ENSG00000162496 | DHRS3 | **2.678136001** | 1.433816549 | 7.054577322 | *2.372156814* |
| ENSG00000162946 | DISC1 | **1.816025928** | 0.020967276 | 0.073618944 | *0.035555062* |
| ENSG00000104371 | DKK4 | **4.491721027** | 0.150715496 | 5.780046188 | *0.737406643* |
| ENSG00000176124 | DLEU1 | **-1.462209642** | 1.127840726 | 0.284982949 | *1.21842176* |
| ENSG00000198719 | DLL1 | **-1.112849043** | 5.135121439 | 2.263690077 | *6.785660085* |
| ENSG00000176399 | DMRTA1 | **1.34603425** | 16.69531317 | 27.31599467 | *14.98857905* |
| ENSG00000167646 | DNAAF3 | **2.625226544** | 0.05335133 | 1.302931546 | *0.086189096* |
| ENSG00000007174 | DNAH9 | **4.070985716** | 0.00485465 | 14.79660566 | *0.011478741* |
| ENSG00000128590 | DNAJB9 | **1.047636243** | 163.8188184 | 219.150908 | *133.9976193* |
| ENSG00000178401 | DNAJC22 | **5.469048076** | 0.318217086 | 9.102236651 | *0.306903726* |
| ENSG00000150760 | DOCK1 | **-1.074177161** | 75.22812648 | 138.532352 | *75.09303315* |
| ENSG00000135905 | DOCK10 | **-1.023416694** | 8.02935728 | 7.40763636 | *5.374061631* |
| ENSG00000183784 | DOCK8-AS1 | **1.285327783** | 6.008952924 | 21.38672967 | *9.30355969* |
| ENSG00000167130 | DOLPP1 | **-1.078407169** | 40.74493395 | 13.40335708 | *37.86191918* |
| ENSG00000227855 | DPY19L2P3 | **1.128179173** | 0.970617335 | 6.532768429 | *1.161275963* |
| ENSG00000178904 | DPY19L3 | **-1.161723181** | 7.577563687 | 9.225337012 | *6.891981423* |
| ENSG00000113657 | DPYSL3 | **1.177101418** | 0.382296532 | 1.248395336 | *0.422221674* |
| ENSG00000151640 | DPYSL4 | **2.111635521** | 1.378585614 | 5.018585553 | *3.597443565* |
| ENSG00000136048 | DRAM1 | **-1.431686812** | 10.43225311 | 2.73773843 | *6.428134582* |
| ENSG00000149295 | DRD2 | **1.092326642** | 0.327291031 | 2.374434717 | *0.439126654* |
| ENSG00000102385 | DRP2 | **2.573132542** | 0.009133664 | 1.421794562 | *0.014060729* |
| ENSG00000134762 | DSC3 | **3.981436288** | 0.037095093 | 0.567411539 | *0.063828639* |
| ENSG00000046604 | DSG2 | **-1.124581542** | 276.7168628 | 83.41258683 | *239.5006012* |
| ENSG00000135144 | DTX1 | **-1.162288903** | 26.92127803 | 8.195129993 | *30.61640062* |
| ENSG00000178498 | DTX3 | **-1.035012286** | 31.91259715 | 52.29886166 | *35.18202077* |
| ENSG00000163840 | DTX3L | **1.138854045** | 3.4813539 | 8.996465909 | *4.467121541* |
| ENSG00000110042 | DTX4 | **4.036598304** | 0.015861054 | 5.138065957 | *0.02044074* |
| ENSG00000111266 | DUSP16 | **1.224813436** | 35.44024465 | 55.75716225 | *25.9533203* |
| ENSG00000108861 | DUSP3 | **-1.103643811** | 70.78118924 | 21.46692238 | *61.56498608* |
| ENSG00000120875 | DUSP4 | **1.229399833** | 41.78614575 | 92.55951156 | *29.67357451* |
| ENSG00000139318 | DUSP6 | **2.482024091** | 7.562477997 | 38.57646075 | *5.628910761* |
| ENSG00000130829 | DUSP9 | **-1.449662087** | 136.6885782 | 65.95816204 | *109.7451083* |
| ENSG00000221818 | EBF2 | **2.287043847** | 0.06675733 | 14.6391605 | *0.101693334* |
| ENSG00000108001 | EBF3 | **1.15969114** | 22.01580538 | 48.80781053 | *26.01050783* |
| ENSG00000171551 | ECEL1 | **3.193570933** | 4.568611641 | 27.80866137 | *6.24195361* |
| ENSG00000244280 | ECEL1P2 | **-1.711949758** | 1.643963116 | 16.79985126 | *1.300552836* |
| ENSG00000106823 | ECM2 | **3.161777765** | 0.025543764 | 13.68348821 | *0.007062312* |
| ENSG00000158813 | EDA | **1.25607622** | 1.946571806 | 3.205513205 | *2.387017058* |
| ENSG00000131080 | EDA2R | **2.187525163** | 0.177928151 | 0.687875288 | *0.112355898* |
| ENSG00000124205 | EDN3 | **4.13693942** | 0.015134629 | 1.745947457 | *0.090290671* |
| ENSG00000224023 | EDRF1-DT | **-1.027538808** | 2.155830408 | 0.86391929 | *1.293520092* |

| | | | | | |
|---|---|---|---|---|---|
| ENSG00000101210 | EEF1A2 | **-1.592799799** | 239.361135 | 51.2252758 | *257.0358485* |
| ENSG00000034239 | EFCAB1 | **2.18210266** | 0.352791079 | 1.416509366 | *0.674264702* |
| ENSG00000169242 | EFNA1 | **1.663356913** | 0.614815842 | 28.40369123 | *1.016751153* |
| ENSG00000243364 | EFNA4 | **1.424073286** | 4.22068455 | 8.087751769 | *5.287556078* |
| ENSG00000146648 | EGFR | **1.666542947** | 0.390704239 | 1.601639611 | *0.808641641* |
| ENSG00000129521 | EGLN3 | **1.367932271** | 0.327954594 | 4.25419879 | *0.428432988* |
| ENSG00000120738 | EGR1 | **1.389124881** | 55.67643445 | 101.7970034 | *22.15764333* |
| ENSG00000135625 | EGR4 | **2.492753623** | 1.196008938 | 4.449513481 | *1.671398782* |
| ENSG00000243056 | EIF4EBP3 | **-4.76099767** | 1.170501123 | 42.85033891 | *0.368472434* |
| ENSG00000111145 | ELK3 | **1.859624862** | 1.313986892 | 5.407838148 | *0.705733729* |
| ENSG00000155849 | ELMO1 | **4.145629208** | 0.019118004 | 0.232983872 | *0.007077679* |
| ENSG00000062598 | ELMO2 | **-1.114763859** | 54.47801566 | 18.11678646 | *56.18612813* |
| ENSG00000110675 | ELMOD1 | **1.695228863** | 0.529443141 | 1.194338895 | *0.422139209* |
| ENSG00000180385 | EMC3-AS1 | **1.556486919** | 12.53982577 | 32.64727413 | *16.90406477* |
| ENSG00000066629 | EML1 | **-1.83144294** | 11.53714689 | 2.678667898 | *11.57815783* |
| ENSG00000171617 | ENC1 | **1.389772328** | 1.767094681 | 15.34756445 | *1.581216727* |
| ENSG00000138792 | ENPEP | **2.612865634** | 0.099881281 | 1.562692629 | *0.191326815* |
| ENSG00000163378 | EOGT | **1.070088966** | 2.401555098 | 4.666229686 | *2.656780126* |
| ENSG00000145242 | EPHA5 | **4.408699388** | 0.025955549 | 0.531434927 | *0.019549674* |
| ENSG00000250846 | EPHA5-AS1 | **3.347148382** | 0.235606475 | 3.142857862 | *0.104956219* |
| ENSG00000080224 | EPHA6 | **2.930064089** | 0.023436164 | 0.309901743 | *0.106063697* |
| ENSG00000070886 | EPHA8 | **-1.257956989** | 18.23021782 | 5.898576225 | *11.39160239* |
| ENSG00000182580 | EPHB3 | **1.361513248** | 2.233113526 | 3.803157014 | *3.856938133* |
| ENSG00000106123 | EPHB6 | **4.235649023** | 0 | 2.572286973 | *0.002428904* |
| ENSG00000172031 | EPHX4 | **2.923002506** | 0.087865585 | 2.313290537 | *0.125339718* |
| ENSG00000273604 | EPOP | **1.762533924** | 1.010327018 | 2.211481178 | *2.334804596* |
| ENSG00000227070 | EPS15-AS1 | **2.172371545** | 3.400790268 | 10.13925281 | *2.925666779* |
| ENSG00000133106 | EPSTI1 | **2.41443765** | 0.578978213 | 3.172126002 | *0.633896606* |
| ENSG00000164308 | ERAP2 | **1.447274751** | 0.105828176 | 5.239007158 | *0.146998438* |
| ENSG00000204334 | ERICH2 | **1.188461594** | 5.333403664 | 9.836382464 | *3.823319114* |
| ENSG00000178965 | ERICH3 | **1.863145956** | 0.058570514 | 0.258620066 | *0.122156652* |
| ENSG00000177459 | ERICH5 | **-1.057712877** | 2.771599505 | 4.083908873 | *1.963037133* |
| ENSG00000178607 | ERN1 | **3.662782791** | 12.24746821 | 99.96365944 | *8.186045874* |
| ENSG00000213462 | ERV3-1 | **1.463461459** | 2.999709059 | 5.840283694 | *2.83217016* |
| ENSG00000158220 | ESYT3 | **-1.119352518** | 0.919090886 | 53.73562407 | *0.971886546* |
| ENSG00000157557 | ETS2 | **-1.449428874** | 96.49346751 | 25.50298459 | *60.09125478* |
| ENSG00000115363 | EVA1A | **4.75038938** | 0.130785142 | 2.352797975 | *0.485694531* |
| ENSG00000110723 | EXPH5 | **1.595873381** | 0.46327777 | 13.05387584 | *0.499288959* |
| ENSG00000104313 | EYA1 | **4.409246437** | 0.003003243 | 1.274314365 | *0.005556599* |
| ENSG00000131187 | F12 | **1.867822707** | 0.077099418 | 0.612044024 | *0.291078179* |
| ENSG00000181104 | F2R | **1.547647146** | 0.428995611 | 0.843470544 | *0.694037928* |
| ENSG00000117525 | F3 | **1.042088217** | 6.200014367 | 8.670377698 | *8.522642743* |
| ENSG00000198734 | F5 | **2.104570179** | 0.127628861 | 0.825346173 | *0.13061295* |
| ENSG00000185010 | F8 | **1.139718255** | 0.221029815 | 4.39938399 | *0.347080632* |
| ENSG00000103089 | FA2H | **-1.278744883** | 10.57241702 | 2.831800672 | *11.42151682* |
| ENSG00000163586 | FABP1 | **-1.28727969** | 0.919638243 | 0.434591668 | *2.745032646* |
| ENSG00000164687 | FABP5 | **2.255242869** | 5.036906727 | 16.88156461 | *9.110468846* |

| | | | | | |
|---|---|---|---|---|---|
| ENSG00000149485 | FADS1 | **-1.299823971** | 91.00229559 | 24.30362292 | *94.76328749* |
| ENSG00000135472 | FAIM2 | **1.037844902** | 3.806349112 | 13.39500705 | *4.693747858* |
| ENSG00000248019 | FAM13A-AS1 | **-1.483387937** | 0.587146732 | 12.44296402 | *0.346882199* |
| ENSG00000148541 | FAM13C | **-1.071503576** | 1.599039142 | 3.176942799 | *1.132174134* |
| ENSG00000170074 | FAM153A | **1.476493239** | 0.05482995 | 0.145665901 | *0.085731788* |
| ENSG00000164142 | FAM160A1 | **1.693844169** | 0.417103394 | 1.128854205 | *0.521249449* |
| ENSG00000154319 | FAM167A | **-1.628632806** | 52.15537313 | 10.90012222 | *49.25708327* |
| ENSG00000283597 | FAM169B | **1.355932766** | 0.283823682 | 0.909804698 | *0.381139615* |
| ENSG00000148468 | FAM171A1 | **1.186148452** | 20.10194089 | 35.17307173 | *24.18252159* |
| ENSG00000104059 | FAM189A1 | **3.527652927** | 0.345043682 | 2.90552925 | *0.259747471* |
| ENSG00000204860 | FAM201A | **1.256387656** | 2.123104739 | 19.22063348 | *3.127997169* |
| ENSG00000108950 | FAM20A | **2.508814731** | 0.056821234 | 1.505497738 | *0.159802821* |
| ENSG00000005238 | FAM214B | **-1.010185383** | 2.937489728 | 2.620644492 | *2.552191816* |
| ENSG00000235118 | FAM237A | **3.868437421** | 0.056766693 | 0.75467321 | *0.318905242* |
| ENSG00000228055 | FAM245A | **-1.228334341** | 9.956058013 | 3.238077784 | *7.230841696* |
| ENSG00000231527 | FAM27C | **3.026294968** | 0.113100789 | 0.876659382 | *0.065635238* |
| ENSG00000185112 | FAM43A | **1.521043666** | 5.816802441 | 12.26214946 | *5.214586735* |
| ENSG00000183114 | FAM43B | **-2.24120489** | 5.602041506 | 1.055086379 | *9.552996347* |
| ENSG00000158483 | FAM86C1 | **-1.282295538** | 3.241385543 | 6.522675295 | *3.260073399* |
| ENSG00000171084 | FAM86JP | **-1.007067681** | 2.26610217 | 1.180314391 | *2.08662479* |
| ENSG00000182118 | FAM89A | **1.301918103** | 5.454283297 | 8.992244802 | *8.737719433* |
| ENSG00000283486 | FAM95C | **2.161412867** | 0.154119026 | 0.845919576 | *0.245618526* |
| ENSG00000083857 | FAT1 | **3.201075957** | 0.419856434 | 7.411504545 | *0.733738541* |
| ENSG00000165323 | FAT3 | **2.192077587** | 0.085124794 | 0.442965499 | *0.102226792* |
| ENSG00000196159 | FAT4 | **1.701629185** | 5.501325683 | 13.04491068 | *4.752271171* |
| ENSG00000166147 | FBN1 | **1.626564342** | 0.436831335 | 1.03435689 | *0.610697853* |
| ENSG00000183580 | FBXL7 | **1.084917133** | 4.986313433 | 12.7580952 | *7.282990438* |
| ENSG00000116661 | FBXO2 | **-1.456688985** | 22.06715013 | 5.611588996 | *18.76102365* |
| ENSG00000163497 | FEV | **5.917420466** | 1.859943975 | 75.65375937 | *1.743479276* |
| ENSG00000186188 | FFAR4 | **-1.319436346** | 0.791837867 | 2.947627348 | *0.331552333* |
| ENSG00000171564 | FGB | **-1.80689622** | 0.831385123 | 38.48202747 | *0.484306339* |
| ENSG00000127084 | FGD3 | **1.490286758** | 0.312540519 | 0.724062846 | *0.309972497* |
| ENSG00000156427 | FGF18 | **-1.788799553** | 237.9293097 | 45.72513173 | *156.6987423* |
| ENSG00000138685 | FGF2 | **1.037349182** | 6.598826677 | 8.946662009 | *6.338709159* |
| ENSG00000219693 | FGF7P8 | **1.645206231** | 2.863339983 | 28.61339305 | *3.527949217* |
| ENSG00000107831 | FGF8 | **4.912941679** | 0 | 4.798576463 | *0* |
| ENSG00000174721 | FGFBP3 | **1.024949544** | 14.53355749 | 22.76306639 | *11.41505326* |
| ENSG00000127418 | FGFRL1 | **3.170433481** | 0.372115766 | 2.116152715 | *0.581404045* |
| ENSG00000134775 | FHOD3 | **1.317014713** | 1.757577545 | 11.86770061 | *1.855444805* |
| ENSG00000261308 | FIGNL2 | **1.031908497** | 3.455367604 | 6.022210523 | *3.893646282* |
| ENSG00000168386 | FILIP1L | **2.032512558** | 0.033471851 | 1.571948394 | *0.066510001* |
| ENSG00000179431 | FJX1 | **2.017633098** | 0.358148881 | 3.154616306 | *0.907910806* |
| ENSG00000141756 | FKBP10 | **1.348434466** | 0.315770963 | 0.570148617 | *0.527768139* |
| ENSG00000198225 | FKBP1C | **-1.271849546** | 3.519822927 | 1.349095968 | *2.347380041* |
| ENSG00000128591 | FLNC | **1.35572199** | 0.069490874 | 0.390769548 | *0.137054903* |
| ENSG00000185070 | FLRT2 | **8.190618487** | 0.001715489 | 0.964416275 | *0.00142549* |
| ENSG00000161791 | FMNL3 | **1.547550372** | 0.621993057 | 1.275663071 | *0.965514918* |

| | | | | | |
|---|---|---|---|---|---|
| ENSG00000131781 | FMO5 | **1.171983095** | 0.782839862 | 1.384389789 | *0.644495874* |
| ENSG00000115414 | FN1 | **1.286186321** | 2.247305373 | 4.119248702 | *2.891349122* |
| ENSG00000115226 | FNDC4 | **1.666053647** | 2.465232785 | 5.486130975 | *2.789887999* |
| ENSG00000170345 | FOS | **1.447013012** | 2.938351213 | 7.128814733 | *2.573736359* |
| ENSG00000075426 | FOSL2 | **2.502472808** | 0.634228982 | 4.965781113 | *0.78279197* |
| ENSG00000054598 | FOXC1 | **4.212548295** | 0.016948777 | 2.876494331 | *0.038990441* |
| ENSG00000251493 | FOXD1 | **1.970952767** | 18.75578341 | 48.36133711 | *32.74370439* |
| ENSG00000237424 | FOXD2-AS1 | **1.714259615** | 3.178240919 | 6.920883794 | *5.016496756* |
| ENSG00000128573 | FOXP2 | **1.979813717** | 0.664772501 | 27.57599833 | *0.910766818* |
| ENSG00000138759 | FRAS1 | **1.682804537** | 3.854358924 | 11.77931893 | *4.721264065* |
| ENSG00000172159 | FRMD3 | **2.950321206** | 1.318667731 | 7.522752085 | *2.297470352* |
| ENSG00000139926 | FRMD6 | **1.09903002** | 0.682941923 | 4.861672782 | *0.7429843* |
| ENSG00000070601 | FRMPD1 | **1.767226804** | 2.431172791 | 8.809895265 | *3.098423822* |
| ENSG00000147234 | FRMPD3 | **-3.869157489** | 1.071325058 | 0.560861603 | *0.492192947* |
| ENSG00000075618 | FSCN1 | **1.360722038** | 0.338380475 | 3.2066313 | *0.487140208* |
| ENSG00000134363 | FST | **4.106329175** | 0.010212281 | 0.153439303 | *0.00806001* |
| ENSG00000163430 | FSTL1 | **1.669358042** | 0.192639864 | 0.731774029 | *0.376216056* |
| ENSG00000160282 | FTCD | **-1.168957145** | 9.557916258 | 2.981562569 | *12.55337373* |
| ENSG00000237338 | FTCD-AS1 | **2.319473443** | 0.67772297 | 1.904716724 | *0.248364662* |
| ENSG00000260459 | FTLP14 | **1.033884044** | 10.34406309 | 15.02658819 | *14.02038166* |
| ENSG00000179163 | FUCA1 | **-1.072842727** | 304.7366482 | 96.76276333 | *296.6225656* |
| ENSG00000137731 | FXYD2 | **-3.337433129** | 4.001243027 | 7.246137555 | *7.149740806* |
| ENSG00000137726 | FXYD6 | **-1.618698785** | 26.19440471 | 53.63375751 | *34.2804617* |
| ENSG00000111432 | FZD10 | **4.072680971** | 0 | 0.262959693 | *0.008736692* |
| ENSG00000250208 | FZD10-AS1 | **4.181606773** | 0.004818929 | 2.727273436 | *0.007714591* |
| ENSG00000164930 | FZD6 | **1.332761568** | 5.373596377 | 8.981543396 | *5.492854339* |
| ENSG00000155760 | FZD7 | **2.083579107** | 4.935845376 | 13.44970341 | *5.258881357* |
| ENSG00000177283 | FZD8 | **1.936230077** | 0.274336124 | 5.026505436 | *0.289199845* |
| ENSG00000152254 | G6PC2 | **1.65298237** | 19.49439322 | 47.85964383 | *42.79910265* |
| ENSG00000109458 | GAB1 | **1.190065652** | 3.354457922 | 5.488071317 | *3.647121501* |
| ENSG00000136928 | GABBR2 | **1.306200369** | 0.560053752 | 20.64388528 | *1.705566603* |
| ENSG00000163288 | GABRB1 | **1.145570943** | 1.008529181 | 3.910001302 | *1.059074845* |
| ENSG00000187730 | GABRD | **6.787604122** | 0.005050785 | 0.799696615 | *0.012769563* |
| ENSG00000113327 | GABRG2 | **2.873647787** | 0.00516336 | 0.795598995 | *0.009642262* |
| ENSG00000182256 | GABRG3 | **2.87677246** | 0.012521275 | 0.284649687 | *0.014468475* |
| ENSG00000099860 | GADD45B | **1.257271685** | 0.780086049 | 1.202015882 | *0.601623291* |
| ENSG00000069482 | GAL | **4.247666174** | 0.139427481 | 1.676481 | *0.129370486* |
| ENSG00000054983 | GALC | **-1.536097685** | 17.79858587 | 4.632975348 | *15.45404709* |
| ENSG00000108479 | GALK1 | **3.821472615** | 0.230447423 | 3.192697226 | *0.564079948* |
| ENSG00000115339 | GALNT3 | **-1.535640727** | 33.15384826 | 9.711103779 | *30.67313959* |
| ENSG00000257594 | GALNT4 | **1.621113668** | 0.349740124 | 1.886036459 | *0.287594147* |
| ENSG00000174473 | GALNTL6 | **7.305971266** | 0 | 3.897543777 | *0.007878978* |
| ENSG00000182687 | GALR2 | **1.339786845** | 0.963416036 | 1.891953473 | *1.11067625* |
| ENSG00000148935 | GAS2 | **1.962145258** | 0.865673951 | 2.297190266 | *1.230291837* |
| ENSG00000184502 | GAST | **-1.90991237** | 5.628288545 | 2.085602231 | *4.201552237* |
| ENSG00000249948 | GBA3 | **4.731349212** | 0.171633564 | 4.035079693 | *0.226581726* |
| ENSG00000148288 | GBGT1 | **1.185516271** | 0.371123929 | 0.707310375 | *0.497233142* |

| | | | | | |
|---|---|---|---|---|---|
| ENSG00000145321 | GC | **-3.166497112** | 137.14585 | 11.26859508 | *136.7926808* |
| ENSG00000115263 | GCG | **5.649286545** | 0.080129434 | 2.871525145 | *0.195568102* |
| ENSG00000023909 | GCLM | **1.076737044** | 72.95932909 | 107.0151986 | *75.73048085* |
| ENSG00000147174 | GCNA | **-1.108084214** | 1.365759867 | 1.711757821 | *0.744919756* |
| ENSG00000111846 | GCNT2 | **1.542831538** | 3.72381361 | 57.40402422 | *3.195367983* |
| ENSG00000205318 | GCNT2P1 | **4.343670231** | 0 | 0.672684856 | *0.265915056* |
| ENSG00000176928 | GCNT4 | **1.069688881** | 6.689173566 | 12.40823945 | *6.612755361* |
| ENSG00000266524 | GDF10 | **1.939823853** | 0.361424241 | 1.19342728 | *0.687969924* |
| ENSG00000130513 | GDF15 | **1.186449335** | 22.05402392 | 38.30391818 | *9.935306238* |
| ENSG00000156466 | GDF6 | **2.922607858** | 0.046664216 | 0.80389569 | *0.115046725* |
| ENSG00000143869 | GDF7 | **7.040976269** | 0 | 16.75244351 | *0.012737956* |
| ENSG00000158555 | GDPD5 | **1.29356667** | 1.263711189 | 2.016738953 | *1.446133539* |
| ENSG00000146013 | GFRA3 | **1.286127106** | 0.749929183 | 1.357866017 | *0.711245025* |
| ENSG00000251000 | GGCTP1 | **2.476671713** | 0.975754324 | 4.57918466 | *1.355425391* |
| ENSG00000112964 | GHR | **-1.051914638** | 4.696726348 | 2.13674163 | *6.372040602* |
| ENSG00000169562 | GJB1 | **-2.415892575** | 0.482415066 | 2.070220409 | *0.471581732* |
| ENSG00000159248 | GJD2 | **-1.224314156** | 71.5625691 | 20.18387313 | *34.31542968* |
| ENSG00000178445 | GLDC | **1.255009587** | 0.177975478 | 0.316167607 | *0.231237597* |
| ENSG00000111087 | GLI1 | **3.657780457** | 0.004560016 | 10.81585041 | *0.007794522* |
| ENSG00000074047 | GLI2 | **7.112351844** | 0 | 0.222884159 | *0* |
| ENSG00000106571 | GLI3 | **3.54089191** | 0.223314139 | 1.732957809 | *0.271487722* |
| ENSG00000122694 | GLIPR2 | **1.61003652** | 13.16321553 | 26.84126052 | *11.35827378* |
| ENSG00000107249 | GLIS3 | **-1.187929675** | 9.04750525 | 3.401999855 | *8.707122373* |
| ENSG00000112164 | GLP1R | **-2.243772453** | 1.402514631 | 13.02131547 | *1.696332057* |
| ENSG00000145451 | GLRA3 | **3.118649724** | 1.179471497 | 7.973991662 | *0.966771378* |
| ENSG00000151948 | GLT1D1 | **2.653303173** | 0.035435661 | 0.27183732 | *0.036418142* |
| ENSG00000120820 | GLT8D2 | **1.868570483** | 1.064125349 | 6.071115229 | *1.05992286* |
| ENSG00000182890 | GLUD2 | **1.135863597** | 2.898654073 | 4.509884507 | *4.885435968* |
| ENSG00000089639 | GMIP | **1.220821573** | 1.968335718 | 4.224474628 | *3.149844727* |
| ENSG00000205835 | GMNC | **1.80889747** | 0.124339857 | 2.05337861 | *0.078036829* |
| ENSG00000232284 | GNG12-AS1 | **3.342583424** | 0.0249614 | 1.774267647 | *0.067738467* |
| ENSG00000188626 | GOLGA8M | **4.97648228** | 0 | 0.538333124 | *0.071161775* |
| ENSG00000179399 | GPC5 | **1.134997471** | 3.36116051 | 4.833743051 | *4.897097997* |
| ENSG00000167588 | GPD1 | **-1.001531306** | 2.063761808 | 0.867029215 | *2.54334716* |
| ENSG00000112293 | GPLD1 | **1.02601947** | 1.602727185 | 4.605798655 | *1.571557097* |
| ENSG00000150625 | GPM6A | **2.685855275** | 0.012270719 | 0.335718132 | *0.013332302* |
| ENSG00000046653 | GPM6B | **-1.670959759** | 0.234448019 | 1.200777827 | *0.348439848* |
| ENSG00000136235 | GPNMB | **1.235220739** | 0.310324506 | 0.510832317 | *0.246447247* |
| ENSG00000132975 | GPR12 | **4.110698072** | 0.061677006 | 0.775307632 | *0.17785641* |
| ENSG00000101850 | GPR143 | **1.044521507** | 6.848605242 | 9.262358239 | *6.405972614* |
| ENSG00000164849 | GPR146 | **-1.008071534** | 8.670247658 | 2.909900208 | *7.549151539* |
| ENSG00000173302 | GPR148 | **1.344148345** | 1.279730438 | 7.199132195 | *1.479225473* |
| ENSG00000178015 | GPR150 | **4.042561356** | 0.102196693 | 2.592018941 | *0.633225598* |
| ENSG00000166073 | GPR176 | **-1.19432034** | 46.42980873 | 13.90373525 | *53.34836592* |
| ENSG00000277399 | GPR179 | **-1.002710578** | 1.78507901 | 1.242204187 | *2.029918032* |
| ENSG00000181773 | GPR3 | **1.318683444** | 4.955479574 | 15.02718255 | *4.273781402* |
| ENSG00000170075 | GPR37L1 | **-6.578906305** | 5.421761877 | 0.35616939 | *0.035308191* |

| | | | | | |
|---|---|---|---|---|---|
| ENSG00000146360 | GPR6 | **1.019563389** | 2.362603908 | 7.204688209 | *2.599073* |
| ENSG00000138271 | GPR87 | **5.864054323** | 0 | 0.616937832 | *0.061281342* |
| ENSG00000167191 | GPRC5B | **1.120279605** | 2.283369155 | 4.886895138 | *3.518423506* |
| ENSG00000204175 | GPRIN2 | **-1.393285143** | 5.203390736 | 1.448196704 | *6.668264861* |
| ENSG00000160360 | GPSM1 | **-1.280890479** | 43.88517383 | 13.72580923 | *55.89917837* |
| ENSG00000167701 | GPT | **-1.063886612** | 1.970470414 | 1.313706664 | *1.822840895* |
| ENSG00000176153 | GPX2 | **-3.298261133** | 10.99377314 | 6.487669301 | *4.034104036* |
| ENSG00000161835 | GRASP | **-2.272672278** | 21.93570611 | 3.289362179 | *26.75040365* |
| ENSG00000141449 | GREB1L | **1.472763022** | 0.155205079 | 0.727522536 | *0.289874176* |
| ENSG00000180875 | GREM2 | **1.118449874** | 1.380188549 | 3.433877146 | *1.231564279* |
| ENSG00000171189 | GRIK1 | **2.87233249** | 0.014077101 | 0.206523204 | *0.010365772* |
| ENSG00000163873 | GRIK3 | **-2.576411384** | 4.527897398 | 1.546724884 | *7.651486889* |
| ENSG00000183454 | GRIN2A | **-1.803462** | 0.2207197 | 0.081716818 | *0.300004537* |
| ENSG00000164082 | GRM2 | **1.335722169** | 0.537650387 | 1.049854249 | *1.233449899* |
| ENSG00000196277 | GRM7 | **4.90929282** | 0.006361219 | 0.150955494 | *0.003580437* |
| ENSG00000179603 | GRM8 | **1.052854758** | 0.536938514 | 1.198421408 | *1.130527959* |
| ENSG00000126010 | GRPR | **3.939428638** | 0.544922835 | 5.051552931 | *1.141896051* |
| ENSG00000139835 | GRTP1 | **3.435867098** | 0.195889611 | 1.711379559 | *0.528000834* |
| ENSG00000084207 | GSTP1 | **1.8789078** | 0.144869666 | 3.576821756 | *0.113851847* |
| ENSG00000152402 | GUCY1A2 | **1.769547325** | 2.498549124 | 6.464861548 | *3.552988516* |
| ENSG00000144366 | GULP1 | **2.10298869** | 1.769487688 | 5.181439658 | *2.439923536* |
| ENSG00000124575 | H1-3 | **3.604576556** | 0.311396131 | 5.036572338 | *0.621819219* |
| ENSG00000130600 | H19 | **1.290749974** | 19.00089479 | 32.66951633 | *13.00224351* |
| ENSG00000275221 | H2AC15 | **1.192876122** | 25.83728225 | 39.31192487 | *29.08706288* |
| ENSG00000124635 | H2BC11 | **1.128597643** | 9.640815318 | 30.26189227 | *8.588577483* |
| ENSG00000274641 | H2BC17 | **2.537994977** | 0.753160976 | 22.91462551 | *1.542718344* |
| ENSG00000220323 | H2BC19P | **-1.01218912** | 55.97472634 | 23.98585012 | *45.86368024* |
| ENSG00000197153 | H3C12 | **1.626204772** | 1.518925046 | 4.369325922 | *1.841282154* |
| ENSG00000270882 | H4C14 | **-1.05770038** | 3.309263867 | 10.74095587 | *2.238475921* |
| ENSG00000138796 | HADH | **-1.131948751** | 90.0254787 | 29.03803378 | *85.94359069* |
| ENSG00000224189 | HAGLR | **2.715925512** | 0.023405438 | 0.6904486 | *0.073321914* |
| ENSG00000226363 | HAGLROS | **4.320188496** | 0 | 14.06148631 | *0.157804471* |
| ENSG00000116478 | HDAC1 | **-1.348561115** | 197.5077273 | 51.75189014 | *230.3476305* |
| ENSG00000228624 | HDAC2-AS2 | **1.02019092** | 0.358254953 | 0.672251308 | *0.501324551* |
| ENSG00000061273 | HDAC7 | **3.036348168** | 0.030277231 | 25.05602388 | *0.07580208* |
| ENSG00000138411 | HECW2 | **1.199948787** | 3.850715678 | 6.055123919 | *4.325371618* |
| ENSG00000259845 | HERC2P10 | **4.184014131** | 0 | 0.336294132 | *0.195089044* |
| ENSG00000138642 | HERC6 | **1.135228637** | 1.464261219 | 5.09992779 | *2.299426714* |
| ENSG00000114315 | HES1 | **1.698194005** | 2.995385576 | 6.372130525 | *3.722729702* |
| ENSG00000188290 | HES4 | **2.488282412** | 0.116723472 | 1.446406316 | *0.292022032* |
| ENSG00000144485 | HES6 | **1.093698248** | 68.73722448 | 98.25340256 | *92.4567607* |
| ENSG00000164683 | HEY1 | **1.122243015** | 2.639936187 | 3.875262194 | *2.62730671* |
| ENSG00000135547 | HEY2 | **1.175838122** | 0.614658454 | 50.83327268 | *1.220992291* |
| ENSG00000113924 | HGD | **1.638562745** | 3.759489828 | 9.551667003 | *5.999920934* |
| ENSG00000152804 | HHEX | **4.52105317** | 3.566666589 | 53.81832931 | *5.483056376* |
| ENSG00000177374 | HIC1 | **1.101886127** | 5.80490137 | 11.99816681 | *4.609387172* |
| ENSG00000159399 | HK2 | **4.168064479** | 0.030497494 | 27.91481833 | *0.075929809* |

| | | | | | |
|---|---|---|---|---|---|
| ENSG00000206503 | HLA-A | **1.503950055** | 282.549366 | 522.8491327 | *261.507936* |
| ENSG00000234745 | HLA-B | **2.031979835** | 0.099065823 | 0.476606447 | *0.11265723* |
| ENSG00000204525 | HLA-C | **1.322631521** | 1.368831957 | 275.1331127 | *2.184790037* |
| ENSG00000204592 | HLA-E | **1.19710588** | 198.0078681 | 294.2993699 | *214.9537063* |
| ENSG00000206341 | HLA-H | **2.004657279** | 4.015452384 | 11.85656724 | *3.664746516* |
| ENSG00000149948 | HMGA2 | **3.037995371** | 0.008268552 | 154.2642897 | *0.011830926* |
| ENSG00000100292 | HMOX1 | **2.594461765** | 0.563831154 | 7.457633325 | *0.308466222* |
| ENSG00000221887 | HMSD | **1.213147568** | 0.718398666 | 1.135767852 | *1.468229435* |
| ENSG00000258900 | HNRNPCP1 | **1.224553451** | 4.188335818 | 7.902406878 | *4.852282954* |
| ENSG00000051128 | HOMER3 | **2.258412073** | 0.17845534 | 1.062962394 | *0.180770843* |
| ENSG00000171476 | HOPX | **1.873350389** | 0.793556716 | 4.879978621 | *1.242722159* |
| ENSG00000159184 | HOXB13 | **1.775445898** | 0.322040272 | 1.028002604 | *0.135988033* |
| ENSG00000128645 | HOXD1 | **3.545437983** | 0.21166969 | 2.635966575 | *0.520776612* |
| ENSG00000113749 | HRH2 | **1.9497527** | 0.144882186 | 0.742630243 | *0.208567882* |
| ENSG00000135116 | HRK | **2.518103882** | 3.436570919 | 13.84422754 | *5.826759018* |
| ENSG00000171004 | HS6ST2 | **3.48483383** | 0.10948531 | 0.94004071 | *0.152348877* |
| ENSG00000099377 | HSD3B7 | **1.064089334** | 2.210837773 | 9.494434654 | *3.659907466* |
| ENSG00000173110 | HSPA6 | **2.620616701** | 1.436403003 | 6.171416853 | *1.251185508* |
| ENSG00000142798 | HSPG2 | **1.366451471** | 5.245487878 | 10.82637991 | *5.349635151* |
| ENSG00000178394 | HTR1A | **6.179318924** | 0 | 3.896459212 | *0.280133741* |
| ENSG00000135312 | HTR1B | **2.083548803** | 3.867055215 | 14.4987648 | *3.509197234* |
| ENSG00000179546 | HTR1D | **1.179187729** | 4.138486119 | 7.075998397 | *4.795529598* |
| ENSG00000148680 | HTR7 | **1.730107467** | 0.973636688 | 7.785653403 | *1.021850275* |
| ENSG00000242028 | HYPK | **-1.002593135** | 2.717370513 | 3.377813464 | *2.099507898* |
| ENSG00000121351 | IAPP | **-1.975311015** | 2403.321407 | 390.6421582 | *1331.948482* |
| ENSG00000125968 | ID1 | **3.925287005** | 2.344951463 | 22.10334405 | *2.006512955* |
| ENSG00000235092 | ID2-AS1 | **2.524805072** | 0.555855493 | 216.6147661 | *0.818027407* |
| ENSG00000117318 | ID3 | **3.008332244** | 0.544932127 | 16.21638606 | *0.609691845* |
| ENSG00000137331 | IER3 | **-2.536549287** | 6.262852819 | 1.819791394 | *1.49561319* |
| ENSG00000165948 | IFI27L1 | **1.039445493** | 4.488374476 | 7.308147329 | *6.368559537* |
| ENSG00000073792 | IGF2BP2 | **2.181204066** | 0.031021535 | 0.411680809 | *0.020822299* |
| ENSG00000146678 | IGFBP1 | **1.991393206** | 0.101655806 | 3.465117675 | *0.172166007* |
| ENSG00000146674 | IGFBP3 | **3.388816481** | 0.179589772 | 1.293306026 | *0.191937763* |
| ENSG00000141753 | IGFBP4 | **1.466818968** | 0.721122007 | 1.523536324 | *1.6137503* |
| ENSG00000167779 | IGFBP6 | **1.032467131** | 1.628099032 | 2.705725461 | *2.039863633* |
| ENSG00000152580 | IGSF10 | **1.015441012** | 7.131355761 | 10.00557167 | *6.112156836* |
| ENSG00000095574 | IKZF5 | **-1.065667747** | 32.01448419 | 11.34958025 | *26.18654049* |
| ENSG00000110324 | IL10RA | **-1.002453288** | 3.477451909 | 5.891324391 | *3.98470416* |
| ENSG00000168811 | IL12A | **4.504089461** | 0 | 5.060397341 | *0* |
| ENSG00000081985 | IL12RB2 | **2.896016775** | 0.050681012 | 0.914875478 | *0.083851374* |
| ENSG00000163701 | IL17RE | **1.37347347** | 0.262870456 | 0.506029794 | *0.362896108* |
| ENSG00000115590 | IL1R2 | **2.737776325** | 0.154768013 | 0.767660026 | *0.451509954* |
| ENSG00000162594 | IL23R | **1.409310482** | 0.831894131 | 1.634126346 | *0.897858502* |
| ENSG00000077238 | IL4R | **5.266269695** | 0 | 0.382896732 | *0.002728909* |
| ENSG00000091181 | IL5RA | **4.209158496** | 0.019892283 | 1.006255622 | *0.019638152* |
| ENSG00000143195 | ILDR2 | **1.249673102** | 1.071023075 | 1.645088149 | *0.95010809* |
| ENSG00000154059 | IMPACT | **-1.143064653** | 32.97941299 | 10.02205184 | *30.55408921* |

| | | | | | |
|---|---|---|---|---|---|
| ENSG00000163362 | INAVA | **2.228416701** | 0.0970475 | 1.205031425 | *0.076243748* |
| ENSG00000123999 | INHA | **1.892328778** | 0.816484984 | 6.953217902 | *1.162311515* |
| ENSG00000163083 | INHBB | **4.613548848** | 0 | 0.30492877 | *0* |
| ENSG00000139269 | INHBE | **2.506533822** | 0.214734464 | 1.643052668 | *0.193541087* |
| ENSG00000068383 | INPP5A | **-1.080568608** | 20.2886569 | 6.42952897 | *21.41542787* |
| ENSG00000129965 | INS-IGF2 | **1.165069685** | 1.348567506 | 2.099742282 | *0.881699325* |
| ENSG00000205363 | INSYN1 | **1.347402763** | 9.377553602 | 18.18019981 | *10.58671303* |
| ENSG00000260469 | INSYN1-AS1 | **1.20819862** | 20.21560875 | 32.75573403 | *18.35712777* |
| ENSG00000188916 | INSYN2A | **1.534073501** | 1.796065621 | 11.87433247 | *1.71926058* |
| ENSG00000074706 | IPCEF1 | **4.476963967** | 0.010148088 | 14.80030463 | *0.017497384* |
| ENSG00000185507 | IRF7 | **-1.189305041** | 0.500631808 | 1.912167318 | *0.607894402* |
| ENSG00000159556 | ISL2 | **4.874830434** | 0 | 0.334358748 | *0* |
| ENSG00000164171 | ITGA2 | **1.66407478** | 1.63795539 | 3.457258654 | *2.552801738* |
| ENSG00000115232 | ITGA4 | **4.637215846** | 0.017679464 | 0.283733968 | *0.019128316* |
| ENSG00000135424 | ITGA7 | **2.369606433** | 2.0235323 | 8.672264918 | *3.274131985* |
| ENSG00000077943 | ITGA8 | **2.744092027** | 0.026942942 | 0.287928318 | *0.031020531* |
| ENSG00000144668 | ITGA9 | **4.267719311** | 0.159109346 | 5.432596536 | *0.245066891* |
| ENSG00000115221 | ITGB6 | **3.044473849** | 0.006650734 | 0.086497742 | *0.008815955* |
| ENSG00000105855 | ITGB8 | **2.839297848** | 0.75505874 | 4.499910451 | *0.711599033* |
| ENSG00000055957 | ITIH1 | **-3.533433115** | 1.23129152 | 0.086173845 | *0.98707862* |
| ENSG00000231249 | ITPR1-DT | **1.342234672** | 21.83355667 | 40.0678649 | *16.62921183* |
| ENSG00000123104 | ITPR2 | **1.616023119** | 4.529823492 | 8.961028682 | *7.137006463* |
| ENSG00000148841 | ITPRIP | **3.38353127** | 0.140908097 | 17.43118779 | *0.198434665* |
| ENSG00000154721 | JAM2 | **2.650582951** | 0.188320985 | 5.536521419 | *0.082515906* |
| ENSG00000140044 | JDP2 | **1.590820269** | 0.075865302 | 0.631747772 | *0.052893439* |
| ENSG00000177272 | KCNA3 | **1.616930597** | 54.66289354 | 110.2823953 | *49.04028871* |
| ENSG00000169282 | KCNAB1 | **3.113525467** | 0.005662996 | 0.127989445 | *0.010370164* |
| ENSG00000116396 | KCNC4 | **-2.190995578** | 1.027466187 | 55.95552122 | *0.902265561* |
| ENSG00000171385 | KCND3 | **-1.676266535** | 3.514014421 | 0.730240778 | *9.049667317* |
| ENSG00000152049 | KCNE4 | **2.659181047** | 0.24608306 | 0.964908793 | *0.823300875* |
| ENSG00000162975 | KCNF1 | **1.78532374** | 1.207817023 | 2.98721815 | *1.562403067* |
| ENSG00000026559 | KCNG1 | **2.406220731** | 0.089317455 | 0.963349467 | *0.214795354* |
| ENSG00000178342 | KCNG2 | **1.735701879** | 12.81916478 | 29.54772321 | *14.29618839* |
| ENSG00000171126 | KCNG3 | **-1.543462334** | 26.75657567 | 6.30407323 | *23.4168137* |
| ENSG00000143473 | KCNH1 | **-1.90569615** | 0.792383913 | 14.33584527 | *0.766886854* |
| ENSG00000135519 | KCNH3 | **-1.031645529** | 26.70511368 | 11.28093655 | *33.45453066* |
| ENSG00000140015 | KCNH5 | **3.627115384** | 0.097434237 | 0.832670789 | *0.230844882* |
| ENSG00000115041 | KCNIP3 | **2.044800663** | 0.03746237 | 4.628811689 | *0.064725728* |
| ENSG00000267365 | KCNJ2-AS1 | **1.511789695** | 8.723973357 | 16.78002795 | *7.065883873* |
| ENSG00000168135 | KCNJ4 | **1.181670328** | 1.544077518 | 2.274797169 | *1.843228932* |
| ENSG00000121361 | KCNJ8 | **2.159599753** | 10.26012997 | 38.6793866 | *19.24690469* |
| ENSG00000135750 | KCNK1 | **2.752894967** | 1.497611947 | 7.902877816 | *2.1514737* |
| ENSG00000100433 | KCNK10 | **1.022609574** | 0.697207403 | 15.62150866 | *0.728103047* |
| ENSG00000184261 | KCNK12 | **1.699130292** | 0.266853178 | 3.829567811 | *0.375702663* |
| ENSG00000082482 | KCNK2 | **1.448777503** | 0.323535355 | 0.946782011 | *0.505431425* |
| ENSG00000171303 | KCNK3 | **2.292992066** | 0.871319686 | 2.966025294 | *1.412354602* |
| ENSG00000099337 | KCNK6 | **1.670319034** | 1.052060169 | 2.586595797 | *1.498395758* |

| | | | | | |
|---|---|---|---|---|---|
| ENSG00000169427 | KCNK9 | **1.748899525** | 1.007708527 | 3.720263678 | *1.771986295* |
| ENSG00000156113 | KCNMA1 | **-1.126800863** | 5.244062389 | 2.648675115 | *4.954141513* |
| ENSG00000053918 | KCNQ1 | **1.272809214** | 0.986852055 | 2.697442601 | *1.993418836* |
| ENSG00000075043 | KCNQ2 | **-1.025889757** | 2.056085028 | 1.46015105 | *2.642643346* |
| ENSG00000170745 | KCNS3 | **1.149588761** | 1.399863473 | 2.833949467 | *1.900223955* |
| ENSG00000164794 | KCNV1 | **1.697817235** | 2.122837061 | 4.822043667 | *2.42297843* |
| ENSG00000135253 | KCP | **1.487131879** | 0.051815264 | 1.18313168 | *0.186286915* |
| ENSG00000178695 | KCTD12 | **1.216376608** | 225.8542441 | 344.8926526 | *316.2453724* |
| ENSG00000153885 | KCTD15 | **3.834871728** | 0.09648956 | 0.921737637 | *0.248813421* |
| ENSG00000183775 | KCTD16 | **1.883447275** | 5.185573027 | 187.9200939 | *5.10064226* |
| ENSG00000100196 | KDELR3 | **2.682575385** | 1.237273493 | 5.875264846 | *1.267793742* |
| ENSG00000162522 | KIAA1522 | **1.138651453** | 8.697844074 | 18.79495695 | *10.71210785* |
| ENSG00000117245 | KIF17 | **1.430594996** | 0.240586828 | 2.82351868 | *0.378690021* |
| ENSG00000162849 | KIF26B | **1.861946881** | 0.291727479 | 7.038419534 | *0.464800898* |
| ENSG00000101350 | KIF3B | **-1.290212679** | 395.1538338 | 105.4649732 | *315.2734464* |
| ENSG00000155980 | KIF5A | **-1.315387457** | 28.08365251 | 7.847558063 | *21.23926416* |
| ENSG00000166813 | KIF7 | **2.928821971** | 0.118367188 | 54.92599402 | *0.275658099* |
| ENSG00000227398 | KIF9-AS1 | **1.278773648** | 5.80074011 | 12.67808734 | *6.021236521* |
| ENSG00000149571 | KIRREL3 | **-1.731542689** | 1.171125235 | 0.457725814 | *0.833579023* |
| ENSG00000170498 | KISS1 | **1.990977034** | 5.60840258 | 18.88548447 | *6.71201728* |
| ENSG00000116014 | KISS1R | **1.06998559** | 2.336278868 | 3.185176136 | *1.862684223* |
| ENSG00000157404 | KIT | **2.002956764** | 3.505522685 | 17.35767612 | *9.140034595* |
| ENSG00000133116 | KL | **1.211580536** | 3.611287155 | 7.612040053 | *5.133421002* |
| ENSG00000104892 | KLC3 | **1.289109976** | 0.5603239 | 5.466249906 | *0.946047995* |
| ENSG00000163884 | KLF15 | **1.363222531** | 1.424884988 | 5.002431945 | *2.007614099* |
| ENSG00000231160 | KLF3-AS1 | **1.317904033** | 0.354494467 | 0.979553318 | *0.345233226* |
| ENSG00000119138 | KLF9 | **1.232086769** | 0.814547731 | 2.343888237 | *1.2327716* |
| ENSG00000128607 | KLHDC10 | **-1.104888922** | 93.87311375 | 28.80963884 | *77.25599035* |
| ENSG00000130487 | KLHDC7B | **4.029413447** | 0.024297716 | 0.780169545 | *0.106422459* |
| ENSG00000197705 | KLHL14 | **1.031876946** | 0.451700039 | 15.14532987 | *0.787111767* |
| ENSG00000129437 | KLK14 | **2.174434106** | 0.215960391 | 0.874749386 | *0.212762619* |
| ENSG00000171346 | KRT15 | **3.06164011** | 0.026322873 | 0.483019029 | *0.099576531* |
| ENSG00000236670 | KRT18P5 | **-1.860088247** | 1.692305533 | 0.719047008 | *1.036012474* |
| ENSG00000204889 | KRT40 | **1.431042272** | 30.26224635 | 53.24845971 | *29.9332811* |
| ENSG00000224928 | KRT8P30 | **2.448718829** | 6.42800598 | 23.41357022 | *5.234518269* |
| ENSG00000233560 | KRT8P39 | **2.097925205** | 0.286903907 | 28.18072549 | *0.75794304* |
| ENSG00000174611 | KY | **-1.026868829** | 2.893280432 | 12.45573464 | *2.784582726* |
| ENSG00000154655 | L3MBTL4 | **2.467323467** | 0.718621924 | 2.726633513 | *0.803600738* |
| ENSG00000246366 | LACTB2-AS1 | **3.735290527** | 0.01052901 | 0.47122816 | *0.053284769* |
| ENSG00000101680 | LAMA1 | **-1.569434778** | 6.040923452 | 2.862732165 | *4.598863272* |
| ENSG00000172037 | LAMB2 | **1.371949294** | 17.38313182 | 29.24972908 | *19.51272463* |
| ENSG00000150457 | LATS2 | **2.240775388** | 0.410304448 | 1.939180734 | *0.154509313* |
| ENSG00000204381 | LAYN | **1.497028621** | 0.683149769 | 16.42307463 | *1.222927438* |
| ENSG00000179528 | LBX2 | **1.69718247** | 0.482777053 | 1.53448624 | *0.774806208* |
| ENSG00000136167 | LCP1 | **1.191162913** | 0.279668919 | 1.101107726 | *0.420780818* |
| ENSG00000179241 | LDLRAD3 | **-1.147821867** | 22.25731714 | 7.116803152 | *24.96791862* |
| ENSG00000138795 | LEF1 | **1.986706066** | 0.0201139 | 0.293098095 | *0.007033083* |

| | | | | | |
|---|---|---|---|---|---|
| ENSG00000116678 | LEPR | **2.966403369** | 0.322312761 | 5.058048401 | *0.663032205* |
| ENSG00000104660 | LEPROTL1 | **-1.124237366** | 65.96822811 | 19.86936995 | *61.08814103* |
| ENSG00000153012 | LGI2 | **2.093181297** | 2.612829488 | 8.312318292 | *2.114835393* |
| ENSG00000145685 | LHFPL2 | **1.739308147** | 1.163229047 | 12.55276518 | *1.617807382* |
| ENSG00000225329 | LHFPL3-AS2 | **-2.06952093** | 2.827785306 | 3.872255043 | *2.889215514* |
| ENSG00000273706 | LHX1 | **4.227102812** | 0 | 1.375156645 | *0.007436953* |
| ENSG00000064042 | LIMCH1 | **1.323207742** | 0.570638615 | 1.239408382 | *0.692141878* |
| ENSG00000231721 | LINC-PINT | **1.111765389** | 0.683863914 | 0.214645008 | *0.605392555* |
| ENSG00000225880 | LINC00115 | **1.523094148** | 5.921305342 | 8.548111821 | *6.530166087* |
| ENSG00000196668 | LINC00173 | **1.917917463** | 0.386601585 | 0.978750863 | *0.731408385* |
| ENSG00000231535 | LINC00278 | **-1.003749284** | 0.495773355 | 5.933516999 | *0.788656789* |
| ENSG00000203441 | LINC00449 | **1.780384816** | 10.8264016 | 3.577674383 | *7.33850177* |
| ENSG00000215117 | LINC00588 | **3.530391632** | 9.497732341 | 21.60666642 | *9.804776226* |
| ENSG00000248307 | LINC00616 | **1.038375348** | 0.124597457 | 2.994950734 | *0.183731852* |
| ENSG00000260941 | LINC00622 | **1.148281977** | 3.928989112 | 16.21799281 | *2.049630935* |
| ENSG00000261824 | LINC00662 | **1.030369979** | 14.23467371 | 21.04401757 | *11.60423981* |
| ENSG00000266256 | LINC00683 | **1.492501256** | 3.34693922 | 7.377176184 | *4.168082063* |
| ENSG00000254275 | LINC00824 | **3.930495858** | 0.819002672 | 12.0440346 | *2.744081162* |
| ENSG00000224805 | LINC00853 | **1.534585661** | 0.104393075 | 3.500629181 | *0* |
| ENSG00000232229 | LINC00865 | **1.367464017** | 10.1625246 | 19.54921079 | *8.866100884* |
| ENSG00000240875 | LINC00886 | **-1.64208912** | 2.393467679 | 4.582492741 | *1.883711291* |
| ENSG00000235703 | LINC00894 | **1.897112411** | 0.591041018 | 10.34693178 | *0.437927876* |
| ENSG00000263812 | LINC00908 | **1.981572787** | 1.476833608 | 5.557621982 | *2.15459228* |
| ENSG00000204054 | LINC00963 | **4.824636561** | 0.160891116 | 0.478292357 | *0.124659807* |
| ENSG00000253138 | LINC00967 | **-1.033956519** | 0 | 1.986103568 | *0.023093506* |
| ENSG00000223941 | LINC01014 | **-5.105392495** | 26.5140194 | 7.644394726 | *20.27980955* |
| ENSG00000249346 | LINC01016 | **1.607350268** | 0.181698666 | 0.118740298 | *0.277393322* |
| ENSG00000232715 | LINC01022 | **2.819396371** | 3.720649228 | 12.33856652 | *2.692572487* |
| ENSG00000222033 | LINC01124 | **1.987973128** | 3.620037544 | 15.89230243 | *6.430196184* |
| ENSG00000279873 | LINC01126 | **2.704313288** | 1.6048902 | 8.209691071 | *2.78804382* |
| ENSG00000267272 | LINC01140 | **1.131613889** | 0.011196414 | 9.396544335 | *0.030555828* |
| ENSG00000256124 | LINC01152 | **7.554915129** | 0.571421037 | 3.137446157 | *0.777745132* |
| ENSG00000280776 | LINC01202 | **1.150463176** | 0 | 4.985535657 | *0.126436404* |
| ENSG00000224957 | LINC01266 | **1.399500016** | 0.631088298 | 1.330427487 | *0.77793086* |
| ENSG00000260924 | LINC01311 | **4.02933589** | 2.540172302 | 6.896029491 | *2.411213464* |
| ENSG00000281327 | LINC01338 | **1.009222396** | 0.1149528 | 1.463636661 | *0.392982061* |
| ENSG00000228065 | LINC01515 | **1.488727182** | 1.292520876 | 4.21988526 | *1.748945746* |
| ENSG00000272138 | LINC01607 | **3.115003591** | 1.075787157 | 2.725522007 | *1.225792041* |
| ENSG00000283503 | LINC01902 | **-1.910118239** | 0.055973063 | 1.220515536 | *0.125251495* |
| ENSG00000260804 | LINC01963 | **1.901349883** | 15.42231376 | 3.559725935 | *12.69798117* |
| ENSG00000262585 | LINC01979 | **2.175199794** | 0.31552709 | 0.937303851 | *0.356749851* |
| ENSG00000238755 | LINC02006 | **-2.572705685** | 0.126911851 | 2.002891506 | *0.200628082* |
| ENSG00000231574 | LINC02015 | **2.586131791** | 0.632467241 | 0.46614527 | *0.297389843* |
| ENSG00000232233 | LINC02043 | **-1.041186371** | 0.154850623 | 0.868732111 | *0.366670916* |
| ENSG00000258884 | LINC02321 | **2.371971144** | 28.02706437 | 9.988758852 | *29.79367813* |
| ENSG00000248505 | LINC02380 | **-1.653247141** | 0.402378884 | 1.312683476 | *0.761014756* |
| ENSG00000248703 | LINC02415 | **2.393862347** | 27.67626284 | 9.329613714 | *24.21246949* |

| | | | | | |
|---|---|---|---|---|---|
| ENSG00000214039 | LINC02418 | **4.753730308** | 0.460410021 | 2.428648662 | *0.68888182* |
| ENSG00000250436 | LINC02499 | **2.600460163** | 0 | 3.344494651 | *0.073240038* |
| ENSG00000226453 | LINC02542 | **3.572550066** | 0.513470002 | 3.141908208 | *0.369922396* |
| ENSG00000223764 | LINC02593 | **3.741481009** | 0.079516741 | 0.745151037 | *0.029151495* |
| ENSG00000224222 | LINC02636 | **-1.278892459** | 0.138645362 | 2.200443197 | *0.485905443* |
| ENSG00000264404 | LINC02675 | **1.654845272** | 22.44517827 | 6.724519396 | *19.35479174* |
| ENSG00000231483 | LINC02678 | **1.507621771** | 1.455800144 | 3.427888133 | *1.860059643* |
| ENSG00000212719 | LINC02693 | **-2.909750651** | 6.776717351 | 14.93744355 | *7.65829469* |
| ENSG00000255406 | LINC02730 | **-1.086656825** | 11.0303174 | 2.444244864 | *8.624059377* |
| ENSG00000204241 | LINC02731 | **2.124733461** | 0.703737273 | 6.976677868 | *0.711360103* |
| ENSG00000187013 | LINC02875 | **-1.146016593** | 1.67924045 | 4.694389912 | *1.718698248* |
| ENSG00000166035 | LIPC | **1.312535036** | 2.218312748 | 3.677998449 | *8.013277069* |
| ENSG00000163898 | LIPH | **2.163056796** | 0.206313527 | 3.767921201 | *0.089456307* |
| ENSG00000189067 | LITAF | **1.367589505** | 1.308650088 | 4.101546223 | *2.006437433* |
| ENSG00000185621 | LMLN | **-1.405544466** | 3.042339215 | 1.087673276 | *3.475742336* |
| ENSG00000135363 | LMO2 | **1.811919029** | 30.12588295 | 69.93075301 | *24.5146777* |
| ENSG00000134013 | LOXL2 | **2.068040654** | 0.713247877 | 2.245421648 | *0.52917942* |
| ENSG00000213071 | LPAL2 | **4.100357946** | 0 | 35.76792719 | *0.040648727* |
| ENSG00000198121 | LPAR1 | **1.489296003** | 10.14552011 | 19.46589755 | *12.12846191* |
| ENSG00000064547 | LPAR2 | **1.46261536** | 0.109827668 | 0.205180501 | *0.179501513* |
| ENSG00000139679 | LPAR6 | **4.607367492** | 0 | 9.722776295 | *0* |
| ENSG00000121207 | LRAT | **-1.101920514** | 0.602145595 | 0.315488099 | *0.656584743* |
| ENSG00000173621 | LRFN4 | **-1.2113962** | 21.13155054 | 5.877459715 | *19.04086011* |
| ENSG00000168702 | LRP1B | **3.696146573** | 0.030555551 | 0.342972432 | *0.052316133* |
| ENSG00000010626 | LRRC23 | **-1.171561471** | 3.956620189 | 4.368853324 | *3.549480519* |
| ENSG00000114248 | LRRC31 | **1.921655845** | 0.687531173 | 1.834871641 | *0.60867008* |
| ENSG00000176809 | LRRC37A3 | **1.573798665** | 1.593795978 | 3.697844077 | *1.620699323* |
| ENSG00000148948 | LRRC4C | **3.52248477** | 0.172718768 | 2.187943144 | *0.102118402* |
| ENSG00000188906 | LRRK2 | **4.407514442** | 0 | 1.550972588 | *0.003322625* |
| ENSG00000162951 | LRRTM1 | **3.070869383** | 0.575270617 | 3.791990478 | *0.570707644* |
| ENSG00000198739 | LRRTM3 | **2.169940612** | 0.544552391 | 1.552985354 | *0.880006748* |
| ENSG00000185565 | LSAMP | **2.004283986** | 3.274182667 | 10.1537287 | *6.783650895* |
| ENSG00000168056 | LTBP3 | **1.511952269** | 6.206595223 | 12.49101894 | *6.562971303* |
| ENSG00000171357 | LURAP1 | **1.491132524** | 2.1784371 | 8.581070662 | *2.259501332* |
| ENSG00000187398 | LUZP2 | **1.73120951** | 0.268420033 | 6.471331743 | *0.331347568* |
| ENSG00000254087 | LYN | **1.40761839** | 0.369065588 | 2.517212487 | *0.682824375* |
| ENSG00000180155 | LYNX1 | **1.536020536** | 0.108663601 | 0.476467497 | *0.256956331* |
| ENSG00000150551 | LYPD1 | **-3.776703032** | 6.802069653 | 0.584391667 | *1.829540461* |
| ENSG00000159871 | LYPD5 | **1.154514874** | 0.341146689 | 0.616074768 | *0.28464898* |
| ENSG00000183833 | MAATS1 | **-1.493664697** | 1.080202788 | 0.469147316 | *1.396953468* |
| ENSG00000180660 | MAB21L1 | **2.301609651** | 2.86423753 | 9.033158375 | *2.480914578* |
| ENSG00000099866 | MADCAM1 | **1.055110558** | 0.313871951 | 0.527225666 | *0.362037335* |
| ENSG00000234456 | MAGI2-AS3 | **1.46624206** | 0.070616793 | 5.340077168 | *0.218412592* |
| ENSG00000111837 | MAK | **-1.127894889** | 0.359450118 | 0.324881101 | *0.260829992* |
| ENSG00000204706 | MAMDC2-AS1 | **1.163487178** | 0.134437731 | 0.274279534 | *0.183463191* |
| ENSG00000184384 | MAML2 | **1.197792162** | 6.51561916 | 10.22282105 | *5.264730341* |
| ENSG00000013619 | MAMLD1 | **-1.4290033** | 5.40824819 | 1.372485813 | *8.210058503* |

| | | | | | |
|---|---|---|---|---|---|
| ENSG00000267799 | MAN1A2P1 | **-1.597770304** | 22.13864177 | 9.147055405 | *17.92602342* |
| ENSG00000069535 | MAOB | **1.894245647** | 10.49978956 | 25.84968424 | *7.723518502* |
| ENSG00000197442 | MAP3K5 | **1.099170727** | 0.438834129 | 3.39505499 | *0.598587498* |
| ENSG00000142733 | MAP3K6 | **1.211158668** | 1.257439808 | 15.4745074 | *2.011891736* |
| ENSG00000156265 | MAP3K7CL | **1.102916543** | 2.301607672 | 3.408349575 | *1.968751895* |
| ENSG00000012983 | MAP4K5 | **-1.084697411** | 22.87809004 | 7.986532092 | *20.41209518* |
| ENSG00000259438 | MAPK6-DT | **1.849405746** | 1.661437424 | 5.594621287 | *1.884897618* |
| ENSG00000173838 | MARCHF10 | **1.588422506** | 0.089271807 | 3.702299449 | *0.086627884* |
| ENSG00000127241 | MASP1 | **4.062977407** | 0 | 2.085335387 | *0* |
| ENSG00000076770 | MBNL3 | **1.04982055** | 0.356611128 | 0.583436941 | *0.579227306* |
| ENSG00000172197 | MBOAT1 | **2.835346959** | 0.157919724 | 0.787461677 | *1.326729574* |
| ENSG00000177669 | MBOAT4 | **1.867114835** | 1.768234205 | 4.114368579 | *2.873194342* |
| ENSG00000076706 | MCAM | **1.790090194** | 0.188704182 | 0.725463452 | *0.278025154* |
| ENSG00000171444 | MCC | **1.335300538** | 5.10488527 | 11.0715962 | *5.515734317* |
| ENSG00000178460 | MCMDC2 | **-1.040200015** | 0.367189389 | 0.349175619 | *0.341943701* |
| ENSG00000175471 | MCTP1 | **1.092623639** | 0.528802614 | 4.898559637 | *0.694083577* |
| ENSG00000135272 | MDFIC | **2.371485364** | 0.855749673 | 2.871363576 | *1.186649366* |
| ENSG00000110492 | MDK | **1.069109609** | 2.778864962 | 4.150043516 | *6.745136775* |
| ENSG00000103313 | MEFV | **2.234254564** | 0.010902788 | 1.578238099 | *0.011716536* |
| ENSG00000225746 | MEG8 | **-1.430291455** | 7.505727924 | 3.940812322 | *5.763436046* |
| ENSG00000223403 | MEG9 | **-1.089924942** | 14.65687435 | 4.901410194 | *11.6802813* |
| ENSG00000145794 | MEGF10 | **2.905837563** | 0.008836896 | 0.874607509 | *0.023147619* |
| ENSG00000134138 | MEIS2 | **-2.039140839** | 31.4052422 | 6.946924176 | *29.93344873* |
| ENSG00000105419 | MEIS3 | **2.779945612** | 0.02799813 | 0.165841116 | *0.053137026* |
| ENSG00000141434 | MEP1B | **2.348338807** | 0.280441174 | 3.437038297 | *0.304447085* |
| ENSG00000103260 | METRN | **1.420678494** | 12.11415068 | 20.69205504 | *15.66181432* |
| ENSG00000176845 | METRNL | **4.229360351** | 0.09262128 | 1.668241594 | *0.022628318* |
| ENSG00000053328 | METTL24 | **1.440031142** | 1.820392088 | 14.53522996 | *1.78580504* |
| ENSG00000170439 | METTL7B | **2.033041191** | 0.774569558 | 2.496087391 | *1.200036036* |
| ENSG00000166482 | MFAP4 | **2.588947347** | 14.9689512 | 60.58432252 | *12.89438056* |
| ENSG00000140545 | MFGE8 | **2.299314003** | 3.993332302 | 13.91333269 | *6.868329288* |
| ENSG00000071073 | MGAT4A | **-1.399023111** | 131.5082673 | 62.53865037 | *100.2411842* |
| ENSG00000182050 | MGAT4C | **1.64762717** | 0.955692663 | 8.636823114 | *0.529299873* |
| ENSG00000167889 | MGAT5B | **3.158534734** | 0.031089119 | 16.95275879 | *0.095557321* |
| ENSG00000074416 | MGLL | **1.39089912** | 0.27842224 | 1.429208991 | *0.242464315* |
| ENSG00000169330 | MINAR1 | **1.012378281** | 0.506677547 | 0.745591714 | *0.582957227* |
| ENSG00000100253 | MIOX | **3.571500468** | 0.043102068 | 0.638223255 | *0.113536401* |
| ENSG00000174403 | MIR1-1HG-AS1 | **-1.020468226** | 22.61582637 | 7.74895202 | *34.30386464* |
| ENSG00000225206 | MIR137HG | **1.066050276** | 2.285458189 | 3.262600016 | *1.843280459* |
| ENSG00000207607 | MIR200A | **-1.185259903** | 75.88773244 | 25.09991614 | *69.12426588* |
| ENSG00000224592 | MIR3659HG | **2.889211984** | 0.177864342 | 2.318441339 | *0.302032816* |
| ENSG00000266325 | MIR3665 | **2.151360028** | 7.111683739 | 31.59929988 | *17.63184226* |
| ENSG00000223749 | MIR503HG | **1.069969611** | 7.386300353 | 10.34182191 | *6.262678616* |
| ENSG00000207561 | MIR635 | **1.296639453** | 77.41780212 | 133.07446 | *70.52128248* |
| ENSG00000207959 | MIR656 | **-1.28123299** | 44.05459629 | 17.11900676 | *32.41715549* |
| ENSG00000207703 | MIR7-2 | **-1.275364167** | 62.06152267 | 82.84294553 | *32.28020135* |
| ENSG00000267374 | MIR924HG | **1.131574901** | 6.471333873 | 15.45529605 | *10.68749424* |

| | | | | | |
|---|---|---|---|---|---|
| ENSG00000099812 | MISP | **-1.118911686** | 86.19540368 | 35.08315777 | *77.36302711* |
| ENSG00000185155 | MIXL1 | **1.949729844** | 0.745242214 | 6.960778103 | *0.812068387* |
| ENSG00000110917 | MLEC | **-1.176348495** | 197.4348337 | 70.61563738 | *198.6952843* |
| ENSG00000146147 | MLIP | **2.02032104** | 0.021570026 | 0.960509757 | *0.023884012* |
| ENSG00000231969 | MMADHC-DT | **1.086135048** | 3.424387675 | 32.28695223 | *3.864441392* |
| ENSG00000136297 | MMD2 | **2.571065894** | 0.105930631 | 0.524758296 | *0.062053149* |
| ENSG00000157227 | MMP14 | **2.894461196** | 0.174604847 | 4.577729422 | *0.252040953* |
| ENSG00000198598 | MMP17 | **-1.576784815** | 9.55871251 | 2.058062893 | *10.24527286* |
| ENSG00000087245 | MMP2 | **3.157181934** | 1.132056781 | 7.134456106 | *2.832074079* |
| ENSG00000137674 | MMP20 | **-1.027865304** | 164.363536 | 53.93616318 | *141.2471062* |
| ENSG00000173542 | MOB1B | **-1.100068509** | 196.6294665 | 63.22459007 | *167.0174587* |
| ENSG00000120162 | MOB3B | **3.049341402** | 1.478011745 | 34.77697046 | *2.859542537* |
| ENSG00000079931 | MOXD1 | **1.858343034** | 0.292824552 | 31.38177576 | *0.548966015* |
| ENSG00000150054 | MPP7 | **-1.038865846** | 12.31170008 | 7.948596959 | *10.47691401* |
| ENSG00000186732 | MPPED1 | **1.558923698** | 0.146666913 | 0.589461766 | *0.23609566* |
| ENSG00000135324 | MRAP2 | **2.251242677** | 1.985714547 | 8.274618904 | *3.076843155* |
| ENSG00000011028 | MRC2 | **3.547875987** | 0.012891362 | 0.293734142 | *0.023717447* |
| ENSG00000227877 | MRLN | **1.445225239** | 5.041993064 | 12.32329774 | *7.368852558* |
| ENSG00000227502 | MROCKI | **1.165085772** | 2.333880664 | 3.575882466 | *4.394973048* |
| ENSG00000166959 | MS4A8 | **1.968195495** | 0.186101751 | 4.836891219 | *0.373833233* |
| ENSG00000204410 | MSH5 | **-1.10199961** | 0.48395005 | 1.776440029 | *0.485166849* |
| ENSG00000102854 | MSLN | **-1.215340185** | 0.359260968 | 0.34362634 | *0.347984449* |
| ENSG00000163132 | MSX1 | **5.037445411** | 0 | 0.391042638 | *0.09757359* |
| ENSG00000065911 | MTHFD2 | **1.166655189** | 36.42047763 | 54.1583683 | *33.33491876* |
| ENSG00000173702 | MUC13 | **1.347070176** | 0.53933355 | 1.01960137 | *0.52879916* |
| ENSG00000169550 | MUC15 | **5.97157169** | 0 | 26.76143044 | *0.009277261* |
| ENSG00000101057 | MYBL2 | **1.0109275** | 10.32494302 | 14.18719594 | *25.99007362* |
| ENSG00000086967 | MYBPC2 | **1.265880334** | 0.463398523 | 0.770502498 | *0.990772861* |
| ENSG00000136997 | MYC | **2.893525102** | 0.224721484 | 7.987648235 | *0.520947443* |
| ENSG00000116990 | MYCL | **1.182272256** | 0.959641129 | 1.808792172 | *1.105051095* |
| ENSG00000074842 | MYDGF | **1.003725955** | 147.6114265 | 194.4038747 | *171.2822766* |
| ENSG00000109063 | MYH3 | **-1.742880691** | 0.570447612 | 0.846173736 | *0.614701464* |
| ENSG00000101335 | MYL9 | **-1.08568516** | 5.52290211 | 99.87541 | *8.490129632* |
| ENSG00000145949 | MYLK4 | **1.080668882** | 0.336388744 | 0.537328672 | *0.275517066* |
| ENSG00000041515 | MYO16 | **1.546235083** | 2.929472472 | 6.567033152 | *2.921061585* |
| ENSG00000164591 | MYOZ3 | **1.232946015** | 0.271493648 | 0.612816068 | *0.444736241* |
| ENSG00000139597 | N4BP2L1 | **2.020655125** | 0.456263194 | 4.224135406 | *0.693453834* |
| ENSG00000281026 | N4BP2L2-IT2 | **-1.008825121** | 1.206064906 | 0.563952164 | *0.814769413* |
| ENSG00000166886 | NAB2 | **1.174308965** | 12.9092349 | 19.78906718 | *9.004790761* |
| ENSG00000237886 | NALT1 | **-2.244563829** | 4.350922722 | 0.835365208 | *3.147775263* |
| ENSG00000231697 | NANOGP5 | **-3.389609855** | 1.285147447 | 9.602152728 | *0.395995642* |
| ENSG00000186310 | NAP1L3 | **1.731058811** | 1.002768238 | 2.410162602 | *1.136709969* |
| ENSG00000125814 | NAPB | **-1.115020198** | 40.48360381 | 12.25233545 | *42.25927574* |
| ENSG00000134440 | NARS1 | **-1.275228975** | 151.218914 | 42.11995207 | *139.2507429* |
| ENSG00000067798 | NAV3 | **1.777613265** | 1.11287337 | 8.68646436 | *0.988034608* |
| ENSG00000271425 | NBPF10 | **1.638397395** | 0.111263182 | 20.98776106 | *0.075071927* |
| ENSG00000270629 | NBPF14 | **2.501786174** | 0.122124513 | 1.739655173 | *0.104231553* |

| | | | | | |
|---|---|---|---|---|---|
| ENSG00000149294 | NCAM1 | **-1.460611235** | 26.3516816 | 6.408143885 | *25.6795219* |
| ENSG00000173376 | NDNF | **5.358146455** | 0.040697419 | 1.346101908 | *0.023280109* |
| ENSG00000104419 | NDRG1 | **-1.188241381** | 27.19989834 | 11.02320856 | *21.19333519* |
| ENSG00000164100 | NDST3 | **1.715101514** | 0.295824644 | 1.192009743 | *0.291189582* |
| ENSG00000138653 | NDST4 | **3.701983557** | 0.020658414 | 4.070008881 | *0.006496271* |
| ENSG00000185633 | NDUFA4L2 | **2.07832575** | 0.943075167 | 2.830004179 | *0.599298641* |
| ENSG00000183091 | NEB | **-1.574339012** | 0.143724299 | 0.151801558 | *0.157405695* |
| ENSG00000123119 | NECAB1 | **-2.452424582** | 4.278622746 | 1.970198002 | *5.51234397* |
| ENSG00000103154 | NECAB2 | **1.509735326** | 10.96220622 | 20.55027509 | *13.96098453* |
| ENSG00000130202 | NECTIN2 | **1.50415396** | 3.720052013 | 7.241026464 | *6.079286884* |
| ENSG00000069869 | NEDD4 | **1.405108725** | 1.770015401 | 13.31547492 | *1.984356323* |
| ENSG00000111859 | NEDD9 | **-1.192122809** | 2.868718652 | 4.334515779 | *1.594753626* |
| ENSG00000100285 | NEFH | **-2.714313145** | 2.164217814 | 1.830145307 | *2.491554069* |
| ENSG00000172260 | NEGR1 | **3.215815794** | 0.289692402 | 2.123086845 | *0.327371841* |
| ENSG00000165973 | NELL1 | **2.617465373** | 0.02727027 | 0.204578203 | *0.08437855* |
| ENSG00000235470 | NEURL1-AS1 | **3.870435322** | 0.131282662 | 2.142314102 | *0.453060402* |
| ENSG00000050030 | NEXMIF | **-2.13366519** | 15.52100032 | 2.386939334 | *11.84276707* |
| ENSG00000141905 | NFIC | **1.085333582** | 2.133234635 | 3.653907193 | *2.292373112* |
| ENSG00000144802 | NFKBIZ | **-1.632924837** | 15.05545011 | 4.221640305 | *8.687404654* |
| ENSG00000272145 | NFYC-AS1 | **1.157873356** | 6.11518042 | 9.667184883 | *7.654913081* |
| ENSG00000177551 | NHLH2 | **1.575432912** | 0.50354674 | 2.656820251 | *0.480396915* |
| ENSG00000188158 | NHS | **1.637940458** | 0.874673728 | 7.065684404 | *1.207177157* |
| ENSG00000135540 | NHSL1 | **1.074893516** | 16.37213192 | 23.17637076 | *13.616742* |
| ENSG00000135842 | NIBAN1 | **1.226060665** | 0.408823835 | 1.591214289 | *0.782600191* |
| ENSG00000116962 | NID1 | **1.34218595** | 1.630357101 | 14.12939096 | *4.263490353* |
| ENSG00000087303 | NID2 | **3.838476601** | 0.028430038 | 0.486298005 | *0.043655508* |
| ENSG00000170113 | NIPA1 | **-1.292150267** | 23.45000459 | 7.792745408 | *19.21600642* |
| ENSG00000185942 | NKAIN3 | **2.186434224** | 0.342469809 | 1.083548645 | *0.935579016* |
| ENSG00000101198 | NKAIN4 | **5.975998043** | 0.05540865 | 5.363827896 | *0.145842332* |
| ENSG00000229544 | NKX1-2 | **4.667064682** | 0 | 0.819117666 | *0.034573478* |
| ENSG00000163623 | NKX6-1 | **-1.330111761** | 477.8581814 | 122.3264807 | *486.7205991* |
| ENSG00000165066 | NKX6-3 | **2.895210855** | 29.03959937 | 141.0514913 | *26.11492755* |
| ENSG00000169760 | NLGN1 | **1.880315201** | 0.032654125 | 66.5452768 | *0.061817461* |
| ENSG00000146938 | NLGN4X | **3.679661025** | 0.036532766 | 72.16346273 | *0.050482308* |
| ENSG00000165246 | NLGN4Y | **1.58025762** | 0.723413588 | 1.419545327 | *0.97484185* |
| ENSG00000197696 | NMB | **1.520174175** | 5.539704152 | 10.78917808 | *9.15022164* |
| ENSG00000123609 | NMI | **1.340668748** | 1.518652563 | 3.325390993 | *1.70075619* |
| ENSG00000109255 | NMU | **1.903758172** | 0.272165648 | 5.806190701 | *0.856523317* |
| ENSG00000106100 | NOD1 | **1.065661814** | 1.817395118 | 3.774206035 | *2.675831978* |
| ENSG00000148400 | NOTCH1 | **-2.615333702** | 23.43056211 | 2.744707472 | *17.79843745* |
| ENSG00000134250 | NOTCH2 | **2.403900234** | 0.8590075 | 4.167199141 | *0.766580651* |
| ENSG00000104967 | NOVA2 | **1.095421883** | 0.973778681 | 2.786065074 | *0.683290201* |
| ENSG00000170485 | NPAS2 | **3.308110988** | 0.016291427 | 1.653971126 | *0.034181319* |
| ENSG00000056291 | NPFFR2 | **1.83617382** | 0.803093374 | 2.416017721 | *0.820122412* |
| ENSG00000168743 | NPNT | **1.007406887** | 9.775223563 | 12.96018795 | *14.7155008* |
| ENSG00000163273 | NPPC | **-1.512124572** | 20.540234 | 5.710872366 | *14.99224146* |
| ENSG00000113389 | NPR3 | **2.900240673** | 0.218318444 | 7.56037686 | *0.608485625* |

| | | | | | |
|---|---|---|---|---|---|
| ENSG00000171246 | NPTX1 | **2.677677965** | 0.217916321 | 3.3460067 | *0.239017248* |
| ENSG00000183971 | NPW | **2.037978993** | 15.22289521 | 42.48062076 | *14.90081563* |
| ENSG00000122585 | NPY | **1.691464546** | 2.081520478 | 5.074063479 | *4.762254486* |
| ENSG00000237187 | NR2F1-AS1 | **1.193728743** | 1.025354551 | 21.27212121 | *1.004183049* |
| ENSG00000151623 | NR3C2 | **2.35939567** | 0.096500723 | 2.375625081 | *0.164450408* |
| ENSG00000123358 | NR4A1 | **1.547099327** | 0.599575561 | 2.025433679 | *0.807577066* |
| ENSG00000116833 | NR5A2 | **3.623957605** | 0.016141318 | 0.286192514 | *0.010400964* |
| ENSG00000185737 | NRG3 | **1.691470375** | 0.998902363 | 2.768976378 | *1.134753468* |
| ENSG00000099250 | NRP1 | **3.499393407** | 0.530464622 | 4.020372913 | *0.75701776* |
| ENSG00000171119 | NRTN | **2.37064144** | 2.256543332 | 8.399131911 | *1.730403971* |
| ENSG00000168824 | NSG1 | **1.981014039** | 0.112991288 | 2.242049038 | *0.068186904* |
| ENSG00000170091 | NSG2 | **2.631171388** | 0.243341447 | 5.027226845 | *0.18901996* |
| ENSG00000205309 | NT5M | **1.657646627** | 0.328581989 | 0.930324562 | *0.379981904* |
| ENSG00000065320 | NTN1 | **-1.503472866** | 147.5019218 | 33.53104909 | *173.3736393* |
| ENSG00000074527 | NTN4 | **4.138613913** | 0.010944116 | 0.39357881 | *0.043093111* |
| ENSG00000148053 | NTRK2 | **2.538995856** | 0.008702537 | 18.40438688 | *0.007539167* |
| ENSG00000140538 | NTRK3 | **2.6437707** | 0.036276443 | 0.197258086 | *0.018125249* |
| ENSG00000122824 | NUDT10 | **1.072387834** | 1.201601952 | 1.733586695 | *1.505729748* |
| ENSG00000213965 | NUDT19 | **-1.192598422** | 13.64162227 | 4.215683594 | *19.34326231* |
| ENSG00000272325 | NUDT3 | **-1.064552021** | 87.60239761 | 28.07023039 | *73.57215287* |
| ENSG00000122584 | NXPH1 | **5.228196037** | 0.008847353 | 1.97169722 | *0.047582499* |
| ENSG00000182575 | NXPH3 | **1.976875756** | 0.41310139 | 15.16615722 | *0.730180562* |
| ENSG00000182379 | NXPH4 | **2.626400667** | 2.851808105 | 11.77180743 | *3.388482244* |
| ENSG00000184232 | OAF | **2.084291095** | 0.289298948 | 1.374537944 | *0.43201354* |
| ENSG00000122126 | OCRL | **-1.586582116** | 21.18392085 | 10.3962116 | *21.05531784* |
| ENSG00000118733 | OLFM3 | **4.42243846** | 0 | 0.335972648 | *0.00394608* |
| ENSG00000183801 | OLFML1 | **1.564265466** | 0.551250043 | 3.401042493 | *1.274148996* |
| ENSG00000162745 | OLFML2B | **2.656391873** | 1.319589912 | 5.466867302 | *1.653004368* |
| ENSG00000116774 | OLFML3 | **5.266410524** | 0.033128543 | 1.384189301 | *0.029012529* |
| ENSG00000184221 | OLIG1 | **1.666754624** | 25.83481518 | 55.97912866 | *38.63699335* |
| ENSG00000126861 | OMG | **1.597253828** | 0.157846443 | 0.699711228 | *0.284629776* |
| ENSG00000082556 | OPRK1 | **1.970007568** | 1.027586866 | 30.12432234 | *1.682686021* |
| ENSG00000221882 | OR3A2 | **-2.515691111** | 0.445419571 | 0.197066342 | *0.246487612* |
| ENSG00000197984 | OR51A8P | **1.69181455** | 0.429879833 | 2.34635072 | *0.416400795* |
| ENSG00000091651 | ORC6 | **1.097689382** | 4.342965045 | 6.391923842 | *8.218253795* |
| ENSG00000070882 | OSBPL3 | **1.328243952** | 0.348073779 | 0.949229288 | *0.501400698* |
| ENSG00000165899 | OTOGL | **2.79802454** | 0.008119554 | 2.857690211 | *0.032861916* |
| ENSG00000089723 | OTUB2 | **-1.219575634** | 21.08159424 | 6.075304029 | *14.7573433* |
| ENSG00000253738 | OTUD6B-AS1 | **-1.469786465** | 72.62713861 | 17.43964601 | *59.64385011* |
| ENSG00000115507 | OTX1 | **2.092320137** | 0.162167862 | 3.532732524 | *0.29833949* |
| ENSG00000187950 | OVCH1 | **3.542318566** | 0.03180377 | 8.768787248 | *0.099688559* |
| ENSG00000083454 | P2RX5 | **-2.220388888** | 0.242648142 | 0.346626135 | *0.375455766* |
| ENSG00000099957 | P2RX6 | **1.08000062** | 0.211816597 | 0.354548066 | *0.218993607* |
| ENSG00000169860 | P2RY1 | **1.141853269** | 1.269856688 | 1.905785914 | *1.59696821* |
| ENSG00000090530 | P3H2 | **-1.075980203** | 4.997572812 | 1.811455868 | *2.345278585* |
| ENSG00000110811 | P3H3 | **4.107289846** | 0.040700529 | 1.310579712 | *0.088960249* |
| ENSG00000158006 | PAFAH2 | **-1.368786194** | 27.68476087 | 7.688875368 | *23.22162996* |

| | | | | | |
|---|---|---|---|---|---|
| ENSG00000099260 | PALMD | **1.316451713** | 0.109218491 | 0.380984414 | *0.082303687* |
| ENSG00000073150 | PANX2 | **2.371490969** | 1.715785588 | 9.162182114 | *2.09941725* |
| ENSG00000116183 | PAPPA2 | **1.075591646** | 0.700476707 | 1.108244317 | *0.635493699* |
| ENSG00000116117 | PARD3B | **1.06392839** | 0.665234386 | 4.100799616 | *1.30291015* |
| ENSG00000173193 | PARP14 | **2.268830532** | 0.184733939 | 1.036317825 | *0.22217669* |
| ENSG00000138496 | PARP9 | **1.732083691** | 0.758244569 | 2.090225901 | *0.930999874* |
| ENSG00000152931 | PART1 | **3.350791684** | 0.033724356 | 0.555237756 | *0.052953902* |
| ENSG00000196092 | PAX5 | **1.088364219** | 5.577743447 | 8.558340668 | *7.72915859* |
| ENSG00000163346 | PBXIP1 | **1.019105866** | 258.7214832 | 341.2702125 | *214.2110337* |
| ENSG00000138650 | PCDH10 | **2.141681622** | 0.987233975 | 6.803957839 | *1.641139666* |
| ENSG00000113555 | PCDH12 | **6.547281928** | 0.281493759 | 193.3676754 | *0.102674491* |
| ENSG00000118946 | PCDH17 | **2.080416464** | 4.637909076 | 14.03231043 | *5.119375552* |
| ENSG00000184226 | PCDH9 | **1.967110013** | 1.674882941 | 13.37105832 | *2.150072662* |
| ENSG00000255408 | PCDHA3 | **1.258219925** | 0.567446254 | 7.682766073 | *0.587353729* |
| ENSG00000171815 | PCDHB1 | **1.281987562** | 0.20496361 | 2.529915013 | *0.196307804* |
| ENSG00000113209 | PCDHB5 | **2.42324472** | 0.063095081 | 0.688559126 | *0.083421251* |
| ENSG00000081853 | PCDHGA2 | **1.239510727** | 0.666582213 | 1.150176847 | *0.410130272* |
| ENSG00000179715 | PCED1B | **1.95899648** | 0.196376402 | 0.621405782 | *0.326937125* |
| ENSG00000100889 | PCK2 | **1.421778271** | 4.423487293 | 8.264833391 | *4.209448698* |
| ENSG00000163710 | PCOLCE2 | **2.02318619** | 0.061330279 | 0.465795631 | *0.25894316* |
| ENSG00000248485 | PCP4L1 | **1.186070183** | 2.451849477 | 7.827879847 | *4.344761176* |
| ENSG00000125851 | PCSK2 | **-1.342358698** | 175.5513795 | 46.01866653 | *202.2521487* |
| ENSG00000099139 | PCSK5 | **1.448716558** | 0.435644822 | 2.427388416 | *0.939953831* |
| ENSG00000203497 | PDCD4-AS1 | **1.20308315** | 10.61261976 | 37.24544995 | *17.56307725* |
| ENSG00000112541 | PDE10A | **2.064238344** | 0.035274147 | 0.560760798 | *0.049884247* |
| ENSG00000128655 | PDE11A | **1.631206929** | 0.181910006 | 9.692607702 | *0.183510893* |
| ENSG00000184588 | PDE4B | **1.542339738** | 0.077720738 | 0.164099882 | *0.195864877* |
| ENSG00000134853 | PDGFRA | **3.786529243** | 0.003965188 | 0.233664652 | *0.016343946* |
| ENSG00000004799 | PDK4 | **1.814307769** | 24.50566145 | 56.48316898 | *31.0847274* |
| ENSG00000154553 | PDLIM3 | **2.72565251** | 0.031201278 | 0.160331993 | *0.021705746* |
| ENSG00000186862 | PDZD7 | **-1.010543474** | 2.224247192 | 29.29065585 | *1.919915462* |
| ENSG00000187800 | PEAR1 | **1.845948369** | 0.096552834 | 0.292108081 | *0.070460172* |
| ENSG00000284395 | PERCC1 | **6.591842727** | 1.741058836 | 109.9700943 | *3.184783191* |
| ENSG00000114757 | PEX5L | **1.854721222** | 1.653421662 | 4.045363097 | *1.837582739* |
| ENSG00000158571 | PFKFB1 | **-1.28299365** | 0.677738487 | 54.05044958 | *0.472444624* |
| ENSG00000170525 | PFKFB3 | **1.017816371** | 2.25437239 | 4.942535006 | *2.314421075* |
| ENSG00000119630 | PGF | **-1.372304377** | 44.14805103 | 11.20966342 | *37.30055597* |
| ENSG00000277778 | PGM5P2 | **2.394369942** | 0.138288997 | 1.937481777 | *0.396686349* |
| ENSG00000225398 | PGM5P4 | **1.343154496** | 6.796333706 | 18.06891499 | *8.268824511* |
| ENSG00000102174 | PHEX | **2.566469632** | 0.071764417 | 0.5436364 | *0.052726444* |
| ENSG00000056487 | PHF21B | **2.100596279** | 0.029505215 | 4.737646759 | *0.065302528* |
| ENSG00000092621 | PHGDH | **1.356660825** | 24.82115635 | 41.25033402 | *23.62730198* |
| ENSG00000100100 | PIK3IP1 | **1.926397387** | 0.383557406 | 1.039717821 | *0.243740218* |
| ENSG00000141506 | PIK3R5 | **-1.275449533** | 0.415544451 | 21.80167086 | *0.595521181* |
| ENSG00000179761 | PIPOX | **-1.949778574** | 4.327154236 | 1.1662073 | *5.130143253* |
| ENSG00000090975 | PITPNM2 | **-2.651801754** | 3.268741582 | 0.400038896 | *2.849658279* |
| ENSG00000091622 | PITPNM3 | **-1.295009297** | 0.275369902 | 0.454046856 | *0.181318181* |

| | | | | | |
|---|---|---|---|---|---|
| ENSG00000197181 | PIWIL2 | **2.07448421** | 0.042753955 | 0.28289252 | *0.033161553* |
| ENSG00000170927 | PKHD1 | **1.343543598** | 0.057431576 | 0.1294818 | *0.086110761* |
| ENSG00000205038 | PKHD1L1 | **1.649203932** | 0.437558856 | 0.910664266 | *0.701200602* |
| ENSG00000135549 | PKIB | **1.685683789** | 10.43118349 | 22.10375938 | *10.79783813* |
| ENSG00000160447 | PKN3 | **2.68293453** | 0.294165808 | 1.698428375 | *0.793936536* |
| ENSG00000144837 | PLA1A | **2.394511387** | 0.267403094 | 12.03845461 | *0.495337141* |
| ENSG00000168907 | PLA2G4F | **3.174297761** | 0.035096608 | 0.869092226 | *0.065932998* |
| ENSG00000153246 | PLA2R1 | **1.227514677** | 0.085878429 | 0.549267559 | *0.091345792* |
| ENSG00000181690 | PLAG1 | **1.178475284** | 0.153858 | 0.300216141 | *0.219715656* |
| ENSG00000161714 | PLCD3 | **1.60969817** | 2.171241949 | 4.268987321 | *3.995253436* |
| ENSG00000075651 | PLD1 | **-1.441990431** | 3.177100875 | 0.912927376 | *2.90762396* |
| ENSG00000169499 | PLEKHA2 | **1.075928673** | 0.725019583 | 3.366195007 | *1.121383447* |
| ENSG00000153404 | PLEKHG4B | **1.236336678** | 0.18328143 | 0.667096009 | *0.15177899* |
| ENSG00000152527 | PLEKHH2 | **1.519012998** | 1.158853628 | 2.682161775 | *1.55417257* |
| ENSG00000147872 | PLIN2 | **-3.232663835** | 157.3675042 | 11.25448569 | *207.9029752* |
| ENSG00000117600 | PLPPR4 | **-1.638724627** | 2.143556204 | 1.569988868 | *1.454964863* |
| ENSG00000102024 | PLS3 | **-1.568379056** | 46.34592995 | 15.76063978 | *46.83907492* |
| ENSG00000100979 | PLTP | **1.886695141** | 0.447325943 | 1.258457356 | *0.752939815* |
| ENSG00000162877 | PM20D1 | **-2.249830932** | 11.00581705 | 6.712892403 | *13.51405405* |
| ENSG00000141682 | PMAIP1 | **2.400877783** | 41.47796606 | 141.4993438 | *38.35425771* |
| ENSG00000182013 | PNMA8A | **-1.115155951** | 96.3525055 | 29.70263609 | *88.93929275* |
| ENSG00000108439 | PNPO | **-1.039799099** | 4.417416227 | 77.18820903 | *4.342973475* |
| ENSG00000128567 | PODXL | **-1.076285889** | 5.358099096 | 16.53346116 | *5.986779563* |
| ENSG00000115138 | POMC | **2.437298766** | 0.415268096 | 2.169994071 | *0.537359174* |
| ENSG00000105852 | PON3 | **2.303401506** | 0.151196315 | 1.373409509 | *0.311140319* |
| ENSG00000185668 | POU3F1 | **1.816824048** | 0.346396871 | 1.540123904 | *0.500920113* |
| ENSG00000184486 | POU3F2 | **-1.54647589** | 85.91057181 | 19.48888901 | *80.95846116* |
| ENSG00000152192 | POU4F1 | **2.308843561** | 1.117424955 | 3.976301591 | *2.174030392* |
| ENSG00000242551 | POU5F1P6 | **1.507878975** | 1.665974355 | 12.60568138 | *1.281839718* |
| ENSG00000106536 | POU6F2 | **3.360056471** | 0.023465696 | 2.027011748 | *0.027656267* |
| ENSG00000109819 | PPARGC1A | **1.991104468** | 0.5703706 | 3.263264661 | *0.670040394* |
| ENSG00000155846 | PPARGC1B | **2.613174146** | 0.097871114 | 0.435771416 | *0.281996258* |
| ENSG00000139220 | PPFIA2 | **2.756105149** | 0.077797857 | 1.120143319 | *0.072737556* |
| ENSG00000171497 | PPID | **-1.110963346** | 91.70873726 | 28.42548687 | *84.5262257* |
| ENSG00000137168 | PPIL1 | **-1.168343511** | 86.94662475 | 25.00826507 | *88.60854499* |
| ENSG00000106341 | PPP1R17 | **7.795026691** | 0.060907437 | 21.97925155 | *0.019153028* |
| ENSG00000184203 | PPP1R2 | **-1.350823558** | 39.79252629 | 23.51035896 | *36.818488* |
| ENSG00000074211 | PPP2R2C | **2.446917626** | 0.178018799 | 5.767531804 | *0.065316358* |
| ENSG00000177133 | PRDM16-DT | **-1.164604029** | 3.538395281 | 6.289169635 | *2.820872006* |
| ENSG00000138738 | PRDM5 | **1.563506714** | 1.377998022 | 3.034862846 | *1.633563096* |
| ENSG00000152784 | PRDM8 | **4.399245933** | 0.004233446 | 0.600074939 | *0.034061008* |
| ENSG00000164256 | PRDM9 | **3.873741171** | 0.011361188 | 1.417760708 | *0.0641064* |
| ENSG00000141391 | PRELID3A | **1.49991724** | 0.227629934 | 0.51440159 | *0.621003361* |
| ENSG00000124126 | PREX1 | **1.67587228** | 1.080447122 | 2.202919202 | *1.460159684* |
| ENSG00000116690 | PRG4 | **2.679523018** | 0.030178516 | 0.280799184 | *0.01607836* |
| ENSG00000163637 | PRICKLE2 | **1.457441557** | 0.619447595 | 2.431941696 | *0.81101821* |
| ENSG00000162409 | PRKAA2 | **-1.301649319** | 18.76983048 | 5.266314313 | *16.79892265* |

| | | | | | |
|---|---|---|---|---|---|
| ENSG00000106617 | PRKAG2 | **-1.247480015** | 21.19283609 | 6.550788317 | *17.70146368* |
| ENSG00000171132 | PRKCE | **1.284790668** | 7.576589113 | 14.16732448 | *8.02702903* |
| ENSG00000185532 | PRKG1 | **1.889257124** | 0.026707443 | 2.827650304 | *0.025310364* |
| ENSG00000138669 | PRKG2 | **2.901953553** | 0.025940709 | 6.459981732 | *0.06868224* |
| ENSG00000185345 | PRKN | **1.259029894** | 0.443503887 | 0.748152374 | *0.460627041* |
| ENSG00000113494 | PRLR | **2.593482921** | 0.03048041 | 0.189011583 | *0.029003263* |
| ENSG00000115718 | PROC | **-1.211175819** | 35.7800016 | 10.92553955 | *56.8245678* |
| ENSG00000167525 | PROCA1 | **1.295987179** | 0.825992761 | 1.28288072 | *1.197666076* |
| ENSG00000169618 | PROKR1 | **3.935774744** | 0.015384 | 4.816688384 | *0.015262359* |
| ENSG00000007062 | PROM1 | **2.672604875** | 0.055694422 | 1.019854123 | *0.283050453* |
| ENSG00000184500 | PROS1 | **2.941781731** | 0.995971113 | 5.309114924 | *1.172238481* |
| ENSG00000148426 | PROSER2 | **3.476004645** | 0.199563651 | 1.653835071 | *0.152489824* |
| ENSG00000141127 | PRPSAP2 | **-1.255888774** | 12.72806159 | 5.895688646 | *12.65083318* |
| ENSG00000176532 | PRR15 | **1.145668711** | 29.92684575 | 41.958779 | *29.22693324* |
| ENSG00000224383 | PRR29 | **1.269328942** | 0.222746075 | 2.070860292 | *0.315934345* |
| ENSG00000010438 | PRSS3 | **2.79740031** | 0.198178818 | 24.97309401 | *0.506714771* |
| ENSG00000166450 | PRTG | **1.175551652** | 4.25764978 | 6.644594838 | *3.833180794* |
| ENSG00000105227 | PRX | **1.361891445** | 0.775171451 | 1.749455393 | *1.090167741* |
| ENSG00000135069 | PSAT1 | **1.574451668** | 98.53859722 | 194.3313616 | *82.34205391* |
| ENSG00000146005 | PSD2 | **1.621189926** | 0.203785617 | 1.005463542 | *0.148182152* |
| ENSG00000156011 | PSD3 | **-1.213156583** | 13.91014017 | 102.3020023 | *11.11946173* |
| ENSG00000159792 | PSKH1 | **-1.542749314** | 105.7222426 | 23.7291117 | *114.1589018* |
| ENSG00000140368 | PSTPIP1 | **1.955456627** | 0.365554385 | 2.868214699 | *0.572802318* |
| ENSG00000152229 | PSTPIP2 | **1.42876671** | 0.249113229 | 12.85501273 | *0.452082756* |
| ENSG00000244694 | PTCHD4 | **1.112165633** | 5.050773086 | 7.914853472 | *4.246570504* |
| ENSG00000171522 | PTGER4 | **2.62676466** | 2.27497002 | 9.184424626 | *1.712738862* |
| ENSG00000134247 | PTGFRN | **1.789881617** | 7.544822223 | 20.59545635 | *9.178207551* |
| ENSG00000124212 | PTGIS | **3.701583734** | 0.020464316 | 5.050683696 | *0.043261165* |
| ENSG00000087494 | PTHLH | **1.37509591** | 0.243479538 | 8.821172532 | *0.262919853* |
| ENSG00000184489 | PTP4A3 | **1.746799387** | 3.35086002 | 7.538959142 | *3.690099218* |
| ENSG00000152104 | PTPN14 | **1.268565449** | 0.569599237 | 1.142831956 | *0.286042723* |
| ENSG00000127329 | PTPRB | **1.627094604** | 0.019704477 | 3.751871943 | *0.025080632* |
| ENSG00000225706 | PTPRD-AS1 | **3.161249355** | 0.156221662 | 1.416698814 | *0.13478141* |
| ENSG00000144724 | PTPRG | **3.417039678** | 0.081635858 | 0.603283751 | *0.092882874* |
| ENSG00000139304 | PTPRQ | **4.771053716** | 0 | 0.426446102 | *0.003723162* |
| ENSG00000105426 | PTPRS | **1.64269172** | 0.138350522 | 0.560319752 | *0.123805907* |
| ENSG00000106278 | PTPRZ1 | **1.003442657** | 10.48060713 | 13.8549636 | *12.21201814* |
| ENSG00000225180 | PVALEF | **1.100972982** | 1.161552358 | 1.674694712 | *1.492421213* |
| ENSG00000100504 | PYGL | **1.100965625** | 0.251189176 | 7.21729978 | *0.37879836* |
| ENSG00000171016 | PYGO1 | **1.041326627** | 0.75181069 | 1.948708611 | *0.900076736* |
| ENSG00000131096 | PYY | **1.768789379** | 0.882759603 | 2.315285897 | *0.748814627* |
| ENSG00000041353 | RAB27B | **-1.106790454** | 3.899352 | 1.793474233 | *4.545313213* |
| ENSG00000137502 | RAB30 | **-1.149319774** | 3.41806234 | 1.884597778 | *2.428553498* |
| ENSG00000168461 | RAB31 | **1.061999104** | 1.35114306 | 2.277402947 | *2.306774267* |
| ENSG00000172780 | RAB43 | **1.705228831** | 0.152502995 | 0.830645139 | *0.252755502* |
| ENSG00000166128 | RAB8B | **-1.061462851** | 48.87347093 | 16.10242241 | *37.45488096* |
| ENSG00000123570 | RAB9B | **-1.004168804** | 52.2558967 | 17.18665687 | *51.0707267* |

| | | | | | |
|---|---|---|---|---|---|
| ENSG00000151164 | RAD9B | **-2.152438893** | 0.153098167 | 8.058635761 | *0.12758438* |
| ENSG00000203722 | RAET1G | **3.048665947** | 0.336871306 | 10.43869931 | *0.653967446* |
| ENSG00000175097 | RAG2 | **4.422295047** | 0 | 0.067380322 | *0.007136017* |
| ENSG00000039560 | RAI14 | **2.059444342** | 2.383548864 | 7.523801318 | *3.922860998* |
| ENSG00000204764 | RANBP17 | **1.623185402** | 0.072116621 | 0.175029175 | *0.14855773* |
| ENSG00000136237 | RAPGEF5 | **1.216017079** | 7.457897514 | 14.48557006 | *10.08802594* |
| ENSG00000108551 | RASD1 | **3.531018564** | 4.896912381 | 36.28952965 | *6.412857473* |
| ENSG00000172575 | RASGRP1 | **1.305978473** | 1.290743699 | 7.965323389 | *1.094469318* |
| ENSG00000281358 | RASSF1-AS1 | **1.299451341** | 2.292826916 | 22.72997088 | *3.481362368* |
| ENSG00000101265 | RASSF2 | **1.263057439** | 0.109258626 | 1.246548589 | *0.125108979* |
| ENSG00000167281 | RBFOX3 | **1.555253669** | 0.496276136 | 3.659149786 | *0.306311888* |
| ENSG00000213411 | RBM22P2 | **2.229361593** | 0.299120706 | 1.006748776 | *0.269488519* |
| ENSG00000151962 | RBM46 | **4.642113768** | 0.007624732 | 0.527990134 | *0.067048035* |
| ENSG00000138207 | RBP4 | **-1.159342117** | 16.0443524 | 5.233590803 | *8.929603639* |
| ENSG00000157110 | RBPMS | **1.445954669** | 0.382868813 | 0.740341441 | *0.47218357* |
| ENSG00000159200 | RCAN1 | **-1.915206705** | 9.279660217 | 3.962329818 | *8.887046756* |
| ENSG00000136144 | RCBTB1 | **-1.05573165** | 76.78681667 | 24.42354978 | *80.48953481* |
| ENSG00000189056 | RELN | **2.666693733** | 0.016389758 | 0.862179505 | *0.027196151* |
| ENSG00000165731 | RET | **-1.594597677** | 18.44197528 | 16.43050352 | *28.98647659* |
| ENSG00000183688 | RFLNB | **1.295969882** | 18.06094787 | 28.37534685 | *30.21453822* |
| ENSG00000090104 | RGS1 | **4.835686013** | 0 | 2.0921254 | *0.037028281* |
| ENSG00000076344 | RGS11 | **-1.614719442** | 1.084058923 | 15.64901189 | *1.821954554* |
| ENSG00000147509 | RGS20 | **3.653480154** | 0.448413317 | 3.770663651 | *0.90470148* |
| ENSG00000182732 | RGS6 | **-2.354922999** | 7.053330054 | 1.014822186 | *7.577488569* |
| ENSG00000186479 | RGS7BP | **1.11269062** | 0.62432077 | 2.740705709 | *0.620412283* |
| ENSG00000108370 | RGS9 | **1.500730069** | 10.15374429 | 18.94554129 | *9.547235471* |
| ENSG00000007384 | RHBDF1 | **1.879701213** | 0.074653054 | 0.660748268 | *0.186284795* |
| ENSG00000141314 | RHBDL3 | **1.741803233** | 0.53479432 | 11.02057731 | *1.158714221* |
| ENSG00000168421 | RHOH | **3.402509425** | 0.024847463 | 0.260006386 | *0.0430452* |
| ENSG00000254389 | RHPN1-AS1 | **1.699486576** | 2.355164201 | 5.871142606 | *3.813400758* |
| ENSG00000188026 | RILPL1 | **1.582932244** | 0.670748802 | 1.378886683 | *0.764779213* |
| ENSG00000101098 | RIMS4 | **1.935759254** | 2.470923525 | 8.854642937 | *5.937902183* |
| ENSG00000042062 | RIPOR3 | **3.465727533** | 0.009972824 | 0.751318878 | *0.012779586* |
| ENSG00000183145 | RIPPLY3 | **1.378529682** | 8.492781266 | 16.92335544 | *24.34639437* |
| ENSG00000139405 | RITA1 | **-1.023985908** | 29.46110909 | 9.428683741 | *32.45726053* |
| ENSG00000107018 | RLN1 | **1.510176241** | 0.592945366 | 8.732374664 | *1.010974202* |
| ENSG00000107014 | RLN2 | **1.008840178** | 1.688003807 | 7.175450193 | *2.704008322* |
| ENSG00000255794 | RMST | **2.873684731** | 0.012238271 | 0.587301346 | *0.014924135* |
| ENSG00000201078 | RN7SKP214 | **1.058520236** | 7.877413277 | 12.58962915 | *11.79951982* |
| ENSG00000240250 | RN7SL541P | **-1.549622412** | 11.10251408 | 2.947831457 | *7.614427869* |
| ENSG00000101695 | RNF125 | **1.736004611** | 0.36516793 | 5.463949973 | *0.449968139* |
| ENSG00000170153 | RNF150 | **1.421500192** | 14.05885549 | 25.57396672 | *9.199166548* |
| ENSG00000141576 | RNF157 | **1.735361663** | 1.184782869 | 2.862809289 | *1.649245994* |
| ENSG00000141622 | RNF165 | **2.774165337** | 0.355958584 | 14.02295421 | *0.392723505* |
| ENSG00000165188 | RNF183 | **3.193509962** | 0.016827732 | 1.512878368 | *0.119358094* |
| ENSG00000158286 | RNF207 | **-1.481596937** | 0.856230298 | 1.06315369 | *1.177518601* |
| ENSG00000215277 | RNF212B | **1.179098885** | 0.663217062 | 0.780171144 | *0.563014613* |

| | | | | | |
|---|---|---|---|---|---|
| ENSG00000105982 | RNF32 | **-1.040196758** | 1.566403402 | 0.521790787 | *1.319176934* |
| ENSG00000238365 | RNU7-57P | **-1.524589547** | 57.73823703 | 9.535631331 | *40.33949595* |
| ENSG00000270722 | RNVU1-31 | **4.009812156** | 0.44298884 | 5.102232801 | *2.107687904* |
| ENSG00000147403 | RPL10 | **-1.146705953** | 115.7960868 | 45.83402089 | *129.8893919* |
| ENSG00000178464 | RPL10P16 | **-1.574485119** | 5.037571082 | 3.453866293 | *4.262705621* |
| ENSG00000237169 | RPL12P27 | **-2.755286842** | 2.926872504 | 17.25646031 | *0.825623331* |
| ENSG00000240270 | RPL12P37 | **4.222751293** | 0.132799659 | 2.897795348 | *0.250873396* |
| ENSG00000227939 | RPL3P2 | **-1.153410057** | 3.792979605 | 1.503490064 | *2.925663508* |
| ENSG00000225093 | RPL3P7 | **-1.829581591** | 2.398288379 | 0.841430493 | *1.041890702* |
| ENSG00000141425 | RPRD1A | **-1.168103166** | 83.91880418 | 25.01429186 | *79.97751504* |
| ENSG00000177519 | RPRM | **1.963749551** | 4.779666435 | 12.92303394 | *6.82946898* |
| ENSG00000220848 | RPS18P9 | **-1.523143104** | 27.54760996 | 19.87578427 | *27.48866336* |
| ENSG00000198208 | RPS6KL1 | **-1.289136633** | 10.20359857 | 8.503051488 | *10.07115028* |
| ENSG00000243499 | RPS6P21 | **-2.098115839** | 2.485683097 | 1.980647694 | *2.245897351* |
| ENSG00000216471 | RPSAP43 | **2.063405503** | 0.652584492 | 3.572375143 | *0.466522609* |
| ENSG00000048392 | RRM2B | **-1.026746694** | 72.13417494 | 23.67986487 | *42.8760537* |
| ENSG00000160208 | RRP1B | **-1.585746598** | 51.48159663 | 11.95450062 | *50.14399178* |
| ENSG00000132026 | RTBDN | **1.638600008** | 1.47676969 | 14.63188565 | *2.156380884* |
| ENSG00000254656 | RTL1 | **-1.901208105** | 0.967815317 | 5.702988475 | *1.526610496* |
| ENSG00000186907 | RTN4RL2 | **1.241419347** | 121.5195738 | 187.0100216 | *177.2067334* |
| ENSG00000188011 | RTP5 | **-3.230794005** | 33.24382018 | 2.279582195 | *17.93015447* |
| ENSG00000182631 | RXFP3 | **-1.031376751** | 86.39429569 | 125.6789088 | *69.45582242* |
| ENSG00000163191 | S100A11 | **2.663078358** | 66.15451466 | 277.7897793 | *45.48453533* |
| ENSG00000171643 | S100Z | **2.073954171** | 0.1981741 | 14.50385437 | *0.509642696* |
| ENSG00000170989 | S1PR1 | **5.400114394** | 0 | 137.2811662 | *0* |
| ENSG00000213694 | S1PR3 | **1.90626387** | 0.146327781 | 0.675605584 | *0.190031286* |
| ENSG00000256463 | SALL3 | **5.040691072** | 0.003731146 | 0.211480549 | *0.006288806* |
| ENSG00000187634 | SAMD11 | **2.017013874** | 1.062655569 | 3.008256786 | *0.82527186* |
| ENSG00000167100 | SAMD14 | **2.189994223** | 0.024145998 | 0.114983086 | *0.017547245* |
| ENSG00000203727 | SAMD5 | **-1.465646632** | 124.2692475 | 31.16189515 | *136.0719088* |
| ENSG00000155307 | SAMSN1 | **2.126503977** | 0.065910873 | 0.249039173 | *0.085124524* |
| ENSG00000228956 | SATB1-AS1 | **2.056989254** | 0.05007716 | 14.80093615 | *0.047929464* |
| ENSG00000119042 | SATB2 | **1.532366797** | 0.083821685 | 0.22812693 | *0.138895711* |
| ENSG00000188659 | SAXO2 | **-1.407116594** | 7.899558325 | 2.147334684 | *6.700306586* |
| ENSG00000145284 | SCD5 | **-1.194899004** | 104.9195802 | 29.84777245 | *135.3573931* |
| ENSG00000105711 | SCN1B | **2.809068525** | 0.017327896 | 0.953130152 | *0.025025708* |
| ENSG00000136546 | SCN7A | **1.378393008** | 0.206588498 | 15.83256825 | *0.292276124* |
| ENSG00000169432 | SCN9A | **-1.205440767** | 1.923573062 | 0.584156389 | *2.002570961* |
| ENSG00000196951 | SCOC-AS1 | **-1.026767363** | 2.065083359 | 0.848620998 | *1.797868007* |
| ENSG00000261678 | SCRT1 | **1.338293236** | 28.52664059 | 47.11534884 | *32.4268415* |
| ENSG00000146197 | SCUBE3 | **4.9308822** | 0.258761802 | 5.696539745 | *0.243332462* |
| ENSG00000162512 | SDC3 | **-1.237678186** | 8.541255075 | 26.82311797 | *8.936805044* |
| ENSG00000069188 | SDK2 | **1.261480782** | 3.829247863 | 8.534700184 | *2.812190288* |
| ENSG00000166562 | SEC11C | **1.065611642** | 162.9941693 | 224.2082645 | *176.7007897* |
| ENSG00000214491 | SEC14L6 | **3.485782045** | 0.146408407 | 4.091385039 | *0.30131981* |
| ENSG00000141574 | SECTM1 | **1.089204926** | 6.884728458 | 123.1417533 | *10.45030934* |
| ENSG00000091490 | SEL1L3 | **2.435002268** | 2.714701538 | 10.09093281 | *2.327177076* |

| | | | | | |
|---|---|---|---|---|---|
| ENSG00000188404 | SELL | **4.00498116** | 0.011256135 | 5.23415416 | *0.030012235* |
| ENSG00000110876 | SELPLG | **1.680245678** | 0.163130056 | 5.311276329 | *0.201477642* |
| ENSG00000153993 | SEMA3D | **2.034169775** | 0.527759728 | 1.526496819 | *0.532408109* |
| ENSG00000170381 | SEMA3E | **4.800107835** | 0.020870151 | 0.620350361 | *0.01097318* |
| ENSG00000112902 | SEMA5A | **2.024444671** | 6.733492875 | 19.21104313 | *9.85596748* |
| ENSG00000138758 | SEPTIN11 | **-1.135256831** | 68.05637685 | 20.59449362 | *70.57816972* |
| ENSG00000168528 | SERINC2 | **1.039239019** | 5.090389355 | 15.37441647 | *6.802165813* |
| ENSG00000099937 | SERPIND1 | **1.208940632** | 1.074677381 | 11.71243765 | *2.681319266* |
| ENSG00000106366 | SERPINE1 | **4.679257835** | 0 | 3.645760413 | *0.108357623* |
| ENSG00000135919 | SERPINE2 | **1.084932018** | 11.92387407 | 17.4447857 | *14.97003105* |
| ENSG00000167711 | SERPINF2 | **1.561439793** | 3.73003047 | 7.448018351 | *5.802835477* |
| ENSG00000149257 | SERPINH1 | **1.194930603** | 4.542269146 | 15.30315068 | *5.003379463* |
| ENSG00000180440 | SERTM1 | **-1.495731679** | 1.821126184 | 3.932697517 | *1.636025664* |
| ENSG00000260802 | SERTM2 | **3.17221111** | 8.051981747 | 50.1024952 | *10.127639* |
| ENSG00000149212 | SESN3 | **1.327266852** | 115.1982519 | 189.7260317 | *92.18191743* |
| ENSG00000104332 | SFRP1 | **1.141426569** | 23.33630338 | 57.82238313 | *38.25404427* |
| ENSG00000107819 | SFXN3 | **-1.247395602** | 20.4154529 | 101.4667794 | *17.04604691* |
| ENSG00000231304 | SGO1-AS1 | **1.233993026** | 2.602451664 | 21.86109455 | *2.488165156* |
| ENSG00000167037 | SGSM1 | **-1.668648199** | 2.379987982 | 3.194090841 | *2.277898115* |
| ENSG00000189410 | SH2D5 | **1.929321857** | 0.057369246 | 2.01398911 | *0.199274585* |
| ENSG00000100092 | SH3BP1 | **1.027078719** | 1.83937597 | 2.64009074 | *2.59267839* |
| ENSG00000130147 | SH3BP4 | **1.106187464** | 0.417566046 | 0.656455377 | *0.838126938* |
| ENSG00000131370 | SH3BP5 | **2.804046666** | 0.770224456 | 4.994869395 | *1.256932632* |
| ENSG00000147010 | SH3KBP1 | **-1.139303228** | 30.76727458 | 9.470332713 | *38.94507336* |
| ENSG00000174705 | SH3PXD2B | **-1.098676454** | 32.11071584 | 11.58435706 | *31.33060659* |
| ENSG00000172985 | SH3RF3 | **1.07065302** | 1.369753926 | 6.405197828 | *1.519583089* |
| ENSG00000125089 | SH3TC1 | **1.320903818** | 3.356616532 | 10.43298968 | *2.37811969* |
| ENSG00000129214 | SHBG | **1.976688155** | 0.091169292 | 1.320337558 | *0.152054222* |
| ENSG00000148082 | SHC3 | **2.013964367** | 1.204949475 | 5.859764824 | *0.734268401* |
| ENSG00000185634 | SHC4 | **1.262541946** | 0.47199778 | 0.838485313 | *0.35526026* |
| ENSG00000171241 | SHCBP1 | **1.121438819** | 3.277836732 | 6.611337629 | *5.499572312* |
| ENSG00000180730 | SHISA2 | **-1.761147293** | 25.52230581 | 5.411882425 | *24.90491319* |
| ENSG00000234965 | SHISA8 | **4.034377135** | 0.036490652 | 2.494495189 | *0.128035364* |
| ENSG00000138944 | SHISAL1 | **1.835941536** | 0.513096707 | 3.648082847 | *1.137347939* |
| ENSG00000182199 | SHMT2 | **1.034976666** | 15.17811136 | 20.54824715 | *15.8090927* |
| ENSG00000215475 | SIAH3 | **4.54485747** | 0.019556135 | 0.888225658 | *0.034963483* |
| ENSG00000126778 | SIX1 | **3.142083836** | 0.030359866 | 10.59834302 | *0.112391701* |
| ENSG00000170577 | SIX2 | **1.928360463** | 10.82514284 | 26.14720591 | *26.01952683* |
| ENSG00000177045 | SIX5 | **1.516660392** | 3.76895465 | 7.252746985 | *3.844472406* |
| ENSG00000113558 | SKP1 | **-1.430345173** | 45.61480475 | 25.60283784 | *45.20273971* |
| ENSG00000140199 | SLC12A6 | **-1.064730428** | 11.22802632 | 7.15550115 | *9.312583901* |
| ENSG00000088386 | SLC15A1 | **-2.376159145** | 1.367150654 | 5.866080323 | *1.448482689* |
| ENSG00000147100 | SLC16A2 | **2.352011348** | 8.683863939 | 30.47412706 | *12.82408455* |
| ENSG00000118596 | SLC16A7 | **1.300580423** | 3.112314459 | 4.857899031 | *3.497218737* |
| ENSG00000124568 | SLC17A1 | **1.317764381** | 1.184558152 | 17.14945441 | *3.978574059* |
| ENSG00000091664 | SLC17A6 | **3.29058233** | 0.045056054 | 3.044304948 | *0.080643486* |
| ENSG00000165646 | SLC18A2 | **4.93877487** | 0.328371975 | 7.417590738 | *0.635366019* |

| | | | | | |
|---|---|---|---|---|---|
| ENSG00000106688 | SLC1A1 | **-1.136082117** | 1.076194135 | 0.442363833 | *1.111030884* |
| ENSG00000110436 | SLC1A2 | **3.270737721** | 0.002768228 | 3.38087295 | *0.002242232* |
| ENSG00000105281 | SLC1A5 | **1.513966318** | 0.543126442 | 1.137830253 | *1.301032552* |
| ENSG00000168065 | SLC22A11 | **1.571455132** | 1.150581163 | 2.270483467 | *0.859702541* |
| ENSG00000197208 | SLC22A4 | **1.8552556** | 1.000386321 | 2.818234284 | *1.200760242* |
| ENSG00000155886 | SLC24A2 | **-1.540700845** | 1.373266919 | 1.446957209 | *0.832796897* |
| ENSG00000147454 | SLC25A37 | **-1.002919761** | 16.06564892 | 6.680240415 | *17.23498879* |
| ENSG00000077713 | SLC25A43 | **-1.698454107** | 9.268163965 | 2.044917523 | *9.944314827* |
| ENSG00000169100 | SLC25A6 | **-1.428108162** | 690.2138309 | 172.7518554 | *777.2858947* |
| ENSG00000091137 | SLC26A4 | **3.485617032** | 0.279790138 | 2.942308406 | *0.246967701* |
| ENSG00000167114 | SLC27A4 | **-1.418745396** | 113.5511842 | 111.0900652 | *113.6613347* |
| ENSG00000146411 | SLC2A12 | **2.363971765** | 3.419784614 | 12.78064115 | *4.615981775* |
| ENSG00000151229 | SLC2A13 | **-1.096266733** | 75.19726774 | 36.58382511 | *76.3803175* |
| ENSG00000163581 | SLC2A2 | **-1.798959327** | 0.256761396 | 5.687517077 | *0.183911191* |
| ENSG00000115194 | SLC30A3 | **2.87905998** | 0.086466728 | 12.32992267 | *0.062299677* |
| ENSG00000101438 | SLC32A1 | **-2.238104318** | 57.4396998 | 8.333184366 | *37.74698121* |
| ENSG00000196376 | SLC35F1 | **2.057635387** | 2.04959619 | 5.843393795 | *5.270607157* |
| ENSG00000110660 | SLC35F2 | **1.334079935** | 0.481686739 | 4.505043781 | *0.637949362* |
| ENSG00000111371 | SLC38A1 | **7.177613931** | 0 | 2.922057532 | *0.02241958* |
| ENSG00000139540 | SLC39A5 | **1.347663623** | 1.132741696 | 2.373047807 | *3.053113498* |
| ENSG00000133065 | SLC41A1 | **-1.697912746** | 90.538418 | 18.46758908 | *82.96187964* |
| ENSG00000134802 | SLC43A3 | **1.01780242** | 1.513951069 | 2.876594158 | *3.346110883* |
| ENSG00000143036 | SLC44A3 | **1.04199563** | 10.08249981 | 22.75253021 | *12.7480572* |
| ENSG00000224081 | SLC44A3-AS1 | **1.667855027** | 0.363225696 | 1.767561951 | *0.410992333* |
| ENSG00000137968 | SLC44A5 | **1.714593341** | 1.135702661 | 9.292859197 | *1.160140767* |
| ENSG00000080493 | SLC4A4 | **2.394877886** | 0.242181755 | 1.25174555 | *0.392663548* |
| ENSG00000188687 | SLC4A5 | **-1.676492739** | 0.061207048 | 1.288383302 | *0.033814446* |
| ENSG00000256870 | SLC5A8 | **1.649778244** | 0.529417395 | 1.326720957 | *0.861848545* |
| ENSG00000197106 | SLC6A17 | **2.107296726** | 0.42471145 | 1.160765262 | *0.295720723* |
| ENSG00000103546 | SLC6A2 | **1.625378156** | 0.887531775 | 2.542443182 | *2.578413464* |
| ENSG00000108576 | SLC6A4 | **-1.915180925** | 4.566734341 | 1.417509326 | *2.612987293* |
| ENSG00000151012 | SLC7A11 | **3.982625545** | 3.276649668 | 35.18566465 | *1.591366075* |
| ENSG00000250033 | SLC7A11-AS1 | **4.710142227** | 0 | 0.540435605 | *0.007853569* |
| ENSG00000013293 | SLC7A14 | **1.188591416** | 2.978347515 | 21.4605251 | *3.657956467* |
| ENSG00000103257 | SLC7A5 | **3.755733203** | 2.785033506 | 24.6154064 | *1.845654419* |
| ENSG00000021488 | SLC7A9 | **-1.710509831** | 0.809786401 | 2.318585107 | *0.793105508* |
| ENSG00000183023 | SLC8A1 | **1.156009068** | 3.473082336 | 17.90897353 | *3.555850136* |
| ENSG00000118160 | SLC8A2 | **2.748400816** | 0.302857529 | 1.44272409 | *0.449741491* |
| ENSG00000100678 | SLC8A3 | **2.586432613** | 0.541274475 | 4.587882921 | *0.565277221* |
| ENSG00000227825 | SLC9A7P1 | **2.113316378** | 0.423023531 | 1.834216236 | *0.591705905* |
| ENSG00000172139 | SLC9C1 | **1.303660935** | 0.494804654 | 1.978008306 | *0.512495083* |
| ENSG00000174640 | SLCO2A1 | **3.828680194** | 0.035886648 | 0.966243205 | *0.04665383* |
| ENSG00000173930 | SLCO4C1 | **1.236148204** | 33.11318172 | 51.42096312 | *36.19910087* |
| ENSG00000137571 | SLCO5A1 | **1.523570449** | 0.920716965 | 1.865965798 | *0.695880156* |
| ENSG00000205045 | SLFN12L | **3.032965162** | 0.023605761 | 26.02621968 | *0.074937926* |
| ENSG00000145147 | SLIT2 | **1.357922991** | 0.178938057 | 1.186180072 | *0.17917584* |
| ENSG00000165300 | SLITRK5 | **3.651967225** | 0.051716227 | 0.485337287 | *0.036531502* |

| | | | | | |
|---|---|---|---|---|---|
| ENSG00000184564 | SLITRK6 | **1.837193167** | 42.77879507 | 99.79324222 | *78.83230899* |
| ENSG00000137834 | SMAD6 | **4.86371862** | 0.017818526 | 0.498237295 | *0.085613344* |
| ENSG00000226746 | SMCR5 | **1.349221603** | 0.493616847 | 51.77767636 | *0.409710169* |
| ENSG00000253457 | SMIM18 | **1.127693566** | 3.300891404 | 4.52800714 | *2.543794727* |
| ENSG00000256235 | SMIM3 | **4.607592399** | 0.059179951 | 1.60668387 | *0.909483694* |
| ENSG00000256162 | SMLR1 | **1.349916319** | 2.691041638 | 7.104090333 | *4.616757357* |
| ENSG00000128602 | SMO | **3.154274049** | 1.344518692 | 8.27406228 | *2.493858405* |
| ENSG00000088826 | SMOX | **1.597715948** | 1.057498423 | 4.665169708 | *1.222832655* |
| ENSG00000237194 | SNAI1P1 | **1.500180249** | 2.565736178 | 9.093771022 | *1.981197009* |
| ENSG00000064692 | SNCAIP | **5.569922459** | 0 | 1.063396817 | *0* |
| ENSG00000260260 | SNHG19 | **1.130942728** | 95.20667299 | 140.6890862 | *108.877615* |
| ENSG00000201700 | SNORD113-3 | **-2.421970515** | 68.96415081 | 6.645512362 | *63.60937878* |
| ENSG00000207063 | SNORD116-1 | **-2.04104867** | 18.95550081 | 69.97039271 | *17.41124099* |
| ENSG00000207197 | SNORD116-12 | **-2.325915836** | 23.9276651 | 10.37613295 | *15.09327417* |
| ENSG00000101298 | SNPH | **-1.190170443** | 63.20009165 | 19.70458005 | *37.43911872* |
| ENSG00000172164 | SNTB1 | **3.579014572** | 0.039953846 | 0.932212022 | *0.06403043* |
| ENSG00000235688 | SNTG2-AS1 | **-1.074104264** | 38.05141383 | 20.91634075 | *53.55170807* |
| ENSG00000225345 | SNX18P3 | **1.017273378** | 1.125137362 | 1.710827264 | *0.998122461* |
| ENSG00000234373 | SNX18P7 | **1.85866854** | 1.20346743 | 9.49973948 | *1.632143246* |
| ENSG00000109762 | SNX25 | **-1.482983183** | 24.12150419 | 6.13788691 | *23.63603435* |
| ENSG00000236809 | SNX25P1 | **1.694370699** | 1.583042086 | 4.419046933 | *2.754177029* |
| ENSG00000173548 | SNX33 | **1.194971661** | 3.217282311 | 8.005895867 | *3.533026232* |
| ENSG00000185338 | SOCS1 | **1.621562287** | 0.716051116 | 3.493417041 | *1.164624612* |
| ENSG00000120833 | SOCS2 | **1.649137469** | 0.234080249 | 2.809957431 | *0.410208843* |
| ENSG00000140263 | SORD | **-1.020813616** | 87.24773411 | 27.93724941 | *102.0476914* |
| ENSG00000181449 | SOX2 | **1.283630598** | 0.991170235 | 1.79427668 | *2.323673728* |
| ENSG00000227640 | SOX21-AS1 | **4.902622899** | 0 | 15.31987474 | *0.017939237* |
| ENSG00000124766 | SOX4 | **-1.120446398** | 54.68497225 | 17.11953888 | *84.88194279* |
| ENSG00000125398 | SOX9 | **3.447728781** | 0.229368262 | 1.710627403 | *0.25434148* |
| ENSG00000234494 | SP2-AS1 | **-1.501902809** | 1.041589974 | 8.7247751 | *0.99933803* |
| ENSG00000204335 | SP5 | **2.346381935** | 7.151042002 | 25.25259577 | *13.05557521* |
| ENSG00000196096 | SPAG16-DT | **1.128176756** | 3.310981174 | 4.791183372 | *2.25195433* |
| ENSG00000189419 | SPATA41 | **1.829495823** | 0.211225131 | 12.03518947 | *0.284655759* |
| ENSG00000161888 | SPC24 | **1.322479621** | 2.497899758 | 6.53245292 | *8.493319945* |
| ENSG00000176170 | SPHK1 | **2.172711191** | 0.027857545 | 0.223761891 | *0.024576393* |
| ENSG00000122711 | SPINK4 | **6.340056411** | 0 | 4.032231268 | *0.069749797* |
| ENSG00000107742 | SPOCK2 | **1.577765973** | 2.144868296 | 4.177076136 | *2.580872509* |
| ENSG00000159674 | SPON2 | **1.174072132** | 0.18676236 | 1.621062457 | *0.195216856* |
| ENSG00000136158 | SPRY2 | **3.543824576** | 0.707875493 | 7.389814131 | *0.81059289* |
| ENSG00000171621 | SPSB1 | **1.244472303** | 2.164189056 | 3.720198085 | *2.838448178* |
| ENSG00000196542 | SPTSSB | **1.623832389** | 4.515763704 | 11.76792246 | *6.840286782* |
| ENSG00000137767 | SQOR | **1.009133637** | 0.407238689 | 1.985826365 | *0.529427081* |
| ENSG00000146700 | SSC4D | **-1.45008229** | 0.772163407 | 5.024174714 | *0.57554389* |
| ENSG00000157005 | SST | **5.053400132** | 1355.657948 | 29963.74035 | *938.2751366* |
| ENSG00000217330 | SSXP10 | **4.503630229** | 0 | 1.503823173 | *0* |
| ENSG00000215875 | ST13P20 | **4.607123887** | 0 | 14457.23436 | *0.061235738* |
| ENSG00000232504 | ST3GAL5-AS1 | **1.653075421** | 2.833259078 | 5.251855865 | *3.615896084* |

| | | | | | |
|---|---|---|---|---|---|
| ENSG00000214188 | ST7-OT4 | **3.562396147** | 0.013974454 | 0.32331589 | *0.062967487* |
| ENSG00000101638 | ST8SIA5 | **1.60698223** | 0.560913661 | 4.766687912 | *0.844572783* |
| ENSG00000010327 | STAB1 | **1.220903738** | 0.172089404 | 0.346658638 | *0.187121923* |
| ENSG00000141750 | STAC2 | **4.244277406** | 0 | 0.635326057 | *0.071507637* |
| ENSG00000138378 | STAT4 | **2.429426722** | 0.104259256 | 0.485367867 | *0.135007497* |
| ENSG00000159167 | STC1 | **2.568989041** | 1.737730665 | 6.322427168 | *2.265987354* |
| ENSG00000113739 | STC2 | **1.524337587** | 32.06081127 | 61.84272214 | *43.24854416* |
| ENSG00000105889 | STEAP1B | **1.554365029** | 0.319908539 | 4.423118785 | *0.804225373* |
| ENSG00000115107 | STEAP3 | **-1.418171497** | 0.835686798 | 29.72459222 | *0.975403972* |
| ENSG00000224418 | STK24-AS1 | **1.242017878** | 3.9017238 | 6.15868968 | *4.614653525* |
| ENSG00000165730 | STOX1 | **1.045827048** | 4.489343675 | 6.213947469 | *4.825090867* |
| ENSG00000173320 | STOX2 | **1.088922684** | 5.361333534 | 10.82152787 | *6.174050598* |
| ENSG00000135604 | STX11 | **-3.915687347** | 8.810577956 | 3.414301627 | *1.314669008* |
| ENSG00000111450 | STX2 | **-1.292035958** | 56.62705357 | 19.21176224 | *53.17597186* |
| ENSG00000168952 | STXBP6 | **1.64982045** | 0.773214618 | 1.75510295 | *1.083471874* |
| ENSG00000137573 | SULF1 | **1.47664841** | 5.284533689 | 16.97643575 | *6.678905815* |
| ENSG00000196562 | SULF2 | **4.075756132** | 1.180733618 | 14.0424006 | *0.431784668* |
| ENSG00000130540 | SULT4A1 | **-1.062055327** | 16.80609421 | 10.24199759 | *16.89199588* |
| ENSG00000143502 | SUSD4 | **-1.318542516** | 15.19284479 | 10.48649828 | *14.9161377* |
| ENSG00000165124 | SVEP1 | **3.267355912** | 0.010208859 | 2.655430621 | *0.019152564* |
| ENSG00000147642 | SYBU | **1.167241902** | 0.124454392 | 2.146588039 | *0.136624756* |
| ENSG00000183379 | SYNDIG1L | **2.829642902** | 11.10835791 | 50.89516829 | *16.40341885* |
| ENSG00000176438 | SYNE3 | **-2.202422402** | 6.896479291 | 1.069132159 | *4.561519698* |
| ENSG00000127561 | SYNGR3 | **-1.681822923** | 11.02792853 | 29.39112056 | *9.214876528* |
| ENSG00000171992 | SYNPO | **1.032521199** | 1.595025676 | 2.588822573 | *3.075978889* |
| ENSG00000172403 | SYNPO2 | **1.609600978** | 0.632440017 | 2.207558782 | *0.375535179* |
| ENSG00000143028 | SYPL2 | **3.061791698** | 1.896249586 | 10.96813211 | *2.870574822* |
| ENSG00000067715 | SYT1 | **1.922912332** | 2.190654995 | 6.012697913 | *2.307802278* |
| ENSG00000143858 | SYT2 | **-1.910115536** | 74.91399552 | 18.60344539 | *60.35453406* |
| ENSG00000137501 | SYTL2 | **1.762122798** | 0.205783249 | 3.247488586 | *0.187401642* |
| ENSG00000164674 | SYTL3 | **1.06670066** | 0.736780173 | 7.910515703 | *0.959877807* |
| ENSG00000147526 | TACC1 | **-1.354194359** | 10.33707363 | 2.897745356 | *9.886691817* |
| ENSG00000197780 | TAF13 | **-1.551270948** | 143.7504145 | 32.34725048 | *124.6461131* |
| ENSG00000187325 | TAF9B | **-1.140975942** | 81.8934053 | 25.357413 | *71.75030173* |
| ENSG00000219438 | TAFA5 | **-1.275348964** | 42.61620533 | 28.260007 | *49.64355475* |
| ENSG00000127364 | TAS2R4 | **-1.238397141** | 1.568439371 | 13.13943364 | *0.875087776* |
| ENSG00000184058 | TBX1 | **2.397029335** | 0.045976628 | 5.878975772 | *0.030736513* |
| ENSG00000267280 | TBX2-AS1 | **1.525482178** | 12.53394447 | 24.16332478 | *18.82600259* |
| ENSG00000159450 | TCHH | **1.836769735** | 0.278355989 | 0.699794267 | *0.343002443* |
| ENSG00000176907 | TCIM | **5.682410165** | 0 | 12.24525256 | *0.239489128* |
| ENSG00000203690 | TCP10 | **4.128063137** | 0 | 0.370282028 | *0* |
| ENSG00000166984 | TCP10L2 | **3.527495641** | 0.020747526 | 0.597117255 | *0.027458561* |
| ENSG00000173809 | TDRD12 | **-2.08470261** | 0.829534575 | 0.15826887 | *1.298789647* |
| ENSG00000162782 | TDRD5 | **-1.021938302** | 2.183055033 | 0.795787023 | *2.899610611* |
| ENSG00000007866 | TEAD3 | **1.168857575** | 10.25000839 | 14.82770902 | *12.3139578* |
| ENSG00000163060 | TEKT4 | **1.478246125** | 0.657332985 | 1.564673669 | *0.92553007* |
| ENSG00000145934 | TENM2 | **1.753208693** | 7.389757905 | 24.50657243 | *12.49654593* |

| | | | | | |
|---|---|---|---|---|---|
| ENSG00000158246 | TENT5B | **-1.547993614** | 51.59171447 | 11.69800684 | *62.44291872* |
| ENSG00000226174 | TEX22 | **1.214916122** | 1.935287876 | 11.05695022 | *2.284661188* |
| ENSG00000163424 | TEX55 | **-1.1615663** | 1.534032388 | 6.817838657 | *1.325782367* |
| ENSG00000008196 | TFAP2B | **2.933933362** | 0.584527375 | 4.352647822 | *0.847988384* |
| ENSG00000008197 | TFAP2D | **3.607626831** | 0.057312145 | 0.72600436 | *0.171981561* |
| ENSG00000115112 | TFCP2L1 | **1.203484194** | 0.548415112 | 2.246982916 | *0.985727744* |
| ENSG00000003436 | TFPI | **2.951700188** | 0.383784843 | 2.090174785 | *0.425672446* |
| ENSG00000105825 | TFPI2 | **3.487518809** | 1.073511955 | 8.211070863 | *2.245459214* |
| ENSG00000072274 | TFRC | **-1.328487716** | 64.91731961 | 18.03853663 | *55.24781275* |
| ENSG00000163235 | TGFA | **-2.263496567** | 1.444216832 | 4.316118375 | *0.667830034* |
| ENSG00000105329 | TGFB1 | **4.074018413** | 1.241710776 | 21.65123774 | *1.191387055* |
| ENSG00000140682 | TGFB1I1 | **1.312332081** | 0.283672208 | 0.623355212 | *0.326728135* |
| ENSG00000092969 | TGFB2 | **3.257026007** | 1.216593019 | 15.00912988 | *1.038049* |
| ENSG00000232480 | TGFB2-AS1 | **2.38859641** | 0.899925482 | 3.230114561 | *0.428833302* |
| ENSG00000163513 | TGFBR2 | **3.477776394** | 1.658203567 | 15.77372118 | *3.640315441* |
| ENSG00000260001 | TGFBR3L | **2.316158542** | 0.136377192 | 2.02309419 | *0.355102877* |
| ENSG00000092295 | TGM1 | **-1.294406254** | 0.826782492 | 6.591430828 | *0.888634287* |
| ENSG00000180176 | TH | **-1.47636277** | 0.868669559 | 0.379133016 | *1.830088333* |
| ENSG00000174796 | THAP6 | **-1.14941262** | 23.65232434 | 7.018585697 | *19.99712193* |
| ENSG00000249693 | THEGL | **1.402557861** | 0.56960016 | 0.978980168 | *1.269508419* |
| ENSG00000226856 | THORLNC | **4.60713159** | 0 | 3.993624963 | *0.064044703* |
| ENSG00000136114 | THSD1 | **1.251603022** | 2.101737929 | 3.905773283 | *2.43243713* |
| ENSG00000187720 | THSD4 | **2.789196896** | 0.113202428 | 0.633024595 | *0.150419146* |
| ENSG00000005108 | THSD7A | **3.143300488** | 0.158429356 | 2.548885711 | *0.197169465* |
| ENSG00000140534 | TICRR | **1.175469823** | 1.309258764 | 2.166046423 | *2.840691499* |
| ENSG00000035862 | TIMP2 | **1.711703493** | 0.64521639 | 1.828409316 | *0.712037961* |
| ENSG00000100234 | TIMP3 | **-1.681129756** | 135.2219343 | 28.46659147 | *136.3783216* |
| ENSG00000137251 | TINAG | **2.476238429** | 0.141377009 | 1.19139155 | *0.142311382* |
| ENSG00000174125 | TLR1 | **3.562376159** | 0.006950174 | 14.29144477 | *0.067954717* |
| ENSG00000187554 | TLR5 | **-1.787420034** | 4.905631214 | 1.24531755 | *3.544742372* |
| ENSG00000168955 | TM4SF20 | **-4.802768416** | 0.297176875 | 0.021386052 | *0.232290888* |
| ENSG00000101337 | TM9SF4 | **-1.315163172** | 166.4265215 | 43.43614131 | *156.0470707* |
| ENSG00000284730 | TMDD1 | **1.083508623** | 7.014760119 | 9.652138591 | *7.004095361* |
| ENSG00000144339 | TMEFF2 | **2.718152175** | 0.388026376 | 24.64559764 | *0.787382095* |
| ENSG00000106460 | TMEM106B | **-1.411570083** | 21.92642618 | 10.15678353 | *18.64386966* |
| ENSG00000232258 | TMEM114 | **3.116614415** | 0.122467186 | 1.613072492 | *0.300175723* |
| ENSG00000139173 | TMEM117 | **2.497299962** | 0.285609189 | 4.053878057 | *0.324530955* |
| ENSG00000151952 | TMEM132D | **-1.446397942** | 16.50412369 | 4.384778111 | *10.25822475* |
| ENSG00000146859 | TMEM140 | **2.402883151** | 1.029988349 | 3.973945144 | *0.696346759* |
| ENSG00000249242 | TMEM150C | **-1.05621696** | 9.899349284 | 5.208637503 | *11.03898138* |
| ENSG00000249992 | TMEM158 | **1.504047313** | 18.48049201 | 35.7457426 | *28.34691063* |
| ENSG00000152128 | TMEM163 | **1.448324218** | 0.388429599 | 2.138724061 | *0.633568875* |
| ENSG00000127419 | TMEM175 | **-1.19604215** | 11.40547006 | 21.30350623 | *10.96169726* |
| ENSG00000206432 | TMEM200C | **2.198160534** | 3.59127422 | 10.804129 | *3.368556027* |
| ENSG00000125355 | TMEM255A | **2.190211853** | 0.10826765 | 1.891348944 | *0.235199919* |
| ENSG00000273238 | TMEM271 | **5.481373609** | 0 | 6.214354319 | *0* |
| ENSG00000171227 | TMEM37 | **-1.207419947** | 425.2785593 | 117.8305072 | *384.540644* |

| | | | | | |
|---|---|---|---|---|---|
| ENSG00000151715 | TMEM45B | **2.092346275** | 1.235401258 | 3.918151692 | *1.764755977* |
| ENSG00000175147 | TMEM51-AS1 | **2.037717571** | 0.139837031 | 64.59727463 | *0.187605495* |
| ENSG00000006042 | TMEM98 | **1.385132429** | 16.01469402 | 29.19122097 | *15.57369059* |
| ENSG00000137747 | TMPRSS13 | **-2.876820419** | 0.26398943 | 0.233787707 | *0.140292463* |
| ENSG00000158164 | TMSB15A | **1.03008765** | 68.43761231 | 105.5349404 | *157.4038099* |
| ENSG00000154620 | TMSB4Y | **2.399782214** | 0.524224364 | 1.934331108 | *1.259381036* |
| ENSG00000133687 | TMTC1 | **1.487235866** | 19.44227365 | 81.60333901 | *20.21576463* |
| ENSG00000139921 | TMX1 | **-1.016514889** | 148.3783998 | 48.26242544 | *148.9413332* |
| ENSG00000183578 | TNFAIP8L3 | **1.43287071** | 0.718153889 | 19.57370766 | *0.340565179* |
| ENSG00000120889 | TNFRSF10B | **3.275910415** | 0.133930142 | 25.92783771 | *0.096607477* |
| ENSG00000164761 | TNFRSF11B | **3.961613919** | 0.187921034 | 2.573313523 | *0.108326038* |
| ENSG00000146072 | TNFRSF21 | **2.067380164** | 13.97955079 | 37.97099357 | *6.428825378* |
| ENSG00000145901 | TNIP1 | **-1.694120088** | 20.27635236 | 5.168841825 | *16.94024981* |
| ENSG00000149115 | TNKS1BP1 | **1.189421912** | 10.62653031 | 35.84724265 | *11.88756948* |
| ENSG00000120332 | TNN | **3.421619897** | 0.079753192 | 2.584520791 | *0.171165171* |
| ENSG00000111077 | TNS2 | **1.411125523** | 0.456606618 | 9.003615954 | *0.518111809* |
| ENSG00000173726 | TOMM20 | **-1.229370083** | 142.8290489 | 40.39115755 | *129.3167841* |
| ENSG00000143337 | TOR1AIP1 | **-1.493083177** | 31.15709371 | 7.531074251 | *27.70117607* |
| ENSG00000175274 | TP53I11 | **1.165174834** | 18.61531487 | 47.11778632 | *18.43414665* |
| ENSG00000124251 | TP53TG5 | **3.919541414** | 0.018276772 | 4.041703199 | *0.109719671* |
| ENSG00000078900 | TP73 | **1.715409887** | 0.023841598 | 14.2147287 | *0.127384448* |
| ENSG00000146242 | TPBG | **1.273825954** | 4.758484517 | 7.339061283 | *6.133627228* |
| ENSG00000111907 | TPD52L1 | **1.256375042** | 0.926627813 | 1.425701602 | *1.115009849* |
| ENSG00000139287 | TPH2 | **3.098927961** | 0.25588111 | 5.492713297 | *0.485928409* |
| ENSG00000230359 | TPI1P2 | **-1.116417922** | 11.3232012 | 4.355400712 | *8.675229032* |
| ENSG00000159713 | TPPP3 | **-2.15528933** | 9.100100878 | 2.108580692 | *8.836466087* |
| ENSG00000269113 | TRABD2B | **4.559166345** | 0.008589551 | 1.727502042 | *0.015409975* |
| ENSG00000277734 | TRAC | **4.100357946** | 0 | 1.116709217 | *0.117497765* |
| ENSG00000082512 | TRAF5 | **1.088004847** | 0.329024497 | 0.535908186 | *0.455030666* |
| ENSG00000168016 | TRANK1 | **2.273442108** | 0.165030037 | 0.700048007 | *0.221000357* |
| ENSG00000231165 | TRBV26OR9-2 | **1.142202453** | 8.540978004 | 11.34226532 | *7.211736599* |
| ENSG00000072657 | TRHDE | **2.771685446** | 0.031899 | 0.404962111 | *0.020229413* |
| ENSG00000236333 | TRHDE-AS1 | **1.992467022** | 0.040151263 | 7.536989837 | *0.060683247* |
| ENSG00000173334 | TRIB1 | **1.703486879** | 13.04204862 | 27.12655429 | *12.93312031* |
| ENSG00000255690 | TRIL | **2.715379439** | 0.056745693 | 0.299387419 | *0.099994853* |
| ENSG00000137699 | TRIM29 | **-1.246587687** | 1.433821785 | 15.2557328 | *1.747356981* |
| ENSG00000132256 | TRIM5 | **4.100321974** | 0 | 0.156755329 | *0* |
| ENSG00000166436 | TRIM66 | **-1.000742293** | 1.977433571 | 0.91638949 | *2.01714288* |
| ENSG00000146054 | TRIM7 | **1.088903011** | 1.088783441 | 1.614985232 | *1.167135308* |
| ENSG00000206557 | TRIM71 | **5.960686554** | 0 | 0.432739534 | *0.006602062* |
| ENSG00000100505 | TRIM9 | **1.378162045** | 1.914831331 | 3.945423622 | *2.240116139* |
| ENSG00000121486 | TRMT1L | **-1.025717688** | 80.45534389 | 25.87350087 | *72.05160754* |
| ENSG00000250305 | TRMT9B | **1.113265336** | 0.676763749 | 2.597602929 | *0.854432974* |
| ENSG00000133107 | TRPC4 | **1.040242078** | 3.923647312 | 18.58814279 | *4.258294921* |
| ENSG00000137672 | TRPC6 | **3.205173781** | 0.01028864 | 0.559316021 | *0.026250372* |
| ENSG00000119121 | TRPM6 | **2.334227001** | 0.05622581 | 2.799807403 | *0.104795849* |
| ENSG00000157514 | TSC22D3 | **-1.947692602** | 40.84335807 | 6.830063348 | *32.20742274* |

| | | | | | |
|---|---|---|---|---|---|
| ENSG00000121297 | TSHZ3 | **1.009308236** | 4.678790249 | 6.235367544 | *3.437424078* |
| ENSG00000182704 | TSKU | **1.183576501** | 6.079618619 | 12.48656874 | *7.957029686* |
| ENSG00000117472 | TSPAN1 | **2.078201318** | 1.821319751 | 8.096650556 | *2.54373421* |
| ENSG00000110900 | TSPAN11 | **3.392209564** | 0.015397173 | 4.913418396 | *0.039107365* |
| ENSG00000157570 | TSPAN18 | **3.540365516** | 0.067167501 | 3.114625104 | *0.117346793* |
| ENSG00000175894 | TSPEAR | **-1.46921582** | 9.435775567 | 2.305098902 | *7.936811609* |
| ENSG00000118271 | TTR | **-1.346419662** | 11711.30033 | 3004.935657 | *11804.68237* |
| ENSG00000234711 | TUBB8P11 | **1.617444573** | 1.097474558 | 3.490986997 | *0.635161414* |
| ENSG00000261812 | TUBB8P7 | **1.799439344** | 0.308656025 | 1542.99328 | *0.450154573* |
| ENSG00000104723 | TUSC3 | **-1.052257053** | 140.5553781 | 44.98712308 | *136.8500459* |
| ENSG00000171928 | TVP23B | **-1.118607381** | 25.4147395 | 7.958071068 | *20.16906659* |
| ENSG00000265972 | TXNIP | **1.043795423** | 28.961275 | 62.34857081 | *28.11539137* |
| ENSG00000077721 | UBE2A | **-1.568043033** | 43.57881469 | 13.8751085 | *42.36563757* |
| ENSG00000175063 | UBE2C | **1.07786297** | 17.44426341 | 44.06178763 | *38.46801166* |
| ENSG00000177889 | UBE2N | **-1.153676733** | 67.08653655 | 24.52414349 | *70.38545775* |
| ENSG00000140367 | UBE2Q2 | **-1.186807489** | 37.16453965 | 22.61097486 | *37.86453771* |
| ENSG00000164332 | UBLCP1 | **-1.11474603** | 42.77744293 | 23.40478057 | *41.26354791* |
| ENSG00000280213 | UCKL1-AS1 | **1.359720707** | 0.475928298 | 6.081908913 | *0.539704931* |
| ENSG00000233797 | UFL1-AS1 | **2.076568664** | 1.394528864 | 10.07795661 | *1.353721285* |
| ENSG00000171234 | UGT2B7 | **2.298113533** | 4.064102672 | 13.65164355 | *3.744022972* |
| ENSG00000145626 | UGT3A1 | **1.728192919** | 10.32524997 | 24.16208082 | *14.37637298* |
| ENSG00000168671 | UGT3A2 | **1.635571851** | 0.352629358 | 7.217779635 | *0.968528093* |
| ENSG00000111981 | ULBP1 | **2.424839426** | 4.031261263 | 25.46140154 | *2.937741963* |
| ENSG00000219545 | UMAD1 | **-1.128992297** | 4.754428306 | 1.749437834 | *4.857422782* |
| ENSG00000130477 | UNC13A | **-1.486355873** | 62.73356882 | 21.9896934 | *60.55836416* |
| ENSG00000182168 | UNC5C | **1.764099845** | 0.366030878 | 1.529578518 | *0.481291652* |
| ENSG00000156687 | UNC5D | **-2.138648418** | 12.99510371 | 9.234653424 | *10.72564452* |
| ENSG00000112494 | UNC93A | **2.701742634** | 0.056628861 | 0.669674649 | *0.099375158* |
| ENSG00000110375 | UPK2 | **2.105823187** | 0.256411674 | 1.563932405 | *0.163128907* |
| ENSG00000007001 | UPP2 | **3.25364361** | 0.020530192 | 0.215926037 | *0.090759232* |
| ENSG00000124422 | USP22 | **-1.138757084** | 194.46482 | 58.23780217 | *178.8966996* |
| ENSG00000162738 | VANGL2 | **2.934155008** | 0.073483501 | 0.511631736 | *0.021542445* |
| ENSG00000038427 | VCAN | **5.62843837** | 0.019154874 | 30.17496003 | *0.021980936* |
| ENSG00000128564 | VGF | **2.560402062** | 49.33079503 | 190.5526155 | *56.04748802* |
| ENSG00000026025 | VIM | **5.215552317** | 0.069407789 | 1.932263734 | *0.164345836* |
| ENSG00000106018 | VIPR2 | **-1.155599818** | 0.546563858 | 96.96796418 | *1.115681272* |
| ENSG00000176834 | VSIG10 | **1.347552935** | 4.376500179 | 8.144260544 | *5.310366396* |
| ENSG00000187135 | VSTM2B | **1.706967913** | 1.47965292 | 3.256731957 | *2.389586178* |
| ENSG00000132821 | VSTM2L | **1.742717094** | 2.775554271 | 9.883053566 | *3.671248414* |
| ENSG00000109072 | VTN | **-1.179749414** | 18.48988129 | 6.595248439 | *25.05265724* |
| ENSG00000237499 | WAKMAR2 | **1.14984825** | 0.391278463 | 3.674367121 | *0.528268416* |
| ENSG00000071127 | WDR1 | **-1.002471383** | 66.07322901 | 24.53029145 | *62.19294796* |
| ENSG00000227165 | WDR11-AS1 | **1.261445445** | 0.22215289 | 0.640457905 | *0.171529062* |
| ENSG00000174776 | WDR49 | **4.147438917** | 0.005701382 | 11.14736586 | *0.019870074* |
| ENSG00000168634 | WFDC13 | **-3.179955122** | 1.076613432 | 0.265353035 | *0.480496749* |
| ENSG00000115935 | WIPF1 | **1.703330912** | 2.427091873 | 5.10956029 | *3.01808888* |
| ENSG00000070540 | WIPI1 | **1.392805163** | 16.69839023 | 28.6190812 | *15.15430148* |

| | | | | | |
|---|---|---|---|---|---|
| ENSG00000075290 | WNT8B | **1.936096237** | 0.296135161 | 3.413335522 | *0.515685342* |
| ENSG00000158955 | WNT9B | **1.854026081** | 0.492812311 | 15.89880142 | *0.709383453* |
| ENSG00000075035 | WSCD2 | **-1.162661023** | 2.857817276 | 1.256981524 | *3.354684669* |
| ENSG00000103489 | XYLT1 | **1.03902153** | 7.481217684 | 10.90957877 | *10.67893007* |
| ENSG00000182362 | YBEY | **-1.001808864** | 11.68201867 | 4.218133524 | *10.37074306* |
| ENSG00000181704 | YIPF6 | **-1.31529321** | 60.56475977 | 20.94268344 | *52.59380955* |
| ENSG00000164924 | YWHAZ | **-1.238640082** | 96.2975292 | 28.55111651 | *81.90410836* |
| ENSG00000274775 | Z82195.2 | **1.477927742** | 0.130278831 | 8.184895966 | *0.171845976* |
| ENSG00000261659 | Z92544.1 | **1.264127195** | 9.246698028 | 28.26004617 | *10.50105051* |
| ENSG00000279255 | Z97653.2 | **2.313596517** | 7.403071411 | 23.7575582 | *10.21486945* |
| ENSG00000109906 | ZBTB16 | **2.369372951** | 0.280874973 | 8.196975009 | *0.612102829* |
| ENSG00000184828 | ZBTB7C | **-2.515109221** | 1.416595342 | 13.00748071 | *1.5431191* |
| ENSG00000165424 | ZCCHC24 | **2.187979556** | 0.942927502 | 3.33415332 | *2.510290276* |
| ENSG00000104219 | ZDHHC2 | **-1.603897021** | 78.04912101 | 16.4401186 | *66.15376822* |
| ENSG00000177108 | ZDHHC22 | **3.370668924** | 0.63829781 | 5.706250671 | *0.751989694* |
| ENSG00000185650 | ZFP36L1 | **2.773125781** | 0.285756641 | 10.15971585 | *0.399935428* |
| ENSG00000236869 | ZKSCAN7-AS1 | **1.476153589** | 2.43084258 | 6.008918803 | *2.81449105* |
| ENSG00000165061 | ZMAT4 | **2.98155963** | 0.118891054 | 1.389439809 | *0.251590806* |
| ENSG00000015171 | ZMYND11 | **-1.235296881** | 25.76812845 | 10.03642196 | *23.23037778* |
| ENSG00000152926 | ZNF117 | **2.149305649** | 0.946056243 | 3.127719241 | *0.663051461* |
| ENSG00000179909 | ZNF154 | **2.34158862** | 0.041054214 | 3.855831136 | *0.078552135* |
| ENSG00000264278 | ZNF236-DT | **1.201019978** | 5.257956509 | 8.854214858 | *7.910835877* |
| ENSG00000063587 | ZNF275 | **1.048679121** | 4.08529926 | 5.799985522 | *4.153701446* |
| ENSG00000213801 | ZNF321P | **1.420169585** | 0.542519734 | 5.393498445 | *0.817043598* |
| ENSG00000170954 | ZNF415 | **1.110549233** | 0.332673028 | 3.04577397 | *0.360633822* |
| ENSG00000102935 | ZNF423 | **1.524614982** | 0.82773129 | 2.024785376 | *0.772035963* |
| ENSG00000198342 | ZNF442 | **1.025563392** | 0.714011577 | 1.160642076 | *0.830549831* |
| ENSG00000256229 | ZNF486 | **3.814627217** | 0.014381061 | 0.883132996 | *0.012900081* |
| ENSG00000237149 | ZNF503-AS2 | **-1.071547563** | 15.2756024 | 5.462184185 | *14.01465485* |
| ENSG00000269834 | ZNF528-AS1 | **1.062649987** | 1.799632757 | 2.486181155 | *1.739820755* |
| ENSG00000240225 | ZNF542P | **1.973330991** | 0.293124796 | 3.004318572 | *0.340098615* |
| ENSG00000171970 | ZNF57 | **2.872916106** | 0.050029256 | 1.511234668 | *0.109459092* |
| ENSG00000188171 | ZNF626 | **1.488550033** | 0.191150657 | 0.739566732 | *0.187617441* |
| ENSG00000197497 | ZNF665 | **1.834657746** | 0.113874923 | 0.395333876 | *0.141077779* |
| ENSG00000196172 | ZNF681 | **1.087270405** | 3.25516294 | 4.806925577 | *2.903804078* |
| ENSG00000198429 | ZNF69 | **1.835316719** | 1.622905081 | 3.952821569 | *1.645244876* |
| ENSG00000160352 | ZNF714 | **1.044728053** | 4.204878329 | 7.831508889 | *4.372576853* |
| ENSG00000213967 | ZNF726 | **1.817616841** | 0.468872492 | 3.04887063 | *1.016080479* |
| ENSG00000214189 | ZNF788P | **3.016481482** | 0.052533216 | 3.170474078 | *0.098747742* |
| ENSG00000170396 | ZNF804A | **1.840445564** | 1.41351509 | 3.812921772 | *1.286187176* |
| ENSG00000223547 | ZNF844 | **1.781172667** | 2.09508774 | 4.748295913 | *2.532512037* |
| ENSG00000105750 | ZNF85 | **2.753457756** | 0.035512557 | 1.804510802 | *0.075441317* |
| ENSG00000198155 | ZNF876P | **1.557275058** | 0.38606076 | 3.3477092 | *0.491363742* |
| ENSG00000257446 | ZNF878 | **2.046345511** | 0.364678085 | 0.939542245 | *0.588763734* |
| ENSG00000213793 | ZNF888 | **2.07603148** | 0.850082529 | 2.655810191 | *1.415695559* |
| ENSG00000184635 | ZNF93 | **1.004902441** | 5.159564577 | 7.163764065 | *6.393739058* |
| ENSG00000260233 | ZNRD2-AS1 | **1.06823553** | 10.66978979 | 16.30615525 | *12.03644011* |

| | | | | | |
|---|---|---|---|---|---|
| ENSG00000130182 | ZSCAN10 | **4.407664921** | 0 | 3.712125157 | *0.012427033* |

**Supplemental Table 3**

| **Gene ID** | **Gene name** | **Log2(fold-change)** | **Average TPM in Sham** | **Average TPM in PLIN2OE** | ***Average TPM in PLIN2KD*** |
|---|---|---|---|---|---|
| ENSG00000281530 | AC004461.3 | **-1.357250162** | 4.23999504 | 1.655809406 | *1.593035118* |
| ENSG00000250770 | AC005865.1 | **1.98889788** | 0.220241934 | 0.87077642 | *0.405218114* |
| ENSG00000267727 | AC008738.5 | **1.36475845** | 3.454929899 | 8.864199873 | *2.637621568* |
| ENSG00000280239 | AC011498.7 | **1.295919257** | 1.709380227 | 4.191496536 | *3.467004901* |
| ENSG00000230732 | AC016949.1 | **-1.35185068** | 2.50438709 | 0.977143717 | *2.372850091* |
| ENSG00000284946 | AC068831.8 | **1.306897573** | 0.303233524 | 0.747271136 | *0.443686245* |
| ENSG00000224593 | AC092427.1 | **-2.266554108** | 4.364996913 | 0.914876362 | *1.772556222* |
| ENSG00000241462 | AC100832.1 | **-1.47244482** | 9.142600484 | 3.313371595 | *8.721904477* |
| ENSG00000251023 | AC114980.1 | **-1.22074321** | 1.377582724 | 0.594173594 | *1.169332104* |
| ENSG00000284727 | AC116562.3 | **1.398917443** | 1.49801364 | 3.908182405 | *3.378291936* |
| ENSG00000261758 | AC117382.2 | **-1.159190926** | 1.447977185 | 0.647396259 | *2.044651562* |
| ENSG00000279283 | AC131009.4 | **-1.379835242** | 4.41766334 | 1.712103689 | *2.768106538* |
| ENSG00000278928 | AC136621.1 | **1.786083673** | 1.011745784 | 3.522230912 | *2.268771951* |
| ENSG00000154734 | ADAMTS1 | **1.321927149** | 0.187484702 | 0.465900344 | *5.472211502* |
| ENSG00000173175 | ADCY5 | **1.329029505** | 0.123341184 | 0.312316344 | *0.151610635* |
| ENSG00000121753 | ADGRB2 | **1.126263944** | 0.141678306 | 0.311224196 | *0.398552109* |
| ENSG00000270083 | AL021878.2 | **-1.370081018** | 13.99923947 | 5.407565154 | *13.08453483* |
| ENSG00000275620 | AL121827.2 | **-1.612635764** | 0.430816944 | 0.140956777 | *1.738362763* |
| ENSG00000227678 | AL355581.1 | **1.154844384** | 0.720988146 | 1.60234935 | *0.563589754* |
| ENSG00000167612 | ANKRD33 | **1.057131119** | 0.557011015 | 1.160055927 | *1.533600541* |
| ENSG00000189127 | ANKRD34B | **1.517716703** | 0.403585998 | 1.159367333 | *0.42268324* |
| ENSG00000169604 | ANTXR1 | **2.400602755** | 0.168820402 | 0.892249262 | *2.146278598* |
| ENSG00000266767 | AP000897.2 | **1.181489989** | 1.165013242 | 2.663410023 | *1.636162352* |
| ENSG00000130203 | APOE | **1.173559508** | 1.957487283 | 4.424768129 | *1.673799935* |
| ENSG00000105011 | ASF1B | **1.23281404** | 12.74164254 | 29.97103719 | *21.79183816* |
| ENSG00000066279 | ASPM | **1.103150313** | 5.172207327 | 11.11603092 | *8.41461578* |
| ENSG00000124788 | ATXN1 | **-1.378440304** | 0.568125769 | 0.220102097 | *1.043023175* |
| ENSG00000178999 | AURKB | **1.412844187** | 4.177215703 | 11.12521578 | *7.141131162* |
| ENSG00000105327 | BBC3 | **-1.205826113** | 5.462620607 | 2.377127986 | *7.535782271* |
| ENSG00000023445 | BIRC3 | **-6.199403822** | 10.42018447 | 0.144706032 | *0.14559059* |
| ENSG00000089685 | BIRC5 | **1.245241233** | 6.212223004 | 14.73117079 | *11.74608317* |
| ENSG00000136492 | BRIP1 | **1.071314812** | 1.886978522 | 3.974506191 | *3.107412986* |
| ENSG00000174808 | BTC | **1.396912076** | 0.604557413 | 1.58075937 | *0.853466254* |
| ENSG00000165810 | BTNL9 | **1.298039374** | 1.270501933 | 3.134256378 | *1.500570433* |
| ENSG00000169679 | BUB1 | **1.106482061** | 3.864257679 | 8.322845532 | *6.869850531* |
| ENSG00000156970 | BUB1B | **1.091166666** | 8.988577225 | 19.15847854 | *11.98945791* |
| ENSG00000154102 | C16orf74 | **1.137807899** | 0.457072784 | 0.999762628 | *0.335358703* |
| ENSG00000162706 | CADM3 | **1.475660331** | 0.507987817 | 1.404142173 | *3.87155361* |
| ENSG00000081803 | CADPS2 | **1.235950354** | 0.276216505 | 0.652921495 | *0.493442522* |
| ENSG00000183049 | CAMK1D | **1.362395995** | 0.299564879 | 0.774141233 | *0.792036664* |
| ENSG00000163888 | CAMK2N2 | **1.043383083** | 20.03815467 | 41.48105803 | *50.0612059* |
| ENSG00000183287 | CCBE1 | **2.022502376** | 0.099978073 | 0.406446204 | *3.218687376* |
| ENSG00000144395 | CCDC150 | **1.021820678** | 1.521563588 | 3.092579189 | *7.027418028* |
| ENSG00000115009 | CCL20 | **-22.32200339** | 2.967713822 | 0 | *0* |

| | | | | | |
|---|---|---|---|---|---|
| ENSG00000145386 | CCNA2 | **1.094685981** | 15.39963331 | 32.90524203 | *11.32138988* |
| ENSG00000157456 | CCNB2 | **1.138378573** | 8.390473114 | 18.47036871 | *10.50228489* |
| ENSG00000156535 | CD109 | **1.330028218** | 1.421182092 | 3.595117568 | *1.45082192* |
| ENSG00000026508 | CD44 | **1.2452786** | 0.071676466 | 0.170538492 | *0.038949698* |
| ENSG00000117399 | CDC20 | **1.09774526** | 18.20414817 | 38.98446123 | *28.7211191* |
| ENSG00000158402 | CDC25C | **1.065826122** | 1.797955345 | 3.758263869 | *3.207638932* |
| ENSG00000093009 | CDC45 | **1.274771023** | 1.966306914 | 4.761139114 | *2.321975273* |
| ENSG00000166589 | CDH16 | **1.164441369** | 4.21670042 | 9.484052678 | *1.695437551* |
| ENSG00000170312 | CDK1 | **1.109178235** | 12.35050404 | 26.67047323 | *19.37115405* |
| ENSG00000117266 | CDK18 | **1.428204622** | 0.113583099 | 0.305895595 | *0.179978285* |
| ENSG00000124762 | CDKN1A | **-1.046470119** | 133.6668482 | 64.92046721 | *691.1395526* |
| ENSG00000277449 | CEBPB-AS1 | **-1.380890827** | 1.948632122 | 0.748718543 | *5.566439033* |
| ENSG00000115163 | CENPA | **1.050592267** | 4.838032125 | 10.03240553 | *7.749794589* |
| ENSG00000138778 | CENPE | **1.036257259** | 2.118400186 | 4.357053799 | *2.634029976* |
| ENSG00000102384 | CENPI | **1.186373574** | 2.139869835 | 4.87895831 | *3.817985319* |
| ENSG00000123219 | CENPK | **1.250562727** | 2.595554265 | 6.176874404 | *3.722418982* |
| ENSG00000100162 | CENPM | **1.193385966** | 5.175725437 | 11.85011015 | *8.260961659* |
| ENSG00000128965 | CHAC1 | **-1.072520941** | 37.44371846 | 17.8719427 | *101.8716087* |
| ENSG00000101204 | CHRNA4 | **1.099748447** | 0.099077435 | 0.212861072 | *0.04712137* |
| ENSG00000127586 | CHTF18 | **1.008066277** | 2.362063281 | 4.75972464 | *2.736180378* |
| ENSG00000122966 | CIT | **1.13349948** | 1.496464734 | 3.289851264 | *1.646383753* |
| ENSG00000118432 | CNR1 | **-2.515773655** | 0.183692425 | 0.032773825 | *0.099516324* |
| ENSG00000124749 | COL21A1 | **1.909783451** | 0.085163045 | 0.319802945 | *0.367459006* |
| ENSG00000187498 | COL4A1 | **1.046332722** | 1.552675292 | 3.204093445 | *1.069465529* |
| ENSG00000134871 | COL4A2 | **1.083988535** | 10.61145058 | 22.541473 | *7.976746476* |
| ENSG00000081052 | COL4A4 | **1.302663413** | 0.253959723 | 0.632425352 | *0.193353549* |
| ENSG00000049089 | COL9A2 | **1.177777309** | 0.959107735 | 2.175672698 | *1.087440319* |
| ENSG00000095713 | CRTAC1 | **1.718047501** | 0.399051706 | 1.323316609 | *1.232263846* |
| ENSG00000169862 | CTNND2 | **1.065946034** | 5.931017961 | 12.43618188 | *10.81739551* |
| ENSG00000103811 | CTSH | **1.021410284** | 0.286981784 | 0.581606534 | *0.558178025* |
| ENSG00000163739 | CXCL1 | **-7.342159085** | 232.8164847 | 1.46196609 | *3.019266601* |
| ENSG00000163734 | CXCL3 | **-4.632218082** | 10.32456113 | 0.424059091 | *0.376150603* |
| ENSG00000169429 | CXCL8 | **-21.87106723** | 2.385221565 | 0 | *0.180220548* |
| ENSG00000166394 | CYB5R2 | **1.105418994** | 0.142407605 | 0.305069209 | *0.254107517* |
| ENSG00000186529 | CYP4F3 | **1.890323557** | 0.059648728 | 0.219427634 | *0.236856472* |
| ENSG00000187323 | DCC | **1.344122301** | 1.388463351 | 3.533571063 | *4.079219374* |
| ENSG00000214826 | DDX12P | **1.174446505** | 3.869288742 | 8.743872125 | *3.827194045* |
| ENSG00000035499 | DEPDC1B | **1.100195133** | 4.380893289 | 9.396361571 | *7.371461575* |
| ENSG00000165507 | DEPP1 | **-1.051304609** | 121.0339613 | 58.61968406 | *564.2512541* |
| ENSG00000165023 | DIRAS2 | **-1.232395951** | 3.685715586 | 1.570203703 | *2.067712524* |
| ENSG00000126787 | DLGAP5 | **1.139499472** | 6.529822861 | 14.39467954 | *10.11252904* |
| ENSG00000187957 | DNER | **1.033971336** | 7.073740386 | 14.49130076 | *10.81214874* |
| ENSG00000151640 | DPYSL4 | **1.378393276** | 1.378585614 | 3.597443565 | *5.87823905* |
| ENSG00000245750 | DRAIC | **1.858954345** | 0.084094812 | 0.30435336 | *0.076911499* |
| ENSG00000101412 | E2F1 | **1.103186294** | 145.8455947 | 313.7864139 | *257.7912616* |
| ENSG00000007968 | E2F2 | **1.054893949** | 5.95576412 | 12.38445293 | *5.718146114* |
| ENSG00000129173 | E2F8 | **1.195912146** | 1.28319032 | 2.938721966 | *1.677154218* |

| | | | | | |
|---|---|---|---|---|---|
| ENSG00000233167 | EEF1A1P26 | **-1.652271803** | 7.092721072 | 2.265215295 | *4.80784785* |
| ENSG00000146648 | EGFR | **1.039748364** | 0.390704239 | 0.808641641 | *1.231436087* |
| ENSG00000120738 | EGR1 | **-1.338087071** | 55.67643445 | 22.15764333 | *144.2821892* |
| ENSG00000122877 | EGR2 | **-1.403106987** | 0.418093645 | 0.158572685 | *0.596170172* |
| ENSG00000186998 | EMID1 | **1.271427242** | 4.218315704 | 10.19244872 | *6.542057739* |
| ENSG00000183798 | EMILIN3 | **1.092769557** | 8.113520727 | 17.26693987 | *11.1468712* |
| ENSG00000135638 | EMX1 | **1.373481577** | 1.042835712 | 2.706163201 | *0.85561214* |
| ENSG00000105131 | EPHX3 | **1.453451549** | 0.40088857 | 1.089786175 | *0.717311076* |
| ENSG00000273604 | EPOP | **1.212370125** | 1.010327018 | 2.334804596 | *3.369052374* |
| ENSG00000186871 | ERCC6L | **1.073525606** | 2.321570491 | 4.890664748 | *3.803010304* |
| ENSG00000135476 | ESPL1 | **1.255843516** | 1.37540943 | 3.284313421 | *1.687518263* |
| ENSG00000187017 | ESPN | **1.111450432** | 0.713263588 | 1.543404914 | *0.586915554* |
| ENSG00000115363 | EVA1A | **1.88959972** | 0.130785142 | 0.485694531 | *3.479163978* |
| ENSG00000174371 | EXO1 | **1.106892111** | 1.285210584 | 2.771495204 | *2.261959149* |
| ENSG00000163586 | FABP1 | **1.57657169** | 0.919638243 | 2.745032646 | *0.368929787* |
| ENSG00000189057 | FAM111B | **1.001235861** | 5.130006744 | 10.28862583 | *4.046141286* |
| ENSG00000196550 | FAM72A | **1.025015801** | 1.023994464 | 2.081687384 | *0.94042145* |
| ENSG00000188610 | FAM72B | **1.154397838** | 0.867003658 | 1.933938131 | *0.738457173* |
| ENSG00000263513 | FAM72C | **1.663509898** | 0.367983593 | 1.161712693 | *0.775879253* |
| ENSG00000215784 | FAM72D | **1.305722596** | 1.768900534 | 4.361868329 | *1.99737389* |
| ENSG00000101447 | FAM83D | **1.067116656** | 5.430564727 | 11.38954288 | *9.37627262* |
| ENSG00000170271 | FAXDC2 | **-1.497403076** | 1.405772308 | 0.498663949 | *1.526725643* |
| ENSG00000077942 | FBLN1 | **1.038256495** | 0.955913658 | 1.964926941 | *1.232004278* |
| ENSG00000156509 | FBXO43 | **1.026486144** | 0.796314249 | 1.616942498 | *1.160397604* |
| ENSG00000186188 | FFAR4 | **-1.261674704** | 0.791837867 | 0.331552333 | *0.312983332* |
| ENSG00000111206 | FOXM1 | **1.137649226** | 10.58437114 | 23.32242565 | *14.82475061* |
| ENSG00000147234 | FRMPD3 | **-1.139272073** | 1.071325058 | 0.492192947 | *0.072023014* |
| ENSG00000152254 | G6PC2 | **1.134264334** | 19.49439322 | 42.79910265 | *60.41383229* |
| ENSG00000160219 | GAB3 | **1.047204806** | 0.283924213 | 0.587356742 | *0.326247751* |
| ENSG00000136928 | GABBR2 | **1.609248945** | 0.560053752 | 1.705566603 | *1.361594952* |
| ENSG00000108479 | GALK1 | **1.289699642** | 0.230447423 | 0.564079948 | *3.228644643* |
| ENSG00000172020 | GAP43 | **-1.158393559** | 17.21722837 | 7.7598421 | *16.59945603* |
| ENSG00000130513 | GDF15 | **-1.156283422** | 22.05402392 | 9.935306238 | *49.66504118* |
| ENSG00000181938 | GINS3 | **1.089666447** | 2.645775926 | 5.648666725 | *3.752581178* |
| ENSG00000159248 | GJD2 | **-1.062831757** | 71.5625691 | 34.31542968 | *30.22789832* |
| ENSG00000149328 | GLB1L2 | **1.056118248** | 8.147169123 | 16.96232323 | *5.887545343* |
| ENSG00000170075 | GPR37L1 | **-7.281491269** | 5.421761877 | 0.035308191 | *0.056623019* |
| ENSG00000176153 | GPX2 | **-1.461006583** | 10.99377314 | 4.034104036 | *1.109881047* |
| ENSG00000198785 | GRIN3A | **1.155318517** | 23.19850343 | 51.71113558 | *30.67778668* |
| ENSG00000164082 | GRM2 | **1.194217585** | 0.537650387 | 1.233449899 | *1.343042276* |
| ENSG00000179603 | GRM8 | **1.069760042** | 0.536938514 | 1.130527959 | *1.100818128* |
| ENSG00000075218 | GTSE1 | **1.195298284** | 9.745038575 | 22.36371452 | *14.42941626* |
| ENSG00000148702 | HABP2 | **1.307299129** | 0.39969948 | 0.992276192 | *0.732815265* |
| ENSG00000123485 | HJURP | **1.100000124** | 7.769658868 | 16.66864764 | *12.30012435* |
| ENSG00000186603 | HPDL | **1.410234879** | 1.013547684 | 2.694198599 | *1.740193694* |
| ENSG00000107521 | HPS1 | **1.54670553** | 0.056506899 | 0.164373356 | *0.058315736* |
| ENSG00000101180 | HRH3 | **1.046122645** | 0.835346956 | 1.719693752 | *1.41049907* |

| | | | | | |
|---|---|---|---|---|---|
| ENSG00000115457 | IGFBP2 | **1.367394974** | 2.63850693 | 6.837761518 | *4.24880195* |
| ENSG00000163453 | IGFBP7 | **1.205715704** | 0.758986882 | 1.758286306 | *0.809341711* |
| ENSG00000115590 | IL1R2 | **1.554347464** | 0.154768013 | 0.451509954 | *1.014679439* |
| ENSG00000183856 | IQGAP3 | **1.160243231** | 7.677160639 | 17.17950795 | *7.109155428* |
| ENSG00000096433 | ITPR3 | **1.18809231** | 3.621521981 | 8.279189398 | *3.078090151* |
| ENSG00000171385 | KCND3 | **1.357002691** | 3.514014421 | 9.049667317 | *1.085766576* |
| ENSG00000152049 | KCNE4 | **1.734769674** | 0.24608306 | 0.823300875 | *1.537691542* |
| ENSG00000140015 | KCNH5 | **1.237085845** | 0.097434237 | 0.230844882 | *1.193432988* |
| ENSG00000053918 | KCNQ1 | **1.009443773** | 0.986852055 | 1.993418836 | *2.362764199* |
| ENSG00000135253 | KCP | **1.843003921** | 0.051815264 | 0.186286915 | *0.143470439* |
| ENSG00000153885 | KCTD15 | **1.353782059** | 0.09648956 | 0.248813421 | *1.367311727* |
| ENSG00000138160 | KIF11 | **1.023535849** | 24.10128662 | 49.01962697 | *40.87839108* |
| ENSG00000118193 | KIF14 | **1.197660476** | 4.390719776 | 10.11206336 | *7.017759915* |
| ENSG00000163808 | KIF15 | **1.061391339** | 2.753367822 | 5.754861907 | *4.377099119* |
| ENSG00000121621 | KIF18A | **1.077835255** | 4.521387848 | 9.551011197 | *6.605681556* |
| ENSG00000186185 | KIF18B | **1.337817401** | 6.288653655 | 15.90245522 | *10.17179692* |
| ENSG00000112984 | KIF20A | **1.092263693** | 7.60353196 | 16.21023697 | *10.1862948* |
| ENSG00000137807 | KIF23 | **1.040903846** | 1.963324892 | 4.040197927 | *2.911612016* |
| ENSG00000142945 | KIF2C | **1.395663139** | 4.221433369 | 11.10280042 | *5.26525026* |
| ENSG00000090889 | KIF4A | **1.086801819** | 15.14274517 | 32.19998935 | *22.46663746* |
| ENSG00000237649 | KIFC1 | **1.146557989** | 22.90346287 | 50.79638593 | *40.44758759* |
| ENSG00000157404 | KIT | **1.37959028** | 3.505522685 | 9.140034595 | *13.8607121* |
| ENSG00000136826 | KLF4 | **1.195602602** | 1.880995691 | 4.306159654 | *3.599142449* |
| ENSG00000116678 | LEPR | **1.040530304** | 0.322312761 | 0.663032205 | *2.481576004* |
| ENSG00000254275 | LINC00824 | **1.741079565** | 0.819002672 | 2.744081162 | *2.272411751* |
| ENSG00000281769 | LINC01230 | **1.104987656** | 0.815592365 | 1.760409065 | *1.014830193* |
| ENSG00000231437 | LINC01750 | **1.657092766** | 0.200782116 | 0.637147719 | *0.275037269* |
| ENSG00000235142 | LINC02532 | **1.081001809** | 0.43842759 | 0.924787616 | *0.412819572* |
| ENSG00000254119 | LINC02842 | **1.488535548** | 1.217475583 | 3.435974713 | *1.160826426* |
| ENSG00000166035 | LIPC | **1.852461303** | 2.218312748 | 8.013277069 | *5.43435684* |
| ENSG00000254815 | LMNTD2-AS1 | **1.130732068** | 0.808810533 | 1.787636776 | *1.151316115* |
| ENSG00000138131 | LOXL4 | **1.595545953** | 43.19454411 | 131.1330978 | *23.41185318* |
| ENSG00000185565 | LSAMP | **1.051185333** | 3.274182667 | 6.783650895 | *12.95072195* |
| ENSG00000119681 | LTBP2 | **1.060934302** | 4.153678091 | 8.699646861 | *6.367306061* |
| ENSG00000150551 | LYPD1 | **-1.911351478** | 6.802069653 | 1.829540461 | *0.494710989* |
| ENSG00000182759 | MAFA | **1.51849927** | 3.028655848 | 8.699387193 | *1.813619939* |
| ENSG00000234456 | MAGI2-AS3 | **1.614270355** | 0.070616793 | 0.218412592 | *0.19276967* |
| ENSG00000091436 | MAP3K20 | **1.154823697** | 0.865267145 | 1.931044132 | *1.617043732* |
| ENSG00000065328 | MCM10 | **1.022308189** | 2.670193691 | 5.432615326 | *4.470231415* |
| ENSG00000110492 | MDK | **1.276075743** | 2.778864962 | 6.745136775 | *5.764231025* |
| ENSG00000065833 | ME1 | **-1.050752918** | 7.125185135 | 3.455858538 | *9.577815626* |
| ENSG00000164877 | MICALL2 | **1.825904801** | 0.045594974 | 0.161674105 | *0.081728296* |
| ENSG00000148773 | MKI67 | **1.059774829** | 16.46809276 | 34.37884802 | *23.79872925* |
| ENSG00000087245 | MMP2 | **1.323280592** | 1.132056781 | 2.832074079 | *9.962732913* |
| ENSG00000121211 | MND1 | **1.258304952** | 0.521262959 | 1.249677909 | *0.725524432* |
| ENSG00000138823 | MTTP | **-1.179885889** | 0.590014805 | 0.260358119 | *0.384450496* |
| ENSG00000213347 | MXD3 | **1.230878369** | 3.011392913 | 7.088317085 | *3.490503512* |

| | | | | | |
|---|---|---|---|---|---|
| ENSG00000101057 | MYBL2 | **1.32933743** | 10.32494302 | 25.99007362 | *20.55470376* |
| ENSG00000136997 | MYC | **1.197489891** | 0.224721484 | 0.520947443 | *1.660163909* |
| ENSG00000166866 | MYO1A | **1.082854726** | 2.390492876 | 5.087539968 | *2.336650401* |
| ENSG00000168060 | NAALADL1 | **2.175714298** | 0.199933991 | 0.901736709 | *0.112250659* |
| ENSG00000080986 | NDC80 | **1.107800031** | 9.580690717 | 20.62177257 | *14.99574918* |
| ENSG00000109674 | NEIL3 | **1.028391218** | 2.831173005 | 5.778032257 | *3.425649252* |
| ENSG00000117650 | NEK2 | **1.138512028** | 8.502456265 | 18.75026547 | *13.70465668* |
| ENSG00000116962 | NID1 | **1.379898796** | 1.630357101 | 4.263490353 | *4.088059097* |
| ENSG00000185942 | NKAIN3 | **1.435712451** | 0.342469809 | 0.935579016 | *1.552121009* |
| ENSG00000109255 | NMU | **1.654289685** | 0.272165648 | 0.856523317 | *1.003372242* |
| ENSG00000185269 | NOTUM | **1.132310709** | 1.200750096 | 2.653628917 | *1.181495084* |
| ENSG00000151322 | NPAS3 | **1.193929341** | 0.143285279 | 0.330173662 | *0.203886918* |
| ENSG00000113389 | NPR3 | **1.480030198** | 0.218318444 | 0.608485625 | *1.602503813* |
| ENSG00000122585 | NPY | **1.194308472** | 2.081520478 | 4.762254486 | *6.636155164* |
| ENSG00000143228 | NUF2 | **1.122685375** | 6.544171202 | 14.26983658 | *11.97958942* |
| ENSG00000137804 | NUSAP1 | **1.034711058** | 12.71167358 | 26.10234932 | *12.11811144* |
| ENSG00000183801 | OLFML1 | **1.202421941** | 0.551250043 | 1.274148996 | *1.609315438* |
| ENSG00000090530 | P3H2 | **-1.094778722** | 4.997572812 | 2.345278585 | *2.336387963* |
| ENSG00000185480 | PARPBP | **1.064767625** | 1.559236363 | 3.258714278 | *2.626104586* |
| ENSG00000168078 | PBK | **1.120420609** | 7.694953756 | 16.70400513 | *13.43075897* |
| ENSG00000242419 | PCDHGC4 | **1.127904526** | 1.666597893 | 3.640166317 | *1.259932488* |
| ENSG00000099139 | PCSK5 | **1.102962972** | 0.435644822 | 0.939953831 | *1.177357042* |
| ENSG00000184588 | PDE4B | **1.325939975** | 0.077720738 | 0.195864877 | *0.223834288* |
| ENSG00000140451 | PIF1 | **2.358032745** | 0.204154668 | 1.05014041 | *0.377622728* |
| ENSG00000129195 | PIMREG | **1.008048599** | 1.230617518 | 2.473987356 | *2.311147437* |
| ENSG00000087842 | PIR | **-1.009698326** | 3.423931406 | 1.692346475 | *2.687735043* |
| ENSG00000127564 | PKMYT1 | **1.025625606** | 1.705149647 | 3.477113363 | *2.686653616* |
| ENSG00000160447 | PKN3 | **1.441112123** | 0.294165808 | 0.793936536 | *1.85329361* |
| ENSG00000166851 | PLK1 | **1.172960503** | 6.815553723 | 15.4007481 | *10.4899421* |
| ENSG00000100341 | PNPLA5 | **1.225354449** | 0.604275414 | 1.414962348 | *0.436395866* |
| ENSG00000051341 | POLQ | **1.022353651** | 2.387035125 | 4.860142267 | *2.657747837* |
| ENSG00000155846 | PPARGC1B | **1.517703421** | 0.097871114 | 0.281996258 | *0.597044857* |
| ENSG00000198901 | PRC1 | **1.116286254** | 0.252967132 | 0.547595626 | *0.404097288* |
| ENSG00000057657 | PRDM1 | **-1.602130517** | 0.567182341 | 0.18851187 | *0.418241191* |
| ENSG00000141391 | PRELID3A | **1.447705902** | 0.227629934 | 0.621003361 | *0.63517655* |
| ENSG00000007062 | PROM1 | **2.338122207** | 0.055694422 | 0.283050453 | *0.351316595* |
| ENSG00000182077 | PTCHD3 | **1.316316228** | 0.237519069 | 0.592400788 | *0.433788857* |
| ENSG00000125384 | PTGER2 | **1.235029625** | 0.799502875 | 1.877750754 | *1.419601369* |
| ENSG00000111247 | RAD51AP1 | **1.016972327** | 2.767977291 | 5.601238845 | *4.405551132* |
| ENSG00000085999 | RAD54L | **1.281959619** | 2.21160979 | 5.385621267 | *3.861909012* |
| ENSG00000204764 | RANBP17 | **1.039318683** | 0.072116621 | 0.14855773 | *0.219450193* |
| ENSG00000111344 | RASAL1 | **1.750602545** | 0.041648822 | 0.143774308 | *0.081515038* |
| ENSG00000147509 | RGS20 | **1.01447113** | 0.448413317 | 0.90470148 | *5.568876204* |
| ENSG00000138835 | RGS3 | **1.058268367** | 0.616603042 | 1.284315636 | *0.464099202* |
| ENSG00000141314 | RHBDL3 | **1.107732957** | 0.53479432 | 1.158714221 | *1.771202468* |
| ENSG00000101098 | RIMS4 | **1.261957897** | 2.470923525 | 5.937902183 | *9.302738182* |
| ENSG00000183145 | RIPPLY3 | **1.520719986** | 8.492781266 | 24.34639437 | *21.74661342* |

| | | | | | |
|---|---|---|---|---|---|
| ENSG00000222898 | RN7SKP97 | **1.125990444** | 6.580068647 | 14.43724789 | *9.764839942* |
| ENSG00000171848 | RRM2 | **1.15928687** | 5.265328232 | 11.76541273 | *9.688418639* |
| ENSG00000115884 | SDC1 | **1.222031841** | 2.4877889 | 5.813467253 | *3.032260296* |
| ENSG00000099937 | SERPIND1 | **1.315310849** | 1.074677381 | 2.681319266 | *2.455159433* |
| ENSG00000138944 | SHISAL1 | **1.146183624** | 0.513096707 | 1.137347939 | *1.801505592* |
| ENSG00000170577 | SIX2 | **1.268815872** | 10.82514284 | 26.01952683 | *40.53720624* |
| ENSG00000132874 | SLC14A2 | **1.239781639** | 0.423682776 | 1.006287499 | *0.628378734* |
| ENSG00000124568 | SLC17A1 | **1.743783308** | 1.184558152 | 3.978574059 | *2.916434853* |
| ENSG00000105281 | SLC1A5 | **1.264711882** | 0.543126442 | 1.301032552 | *1.528262603* |
| ENSG00000170482 | SLC23A1 | **1.231823837** | 1.078549139 | 2.519399633 | *0.701586399* |
| ENSG00000196376 | SLC35F1 | **1.357680184** | 2.04959619 | 5.270607157 | *8.4263338* |
| ENSG00000139540 | SLC39A5 | **1.425909421** | 1.132741696 | 3.053113498 | *2.853724159* |
| ENSG00000134802 | SLC43A3 | **1.140329575** | 1.513951069 | 3.346110883 | *3.030597367* |
| ENSG00000103546 | SLC6A2 | **1.53320571** | 0.887531775 | 2.578413464 | *2.708367174* |
| ENSG00000151012 | SLC7A11 | **-1.051185448** | 3.276649668 | 1.591366075 | *51.24497257* |
| ENSG00000101665 | SMAD7 | **-1.355818814** | 4.026418474 | 1.587400816 | *4.613376296* |
| ENSG00000285441 | SOD2 | **-1.059709499** | 2.210446447 | 1.06379642 | *1.877956694* |
| ENSG00000181449 | SOX2 | **1.232740041** | 0.991170235 | 2.323673728 | *2.375330289* |
| ENSG00000076382 | SPAG5 | **1.099721595** | 7.920611377 | 16.9866191 | *11.74188827* |
| ENSG00000161888 | SPC24 | **1.768888081** | 2.497899758 | 8.493319945 | *6.144192708* |
| ENSG00000066923 | STAG3 | **1.17026422** | 0.250446287 | 0.564208277 | *0.266395858* |
| ENSG00000105889 | STEAP1B | **1.325288554** | 0.319908539 | 0.804225373 | *0.934286114* |
| ENSG00000135604 | STX11 | **-2.768498637** | 8.810577956 | 1.314669008 | *0.581047425* |
| ENSG00000196562 | SULF2 | **-1.450490263** | 1.180733618 | 0.431784668 | *19.66125234* |
| ENSG00000105825 | TFPI2 | **1.056406827** | 1.073511955 | 2.245459214 | *11.90848598* |
| ENSG00000163235 | TGFA | **-1.120849545** | 1.444216832 | 0.667830034 | *0.29917582* |
| ENSG00000163513 | TGFBR2 | **1.134705775** | 1.658203567 | 3.640315441 | *18.20157927* |
| ENSG00000180176 | TH | **1.070388478** | 0.868669559 | 1.830088333 | *0.307924141* |
| ENSG00000140534 | TICRR | **1.116241187** | 1.309258764 | 2.840691499 | *2.914969119* |
| ENSG00000167900 | TK1 | **1.523759782** | 3.600645071 | 10.3603826 | *5.3232948* |
| ENSG00000144339 | TMEFF2 | **1.020165123** | 0.388026376 | 0.787382095 | *2.519960653* |
| ENSG00000148483 | TMEM236 | **1.12843506** | 1.960286143 | 4.282656209 | *2.502366238* |
| ENSG00000158164 | TMSB15A | **1.199472517** | 68.43761231 | 157.4038099 | *137.9866718* |
| ENSG00000232810 | TNF | **-22.14052732** | 5.216506876 | 0 | *0.033140042* |
| ENSG00000146072 | TNFRSF21 | **-1.121464689** | 13.97955079 | 6.428825378 | *57.70629914* |
| ENSG00000160949 | TONSL | **1.100783305** | 5.368149128 | 11.52384777 | *8.479424081* |
| ENSG00000131747 | TOP2A | **1.11274897** | 73.52818305 | 159.2127799 | *53.21342025* |
| ENSG00000078900 | TP73 | **2.402028951** | 0.023841598 | 0.127384448 | *0.07780276* |
| ENSG00000071539 | TRIP13 | **1.028943909** | 8.286996892 | 16.91831219 | *12.43275806* |
| ENSG00000135451 | TROAP | **1.569927921** | 2.79201428 | 8.288625048 | *4.271276082* |
| ENSG00000112742 | TTK | **1.003660956** | 4.596554934 | 9.208176938 | *6.237457231* |
| ENSG00000176014 | TUBB6 | **1.103451116** | 6.345004554 | 13.67294362 | *5.54918573* |
| ENSG00000175063 | UBE2C | **1.1403962** | 17.44426341 | 38.46801166 | *36.3051832* |
| ENSG00000168671 | UGT3A2 | **1.460396665** | 0.352629358 | 0.968528093 | *1.082679916* |
| ENSG00000276043 | UHRF1 | **1.101189811** | 3.416468673 | 7.340149527 | *4.750279551* |
| ENSG00000106018 | VIPR2 | **1.022484342** | 0.546563858 | 1.115681272 | *0.239764641* |
| ENSG00000168658 | VWA3B | **-1.259359616** | 0.143232064 | 0.060304101 | *0.093408779* |

| | | | | | |
|---|---|---|---|---|---|
| ENSG00000109906 | ZBTB16 | **1.125528581** | 0.280874973 | 0.612102829 | *1.432312526* |
| ENSG00000165424 | ZCCHC24 | **1.407399065** | 0.942927502 | 2.510290276 | *4.257191667* |
| ENSG00000144331 | ZNF385B | **1.267068723** | 0.235571887 | 0.564381003 | *0.302790577* |
| ENSG00000213967 | ZNF726 | **1.12153791** | 0.468872492 | 1.016080479 | *1.623046389* |
| ENSG00000170044 | ZPLD1 | **1.16482452** | 1.67038048 | 3.761410312 | *3.234239273* |
| ENSG00000122952 | ZWINT | **1.086079814** | 12.59456838 | 26.74124584 | *16.97001939* |

**Supplemental Table 4**

***Based on 1479 upregulated DEGs in PLIN2KD***

| Rank | KEGG pathway term | p-value | DEGs included |
|---|---|---|---|
| 1 | **Neuroactive ligand-receptor interaction** | 1.78E-10 | NPFFR2;GABRB1;ADM;GRIK1;GRPR;GRM2;HTR7;CYSLTR2;GRM7;GRM8;LEPR;PRSS3;EDN3;F2R;GABRG3;PRLR;GABRG2;GAL;CRH;PTGER4;CHRNA3;CALCA;CALCRL;CALCB;CHRNA7;LPAR1;LPAR2;ADRB2;CHRND;C5;GLRA3;CALCR;NPW;GALR2;HRH2;CHRNE;NPY;P2RY1;KISS1R;S1PR1;S1PR3;DRD2;GABRD;GABBR2;NMB;HTR1D;HTR1A;HTR1B;OPRK1;GCG;AGT;KISS1;POMC;PYY;P2RX6;LPAR6;SST;NMU;RLN2;RLN1 |
| 2 | Basal cell carcinoma | 2.07E-06 | APC2;CDKN1A;GADD45B;FZD7;LEF1;FZD6;WNT9B;WNT8B;FZD8;FZD10;GLI1;KIF7;GLI3;GLI2;BMP4;BMP2;SMO |
| 3 | Hippo signaling pathway | 5.79E-06 | SERPINE1;LEF1;WNT8B;FZD10;GLI2;SOX2;RASSF2;MYC;TEAD3;APC2;TGFB2;TGFB1;FZD7;FZD6;WNT9B;FZD8;GDF6;BMP6;GDF7;TGFBR2;BMP4;BMP2;FRMD6;LATS2;PPP2R2C;ID1;FAT4;AJUBA;TP73 |
| 4 | **ECM-receptor interaction** | 2.90E-04 | ITGA4;LAMB2;ITGA2;FN1;HSPG2;COL2A1;RELN;COL1A2;TNN;ITGA8;ITGB8;ITGA7;CD36;ITGB6;AGRN;ITGA9 |
| 5 | Complement and coagulation cascades | 6.01E-04 | C1QA;SERPIND1;PROS1;F12;SERPINE1;F2R;SERPINF2;TFPI;F3;F5;C2;F8;C5;C7;MASP1 |
| 6 | **TGF-beta signaling pathway** | 8.53E-04 | TGFB2;TGFB1;FST;INHBB;GDF6;SMAD6;BMP6;TGFBR2;INHBE;GDF7;BMP4;BMP2;BAMBI;MYC;ID1;ID3 |
| 7 | Pathways in cancer | 0.0010704 | ALK;CDKN1A;IL23R;IL5RA;FZD10;GLI1;FGF2;GLI3;GLI2;CASP7;FGF8;HEY1;MYC;HEY2;IL12A;APC2;PDGFRA;IL4R;DCC;ITGA2;MMP2;F2R;WNT9B;FOS;KIF7;TGFBR2;SMO;TRAF5;KIT;NOTCH2;PTGER4;GSTP1;LEF1;LPAR1;ADCY3;WNT8B;LPAR2;RASGRP1;EGFR;STAT4;HMOX1;PMAIP1;HES1;IL12RB2;EGLN3;TGFB2;TGFB1;GADD45B;LAMB2;FZD7;ZBTB16;FZD6;FN1;FZD8;AGT;BMP4;BMP2;LPAR6;BCL2 |
| 8 | Cell adhesion molecules (CAMs) | 0.0022947 | NLGN4Y;CD274;CADM3;NLGN1;SELPLG;NLGN4X;ITGA4;NEGR1;HLA-B;HLA-C;HLA-A;HLA-E;VCAN;SELL;ITGA8;ITGB8;MADCAM1;LRRC4C;JAM2;NECTIN2;ITGA9 |
| 9 | Protein digestion and absorption | 0.0023783 | CELA3B;COL27A1;COL14A1;ATP1B2;SLC1A5;SLC8A1;SLC8A2;SLC8A3;COL2A1;COL1A2;MEP1B;COL5A1;KCNQ1;COL21A1;PRSS3 |
| 10 | Cushing syndrome | 0.0051092 | APC2;CDKN1A;FZD7;FZD6;LEF1;WNT9B;ITPR2;FZD8;ADCY3;WNT8B;FZD10;EGFR;AGT;POMC;NR4A1;PDE11A;RASD1;CREB3L1;CRH;KCNK2;KCNK3 |
| 11 | **PI3K-Akt signaling pathway** | 0.0081811 | CDKN1A;PKN3;LPAR1;LPAR2;FGF2;EFNA4;EGFR;RELN;FGF8;TNN;CREB3L1;MYC;ITGB8;ITGB6;PCK2;PDGFRA;NTRK2;IL4R;ANGPT2;ITGA4;BDNF;LAMB2;ITGA2;F2R;FN1;PRLR;EFNA1;NR4A1;G6PC2;COL1A2;COL2A1;PPP2R2C;LPAR6;DDIT4;KIT;BCL2;ITGA8;ITGA7;ITGA9 |
| 12 | Human papillomavirus infection | 0.0123981 | PTGER4;NOTCH2;CDKN1A;MAML2;WNT8B;FZD10;EGFR;RELN;HEY1;TNN;CREB3L1;HEY2;ITGB8;HES1;ITGB6;HES4;APC2;HES6;ITGA4;LAMB2;FZD7;ITGA2;FZD6;HLA-B;FN1;WNT9B;HLA-C;FZD8;HLA-A;HLA-E;COL1A2;COL2A1;PPP2R2C;ITGA8;ITGA7;ITGA9 |

| | | | |
|---|---|---|---|
| 13 | Hepatocellular carcinoma | 0.023558 | SHC4;APC2;TGFB2;CDKN1A;SHC3;TGFB1;GADD45B;FZD7;GSTP1;FZD6;LEF1;GAB1;WNT9B;FZD8;WNT8B;FZD10;EGFR;TGFBR2;MYC;HMOX1 |
| 14 | Proteoglycans in cancer | 0.0244333 | CDKN1A;TGFB2;TGFB1;FZD7;MMP2;ITGA2;FZD6;FN1;GAB1;WNT9B;FZD8;WNT8B;ITPR2;FZD10;ANK1;HSPG2;FGF2;EGFR;SMO;CTSL;MYC;COL21A1;FLNC |
| 15 | Phospholipase D signaling pathway | 0.0252886 | SHC4;PDGFRA;PLA2G4F;SHC3;SPHK1;F2R;LPAR1;GAB1;LPAR2;ADCY3;EGFR;AGT;GRM2;GRM7;LPAR6;GRM8;KIT;DGKI |
| 16 | Glycine, serine and threonine metabolism | 0.025815 | MAOB;GLDC;SHMT2;CBS;PSAT1;CBSL;PHGDH |
| 17 | Nicotine addiction | 0.025815 | GABRB1;CHRNA7;CACNA1A;SLC17A6;GABRD;GABRG3;GABRG2 |
| 18 | Cholesterol metabolism | 0.0292097 | CYP27A1;ABCA1;ABCG8;LIPC;APOC1;APOA1;CD36;PLTP |
| 19 | **AGE-RAGE signaling pathway in diabetic complications** | 0.0326901 | EGR1;TGFB2;TGFB1;PRKCE;MMP2;SERPINE1;FN1;F3;AGT;TGFBR2;COL1A2;BCL2;PLCD3 |
| 20 | **Calcium signaling pathway** | 0.0382071 | PDGFRA;CAMK1D;CHRNA7;SPHK1;F2R;CACNA1A;ITPR2;ADCY3;ATP2B2;ADRB2;GRPR;SLC8A1;EGFR;SLC8A2;MYLK4;SLC8A3;P2RX6;CYSLTR2;HTR7;HRH2;PLCD3 |

| **Rank** | **GO-molecular function term** | **p-value** | **DEGs included** |
|---|---|---|---|
| 1 | **transforming growth factor beta receptor binding (GO:0005160)** | 4.21E-08 | GDF10;TGFB2;TGFB1;GDF15;INHBB;GDF6;SMAD6;BMP6;INHBE;GDF7;TGFBR2;BMP4;BMP3;BMP2;BAMBI;INHA |
| 2 | **potassium ion leak channel activity (GO:0022841)** | 7.52E-05 | KCNK6;KCNK10;KCNK9;KCNK12;KCNK1;KCNK2;KCNK3 |
| 3 | collagen binding (GO:0005518) | 7.79E-05 | CCBE1;ECM2;PCOLCE2;COL14A1;ITGA2;FN1;NID1;ANTXR1;NID2;MRC2;ADGRG1;CTSL;SERPINH1 |
| 4 | **BMP receptor binding (GO:0070700)** | 1.01E-04 | BMP4;BMP3;TGFB2;BMP2;TGFB1;BMP6 |
| 5 | **hormone activity (GO:0005179)** | 1.02E-04 | CALCB;EDN3;GCG;INHBB;METRNL;PTHLH;AGT;POMC;PYY;GAL;INS-IGF2;VGF;GPNMB;SST;NPY;CRH;INHA |
| 6 | leak channel activity (GO:0022840) | 2.72E-04 | KCNK6;KCNK10;KCNK9;KCNK12;KCNK1;KCNK2;KCNK3 |
| 7 | **G-protein coupled receptor activity (GO:0004930)** | 3.40E-04 | NPFFR2;PTGER4;CALCRL;NPR3;LPAR1;LPAR2;FZD10;GRPR;TSKU;PROKR1;GRM2;ADGRG1;HTR7;CYSLTR2;GALR2;NPY;GRM8;KISS1R;S1PR1;S1PR3;GABBR2;FZD7;FZD6;F2R;HTR1D;HTR1A;FZD8;HTR1B;OPRK1;GPR6;SFRP1;GPR143;ADGRB2;SMO;LPAR6;ADGRL3 |
| 8 | transmembrane receptor protein serine/threonine kinase binding (GO:0070696) | 6.79E-04 | BMP4;BMP3;TGFB2;BMP2;TGFB1;BMP6 |

| | | | |
|---|---|---|---|
| 9 | insulin-like growth factor II binding (GO:0031995) | 8.69E-04 | IGFBP1;IGFBP4;IGFBP3;IGFBP6 |
| 10 | potassium channel activity (GO:0005267) | 0.0010063 | KCNG1;KCNG2;KCNF1;KCNK6;KCNH5;KCNK10;KCNK9;KCNK12;KCNA3;GRIK1;ABCC9;KCNV1;KCNQ1;KCNS3;KCNK1;KCNK2;KCNK3 |
| 11 | kinase inhibitor activity (GO:0019210) | 0.001101 | SOCS2;CDKN1A;SOCS1;LRRTM3;FLRT2;LRRTM1;BGN;SH3BP5;RHOH;RTN4RL2;TRIB1;LRRC4C |
| 12 | metallopeptidase activity (GO:0008237) | 0.0012624 | ENPEP;ERAP2;MMP2;ECEL1;ADAM2;ADAMTS5;ADAM29;MMP14;ADAMTS3;ADAMTS1;ANPEP;ADAM12;CLCA1;ADAMTS8;TRABD2B;ADAMTS7;PAPPA2 |
| 13 | transmembrane receptor protein tyrosine kinase activity (GO:0004714) | 0.0014941 | ALK;EPHB6;EPHA5;NRP1;PDGFRA;NTRK2;EPHA6;NTRK3;KIT;FGFRL1;EGFR;EPHB3 |
| 14 | **bioactive lipid receptor activity (GO:0045125)** | 0.0016367 | LPAR6;SPHK1;S1PR1;LPAR1 |
| 15 | protein kinase inhibitor activity (GO:0004860) | 0.0034149 | SOCS2;CDKN1A;SOCS1;LRRTM3;FLRT2;LRRTM1;BGN;SH3BP5;SPRY2;RTN4RL2;TRIB1;LRRC4C |
| 16 | Wnt-activated receptor activity (GO:0042813) | 0.0043069 | SFRP1;SMO;FZD7;FZD6;FZD8;FZD10 |
| 17 | filamin binding (GO:0031005) | 0.0043542 | DPYSL4;SYNPO2;DPYSL3;RFLNB |
| 18 | neurexin family protein binding (GO:0042043) | 0.0043542 | NLGN4Y;NLGN1;NLGN4X;SYTL2 |
| 19 | MAP kinase kinase kinase activity (GO:0004709) | 0.0045089 | ALK;EPHB6;EPHA5;PDGFRA;NTRK2;EPHA6;NTRK3;MAP3K7CL;EGFR;KIT;MAP3K6;MAP3K5;EPHB3 |
| 20 | transmembrane receptor protein kinase activity (GO:0019199) | 0.0053423 | ALK;EPHB6;EPHA5;PDGFRA;NTRK2;EPHA6;NTRK3;KIT;EGFR;EPHB3;TGFBR2 |

***Based on 493 downregulated DEGs in PLIN2KD***

| Rank | KEGG pathway term | p-value | DEGs included |
|---|---|---|---|
| 1 | **Insulin secretion** | 0.0053337 | GLP1R;CAMK2D;FXYD2;KCNMA1;SLC2A2;ATP1B3;ADCY1 |
| 2 | **Notch signaling pathway** | 0.006384 | NOTCH1;HDAC1;DTX1;DTX3;DLL1 |
| 3 | Legionellosis | 0.0112783 | CASP3;EEF1A2;CXCL1;CXCL3;TLR5 |
| 4 | **cAMP signaling pathway** | 0.0164149 | GLP1R;ABCC4;GRIN2A;CAMK2D;VIPR2;FXYD2;ATP1B3;ADCY1;CNGA4;PLD1;MYL9 |
| 5 | **Maturity onset diabetes of the young** | 0.0254169 | SLC2A2;IAPP;NKX6-1 |

| | | | |
|---|---|---|---|
| 6 | Amphetamine addiction | 0.026119 | CAMK2D;DDC;GRIN2A;TH;HDAC1 |
| 7 | Valine, leucine and isoleucine degradation | 0.0302381 | ALDH6A1;ACAA2;HADH;ACADSB |
| 8 | Cocaine addiction | 0.0323065 | GPSM1;DDC;GRIN2A;TH |
| 9 | Bile secretion | 0.032402 | ABCC4;FXYD2;ATP1B3;ADCY1;SLC4A5 |
| 10 | Thyroid hormone synthesis | 0.0358677 | GPX2;TTR;FXYD2;ATP1B3;ADCY1 |
| 11 | Oxytocin signaling pathway | 0.0360941 | RCAN1;CAMK2D;PRKAA2;CACNA2D3;PRKAG2;ADCY1;MYL9;PIK3R5 |
| 12 | **Circadian rhythm** | 0.0401694 | PRKAA2;PRKAG2;SKP1 |
| 13 | Synaptic vesicle cycle | 0.0434612 | UNC13A;SLC32A1;SLC1A1;STX2;SLC6A4 |
| 14 | cGMP-PKG signaling pathway | 0.0537306 | NPPC;FXYD2;KCNMA1;ATP1B3;ADCY1;MYL9;PIK3R5;SLC25A6 |
| 15 | Protein digestion and absorption | 0.0716232 | SLC15A1;SLC7A9;FXYD2;SLC1A1;ATP1B3 |
| 16 | Nicotine addiction | 0.0752398 | CHRNB2;GRIN2A;SLC32A1 |
| 17 | **Fat digestion and absorption** | 0.0797599 | FABP1;DGAT2;SLC27A4 |
| 18 | Tryptophan metabolism | 0.0843952 | DDC;CYP1A1;HADH |
| 19 | Carbohydrate digestion and absorption | 0.0939997 | FXYD2;SLC2A2;ATP1B3 |
| 20 | **Fatty acid degradation** | 0.0939997 | ACAA2;HADH;ACADSB |

| **Rank** | **GO-molecular function term** | **p-value** | **DEGs included** |
|---|---|---|---|
| 1 | **potassium channel activity (GO:0005267)** | 1.80E-04 | KCNH3;KCNG3;KCND3;KCNQ2;KCNC4;KCNMA1;GRIK3;CNGA4;TMEM175;KCNH1 |
| 2 | **voltage-gated cation channel activity (GO:0022843)** | 8.16E-04 | KCNH3;KCNG3;KCND3;KCNQ2;KCNC4;KCNMA1;CNGA4;CACNA1E;KCNH1 |
| 3 | **voltage-gated potassium channel activity (GO:0005249)** | 0.0011774 | KCNH3;KCNG3;KCND3;KCNC4;KCNMA1;KCNQ2;CNGA4;KCNH1 |
| 4 | magnesium ion transmembrane transporter activity (GO:0015095) | 0.0015699 | TUSC3;NIPA1;SLC41A1 |
| 5 | delayed rectifier potassium channel activity (GO:0005251) | 0.0053136 | KCNG3;KCNC4;KCNQ2;KCNH1 |

| | | | |
|---|---|---|---|
| 6 | protein binding involved in heterotypic cell-cell adhesion (GO:0086080) | 0.01173 | CXADR;DSG2 |
| 7 | phosphatidylinositol transporter activity (GO:0008526) | 0.0153867 | PITPNM3;PITPNM2 |
| 8 | MAP kinase phosphatase activity (GO:0033549) | 0.0194631 | DUSP3;DUSP9 |
| 9 | metal ion transmembrane transporter activity (GO:0046873) | 0.0228932 | TUSC3;NIPA1;SLC41A1 |
| 10 | inositol trisphosphate phosphatase activity (GO:0046030) | 0.0239364 | INPP5A;OCRL |
| 11 | **gap junction channel activity (GO:0005243)** | 0.0287847 | GJD2;GJB1 |
| 12 | wide pore channel activity (GO:0022829) | 0.0339868 | GJD2;GJB1 |
| 13 | hexose transmembrane transporter activity (GO:0015149) | 0.0395224 | SLC2A13;SLC2A2 |
| 14 | outward rectifier potassium channel activity (GO:0015271) | 0.045372 | KCND3;KCNMA1 |
| 15 | **syntaxin-1 binding (GO:0017075)** | 0.0515168 | UNC13A;SNPH |
| 16 | **tubulin binding (GO:0015631)** | 0.0521357 | CLIP2;TPPP3;KIF3B;KIF5A;RITA1;DCX;EML1;NEFH;NDRG1;SAXO2;FTCD |
| 17 | **microtubule binding (GO:0008017)** | 0.0530228 | CLIP2;KIF3B;KIF5A;DCX;EML1;NEFH;NDRG1;SAXO2;FTCD |
| 18 | ionotropic glutamate receptor activity (GO:0004970) | 0.0579388 | GRIN2A;GRIK3 |
| 19 | CXCR chemokine receptor binding (GO:0045236) | 0.0646206 | CXCL1;CXCL3 |
| 20 | **Notch binding (GO:0005112)** | 0.0646206 | DTX1;DLL1 |

**Supplemental Table 5**

***Based on 237 upregulated DEGs in PLIN2OE***

| Rank | KEGG pathway term | p-value | DEGs included |
|---|---|---|---|
| 1 | **Cell cycle** | 2.08E-11 | PLK1;BUB1B;TTK;CDC25C;PKMYT1;CDC20;CCNA2;CCNB2;CDC45;ESPL1;MYC;CDK1;E2F1;E2F2;BUB1 |
| 2 | Oocyte meiosis | 3.14E-08 | CDC20;CCNB2;STAG3;ESPL1;PLK1;CDK1;ITPR3;CDC25C;PKMYT1;FBXO43;BUB1;ADCY5 |
| 3 | Cellular senescence | 2.18E-05 | CCNA2;CCNB2;MYC;CDK1;E2F1;E2F2;ITPR3;MYBL2;FOXM1;TGFBR2 |
| 4 | Progesterone-mediated oocyte maturation | 2.34E-05 | CCNA2;CCNB2;PLK1;CDK1;CDC25C;PKMYT1;BUB1;ADCY5 |
| 5 | Human T-cell leukemia virus 1 infection | 6.38E-05 | CDC20;CCNA2;CCNB2;ESPL1;MYC;IL1R2;E2F1;BUB1B;E2F2;ADCY5;TGFBR2 |
| 6 | Protein digestion and absorption | 9.73E-05 | COL4A2;COL4A1;KCNQ1;COL4A4;COL21A1;SLC1A5;COL9A2 |
| 7 | Bladder cancer | 1.19E-04 | MYC;MMP2;E2F1;E2F2;EGFR |
| 8 | ECM-receptor interaction | 4.29E-04 | COL4A2;COL4A1;COL4A4;SDC1;COL9A2;CD44 |
| 9 | Small cell lung cancer | 8.39E-04 | COL4A2;COL4A1;MYC;COL4A4;E2F1;E2F2 |
| 10 | Relaxin signaling pathway | 9.23E-04 | COL4A2;COL4A1;COL4A4;MMP2;EGFR;TGFBR2;ADCY5 |
| 11 | p53 signaling pathway | 0.001644 | CCNB2;RRM2;CDK1;GTSE1;TP73 |
| 12 | Pathways in cancer | 0.001715 | DCC;PTGER2;ZBTB16;MMP2;EGFR;ADCY5;TGFBR2;COL4A2;COL4A1;MYC;COL4A4;KIT;E2F1;E2F2;BIRC5 |
| 13 | Neuroactive ligand-receptor interaction | 0.002431 | GABBR2;GRM2;GRIN3A;VIPR2;HRH3;CHRNA4;PTGER2;NPY;NMU;GRM8;LEPR |
| 14 | Colorectal cancer | 0.003581 | DCC;MYC;BIRC5;EGFR;TGFBR2 |
| 15 | Gap junction | 0.003952 | TUBB6;CDK1;ITPR3;EGFR;ADCY5 |
| 16 | AGE-RAGE signaling pathway in diabetic complications | 0.006777 | COL4A2;COL4A1;COL4A4;MMP2;TGFBR2 |
| 17 | Central carbon metabolism in cancer | 0.007399 | MYC;KIT;SLC1A5;EGFR |
| 18 | Proteoglycans in cancer | 0.010253 | MYC;MMP2;SDC1;COL21A1;ITPR3;CD44;EGFR |
| 19 | Glutamatergic synapse | 0.011577 | GRM2;GRIN3A;GRM8;ITPR3;ADCY5 |
| 20 | Glioma | 0.012125 | CAMK1D;E2F1;E2F2;EGFR |

| Rank | GO-molecular function term | p-value | DEGs included |
|---|---|---|---|
| 1 | microtubule motor activity (GO:0003777) | 1.05E-10 | CENPE;KIF18A;KIF18B;KIFC1;KIF4A;KIF14;KIF23;KIF2C;KIF11;KIF20A;KIF15 |
| 2 | microtubule binding (GO:0008017) | 1.54E-09 | SPAG5;PLK1;KIF14;KIF23;KIF11;KIF15;CENPE;KIF18A;KIF18B;KIFC1;KIF4A;NUSAP1;BIRC5;KIF2C;KIF20A;FAM83D |
| 3 | motor activity (GO:0003774) | 5.06E-09 | CENPE;KIF18A;KIF18B;KIFC1;KIF4A;KIF14;KIF23;KIF2C;KIF11;KIF20A;KIF15 |
| 4 | tubulin binding (GO:0015631) | 7.01E-08 | SPAG5;PLK1;KIF14;KIF23;KIF11;KIF15;CENPE;KIF18A;KIF18B;KIFC1;KIF4A;NUSAP1;BIRC5;KIF2C;KIF20A;FAM83D |

| | | | |
|---|---|---|---|
| 5 | ATP-dependent microtubule motor activity, plus-end-directed (GO:0008574) | 2.96E-05 | KIF18A;KIF18B;KIF4A;KIF14;KIF11 |
| 6 | ATPase activity (GO:0016887) | 3.20E-05 | CENPE;KIF18A;KIF18B;KIFC1;KIF4A;KIF14;KIF23;KIF2C;KIF20A;KIF11;KIF15 |
| 7 | protein kinase activity (GO:0004672) | 1.41E-04 | CDK18;CAMK1D;PKN3;PLK1;BUB1B;PKMYT1;EGFR;AURKB;CIT;BTC;CCNA2;KIT;PBK;MAP3K20;CDK1;NEK2;BUB1 |
| 8 | ATP-dependent microtubule motor activity (GO:1990939) | 2.53E-04 | KIF18A;KIF18B;KIF4A;KIF14;KIF11 |
| 9 | cyclin-dependent protein kinase activity (GO:0097472) | 6.74E-04 | CCNA2;CCNB2;CDK18;CDK1 |
| 10 | double-stranded DNA binding (GO:0003690) | 0.001163 | RAD51AP1;NEIL3;UHRF1;MCM10;MND1;EGFR |
| 11 | transforming growth factor beta binding (GO:0050431) | 0.001384 | CD109;LTBP2;TGFBR2 |
| 12 | protein serine/threonine kinase activity (GO:0004674) | 0.00154 | CDK18;CCNB2;CAMK1D;PKN3;PLK1;PBK;MAP3K20;CDK1;NEK2;AURKB;CIT;TGFBR2 |
| 13 | protein kinase binding (GO:0019901) | 0.002439 | TOP2A;PLK1;KIF14;KIF11;CDC25C;FOXM1;EGFR;CCNA2;PRC1;KCNQ1;KIT;KIF20A;FAM83D;TP73 |
| 14 | collagen binding (GO:0005518) | 0.003334 | CCBE1;NID1;ANTXR1;CD44 |
| 15 | voltage-gated potassium channel activity involved in ventricular cardiac muscle cell action potential repolarization (GO:1902282) | 0.003735 | KCND3;KCNQ1 |
| 16 | kinesin binding (GO:0019894) | 0.004325 | KIF18B;PRC1;FAM83D |
| 17 | histone kinase activity (GO:0035173) | 0.004765 | CDK1;AURKB |
| 18 | nucleoside-triphosphatase activity (GO:0017111) | 0.004854 | CENPE;KIF18A;KIF18B;KIFC1;KIF4A;KIF14;RAD54L;KIF23;KIF2C;KIF20A;KIF11;KIF15 |
| 19 | filamin binding (GO:0031005) | 0.00591 | DPYSL4;MICALL2 |
| 20 | cyclin-dependent protein serine/threonine kinase activity (GO:0004693) | 0.006897 | CCNB2;CDK18;CDK1 |

***Based on 48 downregulated DEGs in PLIN2OE***

| Rank | KEGG pathway term | p-value | DEGs included |
|---|---|---|---|
| 1 | **IL-17 signaling pathway** | 2.85E-06 | CXCL8;CCL20;CXCL1;CXCL3;TNF |
| 2 | **Cytokine-cytokine receptor interaction** | 6.07E-06 | CXCL8;CCL20;GDF15;CXCL1;CXCL3;TNF;TNFRSF21 |
| 3 | **TNF signaling pathway** | 6.51E-06 | CCL20;CXCL1;CXCL3;TNF;BIRC3 |
| 4 | Legionellosis | 9.10E-06 | CXCL8;CXCL1;CXCL3;TNF |
| 5 | NOD-like receptor signaling pathway | 6.63E-05 | CXCL8;CXCL1;CXCL3;TNF;BIRC3 |
| 6 | Rheumatoid arthritis | 6.70E-05 | CXCL8;CCL20;CXCL1;TNF |
| 7 | Hepatitis B | 6.26E-04 | CDKN1A;EGR2;CXCL8;TNF |
| 8 | Kaposi sarcoma-associated herpesvirus infection | 0.001024 | CDKN1A;CXCL8;CXCL1;CXCL3 |
| 9 | **Chemokine signaling pathway** | 0.001108 | CXCL8;CCL20;CXCL1;CXCL3 |
| 10 | Colorectal cancer | 0.001155 | CDKN1A;TGFA;BBC3 |
| 11 | Salmonella infection | 0.001155 | CXCL8;CXCL1;CXCL3 |
| 12 | **NF-kappa B signaling pathway** | 0.001538 | CXCL8;TNF;BIRC3 |
| 13 | Amoebiasis | 0.001585 | CXCL8;CXCL1;TNF |
| 14 | AGE-RAGE signaling pathway in diabetic complications | 0.001782 | EGR1;CXCL8;TNF |
| 15 | Human T-cell leukemia virus 1 infection | 0.001866 | EGR1;CDKN1A;EGR2;TNF |
| 16 | Bladder cancer | 0.004357 | CDKN1A;CXCL8 |
| 17 | **Apoptosis** | 0.004893 | TNF;BBC3;BIRC3 |
| 18 | Malaria | 0.006173 | CXCL8;TNF |
| 19 | Glutathione metabolism | 0.007998 | GPX2;CHAC1 |
| 20 | Pathways in cancer | 0.008595 | CDKN1A;CXCL8;TGFA;BBC3;BIRC3 |

| Rank | GO-molecular function term | p-value | DEGs included |
|---|---|---|---|
| 1 | **cytokine activity (GO:0005125)** | 1.84E-06 | CXCL8;CCL20;GDF15;CXCL1;CXCL3;TNF |
| 2 | **chemokine activity (GO:0008009)** | 4.42E-06 | CXCL8;CCL20;CXCL1;CXCL3 |
| 3 | **chemokine receptor binding (GO:0042379)** | 5.71E-06 | CXCL8;CCL20;CXCL1;CXCL3 |
| 4 | **CXCR chemokine receptor binding (GO:0045236)** | 8.62E-06 | CXCL8;CXCL1;CXCL3 |
| 5 | manganese ion binding (GO:0030145) | 0.003191 | ME1;SOD2 |

| | | | |
|---|---|---|---|
| 6 | **transforming growth factor beta receptor binding (GO:0005160)** | 0.005004 | GDF15;SMAD7 |
| 7 | transcription regulatory region DNA binding (GO:0044212) | 0.012258 | EGR1;EGR2;TNF;SMAD7 |
| 8 | malate dehydrogenase activity (GO:0016615) | 0.014316 | ME1 |
| 9 | regulatory region DNA binding (GO:0000975) | 0.016561 | EGR2;TNF;SMAD7 |
| 10 | poly(G) binding (GO:0034046) | 0.016682 | ATXN1 |
| 11 | type I transforming growth factor beta receptor binding (GO:0034713) | 0.019043 | SMAD7 |
| 12 | **I-SMAD binding (GO:0070411)** | 0.023748 | SMAD7 |
| 13 | cyclin-dependent protein serine/threonine kinase inhibitor activity (GO:0004861) | 0.023748 | CDKN1A |
| 14 | activin binding (GO:0048185) | 0.023748 | SMAD7 |
| 15 | gap junction channel activity (GO:0005243) | 0.026092 | GJD2 |
| 16 | tumor necrosis factor receptor binding (GO:0005164) | 0.026092 | TNF |
| 17 | G-protein coupled receptor activity (GO:0004930) | 0.026175 | CNR1;FFAR4;GPR37L1 |
| 18 | arylsulfatase activity (GO:0004065) | 0.02843 | SULF2 |
| 19 | wide pore channel activity (GO:0022829) | 0.02843 | GJD2 |
| 20 | ubiquitin protein ligase binding (GO:0031625) | 0.030648 | CDKN1A;EGR2;SMAD7 |

**Supplemental Table 6**

***Most upregulated in PLIN2KD***

| Rank | Gene name | Gene description | Log2(Fold-change) | Fold-change |
|---|---|---|---|---|
| 1 | PPP1R17 | protein phosphatase 1 regulatory subunit 17 | 7.80 | 222.9 |
| 2 | PCDH12 | protocadherin 12 | 6.50 | 90.5 |
| 3 | **FEV** | FEV transcription factor, ETS family member | 5.90 | 59.7 |
| 4 | DNAJC22 | DnaJ heat shock protein family (Hsp40) member C22 | 5.50 | 45.3 |
| 5 | BAALC-AS2 | BAALC antisense RNA 2 | 5.20 | 36.8 |
| 6 | **SST** | somatostatin | 5.10 | 34.3 |
| 7 | SLC18A2 | solute carrier family 18 member A2 | 4.90 | 29.9 |
| 8 | SCUBE3 | signal peptide, CUB domain and EGF like domain containing 3 | 4.90 | 29.9 |
| 9 | CALCA | calcitonin related polypeptide alpha | 4.80 | 27.9 |
| 10 | C7 | complement C7 | 4.70 | 26 |
| 11 | **HHEX** | hematopoietically expressed homeobox | 4.50 | 22.6 |
| 12 | ASCL1 | achaete-scute family bHLH transcription factor 1 | 4.10 | 17.1 |
| 13 | SLC7A5 | solute carrier family 7 member 5 | 3.80 | 13.9 |
| 14 | **ERN1 (IRE1)** | endoplasmic reticulum to nucleus signaling 1 | 3.70 | 13 |
| 15 | SPRY2 | sprouty RTK signaling antagonist 2 | 3.50 | 11.3 |
| 16 | RASD1 | ras related dexamethasone induced 1 | 3.50 | 11.3 |
| 17 | TFPI2 | tissue factor pathway inhibitor 2 | 3.50 | 11.3 |
| 18 | **DDIT4 (CHOP)** | DNA damage inducible transcript 4 | 3.30 | 9.8 |
| 19 | **BMP2** | bone morphogenetic protein 2 | 3.30 | 9.8 |
| 20 | **TGFB2** | transforming growth factor beta 2 | 3.26 | 9.6 |

***Most down regulated in PLIN2KD***

| Rank | Gene name | Gene description | Log2(Fold-change) | Fold-change |
|---|---|---|---|---|
| 1 | GC | GC vitamin D binding protein | -3.2 | -9.2 |
| 2 | NOTCH1 | notch receptor 1 | -2.6 | -6.1 |
| 3 | SNORD113-3 | small nucleolar RNA, C/D box 113-3 | -2.4 | -5.3 |
| 4 | GRASP | trafficking regulator and scaffold protein tamalin | -2.3 | -4.9 |
| 5 | PM20D1 | peptidase M20 domain containing 1 | -2.2 | -4.6 |
| 6 | SLC32A1 | solute carrier family 32 member 1 | -2.2 | -4.6 |
| 7 | TPPP3 | tubulin polymerization promoting protein family member 3 | -2.2 | -4.6 |
| 8 | UNC5D | unc-5 netrin receptor D | -2.1 | -4.3 |
| 9 | MEIS2 | Meis homeobox 2 | -2 | -4.0 |
| 10 | IAPP | islet amyloid polypeptide | -2 | -4.0 |
| 11 | CCDC85C | coiled-coil domain containing 85C | -2 | -4.0 |
| 12 | CNNM1 | cyclin and CBS domain divalent metal cation transport mediator 1 | -1.9 | -3.7 |
| 13 | TSC22D3 | TSC22 domain family member 3 | -1.9 | -3.7 |
| 14 | RCAN1 | regulator of calcineurin 1 | -1.9 | -3.7 |
| 15 | SYT2 | synaptotagmin 2 | -1.9 | -3.7 |
| 16 | ARRB1 | arrestin beta 1 | -1.9 | -3.7 |

| | | | | |
|---|---|---|---|---|
| 17 | EML1 | EMAP like 1 | -1.8 | -3.5 |
| 18 | FGF18 | fibroblast growth factor 18 | -1.8 | -3.5 |
| 19 | SHISA2 | shisa family member 2 | -1.8 | -3.5 |
| 20 | SLC25A43 | solute carrier family 25 member 43 | -1.70 | -3.2 |

**Supplemental Table 7**

***Most upregulated in PLIN2OE***

| Rank | Gene name | Gene description | Log2(Fold-change) | Fold-change |
|---|---|---|---|---|
| 1 | LIPC | lipase C, hepatic type | 1.9 | 3.6 |
| 2 | SPC24 | SPC24 component of NDC80 kinetochore complex | 1.8 | 3.4 |
| 3 | LOXL4 | lysyl oxidase like 4 | 1.6 | 3 |
| 4 | TROAP | trophinin associated protein | 1.6 | 3 |
| 5 | TK1 | thymidine kinase 1 | 1.5 | 2.9 |
| 6 | RIPPLY3 | ripply transcriptional repressor 3 | 1.5 | 2.9 |
| 7 | MAFA | MAF bZIP transcription factor A | 1.5 | 2.9 |
| 8 | AURKB | aurora kinase B | 1.4 | 2.7 |
| 9 | KIT | KIT proto-oncogene, receptor tyrosine kinase | 1.4 | 2.6 |
| 10 | SLC35F1 | solute carrier family 35 member F1 | 1.4 | 2.6 |
| 11 | KCND3 | potassium voltage-gated channel subfamily D member 3 | 1.4 | 2.6 |
| 12 | KCTD15 | potassium channel tetramerization domain containing 15 | 1.4 | 2.6 |
| 13 | KIF18B | kinesin family member 18B | 1.3 | 2.5 |
| 14 | MYBL2 | MYB proto-oncogene like 2 | 1.3 | 2.5 |
| 15 | MMP2 | matrix metallopeptidase 2 | 1.3 | 2.5 |
| 16 | RAD54L | RAD54 like | 1.3 | 2.4 |
| 17 | MDK | midkine | 1.3 | 2.4 |
| 18 | EMID1 | EMI domain containing 1 | 1.3 | 2.4 |
| 19 | SIX2 | SIX homeobox 2 | 1.3 | 2.4 |
| 20 | RIMS4 | regulating synaptic membrane exocytosis 4 | 1.3 | 2.4 |

***Most down regulated in PLIN2OE***

| Rank | Gene name | Gene description | Log2(Fold-change) | Fold-change |
|---|---|---|---|---|
| 1 | STX11 | syntaxin 11 | -2.8 | 0.1 |
| 2 | LYPD1 | LY6/PLAUR domain containing 1 | -1.9 | 0.3 |
| 3 | EEF1A1P26 | eukaryotic translation elongation factor 1 alpha 1 pseudogene 26 | -1.7 | 0.3 |
| 4 | SULF2 | sulfatase 2 | -1.5 | 0.4 |
| 5 | FAXDC2 | fatty acid hydroxylase domain containing 2 | -1.5 | 0.4 |
| 6 | GPX2 | glutathione peroxidase 2 | -1.5 | 0.4 |
| 7 | CEBPB-AS1 | CEBPB Antisense RNA 1 | -1.4 | 0.4 |
| 8 | **SMAD7** | SMAD family member 7 | -1.4 | 0.4 |
| 9 | EGR1 | early growth response 1 | -1.3 | 0.4 |
| 10 | BBC3 | BCL2 binding component 3 | -1.2 | 0.4 |
| 11 | GAP43 | growth associated protein 43 | -1.2 | 0.4 |
| 12 | GDF15 | growth differentiation factor 15 | -1.2 | 0.4 |
| 13 | TNFRSF21 | TNF receptor superfamily member 21 | -1.1 | 0.5 |
| 14 | P3H2 | prolyl 3-hydroxylase 2 | -1.1 | 0.5 |
| 15 | CHAC1 | ChaC glutathione specific gamma-glutamylcyclotransferase 1 | -1.1 | 0.5 |
| 16 | GJD2 | gap junction protein delta 2 | -1.1 | 0.5 |
| 17 | DEPP1 | DEPP1 autophagy regulator | -1.1 | 0.5 |
| 18 | SLC7A11 | solute carrier family 7 member 11 | -1.1 | 0.5 |

| | | | | |
|---|---|---|---|---|
| 19 | ME1 | malic enzyme 1 | -1.1 | 0.5 |
| 20 | CDKN1A | cyclin dependent kinase inhibitor 1A | -1 | 0.5 |
